# A phenotypic screen identified KEAP1-kelch domain blockade as a mechanism to restore cardiac function in the setting of chronic severe hemodynamic stress

**DOI:** 10.1101/2025.09.24.678104

**Authors:** John J. Upson, Christine G. Schnackenberg, Brian G. Lawhorn, Quanhai Chen, Daniel T. Sweet, Thimmaiah P. Chendrimada, Carl A. Brooks, Mark E. Burgert, Christopher T. Louer, Anna E. Zacco, Ian Moench, Jonathan Basilla, Joseph G. Boyer, Christine Y. Ivashchenko, Sharada Manns, Roberta E. Bernard, Gordon C. Pipes, Stephen H. Eisennagel, Weike Bao, Elizabeth Davenport, Joanne C. Kuziw, John A. Krawiec, Kristeen Hauk, Sarah Agrapides, Mary Rambo, Alan R. Olzinski, Katrina Rivera, Eugene T. Grygielko, Larry J Jolivette, John J. Lepore, Robert N. Willette, John R. Toomey

## Abstract

A bespoke gene expression (GE) based small molecule screen in human induced pluripotent stem cell derived cardiomyocytes (iPSC-CM) was conducted with the aim of identifying drug targets and pathways with the potential to reverse heart failure (HF) pathologic GE and the resultant decompensated HF phenotype. The screen utilized a composite human-murine HF gene expression signature, a target annotated compound set, and a HF gene expression reversal scoring algorithm to identify small molecules and their associated targets as potential modulators of HF pathologic GE. Following hit triage, a lead optimization program, and compound characterization in preclinical rodent models of HF and human iPSC-CM contractility assays, KEAP1 kelch domain blockers were identified as potent efficacious agents in the restoration of contractile function in the setting of oxidant stress and pressure overload induced cardiac dysfunction.

## Introduction

Heart failure (HF) has a complex etiology driven by a spectrum of cardiac stressors of which the most common are hypertension, ischemic injury, vascular disease, or genetic predispositions to cardiac dysfunction. The progression of HF is characterized by structural and molecular remodeling of the heart, which may include ventricular hypertrophy and/or dilation, deranged myocardial metabolism, inflammation, fibrosis, and cell death (*1*). The net effect of this maladaptive remodeling is to reduce cardiac performance which inexorably progresses to pump failure (*2*). The multifactorial origins of HF and the extensive pathologic remodeling of the organ have presented a significant challenge for the discovery of medicines to reverse the progression of the disease. To date, pharmacologic strategies aimed at reducing cardiac afterload, work, and metabolic stress have produced the most success in reversing cardiac remodeling and attenuating the rate of decline. Studies of chronic therapy with renin-angiotensin-aldosterone system inhibitors, β-blockers, and more recently SGLT2 inhibitors, have all reported decreased left ventricular (LV) dilation and end systolic dimensions and have yielded significant improvements in contractile function (*3-6*). From a therapeutic perspective, the combined data have demonstrated that pathologic cardiac remodeling has elements of reversibility.

The molecular reprogramming required to reverse the pathology of a decompensated myocardium has not been elucidated. Pathways and mechanisms related to cardiomyocyte hypertrophy, calcium handling, fatty acid oxidation, apoptosis, inflammation, oxidant stress, and many others have been implicated in reverse remodeling (*7*). Under the assumption that disease associated GE is predominantly maladaptive and a manifestation of non-adapted plasticity (*8*), multiple drug discovery campaigns have ensued where pharmacologic reversal of disease-related GE has been attempted. Utilizing libraries of human cell based drug-induced GE signatures and pattern matching algorithm’s, compounds have been identified to modulate disease GE signatures and disease models for a broad range of disorders, including irritable bowel disease (*9*), muscle atrophy (*10*), cancer (*11*), and diabetes (*12*). In the following report, a bespoke GE based small molecule screen in human induced pluripotent stem cell derived cardiomyocytes (iPSC-CM) was conducted to identify potential cardioprotective mechanisms.

The screen utilized a composite human-murine HF gene expression signature, a target annotated compound set, and a HF gene expression reversal scoring algorithm to identify small molecules and their associated targets as potential modulators of HF pathologic gene expression. Following hit triage, and compound characterization in preclinical rodent models of HF and human iPSC-CM contractility assays, Kelch-like ECH associated protein-1 (KEAP1) kelch domain blockers were identified as potent efficacious agents in the restoration and normalization of cardiac contractile function in the setting of HF with reduced ejection fraction.

## Materials and Methods

### Study design

The aim of the GE based phenotypic screen was to identify drug targets and pathways with the potential to reverse heart failure (HF) pathologic GE and the resultant decompensated HF phenotype. The critical path for screening hit ID followed the steps detailed in the methods of initially scoring hits by algorithm, and then through triaging hits by concentration response, cytotoxicity, SAR/TAR, and an in vivo murine TAC model. In TAC studies, the primary endpoint was ejection fraction (EF) and studies were designed to provide 80% power to detect a 7% EF difference in vehicle versus drug. Statistical analysis, blinding and randomization, and stopping criteria details were as described in the methods. No animals were excluded from analysis unless withdrawn from the study due to severe negative clinical signs. The animal numbers per cohort were as detailed in the figure legends.

### Materials

SKF-92837, GSK3161593A, GSK3175696A, GSK2366910A (bardoxolone methyl), GSK3776143B, GSK3407931B, GSK3525714B (figure S1) were synthesized at GlaxoSmithKline under a collaboration with Astex Pharmaceuticals (Cambridge, UK). The following supplies were purchased from the companies as cited. Thyroid hormone (3, 3’, 5-Triiodo-L-thyronine) from Alfa Aesar (Ward Hill, MA), human induced pluripotent stem cell derived cardiomyocytes (iPSC-CMs), plating media, and maintenance media from Cellular Dynamics International (Madison, WI), gelatin solution from Millipore (Billerica, MA), endothelin-1 from American Peptide Company (Sunnyvale, CA), RLT lysis buffer and RNeasy Plus Mini kits from Qiagen (Valencia, CA), Nanostring custom Plex^2^ code sets from Nanostring Technologies (Seattle, WA), ToxiLight adenylate kinase cytotoxicity kit from Lonza (Walkersville, MD), isoflurane from <u>HANNA Pharmaceutical Supply Co</u>. (Wilmington, DE), Oxidized Protein Western Detection kit and NAD(P)H quinone oxidoreductase 1 (NQO1) activity kits from Abcam (Cambridge, MA), MYBPC3 (M-190) primary antibody from Santa Cruz Biotechnology (Dallas, TX), goat anti-DNP primary antibody from Bethyl Laboratories (Montgomery, TX), IRDye 800CW donkey anti-goat IgG (H+L), IRDye 680LT donkey (polyclonal) anti-rabbit IgG (H+L), and Odyssey blocking buffer from LI-COR (Lincoln, NE), plasma atrial natriuretic peptide (pro-ANP 1-98) immunoassay from Biomedica Immunoassays (Vienna, AT), Masson’s trichrome stain from Poly Scientific R&D Corporation (Bay Shore, NY), Cavitron from Wacker Chemie AG (Burghausen, DE).

### Animal Protocols

All animal procedures were approved by the Institutional Animal Care and Use Committee of GlaxoSmithKline.

### Construction of a human-mouse HF gene expression signature

Affymetrix HU133A microarray data from HF patient myocardium with advanced systolic HF (n=194) or non-failing controls (n=16) were obtained using Gene Expression Omnibus (http://www.ncbi.nlm.nih.gov/geo; accession no. GSE5406). Patients were reported to have had HF due to either ischemic (n=86) or idiopathic dilated (n=108) cardiomyopathy. No subjects received mechanical support with LVADs, and all HF patients had NYHA class 3 to 4 symptoms and LV systolic dysfunction(*13*). All files were imported into Array Studio (version 6.2) and normalized by robust multiarray analysis (RMA). Samples with a median absolute deviation (MAD) score < -5 were identified as outliers and omitted from subsequent analysis. Fold changes, relative to non-failing controls, and p-values were determined using one-way ANOVA with False Discovery Rate-Benjamini Hochberg (FDR-BH) multiplicity adjustment for both the ischemic and idiopathic dilated cardiomyopathy samples, respectively. Probes with FDR-BH p-values < 0.05 were considered significant. The final GE profiles of both ischemic and idiopathic dilated cardiomyopathy were generated by calculating the mean fold change and FDR-BH p-value for any genes represented by multiple probes in the Affymetrix HU133A array. The overlap of the GE signatures of ischemic and idiopathic dilated cardiomyopathy were determined using the Venn Diagram function in Array Studio. Gene changes identified in the overlap were further assessed for directionality agreement and filtered accordingly. The final human HF gene expression profile was comprised of genes that were significantly altered, in the same direction, in both ischemic and idiopathic dilated cardiomyopathy.

Affymetrix microarray data from mouse left ventricle from a 6-week transverse aortic constriction (TAC) model (n=5) or Sham (n=5) mice were collected (see TAC methods for model details). TAC mice demonstrated left ventricular systolic dysfunction, with mean ± SD ejection fraction of 36 ± 5%. Sham mice had normal left ventricular function with mean ejection fraction of 56 ± 4%. All files were imported into Array Studio (version 6.3) and normalized by robust multiarray analysis (RMA). Samples with a median absolute deviation (MAD) score < -5 were identified as outliers and omitted from subsequent analysis. Fold changes relative to sham mice and p-values were determined using one-way ANOVA with false discovery rate Benjamini-Hochberg (FDR-BH) multiplicity adjustment for TAC mice. Probes with FDR-BH p-values < 0.05 were considered significant. The final GE profile of mouse TAC was generated by calculating the mean fold change and FDR-BH p-value for any genes represented by multiple probes in the Affymetrix Mouse 430_2 array.

The overlap of human HF and mouse TAC gene expression was determined using the Venn diagram function in Array Studio. Gene changes identified in the overlap were further assessed for directionality agreement and filtered accordingly. The overlapping signature was further refined by selecting those genes where transcript was significantly detected in human iPSC-CMs under basal conditions and by suitability for inclusion in a Nanostring code set, satisfying the criteria of uniqueness in the genome and mutual independence from the other probes in the design set. Finally, four genes that have been closely associated with maladaptive cardiac remodeling, PPARGC1(*14*), SERCA2A(*15*), NPPB(*16*), and Myh6(*17*), that did not survive the unbiased selection process for top transcripts were added to the signature. The combined filters and additions resulted in an 85-gene signature (Table 1).

**Table 1.**
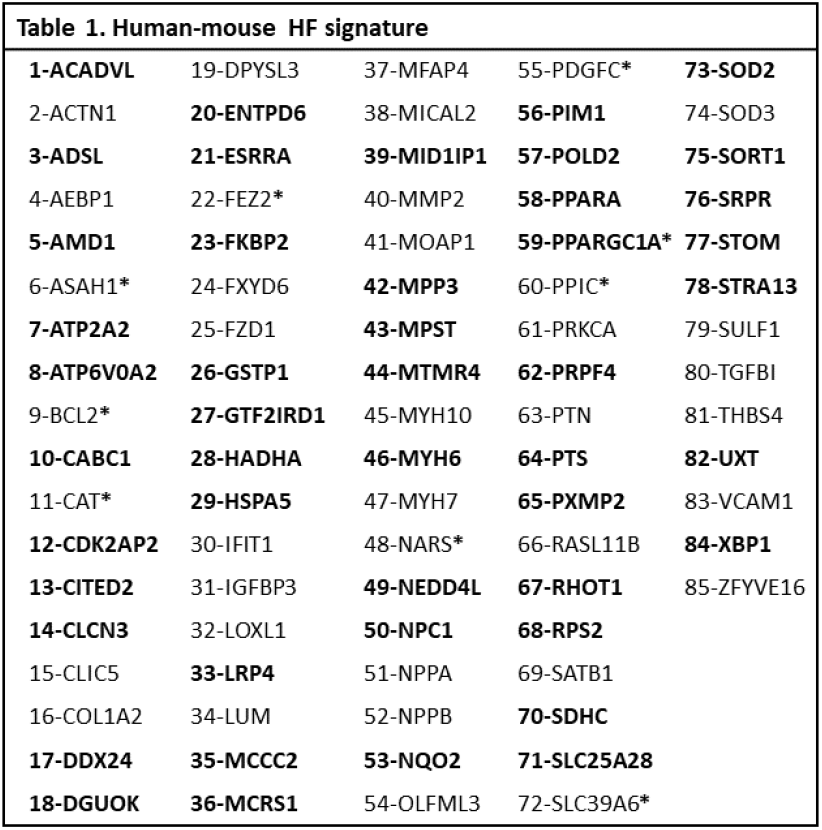
The human-mouse composite HF gene expression signature. The signature was constructed as described in methods. At baseline, genes in bold were decreased in HF and non-bolded genes were increased in HF compared to controls. All genes (except with asterisks) were assigned a positive score by the algorithm when the direction of baseline gene expression was reversed by compounds in human iPSC cardiomyocytes. Genes with asterisks were scored positively when the direction of baseline regulation was augmented by compounds. (See scoring algorithm in methods for further details)

### Annotated compound set

An annotated set of 1050 compounds was assembled from the GlaxoSmithKline compound library and consisted of molecules that were target-annotated, biologically diverse, primary pXC50 > 6.0, 100x selective vs secondary measured activities, high artificial membrane permeable (surrogate for cell penetrance), and structurally diverse. The set of 1050 compounds contained ∼800 unique target annotations and activities.

### Gene expression screen in iPSC-CMs of an annotated compound set

Human iPSC-CMs were plated at 2.5 x 10^4^ cells/well of a 96-well plate and cultured per the manufacturers protocol for 15 days in n=2 replicates. On day-13 cells were switched to serum-free (SF) media [3:1 DMEM (with high glucose):M199 (with Hank’s salt) + 2 mM L-Glutamine]. On day 14 cells were treated with either 0.1% dimethyl sulfoxide (DMSO) vehicle or 10uM of each compound from the annotated compound set in SF media. On day 15 the media was removed, the cells were washed with phosphate buffered saline, and then lysed with RLT buffer, and frozen at - 80°C. Gene expression analysis from frozen lysates was performed using Nanostring nCounter Plex^2^ technology per the manufacturers protocol. The housekeeper genes B2M, HPRT1, OAZ1, and RPLP0 were verified as categorically unaffected by all compounds in the screen and used to normalized GE data by Analysis of Covariance (Array Studio, v7.1). Fold changes relative to vehicle treated cells were calculated by the scoring algorithm as described below.

### Scoring algorithm

The compound scoring algorithm was configured to rank compounds based upon their favorable or unfavorable aggregate effects on the GE signature. For 90% of signature genes, favorable directionality of GE was considered a reversal of native HF GE; whereas, for ∼10% of genes favorable expression was adjudicated as in the same direction as identified in HF. The adjudication process for all genes in the signature was a scientific literature exercise, interrogating the full context of reported functions and roles as pertains to HF. The algorithm used to score and rank order compounds functioned as follows. For each well in the screen, the GE log_2_ fold change relative to vehicle was determined for each gene in the 85-gene signature set and then subsequently transformed into favorable or unfavorable directionality. Genes regulated in a favorable or unfavorable direction were assigned positive or negative values respectively. Weighted gene values were calculated by dividing the individual directed log2 fold change values by the gene specific standard deviation of their respective plate vehicle wells. Directed and weighted log_2_ fold changes were capped based on the absolute values of the directed and weighted changes and the capped values were assigned as follows: (1) absolute values < 2 were assigned as 0; (2) absolute values > 2 were recorded as that specific value but were capped at a maximum value of 4 which limited the effect of large GE changes from a single gene on the final score. The final score was determined by calculating the mean of the capped, directed, and weighted log_2_ fold GE changes across the 85-gene set. A hit score was defined as any score above vehicle-well score variability.

### Cytotoxicity and concentration-response determination

For each compound meeting hit criteria, cytotoxicity and six-point concentration response GE scores were generated. The human iPSC-CMs were cultured as described previously. On day 14, iPSC-CMs were treated with either vehicle (0.1 % DMSO), or a six-point concentration response (0.041, 0.123, 0.370, 1.1, 3.3, & 10 uM) of each compound. On Day 15, media was collected for cytotoxicity determination, and the cells were washed with phosphate buffered saline and then lysed with RLT buffer and frozen at -80°C. Compound induced cytotoxicity was determined by measuring adenylate kinase (AK) released into the media. Cytotoxicity was defined as > 2-fold increase in AK. Gene expression analysis and scoring (per concentration) was performed as described previously.

### Structure activity relationship analysis

For hit compounds demonstrating a concentration-response and a non-cytotoxic profile, a target activity relationship (TAR) and structure activity relationship (SAR) investigation involving chemical or annotated target analogs of hits was conducted. A TAR/SAR investigation would ascertain whether target annotation could predict a positive algorithm score and whether the chemical and structural elements of compounds conferring activity were consistent. Hits demonstrating both TAR and SAR with the algorithm score were deemed progressible, in that a path for chemical optimization had been identified as well as a discreet biological target accountable for the positive algorithm score.

### Pharmacokinetics (PK)/pharmacodynamics (PD)

Adult male C57BL/6 mice were individually housed and acclimated to a standard powdered rodent diet (Lab Diet 5001 meal) for 1-3 days. Water was offered *ad libitum* throughout the study. Compounds were administered over a dose-range for 7 days. Food consumption and body weight were monitored from baseline and on days 1, 5, 6, and 7. Plasma was collected in the morning of Days 1, 4 and 7 for subsequent PK analysis. Analyses of mouse plasma samples for compound concentrations were performed using liquid chromatography/tandem mass spectroscopy. Compound plasma protein binding was determined by the ultracentrifugation method followed by liquid chromatography and tandem mass spectroscopy(*18*).

To measure pharmacodynamic activity (gene expression), hearts were excised on day 7 and the apex of the left ventricle (LV) was removed and sectioned, and the posterior apical quadrant of the LV from each of the respective treatment groups was homogenized in 400uL of TRIzol, and RNA extracted using the RNeasy Plus Mini kit protocol. RNA was quantified using a Nanodrop Spectrophotometer ND-1000 and frozen at -80°C. Gene expression analysis on the isolated RNA samples was performed per the Nanostring nCounter Plex^2^ manufacturers protocol.

### Hemodynamic assessments

Invasive hemodynamic measures were performed on day 7 in PK/PD studies or at the end-of-study for the TAC model. Mice were anesthetized with 2% isoflurane, and then maintained at 1.5%, delivered in 100% oxygen, on a heated surgical table to maintain body temperature within the physiological range (36.6°C - 37.4°C). A 1F Millar catheter (SPR1000) was placed in the right carotid artery. The left jugular vein was isolated and prepared with 4-0 silk suture for later cannulation. After stabilization, the systemic arterial blood pressure was recorded for a minimum of 20 seconds, and then the catheter was advanced into the left ventricle for ventricular pressure measurements. Data were recorded and analyzed using LabChart-7 (AD Instruments).

### Dobutamine challenge

Following the acquisition of baseline hemodynamic data, the jugular vein was cut, and the infusion catheter containing dobutamine (1ug/g body weight) was inserted into the vein and secured with pre-placed 4-0 sutures. Dobutamine was delivered via an infusion pump at the rate of 1uL/g/min for 2 min while LV pressure was recorded continuously. Cardiac reserve was calculated as the difference between dP/dt_max_ measured at baseline and during dobutamine infusion (delta dP/dt_max_).

### Mouse transverse aortic constriction (TAC) left ventricular pressure overload model

Male C57BLK/6J mice (10-12 weeks old) were individually housed and acclimated to a standard powdered rodent diet (Lab Diet 5001 meal) for 1-3 days. Water was offered ad libitum throughout the study. Mice were initially anesthetized in a chamber using an 3% isoflurane with oxygen (1.0 L/min) and then maintained at 1.5% via nose cone (without intubation). A small incision (∼5 mm) was made just left of midline and just above the rib cage at the level of the suprasternal notch. The muscle tissue was retracted to expose the area above the pleural cavity. The lobes of the thymus were separated and retracted to expose the aortic arch. Fine surgical forceps with blunted tips were used to expose a region above and below the aorta. A micro blunt hook tied with 7-0 silk surgical suture was looped under the aorta and the suture pulled through. The suture was tied off against a small piece of blunted 27 G needle and the needle was then removed creating the aortic constriction (a 60-70% constriction of the lumen). The incision was closed using 6-0 silk suture for the muscle layer, then the skin with surgical tissue adhesive. The sham procedure was identical without ligation of aorta. Depending on the individual study designs, drug administration began either on the day of surgery, or 1-2 weeks after surgery (detailed in figure legends) and continued until the end of study. Compounds were administered in the mouse chow unless otherwise noted in figure legends. TAC study duration was either 6 weeks or 10 weeks in duration. All TAC study endpoints were collected at the end of study. Following heart dissections, samples were either snap frozen in liquid nitrogen and stored at -80°C or placed in 10% neutral buffered formalin until analysis.

### Statistical analysis

All data are presented as mean ± sem. For *in vivo* studies, no animals were excluded from analyses unless removed from the study due to severe negative clinical signs or sustained body weight reduction of > 25%. Statistical tests for significance are described in the figure legends, along with replicates (n). P values are captured within the figures themselves (above bars). The data analysis software system used was STATISTICA, StatSoft, Inc. (2013), version 12. www.statsoft.com.

### Blinding and randomization

A multilayered blinding and randomization method was used for all (other than screening) *in vivo* work. Blinding and randomization layers were as follows: (1) Animals were numbered with a study ID and then renumbered with a randomized blinding number; (2) Animals were randomized across days, time of day, and treatment groups for surgery; (3) Surgeons were blinded to all animal dose-group assignment information; (4) Surgeons and the scientists collecting and analyzing all cardiac hemodynamic or imaging endpoints were blinded to the animal ID and excluded from drug administration; (5) All study endpoints *in vitro* and *in vivo* were blinded; (6) A copy of all blinded endpoint data was secured by a study coordinator (transparent raw data capture) prior to release of the blinding key; (7) The blinding key was password protected; (8) No data were unblinded prior to unblinding the primary endpoint.

### Magnetic Resonance Imaging (MRI)

For the initial compound mouse TAC profiling study (table S3, figure 3), cardiac function was determined by MRI at the end-of-study per previously published methods(*19*).

### Echocardiography

For subsequent mouse TAC studies (figures 4 and 5 and S2-S3), echocardiography was performed on a Vevo 2100 high frequency imaging system. Animals were anesthetized with 2-3% isoflurane and sedation maintained with 1% isoflurane with animals on a heating pad. B-mode imaging of three cardiac cycles was used to assess ejection fraction (EF), end diastolic volume (EDV), end systolic volume (ESV), LV mass, and heart rate. ECG was used to measure heart rate during the echocardiography procedure(*20*).

### Cardiac fibrosis

From the mouse TAC studies, the posterior half of the LV was fixed in 10 % neutral buffered formalin, paraffin embedded, cut in 5 um sections, placed on glass slides, deparaffinized, and stained with Masson’s trichrome. Stained slides were digitally scanned and archived using a Leica (Buffalo Grove, IL) image capture device. The images were analyzed for fibrosis using ImageJ 1.49v software (nih.imageJ.gov). The fibrosis percentage was calculated as the aniline stain area divided by total image area. Aorta, aortic valves, and papillary muscles were excluded from the analysis.

### Cardiac myosin binding protein C (cMYBPC) carbonylation

From the mouse TAC studies, the right ventricle was used to determine the extent of cMYBPC carbonylation(*21*). The right ventricle was snap frozen in liquid nitrogen and pulverized. The oxidized protein detection kit (see materials) was used to lyse and derivatize the tissue according to the manufacturer’s recommendations. Derivatized total protein (∼30ug) was run on a 4-12% tris-glycine mini-gel at 150V for 2-hours on ice. Proteins were transferred from the gel to a nitrocellulose membrane, which was subsequently blocked using Odyssey blocking buffer and incubated with anti-mouse cMYBPC (M-190) rabbit polyclonal IgG at a 1:200 dilution, and goat anti-DNP at a 1:500 dilution, at 4°C overnight. Membranes were washed and then incubated with donkey polyclonal anti-rabbit IgG at a 1:20,000 dilution and 800CW donkey anti-goat IgG at a 1: 15,000 dilution at room temperature for 30 min. Membranes were washed and then visualized using an Odyssey Infrared Imaging System (Li-Cor Biosciences).

### NADPH dehydrogenase [quinone]-1 (NQO1) activity

Left ventricular (LV) tissue NQO1 activity was determined using an NQO1 assay kit per manufacturer’s protocol. Pulverized LV was adjusted to a protein concentration of 100ug/ml for the enzyme activity assessment. Fold change values were calculated by normalizing to the mean of the vehicle treated group and values reported as fold change versus the vehicle group.

### iPSC-CM contractility

Human iPSC-CM were seeded onto gelatin coated ePlate CardioECR48 (ACEA Biosciences, San Diego CA) and matured for two weeks according to manufacturer instructions. Cardiomyocyte contractile strength was measured on the xCELLigence CardioECR device which measures changes in electrical impedance of the iPSC-CM monolayer as a function of cellular shape change during contraction. Impedance shape change measurements are directly related to contractile strength. Human iPSC-CMs were paced at 60bpm to normalize variation in beating frequency. Cells were pre-treated for 48 hours with KEAP1 blockers, after which media was changed and fresh compound was added to cells. Tert-butyl hydroperoxide (tBHP) was added 4 hours later to each well and contractile amplitude was measured continuously for 48 hours. The change in contractile amplitude compared to pre-oxidation (before addition of tBHP) over the time course of the experiment was calculated for each well.

## Results

### Phenotypic screening output

Based upon published human HF gene expression data (*13*) and gene expression data acquired from a murine 6-week TAC model (Table S1), a human-mouse composite HF gene expression signature was constructed (Table 1). The selection of signature genes was predominantly unbiased and was based upon the rank order of GE significance in the human and murine HF microarray datasets, the observed gene overlap, and common directionality of GE.

No general attempt to engineer the gene list by perceived mechanistic diversity or - pathologic relevance was made. As described in methods, the compound scoring algorithm was configured to rank order compounds based upon their favorable or unfavorable aggregate effects on the gene expression signature. For 90% of signature genes and in line with the method precedent (*22*), favorable directionality of gene expression was chosen as a reversal of native HF gene expression. For 10% of signature genes, and per a scientific literature adjudication, favorable directionality of GE was selected as in the same direction as that identified in HF.

A select set of 1050 target annotated compounds were screened in duplicate in human iPSC-CMs for their effects on modulating the composite HF signature as specified in the methods. A hit score was defined as any score above the vehicle score variability, which equated to 2.7 standard deviations from the vehicle mean score. The observed hit rate was 9.5% with 100 compounds qualifying as hits (figure 1, table S2). Out of the 100 hits, 63 retested as positive scores at 10uM. After additional interrogation of concentration-responses, cytotoxicity, and testing related analogs, only 2 targets demonstrated both a target activity relationship (TAR) and a structure activity relationship (SAR). The two targets and the compound mechanisms of action were annotated as a thyroid receptor-β (TRβ) agonist and inhibitors of the Kelch-like ECH associated protein-1 (KEAP1). The two KEAP1 inhibitors encompassed different mechanisms of action, with one a known KEAP1 BTB domain electrophile(*23*), and the other a KEAP1 Kelch domain binder(*24*). For higher scoring compounds where SAR did not align with target annotation, multiple attempts to identify the molecular targets by chemical proteomics (*25*) were unsuccessful. Negative scores were found to be significantly driven by cytotoxicity.

**Figure 1.**
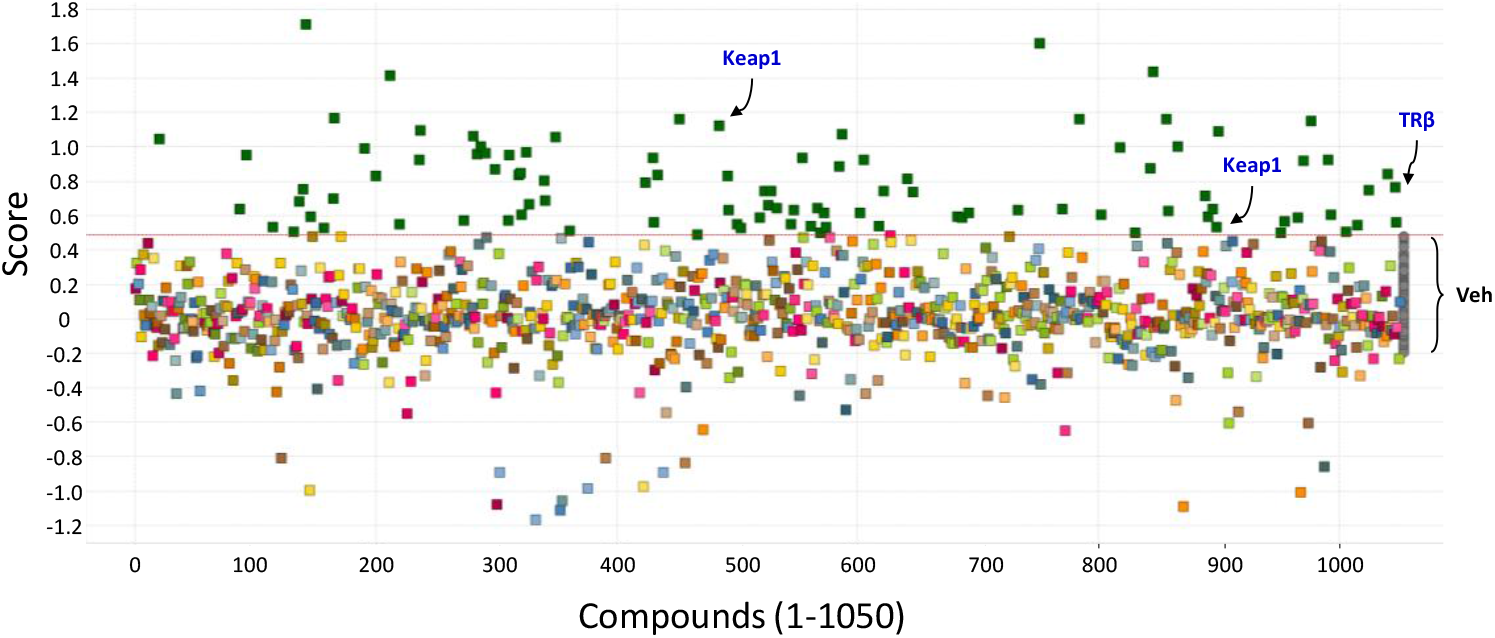
Screening output from the annotated compound set. Dark green symbols, scored at or above vehicle score variability, were selected as hits. The red dotted line represented the hit cut-off GE score, based on vehicle score variability. Vehicle scores represented by the far-right gray symbols.

Gene expression changes and representative visualizations at 10uM for a related KEAP1-Kelch domain blocker and TRβ compounds are shown in figure 2A-B. A concentration-response for the KEAP1 kelch domain blocker is shown in figure 2C. Aside from the aggregate HF gene expression scores, no attempt to weight or triage compounds by a specific subset of gene expression changes were made.

**Figure 2.**
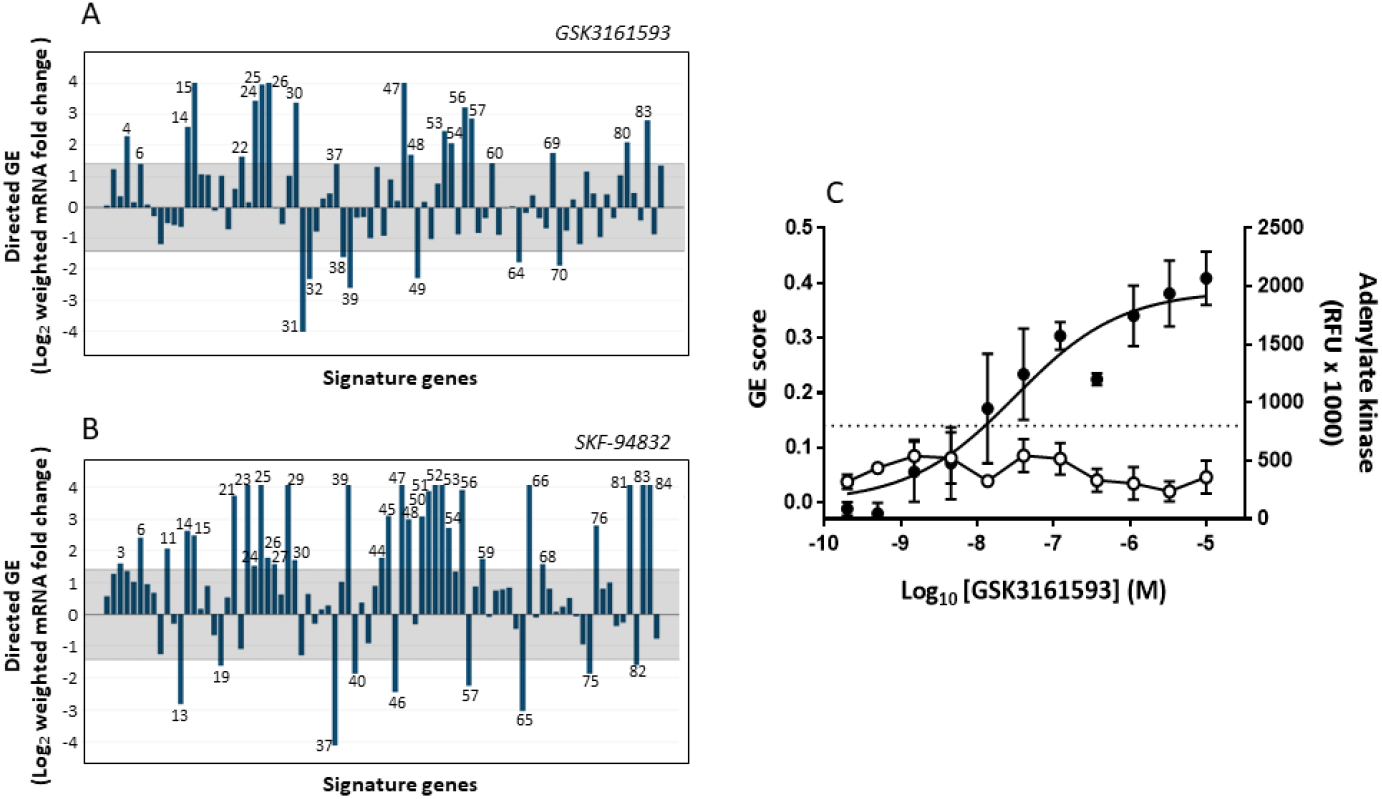
A-B) Visualization of directed GE effects of the KEAP1 kelch domain blocker GSK3161593 (10uM) and TRβ agonist SKF-94832 (10uM) across the HF gene expression signature. Genes contributing to the aggregate score are numbered (see table 1 for gene numbers). Per the algorithm, compound elicited GE changes in a favorable direction were assigned a positive value (up) and a negative value for an unfavorable direction (down). The gray box represents the vehicle-well score variability (see methods for select compound algorithm adjustment) and the minimum log_2_ fold change for inclusion in the score. Note that only 83 of the 85 HF signature genes are shown in figures A-B. The signal for genes BCL2 and LUM fell below the detection limit. **C)** Concentration-dependent GE score and cytotoxicity of KEAP1-kelch domain blocker (GSK3161593) in iPSC-cardiomyocytes. GE scores represented by filled symbols, cytotoxicity by open symbols. The dotted line represents both the hit cut-off GE score (based on vehicle score variability for the specific study) and the cytotoxicity cut-off (based on 2X baseline AK in the cell culture media).

### Compound profiling in murine heart failure (TAC) model

Following the identification of KEAP1 and TRβ as targets underpinning positive GE scores, the hits and/or analogs were profiled in healthy animals in preparation for an *in vivo* efficacy assessment in a murine transverse aortic constriction (TAC) model. Profiling included determination of pharmacokinetics, extent of plasma protein binding, 7-day tolerability assessments with measurements of compound effects on food consumption, body weights, blood pressure, heart rate, and compound effects on a left ventricular GE score in naïve mice. Dose-ranges for the initial mouse TAC efficacy studies were based upon maximum tolerated and neutral hemodynamic doses, while aiming to achieve an unbound (to protein) drug concentration *in vivo* to match or exceed the compound *in vitro* EC_50_ potency determined in the GE iPSC-cardiomyocyte assay, and a positive GE score in left ventricular tissue.

The mouse TAC model was selected as the primary HF screening model based upon its central role in the construction of the human-mouse composite HF gene expression signature. After surveying several tool molecules through the above compound profiling, where most showed poor tolerability and/or inadequate pharmacokinetics, five molecules were progressed for initial TAC studies: TRβ agonist SKF92837, native thyroid hormone (triiodothyronine or T3), KEAP1-kelch blocker GSK3175696A, KEAP1-kelch blocker GSK3161593A, and KEAP1-BTB inhibitor Bardoxolone methyl (GSK2366910A). All five compounds elicited positive efficacy trends toward improving cardiac function (Table S3, SKF92837 and GSK3161593A highlighted in figure 3), whether administered at the time of TAC or one week afterwards (1-week post-TAC for KEAP1 modulators). No significant changes in cardiac hypertrophy or left ventricular end diastolic pressures were observed in these screening studies, and as expected elevated heart rates were observed with TR agonism (SKF92837 30x selective for TRβ vs TRα). Based upon these initial positive functional trends and the chronotropic challenges of TRβ selective agonism (*26*), and the selectivity challenges of KEAP1 electrophiles(*24*), subsequent efforts were directed toward the development of improved KEAP1-kelch domain blocker tools and further interrogation of the KEAP1 mechanism.

**Figure 3.**
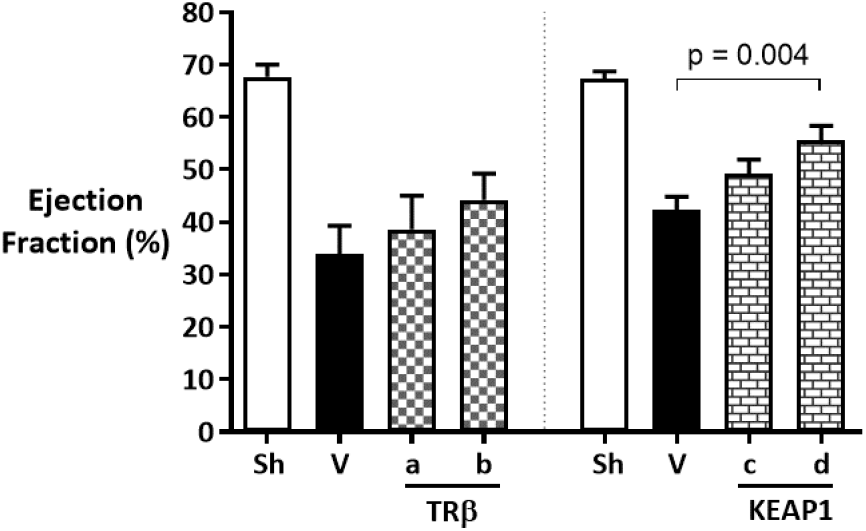
Effects of 2 screening hits, thyroid receptor-β agonist SKF-92837 and KEAP1 blocker GSK3161593A, on ejection fraction in a murine 6-week TAC model. The results are from two independent studies. Figure labeled as follows: For the TRβ study, Sh (sham, n=5), V (TAC, no drug, n=10) a & b (TAC and TRβ agonist SKF-92837, 0.14 & 0.42 mg/kg/day, n=7 & 9 respectively. Drug administration was initiated the day of TAC surgery. For the KEAP1 study, Sh (sham, n=6), V (TAC and Cavitron/DMSO, n=16), c & d (TAC and KEAP1 blocker GSK3161593A 50mg/kg/d, subcutaneous (n=14 each) QD & BID respectively). Drug administration was initiated at 1-week after TAC surgery. KEAP1 study vehicle was 20% Cavitron (hydroxypropyl-β-cyclodextrin) (w/v) with 2% dimethyl sulfoxide, pH 8.5. Ejection fraction was measured by MRI. Data are represented as mean (± SEM). P-values were determined by one-way ANOVA with Dunnett’s multiple comparison test.

A chemical optimization effort focused on the KEAP1-kelch domain blockers was initiated with the aim of developing *in vivo* tool molecules with improved pharmacokinetics (PK) and tolerability (*24, 27*). An early orally bioavailable tool molecule developed in the campaign, GSK3407931B (figure S1) was evaluated in a 10-week TAC model at a maximum tolerated dose. This KEAP1-kelch domain blocker predictably induced NRF2 mediated NQO1 upregulation in the LV and consistent with the screening TAC findings, did not affect LV hypertrophy (figure 4). The compound elicited a full normalization of cardiac contractility as measured by ejection fraction (EF) without reducing afterload (lowering blood pressure) or increasing heart rate (figure 4). EF enhancements were driven by a significant normalization of end systolic volume (ESV), whereas end diastolic volume (EDV) was unaffected by the model or drug treatment (figure S2). Although cardiac fibrosis and pro-atrial natriuretic peptide (proANP) levels were measured, no significant changes were detectable in the model and no baseline influences were observed with the KEAP1 blocker (figure S2). In dobutamine stress testing designed to evaluate contractile reserve, the model demonstrated significant depression of LV developed pressure (dp/dt-max), however no effect was observed following treatment with GSK3407931B (not shown).

**Figure 4.**
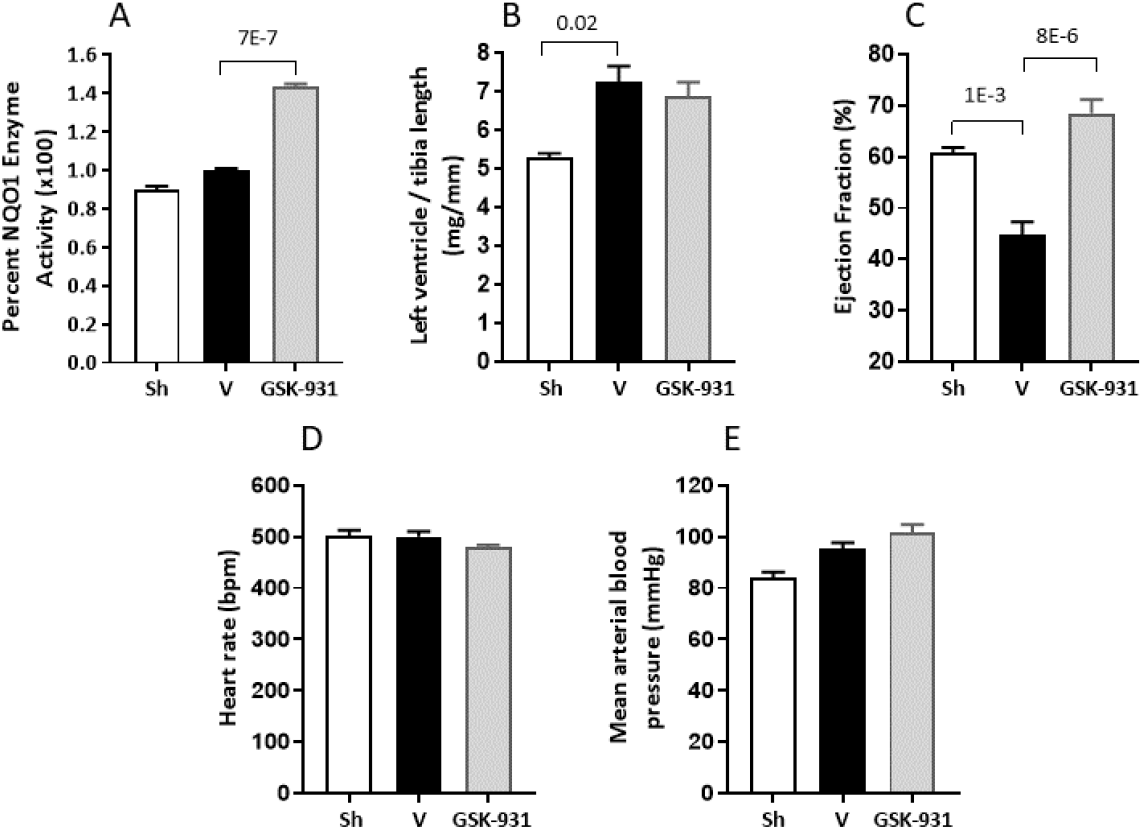
Effect of the KEAP1-kelch domain blocker GSK3407931B on cardiac function, remodeling, and NRF2 pathway activation in the murine 10-week TAC model. Figures labeled as follows: Sh (sham, n=6), V (TAC and no drug, n=16), GSK-931 (TAC and GSK3407931B), 100mg/kg/d, n=13) A) Left ventricular NQO1 enzyme activity B) Left ventricular weight normalized to tibia length C) Ejection fraction, percent of LV volume D) Heart rate, (bpm) beats per min E) mean arterial pressure, millimeters of mercury (mmHg). Drug administration was initiated 2 weeks after TAC surgery. Data are represented as mean (± SEM). P-values were determined by one-way ANOVA with Dunnett’s post hoc multiple comparison test.

To verify the cardiac function findings observed with KEAP1-kelch domain blockade, two additional KEAP1 blockers, GSK3776143B (in dose-response) and GSK3525714B (*28*) (figure S1) along with a repeat of GSK3407931B were evaluated in the murine 10-week TAC model. Over the course of the 8-weeks of compound exposure, the 3 blockers elicited a full restoration of cardiac contractility (figure 5A). For GSK3776143B which was tested in dose-response, the functional effects were dose-related. Part of the apparent EF restoration (approximately 6 out of the 15 EF point differential) measured at end of the study was attributable to the compounds attenuating the continuing EF decline in the model from the 2-week randomization to the 10-week end of study time point (figure 5B). Significant and dose related KEAP1/NRF2 pathway engagement was observed in the study as assessed by LV NQO1 enzyme activity (figure 5C). No effects on heart rate were noted (figure 5F), and as observed previously contractility enhancements were driven predominantly by significant reductions in end systolic volumes (figure S3). Isovolumetric relaxation time, a measure of diastolic function, although significantly prolonged in the model was not improved by KEAP1 blockers in this study (not shown). As a measure of antioxidant activity, protein carbonylation of the sarcomeric protein cMyBPC (*21*) was measured. Two of the 3 compounds significantly suppressed carbonylation of cMyBPC; GSK3525714B trended lower but did not achieve statistical significance. Notably, suppression of cMyBPC carbonylation with GSK3776143B was not dose-related (at the dose-range tested) and near maximal suppression at the lowest dose of 30mg/kg was not associated with an effect on ejection fraction. In aggregate, the data indicated the antioxidation effects of KEAP1-kelch blockade, at least as it pertains to cMyBPC carbonylation, were not tightly correlated with the improvements in contractile performance. No effects on LV hypertrophy or broader cardiac remodeling were observed (figure 5E & S3). Except for GSK3407931B, the compounds did not significantly affect body weight or survival (figure S4). In separate TAC studies, and with the aim of investigating the reported myocardial NRF2-proteostasis connection (*29*), the accumulation of myocardial ubiquitinated protein aggregates was investigated in the model. Although the accumulation of ubiquitinated proteins trended higher in the TAC model, the variability of the measure was too large and could not be demonstrated as significantly different from sham (not shown).

**Figure 5.**
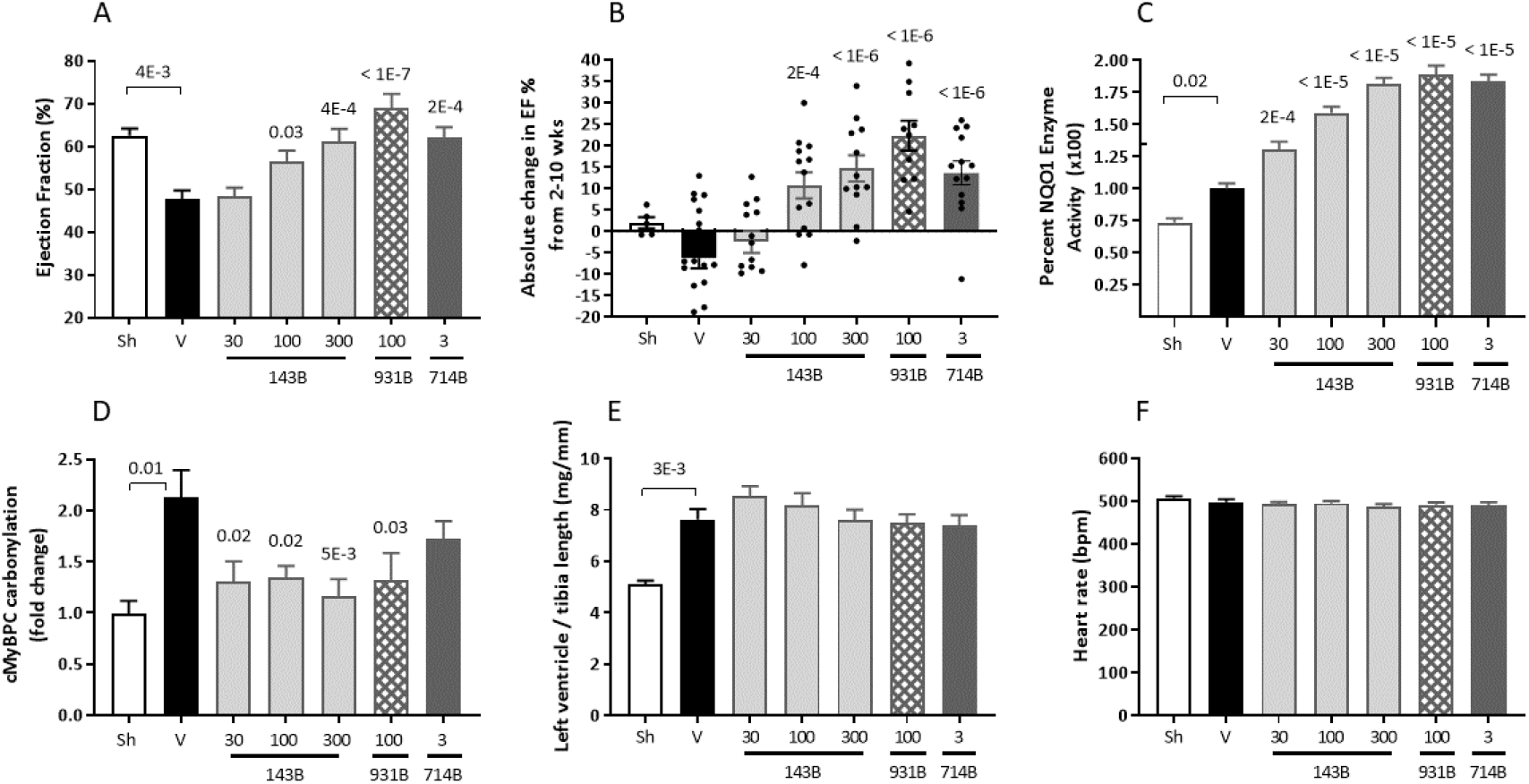
Effect of the KEAP1-kelch domain blockers GSK3776143B (143B), GSK3407931B (931B), and GSK3525714B (714B) on cardiac function, remodeling, NQO1 enzyme activity, and protein oxidation in the murine 10-week TAC model. All compounds were dosed in chow beginning 2-weeks after TAC. Figures labeled as follows: Sh (sham, n=6), V (TAC and chow no drug, n=21), 143B [TAC and GSK3776143B in chow at 30mg/kg/d (n=14), 100 mg/kg/d (n=14) and 300 mg/kg/d (n=13)], 931B [TAC and GSK3407931B in chow at 100 mg/kg/d, n=11], 714B [TAC and GSK3525714B in chow at 3 mg/kg/d, n=13]. A) Ejection fraction, percent of LV volume B) absolute change in ejection fraction from time of animal randomization (at the week 2 echo) to the final EF measure at week-10 C) Left ventricular NQO1 enzyme activity, vehicle value set at 100%, Note: y-axis inward tick was the NQO1 activity achieved by GSK3161593A in the initial TAC findings [figure 3] D) LV cMyBPC carbonylation, fold change E) Left ventricular weights normalized to tibia length F) Heart rate, (bpm) beats per min. Data are represented as mean (± SEM). P-values were determined by one-way ANOVA with Dunnett’s multiple comparison test. All p-values (top of bars) in comparison to vehicle unless otherwise noted.

GSK3776143B was also evaluated in an acute rat myocardial ischemia-reperfusion model where compound was dosed daily for 1-week (in full dose-response) prior to 30-minutes of cardiac ischemia followed by reperfusion and infarct size measured 24 hours later. No effect on left ventricular infarct development (fibrosis area) was observed (not shown). Functional and remodeling measures in this initial ischemic study were not assessed.

To assess the effects of oxidant stress and KEAP1 blockade in the context of human cardiomyocyte contractility, a human iPSC-CM contractility assay was established. Human iPSC-CM were pretreated with GSK3776143B for 48 hours and then exposed to 25uM tert-butyl hydroperoxide (tBHP) for 2 days which induced a significant oxidant cellular stress and depressed cardiomyocyte contraction. Over the 2-days of the combined GSK3776143B and tBHP exposure, hourly contractility endpoints (per methods) were assessed. GSK3776143B, with a concentration dependency, protected iPSC-CMs from tBHP-oxidation-induced loss of contractility (figure 6).

**Figure 6.**
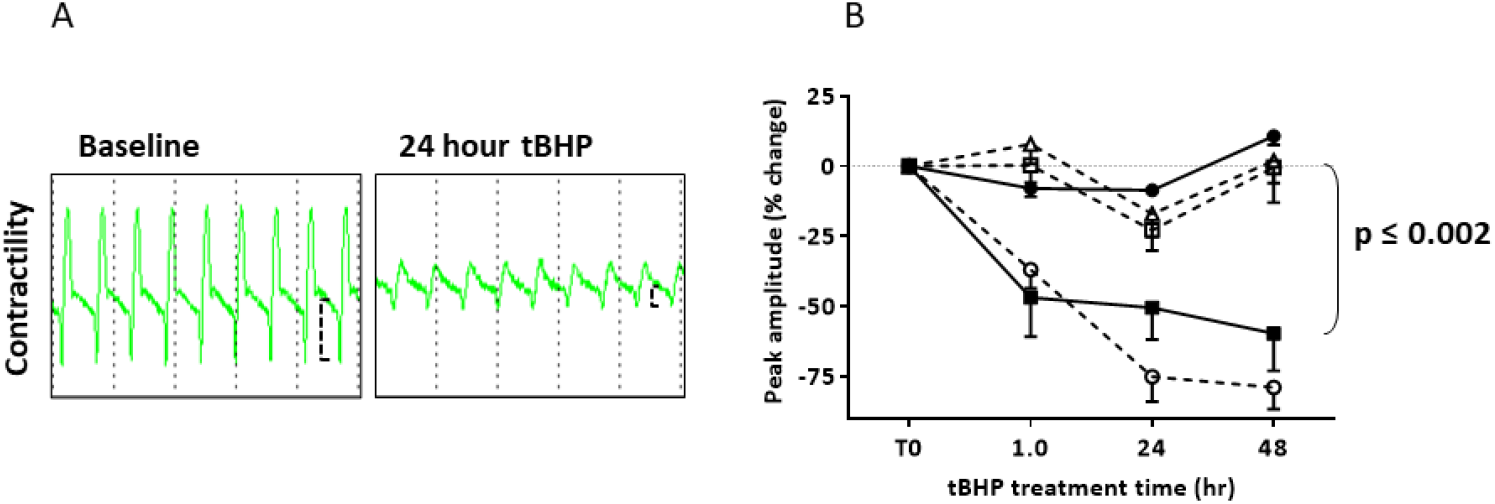
Effect of the KEAP1-kelch domain blocker GSK3776143B on tBHP oxidant stressed human iPSC-CM contractility. A) representative trace of CardioECR contractility demonstrating amplitude change B) Time-course and concentration response of GSK3776143B (+/-) tBHP in CardioECR contractility assay. Symbols represent the following: filled circle (•) no treatment, filled square (◼) treatment with tBHP alone (25 uM), open symbols (○, ◻, Δ) treatment with tBHP and 10, 30, 100 nM GSK3776143B respectively (n=4-6/group). Data represented as means (± SEM), p-value related to 30 and 100 nM GSK3776143B + tBHP in comparison to tBHP alone; determined by one-way ANOVA with Dunnett’s post hoc multiple comparison test.

## Discussion

In the studies described herein, the reversal of a HF transcriptomic signature was used as the basis of a phenotypic screen in search of cardioprotective mechanisms. Conceptually, the HF signature genes were not hypothesized individually as key culprit genes or pathways underpinning the HF phenotype, but rather as sentinel genes, that in aggregate may reflect potentially thousands of other unmeasured nuanced GE changes that collectively manifest as HF. A central finding of these experiments was that this phenotypic screening strategy was able to identify mechanisms that could ameliorate or reverse the decompensated mouse TAC cardiac phenotype. The screen in human iPSC-CMs identified KEAP1-Kelch domain blockers as modulators of a HF GE expression signature and were demonstrated to fully restore cardiac contractile function in a chronic murine LV pressure overload model of HF with reduced ejection fraction. Although the screen output identified multiple small molecules with greater GE effects than KEAP1 modulation, those molecules (other than TRβ agonists) were not successfully connected with a single protein target or mechanism of action to enable follow-up. The findings suggest that more effective GE modulator mechanisms may exist, and that improved methodologies for mining small molecule interactomes may identify additional targets or networks of targets that could influence the HF gene expression signature. The identification of thyroid receptor-β agonist served as an encouraging validation signal for the screen design given the established role of thyroid signaling in adaptive cardiac remodeling (*30*).

KEAP1-kelch domain blockade led to profound improvement in cardiac function in the setting of LV pressure overload. The KEAP1 mode of action predictably led to NRF2 mediated antioxidant endpoints such as the elevated expression of NQO1 and the suppression of protein oxidation. The cardioprotective profile was not associated with suppression of LV hypertrophy, changes in diastolic dysfunction (*31*), or improvements in cardiac reserve. The functional changes appeared to be predominantly related to normalized systolic function and did not influence blood pressure or heart rate. The results most closely resembled the NRF2 cardiomyocyte-specific transgene studies where cardiac overexpression of NRF2 at the time of TAC normalized fractional shortening (*29*) and appeared distinct from neurohumoral mechanisms that unload the heart (*32*). The lack of infarct protection in the setting of acute ischemic injury by KEAP1-kelch compounds was like that reported with the Bardoxolone methyl derivative DH404, which also observed no effect of compound pretreatment on infarct development following acute ischemia-reperfusion(*33*).

The full physiological spectrum of KEAP1 and/or NRF2 biology that underpinned the profound restoration of cardiac contractile efficiency was not elucidated. The transcriptional and mechanistic footprint of NRF2 is substantial. The most well described NRF2 transcriptional activity relates to approximately 250 genes containing an anti-oxidant response element (ARE), however the NRF2 target gene list has been predicted to be in the thousands and its protein interactome in the hundreds (*34, 35*). The working hypothesis was that KEAP1-kelch domain blockade would liberate NRF2 signaling and its attendant activation of antioxidant, anti-inflammatory, and cytoprotective pathways would underpin any observed cardio-protection. This was evidenced in part in the human iPSC-CM studies, where the compounds afforded total protection of contractility while under an intense oxidation and cytotoxicity challenge with tBHP.

Although loss-of-function mutations in the sarcomeric protein cMyBPC have been identified as a major cause of hypertrophic cardiomyopathy (*36, 37*), and carbonylation of cMyBPC protein has been demonstrated to impair its function (*21*), the correlation of cMyBPC carbonylation and LV contractile efficiency in the TAC model was not strong. After 8 weeks of exposure to KEAP1-kelch blockers, full suppression of cMyBPC oxidation did not result in a significant improvement in EF (figure 5, 30mg/kg GSK3776143B). Ultimately, in these TAC studies, carbonylation of cMyBPC was tracked as a reliable surrogate measure of cardiac protein oxidation and activation of the KEAP1-NRF2 anti-oxidation pathway. The relevance of the cMyBPC carbonylation single endpoint as it pertains to the molecular remodeling required to restore degraded cardiac contractile efficiency remains to be established. Given the reported widespread redox regulation of sarcomeric Ca^++^ handling proteins (*38*) and oxidant modification of myofilament proteins (*39*), the KEAP1-NRF2 blocker anti-oxidant activity could reasonably be expected to have a significantly more complex and comprehensive influence on sarcomeric homeostasis. Characterization of the redox regulated sarcomeric proteome as a function of KEAP1-kelch blockade will be the subject of follow-up investigations.

The findings in this report indicated that KEAP1-kelch domain blockade could restore depressed and deteriorating cardiac function in the setting of severe hemodynamic stress without reducing blood pressure or increasing heart rate. Further investigations will be aimed at characterizing the time at which the EF enhancements emerge and whether there is a measurable influence on cardiac function in the setting of ischemic injury. The compounds utilized in these studies selectively bind to the KEAP1-kelch domain with at least one of the consequences being the activation of the NRF2 pathway (*28*). The same KEAP1-kelch domains that bind to NRF2 have also been shown to interact with a complex proteome (*40*). The degree to which these non-NRF2 interactions are influenced by small molecule KEAP1-kelch domain blockade, and whether these mechanisms impact the observed cardiac phenotype remains to be determined.

## List of supplementary materials

Supplementary figures S1-S4. See separate file.

Supplementary table S1. See separate files.

Supplementary tables S2-S3. See separate file.

## Acknowledgements

KEAP compounds described herein are part of a collaboration agreement with Astex Pharmaceuticals. We’d like to thank James Callahan and Jeff Kerns for supply of GSK3161593A, GSK3175696A and GSK3407931B and for helpful consultation.

## Author contributions

Conceptualization: JRT, RNW, JJU, BGL, JJL, ETG, LJJ

Methodology: CGS, QC, DTS, REB, WB

Software: CTL

Formal analysis: MEB, JGB

Investigation: JJU, CGS, BGL, QC, DTS, TPC, CAB, MEB, CTL, AEZ, IM, JB, JGB, CYI, SM, REB, GCP, SE, WB, ED, JCK, JAK, KH, SA, MR, ARO, KR.

Funding acquisition: RNW, JJL

Project administration: ETG

Supervision: RNW, JRT, CGS

Writing: JRT

Writing / review / editing: RNW, CGS, BGL, ARO, JJU

## Data and materials availability

All data are available in the main text or the supplementary materials.

