## Supplemental figures S1-S4 for "A phenotypic screen identified KEAP1-kelch domain blockade as a mechanism to restore cardiac function in the setting of chronic severe hemodynamic stress"

**Supplementary figures S1-S4 for:** Identification of KEAP1-kelch domain blockers as cardio-protective agents in the setting of chronic severe hemodynamic stress

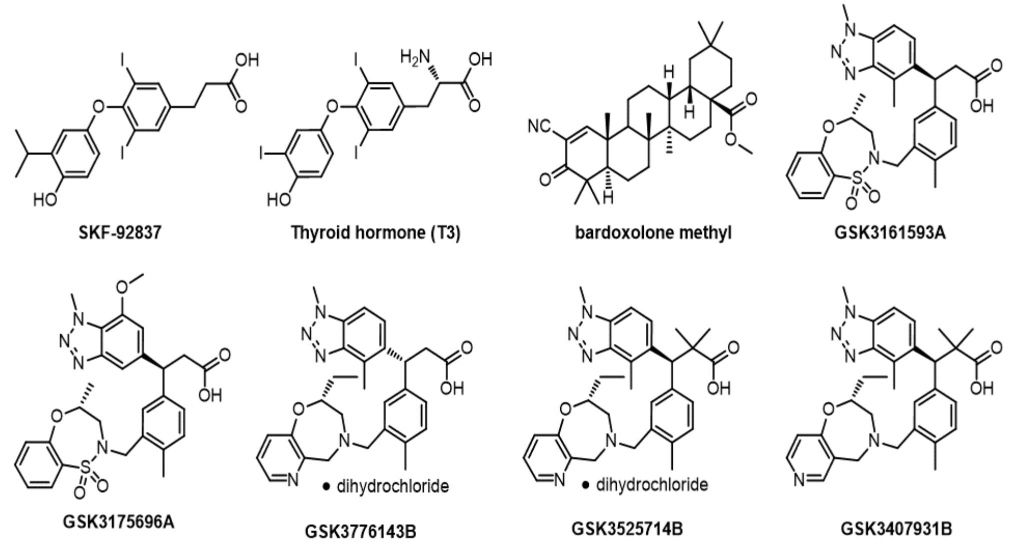


**Figure S1:** ***Compound structures identified in the phenotypic screen and in chemical optimization.***

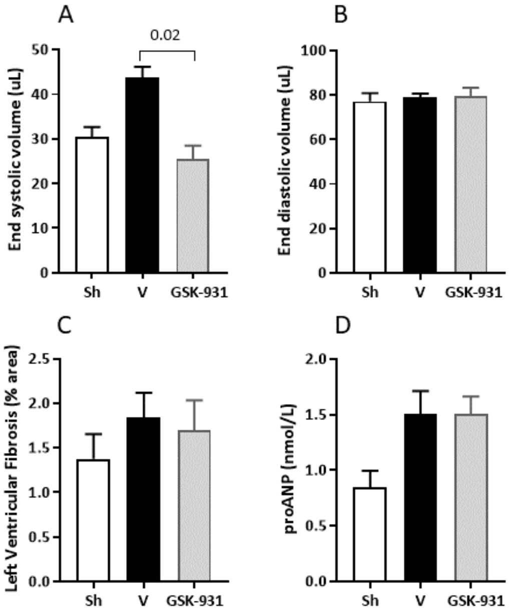


**Figure S2:** ***Effect of the KEAP1-kelch domain blocker GSK3407931B on cardiac function, fibrosis, and remodeling in the murine 10-week TAC model.*** Figures labeled as follows: Sh (sham, no TAC, n=6), V (TAC and no drug, n=16), GSK-931 (TAC and GSK3407931B, 100mg/kg/d, n=13). A) End systolic volume B) End diastolic volume C) LV fibrosis, percent of total LV area D) Plasma proANP. Drug administration was initiated 2-weeks after TAC surgery in the chow. Data are represented as mean (± SEM). P-values were determined by one-way ANOVA with Dunnett's multiple comparison test.

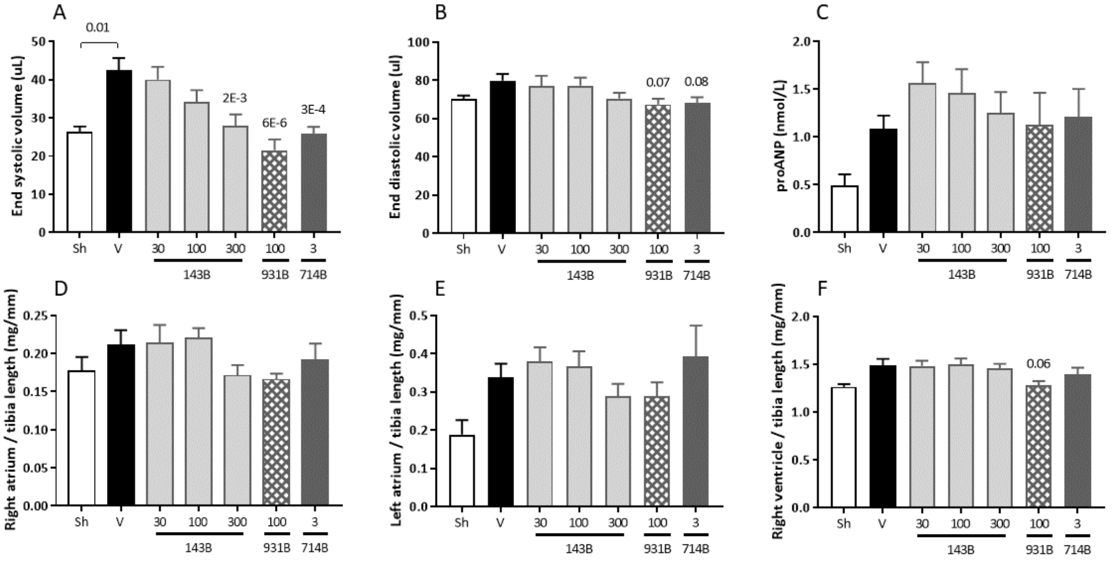


**Figure S3:** ***Effect of the KEAP1-kelch domain blockers GSK3776143B (143B), GSK3407931B (931B), and GSK3525714B (714B) on cardiac function and remodeling in the murine 10-week TAC model.*** Figures labeled as follows: Sh (sham, no TAC, n=6), V (TAC no drug, n=21), 143B [TAC and GSK3773143B at 30mg/kg/d (n=14), 100 mg/kg/d (n=14) and 300 mg/kg/d (n=13)], 931B [TAC and GSK3407931B at 100 mg/kg/d, n=11], 714B [TAC and GSK3525714B at 3 mg/kg/d, n=13]. A) End systolic volume B) End diastolic volume C) Plasma pro-ANP D) Right atrium weights normalized to tibia length E) Left atrium weights normalized to tibia length F) Right ventricle weights normalized to tibia length. Drug administration was initiated 2-weeks after TAC surgery in the chow. Data are represented as mean (± SEM). P-values (top of bars) were determined by one-way ANOVA with Dunnett's multiple comparison test. All p-values in comparison to vehicle unless otherwise noted.

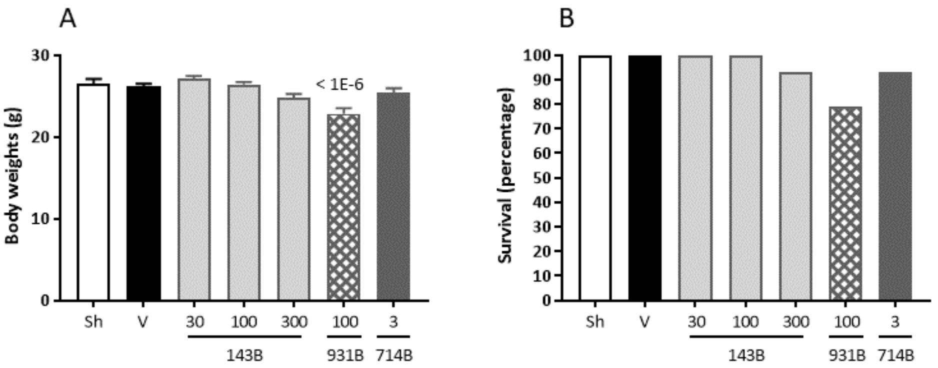


**Figure S4:** ***Effect of the KEAP1-kelch domain blockers GSK3776143B (143B), GSK3407931B (931B), and GSK3525714B (714B) on body weight and survival in the murine 10-week TAC model.*** Figures labeled as follows: Sh (sham, no TAC, n=6), V (TAC no drug, n=21), 143B [TAC and GSK3773143B at 30mg/kg/d (n=14), 100 mg/kg/d (n=14) and 300 mg/kg/d (n=13)], 931B [TAC and GSK3407931B at 100 mg/kg/d, n=11], 714B [TAC and GSK3525714B at 3 mg/kg/d, n=13]. A) body weights (grams, mean, ± SEM) at end of study. P-values (in comparison to vehicle on top of bars) were determined by one-way ANOVA with Dunnett's multiple comparison test. B) Percent survival.
