## Supplemental table S1 for "A phenotypic screen identified KEAP1-kelch domain blockade as a mechanism to restore cardiac function in the setting of chronic severe hemodynamic stress"

Supplementary table S1

| Probe Set | Probe set suffix | Fold change | TAC vs SHAM (p value) | Gene Symbol | Gene Description |
|---|---|---|---|---|---|
| 1435135 | at | -1.11 | 0.890 | Aadacl1 | arylacetamide deacetylase-like 1 |
| 1452988 | at | -1.09 | 0.192 | Aak1 | AP2 associated kinase 1 |
| 1454754 | a_at | -1.06 | 0.469 | Aamp | angio-associated migratory protein |
| 1423685 | at | 1.10 | 0.467 | Aars | alanyl-tRNA synthetase |
| 1436017 | at | 1.13 | 0.200 | Aarsl | alanyl-tRNA synthetase like |
| 1456625 | at | -1.02 | 0.855 | Aasdhppt | aminoadipate-semialdehyde dehydrogenase-phosphopantetheinyl transferase |
| 1432394 | a_at | -1.10 | 0.096 | Aatf | apoptosis antagonizing transcription factor |
| 1433855 | at | 1.44 | 0.006 | Abat | 4-aminobutyrate aminotransferase |
| 1421840 | at | -1.05 | 0.415 | Abca1 | ATP-binding cassette, sub-family A (ABC1), member 1 |
| 1451731 | at | 1.05 | 0.541 | Abca3 | ATP-binding cassette, sub-family A (ABC1), member 3 |
| 1427371 | at | 1.24 | 0.002 | Abca8a | ATP-binding cassette, sub-family A (ABC1), member 8a |
| 1440879 | at | 1.25 | 0.004 | Abca9 | ATP-binding cassette transporter sub-family A member 9 |
| 1416402 | at | -1.10 | 0.045 | Abcb10 | ATP-binding cassette, sub-family B (MDR/TAP), member 10 |
| 1419758 | at | -1.15 | 0.115 | Abcb1a | ATP-binding cassette, sub-family B (MDR/TAP), member 1A |
| 1449818 | at | 1.34 | 0.001 | Abcb4 | ATP-binding cassette, sub-family B (MDR/TAP), member 4 |
| 1422524 | at | -1.01 | 0.858 | Abcb6 | ATP-binding cassette, sub-family B (MDR/TAP), member 6 |
| 1423713 | at | -1.47 | 0.004 | Abcb8 | ATP-binding cassette, sub-family B (MDR/TAP), member 8 |
| 1416263 | at | -1.05 | 0.694 | Abcb9 | ATP-binding cassette, sub-family B (MDR/TAP), member 9 |
| 1452233 | at | 1.37 | 0.000 | Abcc1 | ATP-binding cassette, sub-family C (CFTR/MRP), member 1 |
| 1435684 | at | -1.04 | 0.681 | Abcc5 | ATP-binding cassette, sub-family C (CFTR/MRP), member 5 |
| 1455765 | a_at | -1.20 | 0.015 | Abcc8 | ATP-binding cassette, sub-family C (CFTR/MRP), member 8 |
| 1435751 | at | -1.54 | 0.000 | Abcc9 | ATP-binding cassette, sub-family C (CFTR/MRP), member 9 |
| 1418838 | at | -1.09 | 0.249 | Abcd1 | ATP-binding cassette, sub-family D (ALD), member 1 |
| 1419572 | a_at | 1.05 | 0.673 | Abcd4 | ATP-binding cassette, sub-family D (ALD), member 4 |
| 1442071 | at | 1.06 | 0.557 | Abce1 | ATP-binding cassette, sub-family E (OABP), member 1 |
| 1426747 | at | 1.08 | 0.144 | Abcf3 | ATP-binding cassette, sub-family F (GCN20), member 3 |
| 1423570 | at | 1.45 | 0.011 | Abcg1 | ATP-binding cassette, sub-family G (WHITE), member 1 |
| 1422906 | at | 1.06 | 0.159 | Abcg2 | ATP-binding cassette, sub-family G (WHITE), member 2 |
| 1425699 | a_at | -1.05 | 0.382 | Abhd14a | abhydrolase domain containing 14A |
| 1435653 | at | 1.17 | 0.124 | Abhd2 | abhydrolase domain containing 2 |
| 1416315 | at | 1.15 | 0.040 | Abhd4 | abhydrolase domain containing 4 |
| 1419104 | at | -1.37 | 0.000 | Abhd6 | abhydrolase domain containing 6 |
| 1423178 | at | 1.47 | 0.003 | Abi1 | abl-interactor 1 |
| 1433985 | at | 1.04 | 0.418 | Abi2 | abl-interactor 2 |
| 1423999 | at | -1.12 | 0.312 | Abl1 | v-abl Abelson murine leukemia oncogene 1 |
| 1455682 | at | 1.07 | 0.475 | Abl2 | v-abl Abelson murine leukemia viral oncogene 2 (arg, Abelson-related gene) |
| 1454708 | at | -1.00 | 0.967 | Ablim1 | actin-binding LIM protein 1 |
| 1424751 | at | 1.04 | 0.641 | Abt1 | activator of basal transcription |
| 1417000 | at | -1.10 | 0.168 | Abtb1 | ankyrin repeat and BTB (POZ) domain containing 1 |

Supplementary table S1

| | | | | | |
|---|---|---|---|---|---|
| 1416946 | a_at | -1.13 | 0.033 | Acaa1 | acetyl-Coenzyme A acyltransferase 1 |
| 1428145 | at | -1.88 | 0.005 | Acaa2 | acetyl-Coenzyme A acyltransferase 2 (mitochondrial 3-oxoacyl-Coenzyme A thiolase) |
| 1427052 | at | -1.17 | 0.002 | Acacb | acetyl-Coenzyme A carboxylase beta |
| 1448987 | at | -1.03 | 0.141 | Acadl | acetyl-Coenzyme A dehydrogenase, long-chain |
| 1415984 | at | -1.11 | 0.001 | Acadm | acetyl-Coenzyme A dehydrogenase, medium chain |
| 1460216 | at | -1.09 | 0.409 | Acads | acyl-Coenzyme A dehydrogenase, short chain |
| 1424184 | at | -1.16 | 0.001 | Acadvl | acyl-Coenzyme A dehydrogenase, very long chain |
| 1422479 | at | 1.20 | 0.001 | Acas2 | acetyl-Coenzyme A synthetase 2 (ADP forming) |
| 1416617 | at | -1.17 | 0.009 | Acas2l | acetyl-Coenzyme A synthetase 2 (AMP forming)-like |
| 1424183 | at | -1.09 | 0.304 | Acat1 | acetyl-Coenzyme A acetyltransferase 1 |
| 1425195 | a_at | -1.10 | 0.187 | Acat2 | acetyl-Coenzyme A acetyltransferase 2 |
| 1418073 | at | 1.43 | 0.000 | Acate2 | acyl-Coenzyme A thioesterase 2, mitochondrial |
| 1428668 | at | 1.27 | 0.008 | Acbd3 | acyl-Coenzyme A binding domain containing 3 |
| 1453199 | at | -1.06 | 0.637 | Acbd6 | acyl-Coenzyme A binding domain containing 6 |
| 1435088 | at | -1.13 | 0.059 | Acd | adrenocortical dysplasia |
| 1451911 | a_at | 2.04 | 0.000 | Ace | angiotensin converting enzyme |
| 1416568 | a_at | -1.10 | 0.204 | Acin1 | apoptotic chromatin condensation inducer 1 |
| 1451666 | at | 1.01 | 0.767 | Acly | ATP citrate lyase |
| 1456614 | at | 1.08 | 0.140 | Acn9 | ACN9 homolog (S. cerevisiae) |
| 1451002 | at | -1.10 | 0.031 | Aco2 | aconitase 2, mitochondrial |
| 1416408 | at | 1.11 | 0.409 | Acox1 | acyl-Coenzyme A oxidase 1, palmitoyl |
| 1450720 | at | -1.10 | 0.507 | Acp1 | acid phosphatase 1, soluble |
| 1424654 | at | -1.03 | 0.580 | Acp2 | acid phosphatase 2, lysosomal |
| 1448445 | at | -1.41 | 0.007 | Acp6 | acid phosphatase 6, lysophosphatidic |
| 1422526 | at | -1.21 | 0.052 | Acsl1 | acyl-CoA synthetase long-chain family member 1 |
| 1433531 | at | 1.04 | 0.569 | Acsl4 | acyl-CoA synthetase long-chain family member 4 |
| 1428082 | at | 1.32 | 0.048 | Acsl5 | acyl-CoA synthetase long-chain family member 5 |
| 1451257 | at | -1.31 | 0.001 | Acsl6 | acyl-CoA synthetase long-chain family member 6 |
| 1427735 | a_at | 3.95 | 0.021 | Acta1 | actin, alpha 1, skeletal muscle |
| 1415927 | at | 1.01 | 0.753 | Actc1 | actin, alpha, cardiac |
| 1416569 | at | 1.08 | 0.103 | Actl6a | actin-like 6A |
| 1428585 | at | 1.96 | 0.000 | Actn1 | actinin, alpha 1 |
| 1423449 | a_at | 1.03 | 0.794 | Actn4 | actinin alpha 4 |
| 1417157 | at | 1.01 | 0.790 | Actr10 | ARP10 actin-related protein 10 homolog (S. cerevisiae) |
| 1415873 | a_at | 1.02 | 0.671 | Actr1a | ARP1 actin-related protein 1 homolog A (yeast) |
| 1452057 | at | 1.25 | 0.055 | Actr1b | ARP1 actin-related protein 1 homolog B (yeast) |
| 1452051 | at | 1.12 | 0.014 | Actr3 | ARP3 actin-related protein 3 homolog (yeast) |
| 1428502 | at | 1.14 | 0.108 | Actr6 | ARP6 actin-related protein 6 homolog (yeast) |
| 1423385 | at | 1.03 | 0.404 | Actr8 | ARP8 actin-related protein 8 homolog (S. cerevisiae) |
| 1448460 | at | 1.02 | 0.861 | Acvr1 | activin A receptor, type 1 |
| 1433725 | at | -1.14 | 0.022 | Acvr1b | activin A receptor, type 1B |

Supplementary table S1

| | | | | | |
|---|---|---|---|---|---|
| 1437382 | at | -1.03 | 0.494 | Acvr2 | activin receptor IIA |
| 1420532 | at | -1.25 | 0.125 | Acvrinp1 | activin receptor interacting protein 1 |
| 1435825 | at | 1.21 | 0.041 | Acvrl1 | activin A receptor, type II-like 1 |
| 1419173 | at | -1.62 | 0.000 | Acy1 | aminoacylase 1 |
| 1448539 | a_at | -1.32 | 0.001 | Acy3 | aspartoacylase (aminoacylase) 3 |
| 1450095 | a_at | 1.20 | 0.001 | Acyp1 | acylphosphatase 1, erythrocyte (common) type |
| 1427943 | at | -1.00 | 0.955 | Acyp2 | acylphosphatase 2, muscle type |
| 1416080 | at | 1.11 | 0.376 | Adam15 | a disintegrin and metalloproteinase domain 15 (metargidin) |
| 1418403 | at | 1.02 | 0.953 | Adam19 | a disintegrin and metalloproteinase domain 19 (meltrin beta) |
| 1416094 | at | 1.41 | 0.000 | Adam9 | a disintegrin and metalloproteinase domain 9 (meltrin gamma) |
| 1450716 | at | 1.83 | 0.001 | Adamts1 | a disintegrin-like and metalloprotease (reprolysin type) with thrombospondin type 1 motif, 1 |
| 1427139 | at | 1.24 | 0.038 | Adamts10 | a disintegrin-like and metalloprotease (reprolysin type) with thrombospondin type 1 motif, 10 |
| 1457058 | at | 2.37 | 0.000 | Adamts2 | a disintegrin-like and metalloprotease (reprolysin type) with thrombospondin type 1 motif, 2 |
| 1422561 | at | -1.16 | 0.335 | Adamts5 | a disintegrin-like & metalloprot (reprolysin type) c/ thrombospondin type 1 motif, 5 (aggrecanase-2) |
| 1437785 | at | 1.20 | 0.050 | Adamts9 | a disintegrin-like and metalloprotease (reprolysin type) with thrombospondin type 1 motif, 9 |
| 1434932 | at | 1.04 | 0.752 | Adarb1 | adenosine deaminase, RNA-specific, B1 |
| 1423762 | at | -1.14 | 0.170 | Adck1 | aarF domain containing kinase 1 |
| 1442658 | at | -1.37 | 0.073 | Adck2 | aarF domain containing kinase 2 |
| 1418098 | at | -1.12 | 0.169 | Adcy4 | adenylate cyclase 4 |
| 1418128 | at | -1.50 | 0.000 | Adcy6 | adenylate cyclase 6 |
| 1434454 | at | -1.41 | 0.003 | Adcy9 | adenylate cyclase 9 |
| 1439125 | at | 1.03 | 0.758 | Add1 | adducin 1 (alpha) |
| 1423298 | at | -1.24 | 0.019 | Add3 | adducin 3 (gamma) |
| 1448318 | at | -1.05 | 0.335 | Adfp | adipose differentiation related protein |
| 1416225 | at | -1.59 | 0.005 | Adh1 | alcohol dehydrogenase 1 (class I) |
| 1416185 | a_at | -1.08 | 0.023 | Adh5 | alcohol dehydrogenase 5 (class III), chi polypeptide |
| 1424392 | at | -1.76 | 0.000 | Adhfe1 | alcohol dehydrogenase, iron containing, 1 |
| 1416077 | at | -1.13 | 0.371 | Adm | adrenomedullin |
| 1423069 | at | 1.01 | 0.776 | Adnp | activity-dependent neuroprotective protein |
| 1460710 | at | -1.34 | 0.036 | Adora2a | adenosine A2a receptor |
| 1451079 | at | 1.07 | 0.331 | Adpgk | ADP-dependent glucokinase |
| 1448333 | at | 1.18 | 0.004 | Adprh | ADP-ribosylarginine hydrolase |
| 1455952 | at | 1.14 | 0.076 | Adprhl1 | ADP-ribosylhydrolase like 1 |
| 1460061 | at | -1.89 | 0.003 | Adra1a | adrenergic receptor, alpha 1a |
| 1451992 | at | -1.11 | 0.165 | Adrbk1 | adrenergic receptor kinase, beta 1 |
| 1419263 | a_at | -1.06 | 0.521 | Adrm1 | adhesion regulating molecule 1 |
| 1418372 | at | -1.11 | 0.025 | Adsl | adenylosuccinate lyase |
| 1460726 | at | 1.30 | 0.008 | Adss | adenylosuccinate synthetase, non muscle |
| 1449383 | at | 1.16 | 0.108 | Adssl1 | adenylosuccinate synthetase like 1 |
| 1450637 | a_at | 1.63 | 0.007 | Aebp1 | AE binding protein 1 |
| 1434080 | at | 1.20 | 0.178 | Aebp2 | AE binding protein 2 |

Supplementary table S1

| | | | | | |
|---|---|---|---|---|---|
| 1420619 | a_at | -1.21 | 0.006 | Aes | amino-terminal enhancer of split |
| 1449045 | at | -1.05 | 0.369 | Afg3l1 | AFG3(ATPase family gene 3)-like 1 (yeast) |
| 1427206 | at | 1.12 | 0.176 | Afg3l2 | AFG3(ATPase family gene 3)-like 2 (yeast) |
| 1434665 | at | -1.10 | 0.005 | Aga | aspartylglucosaminidase |
| 1425354 | a_at | 1.30 | 0.003 | Aggf1 | angiogenic factor with G patch and FHA domains 1 |
| 1429083 | at | -1.17 | 0.911 | Agl | amylo-1,6-glucosidase, 4-alpha-glucanotransferase |
| 1421025 | at | -1.08 | 0.355 | Agpat1 | 1-acylglycerol-3-phosphate O-acyltransferase 1 (lysophosphatidic acid acyltransferase, alpha) |
| 1428821 | at | -1.05 | 0.637 | Agpat2 | 1-acylglycerol-3-phosphate O-acyltransferase 2 (lysophosphatidic acid acyltransferase, beta) |
| 1433817 | at | -1.35 | 0.158 | Agpat3 | 1-acylglycerol-3-phosphate O-acyltransferase 3 |
| 1434919 | at | -1.03 | 0.768 | Agps | alkylglycerone phosphate synthase |
| 1426670 | at | 1.18 | 0.004 | Agrn | agrin |
| 1418332 | a_at | -1.18 | 0.009 | Agtpbp1 | ATP/GTP binding protein 1 |
| 1436739 | at | -1.28 | 0.040 | Agtr1 | angiotensin receptor 1 |
| 1460596 | at | -1.12 | 0.134 | Agtrap | angiotensin II, type I receptor-associated protein |
| 1423037 | at | 1.05 | 0.772 | Agtrl1 | angiotensin receptor-like 1 |
| 1417125 | at | 1.11 | 0.085 | Ahcy | S-adenosylhomocysteine hydrolase |
| 1455177 | at | 1.17 | 0.058 | Ahi1 | Abelson helper integration site |
| 1452217 | at | 1.06 | 0.058 | Ahnak | AHNAK nucleoprotein (desmoyokin) |
| 1422631 | at | -1.08 | 0.416 | Ahr | aryl-hydrocarbon receptor |
| 1424147 | at | -1.03 | 0.685 | Ahsa1 | AHA1, activator of heat shock 90kDa protein ATPase homolog 1 (yeast) |
| 1420679 | a_at | -1.22 | 0.015 | Aig1 | androgen-induced 1 |
| 1423128 | at | -1.01 | 0.859 | Aip | aryl-hydrocarbon receptor-interacting protein |
| 1422184 | a_at | -1.03 | 0.554 | Ak1 | adenylate kinase 1 |
| 1448450 | at | -1.36 | 0.301 | Ak2 | adenylate kinase 2 |
| 1421830 | at | -1.03 | 0.630 | Ak3l1 | adenylate kinase 3 alpha-like 1 |
| 1449019 | at | -1.26 | 0.000 | Akap1 | A kinase (PRKA) anchor protein 1 |
| 1455461 | at | 1.07 | 0.360 | Akap10 | A kinase (PRKA) anchor protein 10 |
| 1419706 | a_at | 1.02 | 0.580 | Akap12 | A kinase (PRKA) anchor protein (gravin) 12 |
| 1455870 | at | 1.61 | 0.000 | Akap2 | A kinase (PRKA) anchor protein 2 |
| 1440859 | at | 1.07 | 0.184 | Akap6 | A kinase (PRKA) anchor protein 6 |
| 1450983 | at | -1.14 | 0.494 | Akap8 | A kinase (PRKA) anchor protein 8 |
| 1417734 | at | -1.28 | 0.035 | Akap8l | A kinase (PRKA) anchor protein 8-like |
| 1435769 | at | -1.18 | 0.873 | Akap9 | A kinase (PRKA) anchor protein (yotiao) 9 |
| 1423611 | at | 1.01 | 0.867 | Akp2 | alkaline phosphatase 2, liver |
| 1451035 | a_at | 1.01 | 0.751 | Akr1a4 | aldo-keto reductase family 1, member A4 (aldehyde reductase) |
| 1448319 | at | -1.11 | 0.011 | Akr1b3 | aldo-keto reductase family 1, member B3 (aldose reductase) |
| 1417826 | at | -1.04 | 0.458 | Akr1e1 | aldo-keto reductase family 1, member E1 |
| 1417294 | at | -1.36 | 0.001 | Akr7a5 | aldo-keto reductase family 7, member A5 (aflatoxin aldehyde reductase) |
| 1416657 | at | -1.07 | 0.148 | Akt1 | thymoma viral proto-oncogene 1 |
| 1437736 | at | -1.11 | 0.555 | Akt1s1 | AKT1 substrate 1 (proline-rich) |
| 1424877 | a_at | -1.28 | 0.026 | Alad | aminolevulinate, delta-, dehydratase |

Supplementary table S1

| | | | | | |
|---|---|---|---|---|---|
| 1424126 | at | -1.11 | 0.218 | Alas1 | aminolevulinic acid synthase 1 |
| 1451675 | a_at | -1.19 | 0.206 | Alas2 | aminolevulinic acid synthase 2, erythroid |
| 1415836 | at | 1.41 | 0.036 | Aldh18a1 | aldehyde dehydrogenase 18 family, member A1 |
| 1416468 | at | 1.56 | 0.000 | Aldh1a1 | aldehyde dehydrogenase family 1, subfamily A1 |
| 1451260 | at | 1.06 | 0.608 | Aldh1b1 | aldehyde dehydrogenase 1 family, member B1 |
| 1434987 | at | -1.43 | 0.000 | Aldh2 | aldehyde dehydrogenase 2, mitochondrial |
| 1415776 | at | -1.09 | 0.318 | Aldh3a2 | aldehyde dehydrogenase family 3, subfamily A2 |
| 1452375 | at | -1.21 | 0.080 | Aldh4a1 | aldehyde dehydrogenase 4 family, member A1 |
| 1453065 | at | -1.59 | 0.000 | Aldh5a1 | aldehyde dehydrogenase family 5, subfamily A1 |
| 1448104 | at | -1.16 | 0.017 | Aldh6a1 | aldehyde dehydrogenase family 6, subfamily A1 |
| 1460167 | at | -1.04 | 0.300 | Aldh7a1 | aldehyde dehydrogenase family 7, member A1 |
| 1436689 | a_at | -1.39 | 0.006 | Aldh9a1 | aldehyde dehydrogenase 9, subfamily A1 |
| 1451194 | at | -6.00 | 0.000 | Aldob | aldolase 2, B isoform |
| 1451368 | at | 1.11 | 0.211 | Alg1 | asparagine-linked glycosylation 1 homolog (yeast, beta-1,4-mannosyltransferase) |
| 1424818 | at | -1.20 | 0.055 | Alg12 | asparagine-linked glycosylation 12 homolog (yeast, alpha-1,6-mannosyltransferase) |
| 1421059 | a_at | -1.06 | 0.246 | Alg2 | asparagine-linked glycosylation 2 homolog (yeast, alpha-1,3-mannosyltransferase) |
| 1460742 | at | 1.02 | 0.779 | Alg3 | asparagine-linked glycosylation 3 homolog (yeast, alpha-1,3-mannosyltransferase) |
| 1422699 | at | -1.45 | 0.020 | Alox12 | arachidonate 12-lipoxygenase |
| 1441962 | at | -1.74 | 0.000 | Alox5 | arachidonate 5-lipoxygenase |
| 1452016 | at | 1.37 | 0.003 | Alox5ap | arachidonate 5-lipoxygenase activating protein |
| 1449573 | at | 1.21 | 0.001 | Alpk3 | alpha-kinase 3 |
| 1417783 | at | 1.01 | 0.897 | Als2 | amyotrophic lateral sclerosis 2 (juvenile) homolog (human) |
| 1433769 | at | -1.01 | 0.940 | Als2cl | ALS2 C-terminal like |
| 1434010 | at | -1.26 | 0.063 | Als2cr13 | amyotrophic lateral sclerosis 2 (juvenile) chromosome region, candidate 13 |
| 1433544 | at | 1.16 | 0.092 | Als2cr2 | amyotrophic lateral sclerosis 2 (juvenile) chromosome region, candidate 2 homolog (human) |
| 1435016 | at | 1.52 | 0.001 | Als2cr3 | amyotrophic lateral sclerosis 2 (juvenile) chromosome region, candidate 3 homolog(human) |
| 1433794 | at | 1.11 | 0.063 | Als4 | amyotrophic lateral sclerosis 4 homolog (human) |
| 1458710 | at | -1.00 | 0.979 | Amac1 | acyl-malonyl condensing enzyme 1 |
| 1417208 | at | -1.16 | 0.051 | Amacr | alpha-methylacyl-CoA racemase |
| 1448484 | at | -1.30 | 0.000 | Amd1 | S-adenosylmethionine decarboxylase 1 |
| 1450102 | a_at | 1.16 | 0.000 | Amfr | autocrine motility factor receptor |
| 1455112 | at | 1.11 | 0.211 | Amid | apoptosis-inducing factor (AIF)-like mitochondrion-associated inducer of death |
| 1454890 | at | 2.26 | 0.000 | Amot | angiomotin |
| 1452387 | a_at | -1.16 | 0.082 | Amotl2 | angiomotin like 2 |
| 1422573 | at | 1.05 | 0.297 | Ampd3 | AMP deaminase 3 |
| 1443768 | at | -1.24 | 0.079 | Amt | aminomethyltransferase |
| 1417765 | a_at | -1.68 | 0.000 | Amy1 | amylase 1, salivary |
| 1429375 | at | -1.11 | 0.359 | Anapc10 | anaphase promoting complex subunit 10 |
| 1417326 | a_at | -1.04 | 0.348 | Anapc11 | anaphase promoting complex subunit 11 homolog (yeast) |
| 1452586 | at | -1.25 | 0.000 | Anapc13 | anaphase promoting complex subunit 13 |
| 1415707 | at | 1.21 | 0.003 | Anapc2 | anaphase promoting complex subunit 2 |

Supplementary table S1

| | | | | | |
|---|---|---|---|---|---|
| 1423930 | at | 1.04 | 0.355 | Anapc4 | anaphase promoting complex subunit 4 |
| 1416906 | at | -1.08 | 0.032 | Anapc5 | anaphase-promoting complex subunit 5 |
| 1418458 | at | 1.25 | 0.024 | Anapc7 | anaphase promoting complex subunit 7 |
| 1446656 | at | -1.99 | 0.045 | Angpt1 | angiopoietin 1 |
| 1455090 | at | 1.06 | 0.235 | Angptl2 | angiopoietin-like 2 |
| 1424485 | at | -1.89 | 0.000 | Angptl3 | angiopoietin-like 3 |
| 1450627 | at | -1.16 | 0.068 | Ank | progressive ankylosis |
| 1434264 | at | -1.25 | 0.004 | Ank2 | ankyrin 2, brain |
| 1417685 | at | 1.10 | 0.187 | Ankfy1 | ankyrin repeat and FYVE domain containing 1 |
| 1448008 | at | -1.22 | 0.114 | Ankhd1 | ankyrin repeat and KH domain containing 1 |
| 1429193 | at | 1.08 | 0.106 | Ankib1 | ankyrin repeat and IBR domain containing 1 |
| 1451724 | at | -1.03 | 0.597 | Ankmy2 | ankyrin repeat and MYND domain containing 2 |
| 1420991 | at | 2.27 | 0.000 | Ankrd1 | ankyrin repeat domain 1 (cardiac muscle) |
| 1448199 | at | 1.59 | 0.000 | Ankrd10 | ankyrin repeat domain 10 |
| 1440193 | at | -1.12 | 0.175 | Ankrd12 | ankyrin repeat domain 12 |
| 1433742 | at | 1.12 | 0.092 | Ankrd15 | ankyrin repeat domain 15 |
| 1428239 | at | 1.07 | 0.342 | Ankrd16 | ankyrin repeat domain 16 |
| 1428183 | at | 1.71 | 0.002 | Ankrd23 | ankyrin repeat domain 23 |
| 1425731 | at | -1.50 | 0.000 | Ankrd24 | ankyrin repeat domain 24 |
| 1460559 | at | 1.07 | 0.267 | Ankrd25 | ankyrin repeat domain 25 |
| 1454717 | at | 1.13 | 0.143 | Ankrd27 | ankyrin repeat domain 27 (VPS9 domain) |
| 1423577 | at | -1.32 | 0.001 | Ankrd32 | ankyrin repeat domain 32 |
| 1452204 | at | -1.42 | 0.048 | Anks1 | ankyrin repeat and SAM domain containing 1 |
| 1421918 | at | -1.09 | 0.166 | Anp32a | acidic (leucine-rich) nuclear phosphoprotein 32 family, member A |
| 1417082 | at | 1.19 | 0.002 | Anp32b | acidic nuclear phosphoprotein 32 family, member B |
| 1451446 | at | 1.37 | 0.007 | Antxr1 | anthrax toxin receptor 1 |
| 1426708 | at | -1.33 | 0.000 | Antxr2 | anthrax toxin receptor 2 |
| 1429642 | at | 1.06 | 0.643 | Anubl1 | AN1, ubiquitin-like, homolog (Xenopus laevis) |
| 1448213 | at | 1.61 | 0.000 | Anxa1 | annexin A1 |
| 1418468 | at | -1.27 | 0.009 | Anxa11 | annexin A11 |
| 1419091 | a_at | 1.37 | 0.011 | Anxa2 | annexin A2 |
| 1460330 | at | 1.47 | 0.014 | Anxa3 | annexin A3 |
| 1421223 | a_at | 1.35 | 0.000 | Anxa4 | annexin A4 |
| 1425567 | a_at | 1.38 | 0.000 | Anxa5 | annexin A5 |
| 1415818 | at | -1.14 | 0.007 | Anxa6 | annexin A6 |
| 1416137 | at | 1.27 | 0.017 | Anxa7 | annexin A7 |
| 1449396 | at | -1.05 | 0.487 | Aoc3 | amine oxidase, copper containing 3 |
| 1433649 | at | -1.03 | 0.805 | Aof1 | amine oxidase, flavin containing 1 |
| 1426761 | at | -1.03 | 0.504 | Aof2 | amine oxidase (flavin containing) domain 2 |
| 1419435 | at | -1.07 | 0.425 | Aox1 | aldehyde oxidase 1 |
| 1416279 | at | 1.01 | 0.932 | Ap1b1 | adaptor protein complex AP-1, beta 1 subunit |

Supplementary table S1

| | | | | | |
|---|---|---|---|---|---|
| 1434081 | at | -1.02 | 0.653 | Ap1g1 | adaptor protein complex AP-1, gamma 1 subunit |
| 1457695 | at | 1.21 | 0.000 | Ap1gbp1 | AP1 gamma subunit binding protein 1 |
| 1416307 | at | 1.01 | 0.875 | Ap1m1 | adaptor-related protein complex AP-1, mu subunit 1 |
| 1416087 | at | 1.07 | 0.365 | Ap1s1 | adaptor protein complex AP-1, sigma 1 |
| 1452657 | at | 1.43 | 0.000 | Ap1s2 | adaptor-related protein complex 1, sigma 2 subunit |
| 1460724 | at | 1.14 | 0.151 | Ap2a1 | adaptor protein complex AP-2, alpha 1 subunit |
| 1450894 | a_at | 1.05 | 0.313 | Ap2m1 | adaptor protein complex AP-2, mu1 |
| 1433612 | at | -1.15 | 0.014 | Ap2s1 | adaptor-related protein complex 2, sigma 1 subunit |
| 1417964 | at | 1.01 | 0.887 | Ap3d1 | adaptor-related protein complex 3, delta 1 subunit |
| 1416374 | at | -0.53 | 0.770 | Ap3m1 | adaptor-related protein complex 3, mu 1 subunit |
| 1417527 | at | -1.06 | 0.596 | Ap3m2 | adaptor-related protein complex 3, mu 2 subunit |
| 1422593 | at | 1.33 | 0.005 | Ap3s1 | adaptor-related protein complex 3, sigma 1 subunit |
| 1435718 | at | -1.10 | 0.162 | Ap3s2 | adaptor-related protein complex 3, sigma 2 subunit |
| 1416631 | at | 1.11 | 0.158 | Ap4b1 | adaptor-related protein complex AP-4, beta 1 |
| 1456948 | at | -1.01 | 0.864 | Ap4e1 | adaptor-related protein complex AP-4, epsilon 1 |
| 1451665 | a_at | -1.15 | 0.022 | Ap4s1 | adaptor-related protein complex AP-4, sigma 1 |
| 1452870 | at | -1.05 | 0.425 | Apaf1 | apoptotic protease activating factor 1 |
| 1416182 | at | -1.17 | 0.097 | Apba3 | amyloid beta (A4) precursor protein-binding, family A, member 3 |
| 1423892 | at | -1.89 | 0.000 | Apbb1 | amyloid beta (A4) precursor protein-binding, family B, member 1 |
| 1426719 | at | 1.01 | 0.852 | Apbb2 | amyloid beta (A4) precursor protein-binding, family B, member 2 |
| 1451886 | at | -1.16 | 0.494 | Apeg1 | aortic preferentially expressed gene 1 |
| 1426030 | a_at | -1.11 | 0.185 | Apeh | acylpeptide hydrolase |
| 1416135 | at | 1.27 | 0.007 | Apex1 | apurinic/apyrimidinic endonuclease 1 |
| 1419081 | at | -1.10 | 0.160 | Apg10l | autophagy 10-like (S. cerevisiae) |
| 1451747 | a_at | 1.03 | 0.679 | Apg12l | autophagy 12-like (S. cerevisiae) |
| 1428169 | at | 1.07 | 0.377 | Apg16l | APG16 autophagy 16-like (S. cerevisiae) |
| 1448993 | at | 1.22 | 0.052 | Apg3l | APG3 autophagy 3-like (S. cerevisiae) |
| 1434662 | at | 1.07 | 0.350 | Apg4a | APG4 (ATG4) autophagy-related homolog A (S. cerevisiae) |
| 1424103 | at | 1.02 | 0.688 | Apg4b | APG4 (ATG4) autophagy-related homolog B (S. cerevisiae) |
| 1434014 | at | 1.03 | 0.756 | Apg4c | APG4 (ATG4) autophagy-related homolog C (S. cerevisiae) |
| 1433918 | at | -1.13 | 0.124 | Apg4d | APG4 (ATG4) autophagy-related homolog D (S. cerevisiae) |
| 1415684 | at | 1.05 | 0.626 | Apg5l | autophagy 5-like (S. cerevisiae) |
| 1435784 | at | 1.01 | 0.844 | Apg9l1 | APG9 autophagy 9-like 1 (S. cerevisiae) |
| 1424979 | at | -1.59 | 0.000 | Aph1a | anterior pharynx defective 1a homolog (C. elegans) |
| 1435793 | at | -1.04 | 0.551 | Aph1b | anterior pharynx defective 1b homolog (C. elegans) |
| 1415813 | at | -1.15 | 0.002 | Api5 | apoptosis inhibitor 5 |
| 1418254 | at | -1.16 | 0.001 | Apip | APAF1 interacting protein |
| 1421889 | a_at | -1.18 | 0.062 | Aplp2 | amyloid beta (A4) precursor-like protein 2 |
| 1415727 | at | -1.07 | 0.074 | Apoa1bp | apolipoprotein A-I binding protein |
| 1451755 | a_at | 1.11 | 0.298 | Apobec1 | apolipoprotein B editing complex 1 |
| 1417889 | at | -1.11 | 0.003 | Apobec2 | apolipoprotein B editing complex 2 |

Supplementary table S1

| | | | | | |
|---|---|---|---|---|---|
| 1416371 | at | 2.67 | 0.000 | Apod | apolipoprotein D |
| 1432466 | a_at | -1.07 | 0.468 | Apoe | apolipoprotein E |
| 1423781 | at | -1.05 | 0.299 | Appbp1 | amyloid beta precursor protein binding protein 1 |
| 1423801 | a_at | 1.12 | 0.189 | Aprt | adenine phosphoribosyl transferase |
| 1450921 | at | -1.01 | 0.790 | Aptx | aprataxin |
| 1416203 | at | -1.01 | 0.923 | Aqp1 | aquaporin 1 |
| 1434449 | at | -2.38 | 0.002 | Aqp4 | aquaporin 4 |
| 1455647 | at | -1.12 | 0.444 | Ar | androgen receptor |
| 1419441 | at | -1.04 | 0.093 | Arbp | acidic ribosomal phosphoprotein P0 |
| 1417118 | a_at | -1.34 | 0.000 | Ard1 | N-acetyltransferase ARD1 homolog (S. cerevisiae) |
| 1416458 | at | -1.11 | 0.176 | Arf2 | ADP-ribosylation factor 2 |
| 1434312 | at | 1.21 | 0.002 | Arf6 | ADP-ribosylation factor 6 |
| 1427245 | at | 1.04 | 0.714 | Arfgap1 | ADP-ribosylation factor GTPase activating protein 1 |
| 1426534 | a_at | 1.52 | 0.006 | Arfgap3 | ADP-ribosylation factor GTPase activating protein 3 |
| 1424240 | at | 1.06 | 0.276 | Arfip2 | ADP-ribosylation factor interacting protein 2 |
| 1425507 | at | 1.07 | 0.752 | Arfrp1 | ADP-ribosylation factor related protein 1 |
| 1451309 | at | 1.00 | 0.992 | Arhgap1 | Rho GTPase activating protein 1 |
| 1426027 | a_at | -1.06 | 0.637 | Arhgap10 | Rho GTPase activating protein 10 |
| 1451525 | at | -1.02 | 0.651 | Arhgap12 | Rho GTPase activating protein 12 |
| 1426623 | a_at | 1.06 | 0.394 | Arhgap17 | Rho GTPase activating protein 17 |
| 1438882 | at | -1.04 | 0.755 | Arhgap18 | Rho GTPase activating protein 18 |
| 1429918 | at | -1.41 | 0.003 | Arhgap20 | Rho GTPase activating protein 20 |
| 1428368 | at | 1.01 | 0.804 | Arhgap21 | Rho GTPase activating protein 21 |
| 1419745 | at | 1.08 | 0.418 | Arhgap23 | Rho GTPase activating protein 23 |
| 1424842 | a_at | 1.21 | 0.021 | Arhgap24 | Rho GTPase activating protein 24 |
| 1424249 | a_at | 1.73 | 0.040 | Arhgap9 | Rho GTPase activating protein 9 |
| 1451168 | a_at | 1.16 | 0.202 | Arhgdia | Rho GDP dissociation inhibitor (GDI) alpha |
| 1426454 | at | 1.35 | 0.032 | Arhgdib | Rho, GDP dissociation inhibitor (GDI) beta |
| 1421164 | a_at | -1.02 | 0.824 | Arhgef1 | Rho guanine nucleotide exchange factor (GEF) 1 |
| 1452302 | at | 1.05 | 0.402 | Arhgef10 | Rho guanine nucleotide exchange factor (GEF) 10 |
| 1434926 | at | -1.01 | 0.934 | Arhgef11 | Rho guanine nucleotide exchange factor (GEF) 11 |
| 1457172 | at | 1.02 | 0.821 | Arhgef15 | Rho guanine nucleotide exchange factor (GEF) 15 |
| 1418553 | at | -1.01 | 0.830 | Arhgef18 | rho/rac guanine nucleotide exchange factor (GEF) 18 |
| 1437629 | at | -1.25 | 0.018 | Arhgef19 | Rho guanine nucleotide exchange factor (GEF) 19 |
| 1424250 | a_at | 1.17 | 0.007 | Arhgef3 | Rho guanine nucleotide exchange factor (GEF) 3 |
| 1452304 | a_at | 1.25 | 0.038 | Arhgef5 | Rho guanine nucleotide exchange factor (GEF) 5 |
| 1429012 | at | 1.14 | 0.046 | Arhgef6 | Rac/Cdc42 guanine nucleotide exchange factor (GEF) 6 |
| 1448709 | at | -1.03 | 0.516 | Arid1a | AT rich interactive domain 1A (Swi1 like) |
| 1455979 | at | -1.38 | 0.031 | Arid1b | AT rich interactive domain 1B (Swi1 like) |
| 1454990 | at | -1.14 | 0.219 | Arid2 | AT rich interactive domain 2 (Arid-rfx like) |
| 1456420 | at | -1.13 | 0.350 | Arid4a | AT rich interactive domain 4A (Rbp1 like) |

Supplementary table S1

| | | | | | |
|---|---|---|---|---|---|
| 1435768 | at | -1.09 | 0.245 | Arid4b | AT rich interactive domain 4B (Rbp1 like) |
| 1427189 | at | 1.22 | 0.007 | Arih1 | ariadne ubiquitin-conjugating enzyme E2 binding protein homolog 1 (Drosophila) |
| 1451025 | at | 1.04 | 0.669 | Arl1 | ADP-ribosylation factor-like 1 |
| 1424640 | at | 1.32 | 0.000 | Arl10b | ADP-ribosylation factor-like 10B |
| 1418819 | at | 1.01 | 0.884 | Arl10c | ADP-ribosylation factor-like 10C |
| 1422549 | at | -1.10 | 0.194 | Arl2 | ADP-ribosylation factor-like 2 |
| 1417112 | at | 1.19 | 0.011 | Arl2bp | ADP-ribosylation factor-like 2 binding protein |
| 1450706 | a_at | -1.15 | 0.033 | Arl3 | ADP-ribosylation factor-like 3 |
| 1435092 | at | -1.06 | 0.346 | Arl4 | ADP-ribosylation factor-like 4 |
| 1433951 | at | 1.05 | 0.078 | Arl5 | ADP-ribosylation factor-like 5 |
| 1417331 | a_at | 1.30 | 0.003 | Arl6 | ADP-ribosylation factor-like 6 |
| 1423818 | a_at | 1.10 | 0.117 | Arl6ip1 | ADP-ribosylation factor-like 6 interacting protein 1 |
| 1435594 | at | -1.23 | 0.045 | Arl6ip2 | ADP-ribosylation factor-like 6 interacting protein 2 |
| 1440360 | at | 1.01 | 0.887 | Arl6ip5 | ADP-ribosylation factor-like 6 interacting protein 5 |
| 1424020 | at | 1.14 | 0.166 | Arl6ip6 | ADP-ribosylation factor-like 6 interacting protein 6 |
| 1454788 | at | 1.29 | 0.019 | Arl7 | ADP-ribosylation factor-like 7 |
| 1424373 | at | 1.09 | 0.282 | Armcx3 | armadillo repeat containing, X-linked 3 |
| 1428112 | at | -1.34 | 0.003 | Armet | arginine-rich, mutated in early stage tumors |
| 1437042 | at | 1.19 | 0.033 | Arnt | aryl hydrocarbon receptor nuclear translocator |
| 1425099 | a_at | -1.04 | 0.886 | Arntl | aryl hydrocarbon receptor nuclear translocator-like |
| 1429688 | at | -1.03 | 0.696 | Arntl2 | aryl hydrocarbon receptor nuclear translocator-like 2 |
| 1435605 | at | -1.17 | 0.105 | Arp3b | actin-related protein 3-beta |
| 1416079 | a_at | 1.04 | 0.446 | Arpc1a | actin related protein 2/3 complex, subunit 1A |
| 1416226 | at | 1.32 | 0.046 | Arpc1b | actin related protein 2/3 complex, subunit 1B |
| 1437148 | at | 1.20 | 0.000 | Arpc2 | actin related protein 2/3 complex, subunit 2 |
| 1448279 | at | 1.15 | 0.028 | Arpc3 | actin related protein 2/3 complex, subunit 3 |
| 1423588 | at | 1.06 | 0.554 | Arpc4 | actin related protein 2/3 complex, subunit 4 |
| 1448129 | at | 1.34 | 0.000 | Arpc5 | actin related protein 2/3 complex, subunit 5 |
| 1452044 | at | 1.29 | 0.005 | Arpc5l | actin related protein 2/3 complex, subunit 5-like |
| 1460444 | at | 1.27 | 0.149 | Arrb1 | arrestin, beta 1 |
| 1454617 | at | 1.11 | 0.290 | Arrdc3 | arrestin domain containing 3 |
| 1426818 | at | 1.20 | 0.107 | Arrdc4 | arrestin domain containing 4 |
| 1417655 | a_at | 1.02 | 0.798 | Ars2 | arsenate resistance protein 2 |
| 1451372 | a_at | 1.06 | 0.107 | Art1 | ADP-ribosyltransferase 1 |
| 1452474 | a_at | -1.18 | 0.003 | Art3 | ADP-ribosyltransferase 3 |
| 1449174 | at | -1.57 | 0.000 | Art4 | ADP-ribosyltransferase 4 |
| 1451553 | at | -1.66 | 0.013 | Art5 | ADP-ribosyltransferase 5 |
| 1416942 | at | 1.10 | 0.138 | Arts1 | type 1 tumor necrosis factor receptor shedding aminopeptidase regulator |
| 1431980 | a_at | -1.49 | 0.000 | As3mt | arsenic (+3 oxidation state) methyltransferase |
| 1416735 | at | 1.13 | 0.003 | Asah1 | N-acylsphingosine amidohydrolase 1 |
| 1450726 | at | 1.06 | 0.352 | Asah2 | N-acylsphingosine amidohydrolase 2 |

Supplementary table S1

| | | | | | |
|---|---|---|---|---|---|
| 1452068 | at | -1.29 | 0.082 | Asahl | N-acylsphingosine amidohydrolase (acid ceramidase)-like |
| 1434961 | at | 1.11 | 0.035 | Asb1 | ankyrin repeat and SOCS box-containing protein 1 |
| 1421466 | at | -1.55 | 0.005 | Asb10 | ankyrin repeat and SOCS box-containing protein 10 |
| 1422153 | a_at | -1.23 | 0.017 | Asb11 | ankyrin repeat and SOCS box-containing protein 11 |
| 1419401 | at | -1.01 | 0.828 | Asb13 | ankyrin repeat and SOCS box-containing protein 13 |
| 1449547 | at | -1.45 | 0.000 | Asb14 | ankyrin repeat and SOCS box-containing protein 14 |
| 1439836 | at | -1.34 | 0.019 | Asb15 | ankyrin repeat and SOCS box-containing protein 15 |
| 1440603 | at | -1.16 | 0.185 | Asb18 | ankyrin repeat and SOCS box-containing 18 |
| 1428444 | at | -1.27 | 0.004 | Asb2 | ankyrin repeat and SOCS box-containing protein 2 |
| 1417361 | at | -1.03 | 0.551 | Asb3 | ankyrin repeat and SOCS box-containing protein 3 |
| 1433919 | at | -1.39 | 0.083 | Asb4 | ankyrin repeat and SOCS box-containing protein 4 |
| 1449356 | at | 1.19 | 0.005 | Asb5 | ankyrin repeat and SOCs box-containing protein 5 |
| 1438744 | at | -1.04 | 0.729 | Asb7 | ankyrin repeat and SOCS box-containing protein 7 |
| 1460703 | at | -1.12 | 0.079 | Ascc1 | activating signal cointegrator 1 complex subunit 1 |
| 1457069 | at | 1.19 | 0.023 | Ascc3 | activating signal cointegrator 1 complex subunit 3 |
| 1454772 | at | -1.08 | 0.059 | Ascc3l1 | activating signal cointegrator 1 complex subunit 3-like 1 |
| 1430138 | at | 1.04 | 0.618 | Ase1 | CD3 epsilon associated protein; antisense to Ercc1 |
| 1423511 | at | -1.08 | 0.052 | Asf1a | ASF1 anti-silencing function 1 homolog A (S. cerevisiae) |
| 1420616 | at | 1.04 | 0.424 | Ash2l | ash2 (absent, small, or homeotic)-like (Drosophila) |
| 1418292 | at | -1.31 | 0.001 | Asna1 | arsA (bacterial) arsenite transporter, ATP-binding, homolog 1 |
| 1451095 | at | 1.20 | 0.120 | Asns | asparagine synthetase |
| 1416652 | at | 1.55 | 0.002 | Aspn | asporin |
| 1423912 | at | 1.41 | 0.006 | Aspscr1 | alveolar soft part sarcoma chromosome region, candidate 1 (human) |
| 1460597 | at | 1.02 | 0.798 | Asxl2 | additional sex combs like 2 (Drosophila) |
| 1448763 | at | -1.06 | 0.202 | Atad1 | ATPase family, AAA domain containing 1 |
| 1436174 | at | 1.08 | 0.314 | Atad2 | ATPase family, AAA domain containing 2 |
| 1422461 | at | -1.10 | 0.291 | Atad3a | ATPase family, AAA domain containing 3A |
| 1453267 | at | -1.05 | 0.815 | Atbf1 | AT motif binding factor 1 |
| 1458369 | at | 1.36 | 0.024 | Atcay | ataxia, cerebellar, Cayman type homolog (human) |
| 1420036 | at | -1.03 | 0.699 | Ate1 | arginine-tRNA-protein transferase 1 |
| 1417296 | at | 1.04 | 0.496 | Atf1 | activating transcription factor 1 |
| 1449363 | at | 2.97 | 0.002 | Atf3 | activating transcription factor 3 |
| 1448135 | at | 1.41 | 0.001 | Atf4 | activating transcription factor 4 |
| 1425927 | a_at | 1.37 | 0.015 | Atf5 | activating transcription factor 5 |
| 1428506 | at | 1.32 | 0.000 | Atic | 5-aminoimidazole-4-carboxamide ribonucleotide formyltransferase/IMP cyclohydrolase |
| 1460639 | a_at | 1.04 | 0.643 | Atox1 | ATX1 (antioxidant protein 1) homolog 1 (yeast) |
| 1456388 | at | 1.03 | 0.428 | Atp11a | ATPase, class VI, type 11A |
| 1451388 | a_at | 1.00 | 0.912 | Atp11b | ATPase, Class VI, type 11B |
| 1455083 | at | 1.06 | 0.413 | Atp11c | Atpase, class VI, type 11C |
| 1417475 | at | 1.10 | 0.432 | Atp13a1 | ATPase type 13A1 |
| 1455136 | at | -1.22 | 0.454 | Atp1a2 | ATPase, Na+/K+ transporting, alpha 2 polypeptide |

Supplementary table S1

| | | | | | |
|---|---|---|---|---|---|
| 1451152 | a_at | 1.19 | 0.164 | Atp1b1 | ATPase, Na+/K+ transporting, beta 1 polypeptide |
| 1435148 | at | -1.22 | 0.042 | Atp1b2 | ATPase, Na+/K+ transporting, beta 2 polypeptide |
| 1423126 | at | -1.07 | 0.117 | Atp1b3 | ATPase, Na+/K+ transporting, beta 3 polypeptide |
| 1427250 | at | -1.34 | 0.129 | Atp2a2 | ATPase, Ca++ transporting, cardiac muscle, slow twitch 2 |
| 1420037 | at | -1.01 | 0.254 | Atp5a1 | ATP synthase, H+ transporting, mitochondrial F1 complex, alpha subunit, isoform 1 |
| 1416829 | at | -1.03 | 0.072 | Atp5b | ATP synthase, H+ transporting mitochondrial F1 complex, beta subunit |
| 1438809 | at | -1.27 | 0.001 | Atp5c1 | ATP synthase, H+ transporting, mitochondrial F1 complex, gamma polypeptide 1 |
| 1426742 | at | -1.03 | 0.163 | Atp5f1 | ATP synthase, H+ transporting, mitochondrial F0 complex, subunit b, isoform 1 |
| 1416020 | a_at | -1.05 | 0.028 | Atp5g1 | ATP synthase, H+ transporting, mitochondrial F0 complex, subunit c (subunit 9), isoform 1 |
| 1415980 | at | -1.23 | 0.002 | Atp5g2 | ATP synthase, H+ transporting, mitochondrial F0 complex, subunit c (subunit 9), isoform 2 |
| 1454661 | at | 1.02 | 0.350 | Atp5g3 | ATP synthase, H+ transporting, mitochondrial F0 complex, subunit c (subunit 9), isoform 3 |
| 1423676 | at | -1.07 | 0.003 | Atp5h | ATP synthase, H+ transporting, mitochondrial F0 complex, subunit d |
| 1416143 | at | -1.04 | 0.025 | Atp5j | ATP synthase, H+ transporting, mitochondrial F0 complex, subunit F |
| 1416269 | at | -1.04 | 0.010 | Atp5j2 | ATP synthase, H+ transporting, mitochondrial F0 complex, subunit f, isoform 2 |
| 1422525 | at | -1.11 | 0.001 | Atp5k | ATP synthase, H+ transporting, mitochondrial F1F0 complex, subunit e |
| 1448203 | at | -1.08 | 0.000 | Atp5l | ATP synthase, H+ transporting, mitochondrial F0 complex, subunit g |
| 1416278 | a_at | -1.07 | 0.000 | Atp5o | ATP synthase, H+ transporting, mitochondrial F1 complex, O subunit |
| 1417970 | at | -1.12 | 0.017 | Atp5s | ATP synthase, H+ transporting, mitochondrial F0 complex, subunit s |
| 1423662 | at | 1.17 | 0.005 | Atp6ap2 | ATPase, H+ transporting, lysosomal accessory protein 2 |
| 1417632 | at | -1.01 | 0.871 | Atp6v0a1 | ATPase, H+ transporting, lysosomal V0 subunit a isoform 1 |
| 1434791 | at | -1.28 | 0.000 | Atp6v0a2 | ATPase, H+ transporting, lysosomal V0 subunit a isoform 2 |
| 1416392 | a_at | 1.05 | 0.063 | Atp6v0c | ATPase, H+ transporting, V0 subunit C |
| 1415671 | at | 1.02 | 0.722 | Atp6v0d1 | ATPase, H+ transporting, V0 subunit D isoform 1 |
| 1416328 | a_at | 1.09 | 0.153 | Atp6v0e | ATPase, H+ transporting, V0 subunit |
| 1450634 | at | 1.15 | 0.355 | Atp6v1a1 | ATPase, H+ transporting, V1 subunit A, isoform 1 |
| 1415814 | at | 1.26 | 0.012 | Atp6v1b2 | ATPase, H+ transporting, V1 subunit B, isoform 2 |
| 1416952 | at | 1.07 | 0.005 | Atp6v1d | ATPase, H+ transporting, V1 subunit D |
| 1449711 | at | -1.11 | 0.498 | Atp6v1e1 | ATPase, H+ transporting, V1 subunit E isoform 1 |
| 1423993 | at | -1.16 | 0.000 | Atp6v1f | ATPase, H+ transporting, V1 subunit F |
| 1415826 | at | 1.54 | 0.000 | Atp6v1h | ATPase, H+ transporting, lysosomal 50/57kDa, V1 subunit H |
| 1436921 | at | 1.13 | 0.128 | Atp7a | ATPase, Cu++ transporting, alpha polypeptide |
| 1433965 | at | -1.32 | 0.031 | Atp8a1 | ATPase, aminophospholipid transporter (APLT), class I, type 8A, member 1 |
| 1455396 | at | 1.57 | 0.006 | Atp8b1 | ATPase, class I, type 8B, member 1 |
| 1434026 | at | 1.52 | 0.000 | Atp8b2 | Atpase, class I, type 8B, member 2 |
| 1427604 | a_at | 1.08 | 0.357 | Atp9a | ATPase, class II, type 9A |
| 1450879 | at | 1.38 | 0.009 | Atp9b | ATPas, class II, type 9B |
| 1434934 | at | -1.12 | 0.346 | Atpaf1 | ATP synthase mitochondrial F1 complex assembly factor 1 |
| 1426474 | at | 1.05 | 0.373 | Atpaf2 | ATP synthase mitochondrial F1 complex assembly factor 2 |
| 1448770 | a_at | 1.09 | 0.146 | Atpif1 | ATPase inhibitory factor 1 |
| 1454311 | at | -1.31 | 0.020 | Atrn | attractin |
| 1449666 | at | 1.03 | 0.303 | Atrnl1 | attractin like 1 |

Supplementary table S1

| | | | | | |
|---|---|---|---|---|---|
| 1456832 | at | -1.29 | 0.039 | Atrx | alpha thalassemia/mental retardation syndrome X-linked homolog (human) |
| 1455551 | at | 1.11 | 0.267 | Attp | signaling molecule ATTP |
| 1428279 | a_at | 1.08 | 0.160 | Atxn7l4 | ataxin 7-like 4 |
| 1420776 | a_at | -1.31 | 0.000 | Auh | AU RNA binding protein/enoyl-coenzyme A hydratase |
| 1415742 | at | -1.04 | 0.602 | Aup1 | ancient ubiquitous protein |
| 1438680 | at | -1.36 | 0.028 | Auts2 | autism susceptibility candidate 2 |
| 1423122 | at | 1.17 | 0.024 | Avpi1 | arginine vasopressin-induced 1 |
| 1423586 | at | 1.38 | 0.001 | Axl | AXL receptor tyrosine kinase |
| 1448032 | at | -1.04 | 0.615 | Azi2 | 5-azacytidine induced gene 2 |
| 1452428 | a_at | 1.04 | 0.543 | B2m | beta-2 microglobulin |
| 1434135 | at | 1.24 | 0.323 | B3galnt2 | UDP-GalNAc:betaGlcNAc beta 1,3-galactosaminyltransferase, polypeptide 2 |
| 1435252 | at | 1.17 | 0.074 | B3galt6 | UDP-Gal:betaGal beta 1,3-galactosyltransferase, polypeptide 6 |
| 1416960 | at | 1.05 | 0.239 | B3gat3 | beta-1,3-glucuronyltransferase 3 (glucuronosyltransferase I) |
| 1450026 | a_at | 1.32 | 0.002 | B3gnt1 | UDP-GlcNAc:betaGal beta-1,3-N-acetylglucosaminyltransferase 1 |
| 1452346 | at | -1.10 | 0.207 | B3gnt6 | UDP-GlcNAc:betaGal beta-1,3-N-acetylglucosaminyltransferase 6 |
| 1418014 | a_at | 1.09 | 0.160 | B4galt1 | UDP-Gal:betaGlcNAc beta 1,4- galactosyltransferase, polypeptide 1 |
| 1425934 | a_at | 1.05 | 0.649 | B4galt4 | UDP-Gal:betaGlcNAc beta 1,4-galactosyltransferase, polypeptide 4 |
| 1426303 | at | -1.04 | 0.692 | B4galt7 | xylosylprotein beta1,4-galactosyltransferase, polypeptide 7 (galactosyltransferase I) |
| 1417094 | at | -1.35 | 0.001 | Bach | brain acyl-CoA hydrolase |
| 1440831 | at | -1.10 | 0.093 | Bach1 | BTB and CNC homology 1 |
| 1416394 | at | -1.09 | 0.286 | Bag1 | Bcl2-associated athanogene 1 |
| 1423948 | at | 1.54 | 0.000 | Bag2 | Bcl2-associated athanogene 2 |
| 1422452 | at | 1.36 | 0.001 | Bag3 | Bcl2-associated athanogene 3 |
| 1452628 | at | 1.06 | 0.393 | Bag5 | BCL2-associated athanogene 5 |
| 1445088 | at | -1.20 | 0.087 | Baiap1 | BAI1-associated protein 1 |
| 1424951 | at | 1.17 | 0.025 | Baiap2l1 | BAI1-associated protein 2-like 1 |
| 1418991 | at | 1.51 | 0.000 | Bak1 | BCL2-antagonist/killer 1 |
| 1423753 | at | 1.90 | 0.000 | Bambi | BMP and activin membrane-bound inhibitor, homolog (Xenopus laevis) |
| 1421081 | a_at | -1.20 | 0.007 | Banf1 | barrier to autointegration factor 1 |
| 1426354 | at | 1.04 | 0.671 | Bap1 | Brca1 associated protein 1 |
| 1428572 | at | 1.25 | 0.068 | Basp1 | brain abundant, membrane attached signal protein 1 |
| 1448221 | at | -1.10 | 0.026 | Bat1a | HLA-B-associated transcript 1A |
| 1422799 | at | -1.07 | 0.326 | Bat2 | HLA-B associated transcript 2 |
| 1433890 | a_at | -1.02 | 0.725 | Bat3 | HLA-B-associated transcript 3 |
| 1415732 | at | 1.06 | 0.490 | Bat5 | HLA-B associated transcript 5 |
| 1460692 | at | 1.08 | 0.283 | Bat8 | HLA-B associated transcript 8 |
| 1416837 | at | -1.01 | 0.879 | Bax | Bcl2-associated X protein |
| 1447930 | at | 1.10 | 0.481 | Baz1a | bromodomain adjacent to zinc finger domain 1A |
| 1434824 | at | -1.16 | 0.073 | Baz1b | bromodomain adjacent to zinc finger domain, 1B |
| 1451996 | at | 1.06 | 0.342 | Bbp | beta-amyloid binding protein precursor |
| 1424478 | at | -1.35 | 0.003 | Bbs2 | Bardet-Biedl syndrome 2 homolog (human) |

Supplementary table S1

| | | | | | |
|---|---|---|---|---|---|
| 1430820 | a_at | -1.13 | 0.168 | Bbx | bobby sox homolog (Drosophila) |
| 1417077 | at | -1.11 | 0.184 | Bcap29 | B-cell receptor-associated protein 29 |
| 1416202 | at | -1.08 | 0.020 | Bcap37 | B-cell receptor-associated protein 37 |
| 1415936 | at | -1.18 | 0.228 | Bcar3 | breast cancer anti-estrogen resistance 3 |
| 1425764 | a_at | -1.49 | 0.000 | Bcat2 | branched chain aminotransferase 2, mitochondrial |
| 1448542 | at | 1.07 | 0.099 | Bccip | BRCA2 and CDKN1A interacting protein |
| 1416647 | at | -1.49 | 0.002 | Bckdha | branched chain ketoacid dehydrogenase E1, alpha polypeptide |
| 1427153 | at | -1.79 | 0.000 | Bckdhb | branched chain ketoacid dehydrogenase E1, beta polypeptide |
| 1460644 | at | -1.12 | 0.086 | Bckdk | branched chain ketoacid dehydrogenase kinase |
| 1418970 | a_at | 1.16 | 0.083 | Bcl10 | B-cell leukemia/lymphoma 10 |
| 1456632 | at | -1.16 | 0.344 | Bcl11a | B-cell CLL/lymphoma 11A (zinc finger protein) |
| 1435227 | at | -1.88 | 0.003 | Bcl11b | B-cell leukemia/lymphoma 11B |
| 1457687 | at | 2.36 | 0.000 | Bcl2 | B-cell leukemia/lymphoma 2 |
| 1444565 | at | -1.14 | 0.283 | Bcl2l1 | Bcl2-like 1 |
| 1435448 | at | -1.80 | 0.043 | Bcl2l11 | BCL2-like 11 (apoptosis facilitator) |
| 1423572 | at | 1.26 | 0.006 | Bcl2l2 | Bcl2-like 2 |
| 1421818 | at | 1.41 | 0.003 | Bcl6 | B-cell leukemia/lymphoma 6 |
| 1418421 | at | 1.14 | 0.337 | Bcl6b | B-cell CLL/lymphoma 6, member B |
| 1448247 | at | 1.10 | 0.020 | Bcl7b | B-cell CLL/lymphoma 7B |
| 1449523 | at | 1.02 | 0.822 | Bcl7c | B-cell CLL/lymphoma 7C |
| 1451574 | at | -1.35 | 0.007 | Bcl9 | B-cell CLL/lymphoma 9 |
| 1429438 | at | -1.00 | 0.972 | Bcor | Bcl6 interacting corepressor |
| 1455564 | at | 1.17 | 0.241 | Bcr | breakpoint cluster region homolog |
| 1451541 | at | -1.06 | 0.284 | Bcs1l | BCS1-like (yeast) |
| 1426959 | at | 1.66 | 0.004 | Bdh | 3-hydroxybutyrate dehydrogenase (heart, mitochondrial) |
| 1422168 | a_at | 1.16 | 0.209 | Bdnf | brain derived neurotrophic factor |
| 1460320 | at | 1.33 | 0.000 | Becn1 | beclin 1 (coiled-coil, myosin-like BCL2-interacting protein) |
| 1416866 | at | -1.02 | 0.653 | Bet1 | blocked early in transport 1 homolog (S. cerevisiae) |
| 1426490 | at | -1.05 | 0.281 | Bfar | bifunctional apoptosis regulator |
| 1416405 | at | 2.47 | 0.000 | Bgn | biglycan |
| 1418025 | at | 1.64 | 0.005 | Bhlhb2 | basic helix-loop-helix domain containing, class B2 |
| 1428512 | at | -1.48 | 0.000 | Bhlhb9 | basic helix-loop-helix domain containing, class B9 |
| 1450624 | at | -1.20 | 0.558 | Bhmt | betaine-homocysteine methyltransferase |
| 1441137 | at | 1.19 | 0.328 | Bicc1 | bicaudal C homolog 1 (Drosophila) |
| 1438701 | at | 1.06 | 0.488 | Bicd1 | bicaudal D homolog 1 (Drosophila) |
| 1420362 | a_at | -1.38 | 0.004 | Biklk | Bcl2-interacting killer-like |
| 1417691 | at | 1.20 | 0.232 | Bin3 | bridging integrator 3 |
| 1418854 | at | -1.04 | 0.298 | Birc2 | baculoviral IAP repeat-containing 2 |
| 1421392 | a_at | 1.05 | 0.600 | Birc3 | baculoviral IAP repeat-containing 3 |
| 1420631 | a_at | -1.20 | 0.002 | Blcap | bladder cancer associated protein homolog (human) |
| 1452101 | at | 1.08 | 0.353 | Blmh | bleomycin hydrolase |

Supplementary table S1

| | | | | | |
|---|---|---|---|---|---|
| 1429048 | at | 1.10 | 0.234 | Bloc1s2 | biogenesis of lysosome-related organelles complex-1, subunit 2 |
| 1428580 | at | 1.05 | 0.304 | Blvra | biliverdin reductase A |
| 1451386 | at | -1.28 | 0.036 | Blvrb | biliverdin reductase B (flavin reductase (NADPH)) |
| 1448733 | at | -1.05 | 0.061 | Bmi1 | B lymphoma Mo-MLV insertion region 1 |
| 1458370 | at | 1.27 | 0.075 | Bmp2k | BMP2 inducible kinase |
| 1450759 | at | -1.21 | 0.447 | Bmp6 | bone morphogenetic protein 6 |
| 1429273 | at | -1.80 | 0.001 | Bmper | BMP-binding endothelial regulator |
| 1425491 | at | 1.02 | 0.874 | Bmpr1a | bone morphogenetic protein receptor, type 1A |
| 1419616 | at | 1.03 | 0.733 | Bmpr2 | bone morphogenic protein receptor, type II (serine/threonine kinase) |
| 1433636 | at | 1.02 | 0.730 | Bms1l | BMS1-like, ribosome assembly protein (yeast) |
| 1428669 | at | 1.00 | 0.980 | Bmyc | brain expressed myelocytomatosis oncogene |
| 1427908 | at | -1.06 | 0.241 | Bnip1 | BCL2/adenovirus E1B 19kDa-interacting protein 1, NIP1 |
| 1422470 | at | 1.17 | 0.003 | Bnip3 | BCL2/adenovirus E1B 19kDa-interacting protein 1, NIP3 |
| 1426869 | at | 1.24 | 0.021 | Boc | biregional cell adhesion molecule-related/down-regulated by oncogenes (Cdon) binding protein |
| 1423264 | at | 1.21 | 0.018 | Bop1 | block of proliferation 1 |
| 1415864 | at | 1.24 | 0.017 | Bpgm | 2,3-bisphosphoglycerate mutase |
| 1424242 | at | -1.00 | 0.973 | Bphl | biphenyl hydrolase-like (serine hydrolase, breast epithelial mucin-associated antigen) |
| 1449211 | at | -1.14 | 0.224 | Bpnt1 | bisphosphate 3'-nucleotidase 1 |
| 1425693 | at | -1.10 | 0.430 | Braf | Braf transforming gene |
| 1416738 | at | 1.11 | 0.022 | Brap | BRCA1 associated protein |
| 1434821 | at | -1.08 | 0.254 | Brd1 | bromodomain containing 1 |
| 1437210 | a_at | -1.05 | 0.575 | Brd2 | bromodomain containing 2 |
| 1436960 | at | -1.10 | 0.059 | Brd3 | bromodomain containing 3 |
| 1450711 | at | -1.09 | 0.371 | Brd4 | bromodomain containing 4 |
| 1448521 | at | 1.11 | 0.121 | Brd7 | bromodomain containing 7 |
| 1456351 | at | -1.28 | 0.031 | Brd8 | bromodomain containing 8 |
| 1416924 | at | 1.11 | 0.193 | Bri3 | brain protein I3 |
| 1428743 | at | -1.26 | 0.127 | Bri3bp | Bri3 binding protein |
| 1460443 | at | -1.01 | 0.897 | Brms1l | breast cancer metastasis-suppressor 1-like |
| 1418746 | at | -1.26 | 0.000 | Brp17 | brain protein 17 |
| 1453651 | a_at | -1.03 | 0.121 | Brp44l | brain protein 44-like |
| 1455243 | at | 1.22 | 0.012 | Brpf3 | bromodomain and PHD finger containing, 3 |
| 1456616 | a_at | -1.01 | 0.772 | Bsg | basigin |
| 1424921 | at | 1.04 | 0.726 | Bst2 | bone marrow stromal cell antigen 2 |
| 1455286 | at | 1.11 | 0.013 | Btbd1 | BTB (POZ) domain containing 1 |
| 1428377 | at | -1.69 | 0.025 | Btbd11 | BTB (POZ) domain containing 11 |
| 1429582 | at | -1.05 | 0.309 | Btbd14a | BTB (POZ) domain containing 14A |
| 1427271 | at | 1.06 | 0.153 | Btbd15 | BTB (POZ) domain containing 15 |
| 1424054 | at | -1.17 | 0.126 | Btbd2 | BTB (POZ) domain containing 2 |
| 1433868 | at | -1.09 | 0.051 | Btbd3 | BTB (POZ) domain containing 3 |
| 1454884 | at | 1.00 | 0.953 | Btbd4 | BTB (POZ) domain containing 4 |

Supplementary table S1

| | | | | | |
|---|---|---|---|---|---|
| 1434861 | at | -1.06 | 0.401 | Btbd5 | BTB (POZ) domain containing 5 |
| 1435153 | at | -1.14 | 0.008 | Btbd6 | BTB (POZ) domain containing 6 |
| 1459011 | at | -1.31 | 0.031 | Btbd9 | BTB (POZ) domain containing 9 |
| 1428289 | at | -1.32 | 0.001 | Bteb1 | basic transcription element binding protein 1 |
| 1423839 | a_at | 1.19 | 0.000 | Btf3 | basic transcription factor 3 |
| 1426083 | a_at | 1.02 | 0.661 | Btg1 | B-cell translocation gene 1, anti-proliferative |
| 1416250 | at | 1.39 | 0.565 | Btg2 | B-cell translocation gene 2, anti-proliferative |
| 1449007 | at | 1.22 | 0.021 | Btg3 | B-cell translocation gene 3 |
| 1448473 | at | 1.02 | 0.744 | Bub3 | budding uninhibited by benzimidazoles 3 homolog (S. cerevisiae) |
| 1425884 | at | 1.03 | 0.671 | Bxdc1 | brix domain containing 1 |
| 1422767 | at | 1.25 | 0.057 | Bysl | bystin-like |
| 1416695 | at | 1.63 | 0.001 | Bzrp | benzodiazepine receptor, peripheral |
| 1423456 | at | -1.33 | 0.008 | Bzw2 | basic leucine zipper and W2 domains 2 |
| 1416886 | at | 1.17 | 0.000 | C1d | nuclear DNA binding protein |
| 1428490 | at | -1.22 | 0.142 | C1galt1 | core 1 UDP-galactose:N-acetylgalactosamine-alpha-R beta 1,3-galactosyltransferase |
| 1416655 | at | -1.05 | 0.283 | C1galt1c1 | C1GALT1-specific chaperone 1 |
| 1417381 | at | 1.37 | 0.005 | C1qa | complement component 1, q subcomponent, alpha polypeptide |
| 1417063 | at | 1.30 | 0.020 | C1qb | complement component 1, q subcomponent, beta polypeptide |
| 1448274 | at | 1.02 | 0.492 | C1qbp | complement component 1, q subcomponent binding protein |
| 1449401 | at | 1.33 | 0.014 | C1qg | complement component 1, q subcomponent, gamma polypeptide |
| 1456046 | at | 1.14 | 0.168 | C1qr1 | complement component 1, q subcomponent, receptor 1 |
| 1455582 | at | -1.16 | 0.242 | C1qtnf1 | C1q and tumor necrosis factor related protein 1 |
| 1423954 | at | 1.12 | 0.074 | C3 | complement component 3 |
| 1419483 | at | -1.18 | 0.199 | C3ar1 | complement component 3a receptor 1 |
| 1418021 | at | 3.13 | 0.000 | C4 | complement component 4 (within H-2S) |
| 1437023 | at | 1.07 | 0.238 | C530015C18 | hypothetical protein C530015C18 |
| 1439902 | at | 1.28 | 0.047 | C5r1 | complement component 5, receptor 1 |
| 1437553 | at | -1.06 | 0.392 | C6.1A | c6.1a protein |
| 1418432 | at | 1.15 | 0.217 | Cab39 | calcium binding protein 39 |
| 1426367 | at | 1.03 | 0.683 | Cab39l | calcium binding protein 39-like |
| 1417066 | at | -1.21 | 0.008 | Cabc1 | chaperone, ABC1 activity of bc1 complex like (S. pombe) |
| 1433686 | at | 1.03 | 0.803 | Cabin1 | calcineurin binding protein 1 |
| 1441744 | at | -1.27 | 0.037 | Cacna1c | calcium channel, voltage-dependent, L type, alpha 1C subunit |
| 1456401 | at | -1.33 | 0.025 | Cacnb2 | calcium channel, voltage-dependent, beta 2 subunit |
| 1452047 | at | 1.00 | 0.962 | Cacybp | calcyclin binding protein |
| 1425814 | a_at | 1.31 | 0.009 | Calcrl | calcitonin receptor-like |
| 1433592 | at | 1.05 | 0.221 | Calm1 | calmodulin 1 |
| 1417606 | a_at | 1.15 | 0.023 | Calr | calreticulin |
| 1431255 | at | -1.19 | 0.072 | Calr3 | calreticulin 3 |
| 1415870 | at | 1.25 | 0.000 | Calu | calumenin |
| 1417604 | at | -1.20 | 0.002 | Camk1 | calcium/calmodulin-dependent protein kinase I |

Supplementary table S1

| | | | | | |
|---|---|---|---|---|---|
| 1457311 | at | -1.28 | 0.020 | Camk2a | calcium/calmodulin-dependent protein kinase II alpha |
| 1448676 | at | -1.40 | 0.031 | Camk2b | calcium/calmodulin-dependent protein kinase II, beta |
| 1423941 | at | 1.00 | 0.933 | Camk2g | calcium/calmodulin-dependent protein kinase II gamma |
| 1455401 | at | 1.37 | 0.004 | Camkk2 | calcium/calmodulin-dependent protein kinase kinase 2, beta |
| 1433709 | at | -1.10 | 0.139 | Cant1 | calcium activated nucleotidase 1 |
| 1428935 | at | -1.19 | 0.068 | Canx | calnexin |
| 1423222 | at | -1.06 | 0.426 | Cap2 | CAP, adenylate cyclase-associated protein, 2 (yeast) |
| 1450355 | a_at | 2.12 | 0.005 | Capg | capping protein (actin filament), gelsolin-like |
| 1416257 | at | 1.06 | 0.362 | Capn2 | calpain 2 |
| 1426400 | a_at | 1.05 | 0.373 | Capns1 | calpain, small subunit 1 |
| 1452038 | at | 1.17 | 0.013 | Capza1 | capping protein (actin filament) muscle Z-line, alpha 1 |
| 1423058 | at | -1.04 | 0.595 | Capza2 | capping protein (actin filament) muscle Z-line, alpha 2 |
| 1448752 | at | -1.26 | 0.007 | Car2 | carbonic anhydrase 2 |
| 1453588 | at | -1.48 | | Car3 | carbonic anhydrase 3 |
| 1448949 | at | -1.67 | 0.000 | Car4 | carbonic anhydrase 4 |
| 1427482 | a_at | 1.43 | 0.001 | Car8 | carbonic anhydrase 8 |
| 1449491 | at | 1.01 | 0.843 | Card10 | caspase recruitment domain family, member 10 |
| 1454733 | at | 1.02 | 0.852 | Card4 | caspase recruitment domain 4 |
| 1460588 | at | 1.13 | 0.022 | Carf | calcium response factor |
| 1415975 | at | 1.97 | 0.000 | Carhsp1 | calcium regulated heat stable protein 1 |
| 1416736 | at | -1.10 | 0.040 | Casc3 | cancer susceptibility candidate 3 |
| 1422518 | at | 1.26 | 0.008 | Cask | calcium/calmodulin-dependent serine protein kinase (MAGUK family) |
| 1450691 | at | -1.12 | 0.296 | Caskin2 | cask-interacting protein 2 |
| 1449265 | at | 1.42 | 0.010 | Casp1 | caspase 1 |
| 1449591 | at | 1.49 | 0.000 | Casp11 | caspase 11, apoptosis-related cysteine protease |
| 1418981 | at | 1.80 | 0.003 | Casp12 | caspase 12 |
| 1448165 | at | 1.02 | 0.776 | Casp2 | caspase 2 |
| 1415995 | at | -1.24 | 0.020 | Casp6 | caspase 6 |
| 1448659 | at | -1.10 | 0.143 | Casp7 | caspase 7 |
| 1449217 | at | 1.07 | 0.412 | Casp8ap2 | caspase 8 associated protein 2 |
| 1437537 | at | -1.15 | 0.035 | Casp9 | caspase 9 |
| 1451413 | at | 1.17 | 0.210 | Cast | calpastatin |
| 1451980 | at | -1.08 | 0.345 | Cast1 | O-acetyltransferase |
| 1416430 | at | 1.28 | 0.000 | Cat | catalase |
| 1448149 | at | -1.02 | 0.554 | Catna1 | catenin alpha 1 |
| 1441193 | at | -2.13 | 0.000 | Catna3 | catenin alpha 3 |
| 1420929 | at | -1.47 | 0.060 | Catnal1 | catenin alpha-like 1 |
| 1430533 | a_at | 1.04 | 0.809 | Catnb | catenin beta |
| 1417567 | at | -1.01 | 0.923 | Catnbip1 | catenin beta interacting protein 1 |
| 1425309 | at | 1.10 | 0.142 | Catsper2 | cation channel, sperm associated 2 |
| 1449145 | a_at | 1.05 | 0.180 | Cav1 | caveolin, caveolae protein 1 |

Supplementary table S1

| | | | | | |
|---|---|---|---|---|---|
| 1417327 | at | 1.04 | 0.464 | Cav2 | caveolin 2 |
| 1418413 | at | 1.12 | 0.039 | Cav3 | caveolin 3 |
| 1440310 | at | -1.35 | 0.040 | Cbfa2t1h | CBFA2T1 identified gene homolog (human) |
| 1434757 | at | -1.07 | 0.359 | Cbfa2t2h | core-binding factor, runt domain, alpha subunit 2, translocated to, 2 homolog (human) |
| 1438705 | at | -1.15 | 0.082 | Cbfa2t3h | core-binding factor, runt domain, alpha subunit 2, translocated to, 3 homolog (human) |
| 1460716 | a_at | 1.02 | 0.767 | Cbfb | core binding factor beta |
| 1434829 | at | -1.06 | 0.383 | Cbl | Casitas B-lineage lymphoma |
| 1437203 | at | -1.10 | 0.309 | Cbll1 | Casitas B-lineage lymphoma-like 1 |
| 1424104 | at | 1.04 | 0.245 | Cbpin | CCNDBP1 interactor |
| 1460196 | at | -1.26 | 0.001 | Cbr1 | carbonyl reductase 1 |
| 1418509 | at | 1.33 | 0.037 | Cbr2 | carbonyl reductase 2 |
| 1451531 | at | 1.07 | 0.176 | Cbwd1 | COBW domain containing 1 |
| 1416884 | at | -1.06 | 0.098 | Cbx3 | chromobox homolog 3 (Drosophila HP1 gamma) |
| 1419583 | at | 1.25 | 0.011 | Cbx4 | chromobox homolog 4 (Drosophila Pc class) |
| 1454636 | at | 1.03 | 0.551 | Cbx5 | chromobox homolog 5 (Drosophila HP1a) |
| 1437385 | at | -2.08 | 0.000 | Ccbe1 | collagen and calcium binding EGF domains 1 |
| 1452678 | a_at | -1.20 | 0.005 | Ccbl1 | cysteine conjugate-beta lyase 1 |
| 1455991 | at | -1.56 | 0.000 | Ccbl2 | cysteine conjugate-beta lyase 2 |
| 1424955 | at | 1.05 | 0.455 | Ccdc5 | coiled-coil domain containing 5 |
| 1419282 | at | 2.19 | 0.004 | Ccl12 | chemokine (C-C motif) ligand 12 |
| 1430375 | a_at | -1.38 | 0.003 | Ccl27 | chemokine (C-C motif) ligand 27 |
| 1417266 | at | 1.28 | 0.037 | Ccl6 | chemokine (C-C motif) ligand 6 |
| 1417936 | at | 1.36 | 0.008 | Ccl9 | chemokine (C-C motif) ligand 9 |
| 1442125 | at | 1.15 | 0.397 | Ccm1 | cerebral cavernous malformations 1 |
| 1434649 | at | 1.11 | 0.377 | Ccm2 | cerebral cavernous malformation 2 homolog (human) |
| 1417420 | at | 1.50 | 0.094 | Ccnd1 | cyclin D1 |
| 1416124 | at | 1.38 | 0.068 | Ccnd2 | cyclin D2 |
| 1415907 | at | 1.11 | 0.102 | Ccnd3 | cyclin D3 |
| 1420745 | a_at | 1.15 | 0.034 | Ccndbp1 | cyclin D-type binding-protein 1 |
| 1450016 | at | 1.01 | 0.775 | Ccng1 | cyclin G1 |
| 1416488 | at | -1.04 | 0.640 | Ccng2 | cyclin G2 |
| 1418585 | at | -1.15 | 0.214 | Ccnh | cyclin H |
| 1416427 | at | -1.11 | 0.421 | Ccni | cyclin I |
| 1423622 | a_at | 1.06 | 0.195 | Ccnl1 | cyclin L1 |
| 1454149 | a_at | -1.04 | 0.539 | Ccnl2 | cyclin L2 |
| 1456477 | at | -1.15 | 0.243 | Ccnt1 | cyclin T1 |
| 1424420 | at | 1.07 | 0.032 | Ccpg1 | cell cycle progression 1 |
| 1425837 | a_at | -1.43 | 0.001 | Ccrn4l | CCR4 carbon catabolite repression 4-like (S. cerevisiae) |
| 1448615 | at | -1.01 | 0.878 | Ccs | copper chaperone for superoxide dismutase |
| 1448178 | a_at | 1.10 | 0.134 | Cct3 | chaperonin subunit 3 (gamma) |
| 1415867 | at | 1.04 | 0.351 | Cct4 | chaperonin subunit 4 (delta) |

Supplementary table S1

| | | | | | |
|---|---|---|---|---|---|
| 1417258 | at | 1.16 | 0.012 | Cct5 | chaperonin subunit 5 (epsilon) |
| 1423517 | at | 1.12 | 0.070 | Cct6a | chaperonin subunit 6a (zeta) |
| 1415816 | at | -1.08 | 0.400 | Cct7 | chaperonin subunit 7 (eta) |
| 1415785 | a_at | 1.17 | 0.000 | Cct8 | chaperonin subunit 8 (theta) |
| 1451232 | at | 1.13 | 0.002 | Cd151 | CD151 antigen |
| 1419144 | at | -1.19 | 0.153 | Cd163 | CD163 antigen |
| 1416440 | at | -1.08 | 0.018 | Cd164 | CD164 antigen |
| 1458082 | at | 1.09 | 0.667 | Cd200 | Cd200 antigen |
| 1417439 | at | 1.72 | 0.003 | Cd248 | CD248 antigen, endosialin |
| 1448182 | a_at | 1.13 | 0.187 | Cd24a | CD24a antigen |
| 1419714 | at | -1.17 | 0.295 | Cd274 | CD274 antigen |
| 1420907 | at | -1.01 | 0.753 | Cd2ap | CD2-associated protein |
| 1417223 | at | 1.13 | 0.006 | Cd2bp2 | CD2 antigen (cytoplasmic tail) binding protein 2 |
| 1448919 | at | 1.53 | 0.000 | Cd302 | CD302 antigen |
| 1417100 | at | -1.04 | 0.503 | Cd320 | CD320 antigen |
| 1416072 | at | 1.48 | 0.002 | Cd34 | CD34 antigen |
| 1450884 | at | -1.14 | 0.026 | Cd36 | CD36 antigen |
| 1433741 | at | 1.31 | 0.001 | Cd38 | CD38 antigen |
| 1423760 | at | 1.66 | 0.018 | Cd44 | CD44 antigen |
| 1419554 | at | -1.08 | 0.210 | Cd47 | CD47 antigen (Rh-related antigen, integrin-associated signal transducer) |
| 1460218 | at | 1.56 | 0.016 | Cd52 | CD52 antigen |
| 1448617 | at | 1.36 | 0.000 | Cd53 | CD53 antigen |
| 1418710 | at | -1.25 | 0.001 | Cd59a | CD59a antigen |
| 1449164 | at | 1.30 | 0.070 | Cd68 | CD68 antigen |
| 1457952 | at | -1.22 | 0.008 | Cd80 | CD80 antigen |
| 1416330 | at | -1.17 | 0.007 | Cd81 | CD 81 antigen |
| 1416111 | at | 1.24 | 0.028 | Cd83 | CD83 antigen |
| 1422875 | at | 1.18 | 0.504 | Cd84 | CD84 antigen |
| 1416066 | at | 1.20 | 0.001 | Cd9 | CD9 antigen |
| 1418394 | a_at | 1.10 | 0.324 | Cd97 | CD97 antigen |
| 1423965 | at | -1.10 | 0.230 | Cd99l2 | Cd99 antigen-like 2 |
| 1428166 | at | -1.22 | 0.001 | Cdan1 | congenital dyserythropoietic anemia, type I (human) |
| 1437070 | at | 1.07 | 0.314 | Cdc14b | CDC14 cell division cycle 14 homolog B (S. cerevisiae) |
| 1425554 | a_at | 1.11 | 0.090 | Cdc16 | CDC16 cell division cycle 16 homolog (S. cerevisiae) |
| 1421963 | a_at | 1.20 | 0.031 | Cdc25b | cell division cycle 25 homolog B (S. cerevisiae) |
| 1423987 | at | 1.04 | 0.404 | Cdc26 | cell division cycle 26 |
| 1458282 | at | -1.17 | 0.333 | Cdc27 | cell division cycle 27 homolog (S. cerevisiae) |
| 1440833 | at | -1.02 | 0.813 | Cdc2l5 | cell division cycle 2-like 5 (cholinesterase-related cell division controller) |
| 1455031 | at | -1.01 | 0.877 | Cdc2l6 | cell division cycle 2-like 6 (CDK8-like) |
| 1434879 | at | -1.17 | 0.007 | Cdc34 | cell division cycle 34 homolog (S. cerevisiae) |
| 1416819 | at | -1.05 | 0.492 | Cdc37 | cell division cycle 37 homolog (S. cerevisiae) |

Supplementary table S1

| | | | | | |
|---|---|---|---|---|---|
| 1453342 | at | -1.13 | 0.032 | Cdc40 | cell division cycle 40 homolog (yeast) |
| 1428750 | at | 1.04 | 0.775 | Cdc42ep2 | CDC42 effector protein (Rho GTPase binding) 2 |
| 1450700 | at | -1.13 | 0.204 | Cdc42ep3 | CDC42 effector protein (Rho GTPase binding) 3 |
| 1416511 | a_at | -1.02 | 0.866 | Cdc42ep4 | CDC42 effector protein (Rho GTPase binding) 4 |
| 1428131 | a_at | 1.32 | 0.007 | Cdc42se1 | CDC42 small effector 1 |
| 1423683 | at | 1.22 | 0.355 | Cdca4 | cell division cycle associated 4 |
| 1455164 | at | -1.03 | 0.460 | Cdgap | Cdc42 GTPase-activating protein |
| 1433956 | at | 1.01 | 0.841 | Cdh5 | cadherin 5 |
| 1423657 | at | 1.07 | 0.467 | Cdipt | CDP-diacylglycerol--inositol 3-phosphatidyltransferase (phosphatidylinositol synthase) |
| 1433975 | at | 1.04 | 0.689 | Cdk10 | cyclin-dependent kinase (CDC2-like) 10 |
| 1426885 | a_at | 1.01 | 0.689 | Cdk2ap1 | CDK2 (cyclin-dependent kinase 2)-associated protein 1 |
| 1417161 | at | -1.25 | 0.011 | Cdk2ap2 | CDK2-associated protein 2 |
| 1450674 | at | -1.17 | 0.025 | Cdk5 | cyclin-dependent kinase 5 |
| 1448626 | at | -1.14 | 0.134 | Cdk5rap1 | CDK5 regulatory subunit associated protein 1 |
| 1423067 | at | 1.05 | 0.523 | Cdk5rap3 | CDK5 regulatory subunit associated protein 3 |
| 1435338 | at | 1.43 | 0.012 | Cdk6 | cyclin-dependent kinase 6 |
| 1435299 | at | -1.23 | 0.015 | Cdk7 | cyclin-dependent kinase 7 (homolog of Xenopus MO15 cdk-activating kinase) |
| 1460389 | at | 1.14 | 0.045 | Cdk8 | cyclin-dependent kinase 8 |
| 1417269 | at | 1.17 | 0.044 | Cdk9 | cyclin-dependent kinase 9 (CDC2-related kinase) |
| 1423953 | at | -1.26 | 0.163 | Cdkal1 | CDK5 regulatory subunit associated protein 1-like 1 |
| 1455528 | at | 1.11 | 0.345 | Cdkl1 | cyclin-dependent kinase-like 1 (CDC2-related kinase) |
| 1449229 | a_at | 1.59 | 0.001 | Cdkl2 | cyclin-dependent kinase-like 2 (CDC2-related kinase) |
| 1424638 | at | 1.27 | 0.110 | Cdkn1a | cyclin-dependent kinase inhibitor 1A (P21) |
| 1434045 | at | -1.34 | 0.072 | Cdkn1b | cyclin-dependent kinase inhibitor 1B (P27) |
| 1416868 | at | -1.24 | 0.003 | Cdkn2c | cyclin-dependent kinase inhibitor 2C (p18, inhibits CDK4) |
| 1416253 | at | 1.14 | 0.081 | Cdkn2d | cyclin-dependent kinase inhibitor 2D (p19, inhibits CDK4) |
| 1417430 | at | 1.67 | 0.023 | Cdr2 | cerebellar degeneration-related 2 |
| 1449407 | at | -2.28 | 0.000 | Cdv1 | carnitine deficiency-associated gene expressed in ventricle 1 |
| 1415704 | a_at | 1.33 | 0.001 | Cdv3 | carnitine deficiency-associated gene expressed in ventricle 3 |
| 1423865 | at | 1.01 | 0.862 | Cdw92 | CDW92 antigen |
| 1435374 | at | -1.14 | 0.003 | Cdyl2 | chromodomain protein, Y chromosome-like 2 |
| 1460681 | at | -1.00 | 0.984 | Ceacam2 | CEA-related cell adhesion molecule 2 |
| 1418982 | at | 1.17 | 0.161 | Cebpa | CCAAT/enhancer binding protein (C/EBP), alpha |
| 1456605 | at | 1.47 | 0.018 | Cebpd | CCAAT/enhancer binding protein (C/EBP), delta |
| 1451639 | at | 1.28 | 0.007 | Cebpg | CCAAT/enhancer binding protein (C/EBP), gamma |
| 1420497 | a_at | -1.15 | 0.016 | Cebpz | CCAAT/enhancer binding protein zeta |
| 1435336 | at | -1.78 | 0.000 | Celsr2 | cadherin EGF LAG seven-pass G-type receptor 2 |
| 1450842 | a_at | 1.21 | 0.006 | Cenpa | centromere autoantigen A |
| 1420441 | at | 1.15 | 0.077 | Cenpc1 | centromere autoantigen C1 |
| 1427161 | at | -3.59 | 0.000 | Cenpf | centromere autoantigen F |
| 1425639 | at | -1.25 | 0.063 | Centa2 | centaurin, alpha 2 |

Supplementary table S1

| | | | | | |
|---|---|---|---|---|---|
| 1433561 | at | 1.15 | 0.054 | Centb2 | centaurin, beta 2 |
| 1452291 | at | -1.11 | 0.430 | Centd1 | centaurin, delta 1 |
| 1451282 | at | 1.03 | 0.737 | Centd3 | centaurin, delta 3 |
| 1435432 | at | 1.14 | 0.025 | Centg2 | centaurin, gamma 2 |
| 1423910 | at | 1.04 | 0.657 | Centg3 | centaurin, gamma 3 |
| 1421005 | at | -1.11 | 0.278 | Cep1 | centrosomal protein 1 |
| 1416471 | at | 1.04 | 0.392 | Cept1 | choline/ethanolaminephosphotransferase 1 |
| 1434034 | at | 1.06 | 0.419 | Cerk | ceramide kinase |
| 1449081 | at | -3.30 | 0.000 | Ces3 | carboxylesterase 3 |
| 1418579 | at | 1.03 | 0.592 | Cetn2 | centrin 2 |
| 1417239 | at | -1.07 | 0.065 | Cetn3 | centrin 3 |
| 1450876 | at | 1.88 | 0.000 | Cfh | complement component factor h |
| 1448346 | at | -1.01 | 0.821 | Cfl1 | cofilin 1, non-muscle |
| 1418066 | at | 1.14 | 0.069 | Cfl2 | cofilin 2, muscle |
| 1425686 | at | -1.16 | 0.235 | Cflar | CASP8 and FADD-like apoptosis regulator |
| 1454641 | at | 1.03 | 0.389 | Cggbp1 | CGG triplet repeat binding protein 1 |
| 1452309 | at | -1.17 | 0.079 | Cgnl1 | cingulin-like 1 |
| 1434565 | at | -1.06 | 0.424 | Cgrrf1 | cell growth regulator with ring finger domain 1 |
| 1416962 | at | -1.07 | 0.538 | Chc1 | chromosome condensation 1 |
| 1416389 | a_at | -1.05 | 0.219 | Chc1l | chromosome condensation 1-like |
| 1416938 | at | -1.02 | 0.718 | Chchd1 | coiled-coil-helix-coiled-coil-helix domain containing 1 |
| 1428161 | a_at | 1.05 | 0.232 | Chchd2 | coiled-coil-helix-coiled-coil-helix domain containing 2 |
| 1417233 | at | 1.41 | 0.000 | Chchd4 | coiled-coil-helix-coiled-coil-helix domain containing 4 |
| 1451505 | at | 1.03 | 0.621 | Chchd5 | coiled-coil-helix-coiled-coil-helix domain containing 5 |
| 1423779 | at | 1.29 | 0.070 | Chchd6 | coiled-coil-helix-coiled-coil-helix domain containing 6 |
| 1446844 | at | -1.39 | 0.007 | Chchd7 | coiled-coil-helix-coiled-coil-helix domain containing 7 |
| 1457473 | at | 1.04 | 0.713 | Chd1 | chromodomain helicase DNA binding protein 1 |
| 1449415 | at | 1.00 | 0.979 | Chd1l | chromodomain helicase DNA binding protein 1-like |
| 1438476 | a_at | -1.07 | 0.401 | Chd4 | chromodomain helicase DNA binding protein 4 |
| 1437297 | at | -1.03 | 0.645 | Chd8 | chromodomain helicase DNA binding protein 8 |
| 1451896 | a_at | 1.07 | 0.269 | Cherp | calcium homeostasis endoplasmic reticulum protein |
| 1436925 | at | -1.19 | 0.872 | Ches1 | checkpoint supressor 1 |
| 1433550 | at | 1.26 | 0.008 | Chfr | checkpoint with forkhead and ring finger domains |
| 1444018 | at | 1.09 | 0.358 | Chic2 | cysteine-rich hydrophobic domain 2 |
| 1448448 | a_at | -1.19 | 0.019 | Chkb | choline kinase beta |
| 1447156 | at | 1.19 | 0.143 | Chm | choroidermia |
| 1435926 | at | -1.08 | 0.286 | Chml | choroideremia-like |
| 1418817 | at | 1.23 | 0.010 | Chmp1b | chromatin modifying protein 1B |
| 1425591 | a_at | 1.00 | 0.964 | Chmp2a | chromatin modifying protein 2A |
| 1428253 | at | 1.05 | 0.242 | Chmp2b | chromatin modifying protein 2B |
| 1426240 | at | 1.16 | 0.036 | Chmp4b | chromatin modifying protein 4B |

Supplementary table S1

| | | | | | |
|---|---|---|---|---|---|
| 1432270 | a_at | 1.02 | 0.595 | Chmp5 | chromatin modifying protein 5 |
| 1451300 | a_at | -1.03 | 0.431 | Chmp7 | CHMP family, member 7 |
| 1420545 | a_at | -1.08 | 0.190 | Chn1 | chimerin (chimaerin) 1 |
| 1435574 | at | -1.07 | 0.278 | Chordc1 | cysteine and histidine-rich domain (CHORD)-containing, zinc-binding protein 1 |
| 1422505 | at | -1.04 | 0.309 | Chrac1 | chromatin accessibility complex 1 |
| 1420682 | at | 1.33 | 0.029 | Chrnb1 | cholinergic receptor, nicotinic, beta polypeptide 1 (muscle) |
| 1449147 | at | 1.06 | 0.543 | Chst1 | carbohydrate (keratan sulfate Gal-6) sulfotransferase 1 |
| 1417091 | at | 1.00 | 0.948 | Chuk | conserved helix-loop-helix ubiquitous kinase |
| 1448140 | at | -1.20 | 0.175 | Ciapin1 | cytokine induced apoptosis inhibitor 1 |
| 1448564 | at | 1.26 | 0.094 | Cib1 | calcium and integrin binding 1 (calmyrin) |
| 1415746 | at | -1.06 | 0.331 | Cic | capicua homolog (Drosophila) |
| 1417956 | at | 1.27 | 0.008 | Cidea | cell death-inducing DNA fragmentation factor, alpha subunit-like effector A |
| 1416332 | at | 1.36 | 0.094 | Cirbp | cold inducible RNA binding protein |
| 1448724 | at | -1.23 | 0.026 | Cish | cytokine inducible SH2-containing protein |
| 1452207 | at | -1.37 | 0.038 | Cited2 | Cbp/p300-interacting transactivator, with Glu/Asp-rich carboxy-terminal domain, 2 |
| 1448779 | at | -1.13 | 0.018 | Ciz1 | CDKN1A interacting zinc finger protein 1 |
| 1448533 | at | 1.06 | 0.114 | Ckap1 | cytoskeleton-associated protein 1 |
| 1452181 | at | 1.42 | 0.004 | Ckap4 | cytoskeleton-associated protein 4 |
| 1455106 | a_at | 1.12 | 0.079 | Ckb | creatine kinase, brain |
| 1448316 | at | 1.17 | 0.054 | Cklfsf3 | chemokine-like factor super family 3 |
| 1434337 | at | -1.18 | 0.071 | Cklfsf4 | chemokine-like factor super family 4 |
| 1451114 | at | 1.30 | 0.008 | Cklfsf6 | chemokine-like factor super family 6 |
| 1460253 | at | 1.01 | 0.841 | Cklfsf7 | chemokine-like factor super family 7 |
| 1427964 | at | -1.71 | 0.000 | Cklfsf8 | chemokine-like factor super family 8 |
| 1428722 | at | 1.02 | 0.437 | Ckmt2 | creatine kinase, mitochondrial 2 |
| 1416698 | a_at | 1.21 | 0.005 | Cks1b | CDC28 protein kinase 1b |
| 1417457 | at | 1.87 | 0.000 | Cks2 | CDC28 protein kinase regulatory subunit 2 |
| 1452265 | at | -1.37 | 0.078 | Clasp1 | CLIP associating protein 1 |
| 1424836 | a_at | -1.13 | 0.160 | Clasp2 | CLIP associating protein 2 |
| 1441248 | at | -1.39 | 0.001 | Clcn3 | chloride channel 3 |
| 1448161 | a_at | -1.23 | 0.001 | Clcn4-2 | chloride channel 4-2 |
| 1429400 | at | 1.15 | 0.075 | Clcn5 | chloride channel 5 |
| 1455440 | at | -1.09 | 0.378 | Clcn6 | chloride channel 6 |
| 1452702 | at | -1.05 | 0.601 | Clcn7 | chloride channel 7 |
| 1433782 | at | -1.10 | 0.104 | Cldn12 | claudin 12 |
| 1417839 | at | 1.33 | 0.027 | Cldn5 | claudin 5 |
| 1456318 | at | 1.03 | 0.755 | Clec1a | C-type lectin domain family 1, member a |
| 1419477 | at | 1.14 | 0.224 | Clec2d | C-type lectin domain family 2, member d |
| 1416656 | at | 1.30 | 0.036 | Clic1 | chloride intracellular channel 1 |
| 1439505 | at | 1.70 | 0.002 | Clic5 | chloride intracellular channel 5 |
| 1426124 | a_at | -1.09 | 0.201 | Clk1 | CDC-like kinase 1 |

Supplementary table S1

| | | | | | |
|---|---|---|---|---|---|
| 1417743 | at | 1.09 | 0.357 | Clk2 | CDC-like kinase 2 |
| 1423849 | a_at | -1.06 | 0.450 | Clk3 | CDC-like kinase 3 |
| 1422886 | a_at | -1.20 | 0.099 | Clk4 | CDC like kinase 4 |
| 1434768 | at | -1.06 | 0.394 | Cln2 | ceroid-lipofuscinosis, neuronal 2 |
| 1454837 | at | -1.15 | 0.125 | Cln6 | ceroid-lipofuscinosis, neuronal 6 |
| 1427548 | a_at | 1.10 | 0.492 | Clns1a | chloride channel, nucleotide-sensitive, 1A |
| 1418660 | at | -1.14 | 0.244 | Clock | circadian locomoter output cycles kaput |
| 1425937 | a_at | 1.10 | 0.181 | Clp1 | cardiac lineage protein 1 |
| 1416615 | at | -1.29 | 0.000 | Clpp | caseinolytic protease, ATP-dependent, proteolytic subunit homolog (E. coli) |
| 1416883 | at | 1.06 | 0.232 | Clptm1 | cleft lip and palate associated transmembrane protein 1 |
| 1439342 | at | -1.24 | 0.019 | Clpx | caseinolytic protease X (E.coli) |
| 1421861 | at | -1.22 | 0.003 | Clstn1 | calsyntenin 1 |
| 1434540 | a_at | -1.04 | 0.373 | Clta | clathrin, light polypeptide (Lca) |
| 1453063 | at | 1.30 | 0.033 | Cltb | clathrin, light polypeptide (Lcb) |
| 1454626 | at | 1.07 | 0.047 | Cltc | clathrin, heavy polypeptide (Hc) |
| 1418626 | a_at | 1.12 | 0.008 | Clu | clusterin |
| 1424388 | at | -1.01 | 0.860 | Cluap1 | clusterin associated protein 1 |
| 1421014 | a_at | -1.26 | 0.001 | Clybl | citrate lyase beta like |
| 1426662 | at | 1.14 | 0.010 | Cmas | cytidine monophospho-N-acetylneuraminic acid synthetase |
| 1419220 | at | 1.72 | 0.000 | Cmya1 | cardiomyopathy associated 1 |
| 1436939 | at | -1.01 | 0.809 | Cmya4 | cardiomyopathy associated 4 |
| 1427312 | at | -1.11 | 0.014 | Cmya5 | cardiomyopathy associated 5 |
| 1460177 | at | 1.19 | 0.046 | Cndp2 | CNDP dipeptidase 2 (metallopeptidase M20 family) |
| 1423727 | at | 1.08 | 0.094 | Cnih | cornichon homolog (Drosophila) |
| 1433983 | at | 1.26 | 0.040 | Cnksr3 | Cnksr family member 3 |
| 1450981 | at | 1.14 | 0.022 | Cnn2 | calponin 2 |
| 1426724 | at | 1.39 | 0.001 | Cnn3 | calponin 3, acidic |
| 1420481 | at | -1.25 | 0.011 | Cnnm3 | cyclin M3 |
| 1434251 | at | -1.14 | 0.003 | Cnot1 | CCR4-NOT transcription complex, subunit 1 |
| 1433882 | at | -1.10 | 0.431 | Cnot10 | CCR4-NOT transcription complex, subunit 10 |
| 1454603 | a_at | 1.01 | 0.822 | Cnot2 | CCR4-NOT transcription complex, subunit 2 |
| 1421000 | at | -1.18 | 0.062 | Cnot4 | CCR4-NOT transcription complex, subunit 4 |
| 1426684 | at | 1.06 | 0.589 | Cnot6 | CCR4-NOT transcription complex, subunit 6 |
| 1425481 | at | -1.17 | 0.107 | Cnot6l | CCR4-NOT transcription complex, subunit 6-like |
| 1460665 | a_at | 1.06 | 0.470 | Cnot7 | CCR4-NOT transcription complex, subunit 7 |
| 1451102 | at | -1.08 | 0.017 | Cnot8 | CCR4-NOT transcription complex, subunit 8 |
| 1426326 | at | 1.13 | 0.043 | Cntf | ciliary neurotrophic factor |
| 1423701 | at | -1.11 | 0.150 | Coasy | Coenzyme A synthase |
| 1454795 | at | -1.54 | 0.192 | Cobll1 | Cobl-like 1 |
| 1416932 | at | 1.05 | 0.531 | Cog2 | component of oligomeric golgi complex 2 |
| 1434654 | at | -1.09 | 0.092 | Cog3 | component of oligomeric golgi complex 3 |

Supplementary table S1

| | | | | | |
|---|---|---|---|---|---|
| 1416993 | at | 1.12 | 0.070 | Cog4 | component of oligomeric golgi complex 4 |
| 1426216 | at | 1.00 | 0.982 | Cog6 | component of oligomeric golgi complex 6 |
| 1448755 | at | 1.72 | 0.000 | Col15a1 | procollagen, type XV |
| 1423669 | at | 1.89 | 0.000 | Col1a1 | procollagen, type I, alpha 1 |
| 1423110 | at | 2.36 | 0.000 | Col1a2 | procollagen, type I, alpha 2 |
| 1442977 | at | 1.65 | 0.010 | Col3a1 | procollagen, type III, alpha 1 |
| 1452035 | at | 1.44 | 0.001 | Col4a1 | procollagen, type IV, alpha 1 |
| 1424051 | at | 1.38 | 0.023 | Col4a2 | procollagen, type IV, alpha 2 |
| 1425476 | at | 1.31 | 0.008 | Col4a5 | procollagen, type IV, alpha 5 |
| 1434479 | at | 1.43 | 0.066 | Col5a1 | procollagen, type V, alpha 1 |
| 1422437 | at | 2.51 | 0.000 | Col5a2 | procollagen, type V, alpha 2 |
| 1419703 | at | -1.06 | 0.555 | Col5a3 | procollagen, type V, alpha 3 |
| 1448590 | at | 1.40 | 0.001 | Col6a1 | procollagen, type VI, alpha 1 |
| 1452250 | a_at | 1.48 | 0.004 | Col6a2 | procollagen, type VI, alpha 2 |
| 1424131 | at | 1.35 | 0.001 | Col6a3 | procollagen, type VI, alpha 3 |
| 1455627 | at | 6.02 | 0.000 | Col8a1 | procollagen, type VIII, alpha 1 |
| 1429179 | at | -1.17 | 0.133 | Colec11 | collectin sub-family member 11 |
| 1424121 | at | 1.01 | 0.781 | Commd1 | COMM domain containing 1 |
| 1428491 | at | 1.17 | 0.030 | Commd10 | COMM domain containing 10 |
| 1433594 | at | 1.01 | 0.872 | Commd2 | COMM domain containing 2 |
| 1433539 | at | 1.19 | 0.075 | Commd3 | COMM domain containing 3 |
| 1417006 | at | 1.11 | 0.258 | Commd4 | COMM domain containing 4 |
| 1419989 | at | -1.02 | 0.722 | Commd5 | COMM domain containing 5 |
| 1435863 | at | 1.04 | 0.524 | Commd6 | COMM domain containing 6 |
| 1426765 | at | 1.25 | 0.045 | Commd7 | COMM domain containing 7 |
| 1455671 | at | 1.05 | 0.499 | Commd8 | COMM domain containing 8 |
| 1428155 | at | 1.06 | 0.438 | Commd9 | COMM domain containing 9 |
| 1418701 | at | 1.52 | 0.001 | Comt | catechol-O-methyltransferase |
| 1428635 | at | 1.83 | 0.003 | Comtd1 | catechol-O-methyltransferase domain containing 1 |
| 1423301 | at | -1.03 | 0.541 | Copb1 | coatomer protein complex, subunit beta 1 |
| 1416384 | a_at | -1.12 | 0.077 | Cope | coatomer protein complex, subunit epsilon |
| 1416017 | at | 1.00 | 0.996 | Copg | coatomer protein complex, subunit gamma |
| 1448761 | a_at | 1.11 | 0.021 | Copg2 | coatomer protein complex, subunit gamma 2 |
| 1416678 | at | -1.07 | 0.190 | Cops3 | COP9 (constitutive photomorphogenic) homolog, subunit 3 (Arabidopsis thaliana) |
| 1416163 | at | 1.08 | 0.098 | Cops4 | COP9 (constitutive photomorphogenic) homolog, subunit 4 (Arabidopsis thaliana) |
| 1460171 | at | 1.02 | 0.522 | Cops5 | COP9 (constitutive photomorphogenic) homolog, subunit 5 (Arabidopsis thaliana) |
| 1451366 | at | -1.04 | 0.218 | Cops6 | COP9 (constitutive photomorphogenic) homolog, subunit 6 (Arabidopsis thaliana) |
| 1418721 | at | 1.03 | 0.796 | Cops7b | COP9 (constitutive photomorphogenic) homolog, subunit 7b (Arabidopsis thaliana) |
| 1415934 | at | 1.14 | 0.046 | Cops8 | COP9 (constitutive photomorphogenic) homolog, subunit 8 (Arabidopsis thaliana) |
| 1451825 | a_at | 1.00 | 0.992 | Copz1 | coatomer protein complex, subunit zeta 1 |
| 1418455 | at | -1.12 | 0.225 | Copz2 | coatomer protein complex, subunit zeta 2 |

Supplementary table S1

| | | | | | |
|---|---|---|---|---|---|
| 1436351 | at | -1.11 | 0.059 | Coq3 | coenzyme Q3 homolog, methyltransferase (yeast) |
| 1435231 | at | 1.07 | 0.528 | Coq4 | coenzyme Q4 homolog (yeast) |
| 1426902 | at | -1.25 | 0.067 | Coq6 | coenzyme Q6 homolog (yeast) |
| 1416665 | at | -1.10 | 0.007 | Coq7 | demethyl-Q 7 |
| 1419017 | at | -1.30 | 0.001 | Corin | corin |
| 1455269 | a_at | 1.35 | 0.024 | Coro1a | coronin, actin binding protein 1A |
| 1448351 | at | 1.07 | 0.558 | Coro1b | coronin, actin binding protein 1B |
| 1419911 | at | 1.12 | 0.017 | Coro1c | coronin, actin binding protein 1C |
| 1443513 | at | -1.10 | 0.655 | Cox11 | COX11 homolog, cytochrome c oxidase assembly protein (yeast) |
| 1452146 | a_at | -1.15 | 0.210 | Cox15 | COX15 homolog, cytochrome c oxidase assembly protein (yeast) |
| 1448322 | a_at | -1.03 | 0.139 | Cox4i1 | cytochrome c oxidase subunit IV isoform 1 |
| 1421373 | at | -1.27 | 0.225 | Cox4i2 | cytochrome c oxidase subunit IV isoform 2 |
| 1416902 | a_at | -1.09 | 0.000 | Cox5b | cytochrome c oxidase, subunit Vb |
| 1417417 | a_at | 1.02 | 0.424 | Cox6a1 | cytochrome c oxidase, subunit VI a, polypeptide 1 |
| 1417607 | at | -1.02 | 0.574 | Cox6a2 | cytochrome c oxidase, subunit VI a, polypeptide 2 |
| 1435275 | at | -2.44 | 0.000 | Cox6b2 | cytochrome c oxidase subunit VIb polypeptide 2 |
| 1418709 | at | -1.03 | 0.275 | Cox7a1 | cytochrome c oxidase, subunit VIIa 1 |
| 1448112 | at | -1.18 | 0.002 | Cox7c | cytochrome c oxidase, subunit VIIc |
| 1416112 | at | -1.03 | 0.477 | Cox8a | cytochrome c oxidase, subunit VIIIa |
| 1449218 | at | -1.20 | 0.000 | Cox8b | cytochrome c oxidase, subunit VIIIb |
| 1448735 | at | 1.18 | 0.173 | Cp | ceruloplasmin |
| 1459488 | at | -1.08 | 0.682 | Cpb2 | carboxypeptidase B2 (plasma) |
| 1434547 | at | 1.19 | 0.063 | Cpd | carboxypeptidase D |
| 1415949 | at | 1.05 | 0.337 | Cpe | carboxypeptidase E |
| 1456048 | at | -1.61 | 0.001 | Cpeb3 | cytoplasmic polyadenylation element binding protein 3 |
| 1420618 | at | 1.10 | 0.968 | Cpeb4 | cytoplasmic polyadenylation element binding protein 4 |
| 1453009 | at | -1.01 | 0.900 | Cpm | carboxypeptidase M |
| 1433439 | at | -1.01 | 0.911 | Cpne1 | copine I |
| 1435450 | at | -1.29 | 0.542 | Cpne3 | copine III |
| 1431146 | a_at | 1.36 | 0.002 | Cpne8 | copine VIII |
| 1422492 | at | 1.23 | 0.061 | Cpox | coproporphyrinogen oxidase |
| 1417665 | a_at | 1.04 | 0.752 | Cpsf1 | cleavage and polyadenylation specific factor 1 |
| 1422855 | at | 1.04 | 0.474 | Cpsf3 | cleavage and polyadenylation specificity factor 3 |
| 1424469 | a_at | -1.11 | 0.033 | Cpsf4 | cleavage and polyadenylation specific factor 4 |
| 1417681 | at | 1.01 | 0.619 | Cpsf5 | cleavage and polyadenylation specific factor 5 |
| 1460409 | at | -1.03 | 0.659 | Cpt1a | carnitine palmitoyltransferase 1a, liver |
| 1418328 | at | -1.18 | 0.014 | Cpt1b | carnitine palmitoyltransferase 1b, muscle |
| 1416772 | at | -1.23 | 0.002 | Cpt2 | carnitine palmitoyltransferase 2 |
| 1460248 | at | 1.90 | 0.031 | Cpxm2 | carboxypeptidase X 2 (M14 family) |
| 1443430 | at | -1.19 | 0.180 | Cradd | CASP2 and RIPK1 domain containing adaptor with death domain |
| 1417008 | at | -1.33 | 0.001 | Crat | carnitine acetyltransferase |

Supplementary table S1

| | | | | | |
|---|---|---|---|---|---|
| 1418249 | at | 1.10 | 0.148 | Crcp | calcitonin gene-related peptide-receptor component protein |
| 1419980 | at | 1.26 | 0.005 | Creb3 | cAMP responsive element binding protein 3 |
| 1452381 | at | 1.69 | 0.000 | Creb3l2 | cAMP responsive element binding protein 3-like 2 |
| 1434633 | at | -1.24 | 0.137 | Crebbp | CREB binding protein |
| 1416537 | at | 1.19 | 0.057 | Creld1 | cysteine-rich with EGF-like domains 1 |
| 1418322 | at | 1.41 | 0.033 | Crem | cAMP responsive element modulator |
| 1451355 | at | 1.11 | 0.423 | CRG-L1 | cancer related gene-liver 1 |
| 1422012 | at | -1.64 | 0.001 | Crhr2 | corticotropin releasing hormone receptor 2 |
| 1448406 | at | 1.16 | 0.336 | Cri1 | CREBBP/EP300 inhibitory protein 1 |
| 1454931 | at | -1.09 | 0.160 | Cri2 | CREBBP/EP300 inhibitory protein 2 |
| 1416326 | at | 1.19 | 0.063 | Crip1 | cysteine-rich protein 1 (intestinal) |
| 1417311 | at | -1.17 | 0.000 | Crip2 | cysteine rich protein 2 |
| 1434758 | at | 1.38 | 0.005 | Crispld2 | cysteine-rich secretory protein LCCL domain containing 2 |
| 1460176 | at | 1.02 | 0.816 | Crk | v-crk sarcoma virus CT10 oncogene homolog (avian) |
| 1455964 | at | -1.10 | 0.299 | Crk7 | CDC2-related kinase 7 |
| 1436950 | at | -1.14 | 0.298 | Crkl | v-crk sarcoma virus CT10 oncogene homolog (avian)-like |
| 1460338 | a_at | -1.08 | 0.270 | Crlf3 | cytokine receptor-like factor 3 |
| 1417583 | a_at | 1.01 | 0.862 | Crlz1 | charged amino acid rich leucine zipper 1 |
| 1420850 | at | -1.07 | 0.262 | Crnkl1 | Crn, crooked neck-like 1 (Drosophila) |
| 1450966 | at | 1.08 | 0.180 | Crot | carnitine O-octanoyltransferase |
| 1422563 | at | -1.02 | 0.514 | Crry | complement receptor related protein |
| 1437632 | at | 1.09 | 0.432 | Crsp2 | cofactor required for Sp1 transcriptional activation, subunit 2 |
| 1452064 | at | -1.27 | 0.001 | Crsp3 | cofactor required for Sp1 transcriptional activation, subunit 3 |
| 1425473 | at | -1.00 | 0.990 | Crsp6 | cofactor required for Sp1 transcriptional activation, subunit 6 |
| 1452282 | at | -1.17 | 0.074 | Crsp7 | cofactor required for Sp1 transcriptional activation, subunit 7 |
| 1417564 | at | 1.09 | 0.186 | Crsp9 | cofactor required for Sp1 transcriptional activation, subunit 9 |
| 1448592 | at | 1.02 | 0.661 | Crtap | cartilage associated protein |
| 1433733 | a_at | 1.05 | 0.607 | Cry1 | cryptochrome 1 (photolyase-like) |
| 1426383 | at | -1.11 | 0.365 | Cry2 | cryptochrome 2 (photolyase-like) |
| 1434369 | a_at | 1.11 | 0.010 | Cryab | crystallin, alpha B |
| 1420686 | at | -2.22 | 0.000 | Cryba4 | crystallin, beta A4 |
| 1451848 | a_at | -1.39 | 0.001 | Cryz | crystallin, zeta |
| 1430546 | at | -1.21 | 0.021 | Cryzl1 | crystallin, zeta (quinone reductase)-like 1 |
| 1422578 | at | -1.05 | 0.059 | Cs | citrate synthase |
| 1451012 | a_at | 1.01 | 0.928 | Csda | cold shock domain protein A |
| 1448809 | at | -1.07 | 0.087 | Cse1l | chromosome segregation 1-like (S. cerevisiae) |
| 1460220 | a_at | 1.23 | 0.027 | Csf1 | colony stimulating factor 1 (macrophage) |
| 1419872 | at | 1.22 | 0.019 | Csf1r | colony stimulating factor 1 receptor |
| 1438694 | at | 1.44 | 0.059 | Csng | casein gamma |
| 1449932 | at | 1.31 | 0.324 | Csnk1d | casein kinase 1, delta |
| 1428806 | at | 1.03 | 0.600 | Csnk1g1 | casein kinase 1, gamma 1 |

Supplementary table S1

| | | | | | |
|---|---|---|---|---|---|
| 1423370 | a_at | -1.03 | 0.483 | Csnk1g2 | casein kinase 1, gamma 2 |
| 1429383 | at | 1.23 | 0.756 | Csnk1g3 | casein kinase 1, gamma 3 |
| 1419036 | at | -1.09 | 0.696 | Csnk2a1 | casein kinase II, alpha 1 polypeptide |
| 1460646 | at | -1.09 | 0.298 | Csnk2a2 | casein kinase II, alpha 2, polypeptide |
| 1416728 | at | -1.15 | 0.010 | Csnk2b | casein kinase II, beta subunit |
| 1427256 | at | 2.82 | 0.000 | Cspg2 | chondroitin sulfate proteoglycan 2 |
| 1450950 | at | -1.09 | 0.096 | Cspg6 | chondroitin sulfate proteoglycan 6 |
| 1435792 | at | 1.06 | 0.564 | Csprs | component of Sp100-rs |
| 1425811 | a_at | 1.17 | 0.019 | Csrp1 | cysteine and glycine-rich protein 1 |
| 1420731 | a_at | 1.83 | 0.000 | Csrp2 | cysteine and glycine-rich protein 2 |
| 1452775 | at | -1.16 | 0.022 | Csrp2bp | cysteine and glycine-rich protein 2 binding protein |
| 1460318 | at | 1.07 | 0.007 | Csrp3 | cysteine and glycine-rich protein 3 |
| 1426195 | a_at | 1.05 | 0.030 | Cst3 | cystatin C |
| 1427910 | at | 1.67 | 0.010 | Cst6 | cystatin E/M |
| 1422507 | at | 1.03 | 0.563 | Cstb | cystatin B |
| 1448597 | at | 1.18 | 0.284 | Cstf1 | cleavage stimulation factor, 3' pre-RNA, subunit 1 |
| 1419645 | at | 1.08 | 0.241 | Cstf2 | cleavage stimulation factor, 3' pre-RNA subunit 2 |
| 1415920 | at | -1.07 | 0.149 | Cstf2t | cleavage stimulation factor, 3' pre-RNA subunit 2, tau |
| 1415702 | a_at | -1.04 | 0.515 | Ctbp1 | C-terminal binding protein 1 |
| 1434705 | at | 1.06 | 0.495 | Ctbp2 | C-terminal binding protein 2 |
| 1449042 | at | -1.18 | 0.029 | Ctcf | CCCTC-binding factor |
| 1452055 | at | -1.09 | 0.023 | Ctdsp1 | CTD (carboxy-terminal domain, RNA polymerase II, polypeptide A) small phosphatase 1 |
| 1422510 | at | 1.31 | 0.021 | Ctdspl | CTD (carboxy-terminal domain, RNA polymerase II, polypeptide A) small phosphatase-like |
| 1449065 | at | -3.95 | 0.000 | Cte1 | cytosolic acyl-CoA thioesterase 1 |
| 1449093 | at | -1.64 | 0.007 | Ctf1 | cardiotrophin 1 |
| 1416953 | at | 2.34 | 0.000 | Ctgf | connective tissue growth factor |
| 1448471 | a_at | 1.35 | 0.060 | Ctla2a | cytotoxic T lymphocyte-associated protein 2 alpha |
| 1452352 | at | 1.31 | 0.006 | Ctla2b | cytotoxic T lymphocyte-associated protein 2 beta |
| 1448582 | at | -1.02 | 0.810 | Ctnnbl1 | catenin, beta like 1 |
| 1416274 | at | 1.28 | 0.004 | Ctns | cystinosis, nephropathic |
| 1416563 | at | 1.34 | 0.008 | Ctps | cytidine 5'-triphosphate synthase |
| 1448111 | at | 1.09 | 0.030 | Ctps2 | cytidine 5'-triphosphate synthase 2 |
| 1446834 | at | 1.02 | 0.877 | Ctsc | cathepsin C |
| 1448118 | a_at | 1.15 | 0.000 | Ctsd | cathepsin D |
| 1451019 | at | -1.23 | 0.026 | Ctsf | cathepsin F |
| 1418365 | at | -1.08 | 0.104 | Ctsh | cathepsin H |
| 1457724 | at | -1.24 | 0.077 | Ctsl | cathepsin L |
| 1433501 | at | 1.14 | 0.003 | Ctso | cathepsin O |
| 1448591 | at | 1.58 | 0.000 | Ctss | cathepsin S |
| 1417868 | a_at | 1.71 | 0.001 | Ctsz | cathepsin Z |
| 1423917 | a_at | 1.34 | 0.000 | Cttn | cortactin |

Supplementary table S1

| | | | | | |
|---|---|---|---|---|---|
| 1431145 | a_at | -1.15 | 0.134 | Cuedc2 | CUE domain containing 2 |
| 1427413 | a_at | 1.28 | 0.008 | Cugbp1 | CUG triplet repeat, RNA binding protein 1 |
| 1448174 | at | -1.07 | 0.226 | Cul1 | cullin 1 |
| 1451231 | a_at | -1.00 | 0.966 | Cul2 | cullin 2 |
| 1422794 | at | -1.16 | 0.152 | Cul3 | cullin 3 |
| 1426060 | at | 1.14 | 0.371 | Cul4a | cullin 4A |
| 1417453 | at | 1.06 | 0.442 | Cul4b | cullin 4B |
| 1417039 | a_at | 1.22 | 0.048 | Cul7 | cullin 7 |
| 1418473 | at | -1.08 | 0.204 | Cutc | cutC copper transporter homolog (E.coli) |
| 1434683 | at | -1.11 | 0.523 | Cutl1 | cut-like 1 (Drosophila) |
| 1447500 | at | -1.22 | 0.012 | Cutl2 | cut-like 2 (Drosophila) |
| 1453282 | at | -1.18 | 0.229 | Cxadr | coxsackievirus and adenovirus receptor |
| 1417574 | at | -1.20 | 0.014 | Cxcl12 | chemokine (C-X-C motif) ligand 12 |
| 1418457 | at | -1.02 | 0.852 | Cxcl14 | chemokine (C-X-C motif) ligand 14 |
| 1448995 | at | -1.10 | 0.185 | Cxcl4 | chemokine (C-X-C motif) ligand 4 |
| 1418480 | at | -3.19 | 0.007 | Cxcl7 | chemokine (C-X-C motif) ligand 7 |
| 1418652 | at | -1.09 | 0.542 | Cxcl9 | chemokine (C-X-C motif) ligand 9 |
| 1448710 | at | 1.08 | 0.541 | Cxcr4 | chemokine (C-X-C motif) receptor 4 |
| 1452221 | a_at | -1.19 | 0.123 | Cxxc1 | CXXC finger 1 (PHD domain) |
| 1437351 | at | 1.05 | 0.736 | Cxxc4 | CXXC finger 4 |
| 1448960 | at | 1.02 | 0.757 | Cxxc5 | CXXC finger 5 |
| 1416727 | a_at | -1.09 | 0.045 | Cyb5 | cytochrome b-5 |
| 1417507 | at | 1.97 | 0.001 | Cyb561 | cytochrome b-561 |
| 1444117 | at | -1.30 | 0.000 | Cyb561d1 | cytochrome b-561 domain containing 1 |
| 1454268 | a_at | -1.20 | 0.102 | Cyba | cytochrome b-245, alpha polypeptide |
| 1454895 | at | 1.18 | 0.142 | Cybasc3 | cytochrome b, ascorbate dependent 3 |
| 1416604 | at | -1.03 | 0.315 | Cyc1 | cytochrome c-1 |
| 1416329 | at | 1.25 | 0.000 | Cyfip1 | cytoplasmic FMR1 interacting protein 1 |
| 1449273 | at | -1.23 | 0.007 | Cyfip2 | cytoplasmic FMR1 interacting protein 2 |
| 1423630 | at | 1.53 | 0.000 | Cygb | cytoglobin |
| 1429618 | at | 1.20 | 0.021 | Cyld | cylindromatosis (turban tumor syndrome) |
| 1416612 | at | 1.88 | 0.001 | Cyp1b1 | cytochrome P450, family 1, subfamily b, polypeptide 1 |
| 1435616 | at | 1.25 | 0.003 | Cyp20a1 | cytochrome P450, family 20, subfamily A, polypeptide 1 |
| 1417590 | at | 1.02 | 0.825 | Cyp27a1 | cytochrome P450, family 27, subfamily a, polypeptide 1 |
| 1419040 | at | -1.03 | 0.731 | Cyp2d22 | cytochrome P450, family 2, subfamily d, polypeptide 22 |
| 1442537 | at | -1.35 | 0.003 | Cyp2u1 | cytochrome P450, family 2, subfamily u, polypeptide 1 |
| 1418780 | at | 1.09 | 0.286 | Cyp39a1 | cytochrome P450, family 39, subfamily a, polypeptide 1 |
| 1416194 | at | -1.51 | 0.002 | Cyp4b1 | cytochrome P450, family 4, subfamily b, polypeptide 1 |
| 1450646 | at | 1.15 | 0.001 | Cyp51 | cytochrome P450, family 51 |
| 1457823 | at | 1.37 | 0.563 | Cyr61 | cysteine rich protein 61 |
| 1435701 | at | -1.20 | 0.003 | Cyyr1 | cysteine and tyrosine-rich protein 1 |

Supplementary table S1

| | | | | | |
|---|---|---|---|---|---|
| 1423879 | at | -1.07 | 0.380 | D030056L22 | hypothetical protein D030056L22 |
| 1455334 | at | -1.16 | 0.196 | D330038O06 | hypothetical protein D330038O06 |
| 1455244 | at | -1.13 | 0.002 | Daam1 | dishevelled associated activator of morphogenesis 1 |
| 1420498 | a_at | 1.34 | 0.108 | Dab2 | disabled homolog 2 (Drosophila) |
| 1433558 | at | -1.02 | 0.614 | Dab2ip | disabled homolog 2 (Drosophila) interacting protein |
| 1433743 | at | -1.19 | 0.157 | Dach1 | dachshund 1 (Drosophila) |
| 1418528 | a_at | -1.09 | 0.242 | Dad1 | defender against cell death 1 |
| 1418762 | at | 1.36 | 0.006 | Daf1 | decay accelerating factor 1 |
| 1423872 | a_at | -1.17 | 0.203 | Dag1 | dystroglycan 1 |
| 1423894 | a_at | -1.01 | 0.867 | Dalrd3 | DALR anticodon binding domain containing 3 |
| 1436476 | at | -1.26 | 0.031 | Dand5 | DAN domain family, member 5 |
| 1450848 | at | -1.07 | 0.018 | Dap3 | death associated protein 3 |
| 1426915 | at | -1.42 | 0.001 | Dapk1 | death associated protein kinase 1 |
| 1423800 | at | 1.13 | 0.000 | Dars | aspartyl-tRNA synthetase |
| 1419026 | at | 1.10 | 0.109 | Daxx | Fas death domain-associated protein |
| 1419275 | at | 1.06 | 0.269 | Dazap1 | DAZ associated protein 1 |
| 1415796 | at | 1.11 | 0.006 | Dazap2 | DAZ associated protein 2 |
| 1422432 | at | -1.26 | 0.000 | Dbi | diazepam binding inhibitor |
| 1460334 | at | 1.66 | 0.000 | Dbnl | drebrin-like |
| 1418174 | at | -1.36 | 0.507 | Dbp | D site albumin promoter binding protein |
| 1449118 | at | -1.36 | 0.005 | Dbt | dihydrolipoamide branched chain transacylase E2 |
| 1424271 | at | 1.52 | 0.010 | Dcamkl1 | double cortin and calcium/calmodulin-dependent protein kinase-like 1 |
| 1449291 | a_at | 1.11 | 0.212 | Dcbld1 | discoidin, CUB and LCCL domain containing 1 |
| 1437635 | at | -1.14 | 0.632 | Dcbld2 | discoidin, CUB and LCCL domain containing 2 |
| 1418321 | at | -1.43 | 0.000 | Dci | dodecenoyl-Coenzyme A delta isomerase (3,2 trans-enoyl-Coenyme A isomerase) |
| 1439012 | a_at | 1.30 | 0.003 | Dck | deoxycytidine kinase |
| 1417904 | at | -1.19 | 0.022 | Dclre1a | DNA cross-link repair 1A, PSO2 homolog (S. cerevisiae) |
| 1440383 | at | 1.03 | 0.831 | Dclre1b | DNA cross-link repair 1B, PSO2 homolog (S. cerevisiae) |
| 1424080 | at | -1.11 | 0.204 | Dcps | decapping enzyme, scavenger |
| 1422521 | at | -1.24 | 0.070 | Dctn1 | dynactin 1 |
| 1424461 | at | 1.13 | 0.080 | Dctn2 | dynactin 2 |
| 1416247 | at | -1.14 | 0.035 | Dctn3 | dynactin 3 |
| 1420861 | at | -1.03 | 0.455 | Dctn4 | dynactin 4 |
| 1415748 | a_at | -1.11 | 0.016 | Dctn5 | dynactin 5 |
| 1419456 | at | -1.22 | 0.059 | Dcxr | dicarbonyl L-xylulose reductase |
| 1454995 | at | 2.40 | 0.001 | Ddah1 | dimethylarginine dimethylaminohydrolase 1 |
| 1415735 | at | 1.06 | 0.154 | Ddb1 | damage specific DNA binding protein 1 |
| 1455321 | at | 1.15 | 0.279 | Ddhd1 | DDHD domain containing 1 |
| 1435376 | at | -1.01 | 0.635 | Ddhd2 | DDHD domain containing 2 |
| 1417516 | at | 1.23 | 0.116 | Ddit3 | DNA-damage inducible transcript 3 |
| 1428306 | at | -1.16 | 0.143 | Ddit4 | DNA-damage-inducible transcript 4 |

Supplementary table S1

| | | | | | |
|---|---|---|---|---|---|
| 1416493 | at | -1.10 | 0.144 | Ddost | dolichyl-di-phosphooligosaccharide-protein glycotransferase |
| 1455688 | at | 1.19 | 0.009 | Ddr2 | discoidin domain receptor family, member 2 |
| 1417103 | at | -1.20 | 0.000 | Ddt | D-dopachrome tautomerase |
| 1415915 | at | 1.00 | 0.901 | Ddx1 | DEAD (Asp-Glu-Ala-Asp) box polypeptide 1 |
| 1452838 | at | 1.11 | 0.096 | Ddx10 | DEAD (Asp-Glu-Ala-Asp) box polypeptide 10 |
| 1417927 | at | 1.20 | 0.004 | Ddx19a | DEAD (Asp-Glu-Ala-Asp) box polypeptide 19a |
| 1416751 | a_at | -1.02 | 0.750 | Ddx20 | DEAD (Asp-Glu-Ala-Asp) box polypeptide 20 |
| 1448270 | at | 1.15 | 0.007 | Ddx21 | DEAD (Asp-Glu-Ala-Asp) box polypeptide 21 |
| 1441631 | at | -1.63 | 0.002 | Ddx24 | DEAD (Asp-Glu-Ala-Asp) box polypeptide 24 |
| 1424161 | at | 1.09 | 0.276 | Ddx27 | DEAD (Asp-Glu-Ala-Asp) box polypeptide 27 |
| 1428702 | at | -1.13 | 0.025 | Ddx28 | DEAD (Asp-Glu-Ala-Asp) box polypeptide 28 |
| 1423643 | at | 1.37 | 0.000 | Ddx39 | DEAD (Asp-Glu-Ala-Asp) box polypeptide 39 |
| 1416467 | at | -1.03 | 0.668 | Ddx3x | DEAD/H (Asp-Glu-Ala-Asp/His) box polypeptide 3, X-linked |
| 1426439 | at | 1.07 | 0.453 | Ddx3y | DEAD (Asp-Glu-Ala-Asp) box polypeptide 3, Y-linked |
| 1427022 | at | 1.01 | 0.884 | Ddx42 | DEAD (Asp-Glu-Ala-Asp) box polypeptide 42 |
| 1423752 | at | -1.08 | 0.172 | Ddx47 | DEAD (Asp-Glu-Ala-Asp) box polypeptide 47 |
| 1417242 | at | 1.00 | 0.899 | Ddx48 | DEAD (Asp-Glu-Ala-Asp) box polypeptide 48 |
| 1419653 | a_at | -1.02 | 0.443 | Ddx5 | DEAD (Asp-Glu-Ala-Asp) box polypeptide 5 |
| 1417875 | at | 1.11 | 0.007 | Ddx50 | DEAD (Asp-Glu-Ala-Asp) box polypeptide 50 |
| 1434607 | at | -1.05 | 0.220 | Ddx52 | DEAD (Asp-Glu-Ala-Asp) box polypeptide 52 |
| 1423815 | at | 1.11 | 0.142 | Ddx56 | DEAD (Asp-Glu-Ala-Asp) box polypeptide 56 |
| 1439122 | at | -1.23 | 0.253 | Ddx6 | DEAD (Asp-Glu-Ala-Asp) box polypeptide 6 |
| 1448446 | at | 1.01 | 0.891 | Deaf1 | deformed epidermal autoregulatory factor 1 (Drosophila) |
| 1427955 | a_at | -1.15 | 0.000 | Deb1 | differentially expressed in B16F10 1 |
| 1419367 | at | -1.27 | 0.000 | Decr1 | 2,4-dienoyl CoA reductase 1, mitochondrial |
| 1434994 | at | -1.05 | 0.568 | Dedd | death effector domain-containing |
| 1415853 | at | 1.26 | 0.061 | Def8 | differentially expressed in FDCP 8 |
| 1423345 | at | 1.12 | 0.361 | Degs1 | degenerative spermatocyte homolog 1 (Drosophila) |
| 1452659 | at | -1.01 | 0.809 | Dek | DEK oncogene (DNA binding) |
| 1420368 | at | -1.17 | 0.034 | Denr | density-regulated protein |
| 1438892 | at | -1.12 | 0.274 | Dep1 | diabetic embryopathy 1 |
| 1428622 | at | -1.21 | 0.118 | Depdc6 | DEP domain containing 6 |
| 1448438 | at | -1.06 | 0.991 | Derl2 | Der1-like domain family, member 2 |
| 1426731 | at | 1.23 | 0.004 | Des | desmin |
| 1444774 | at | -1.15 | 0.174 | Det1 | de-etiolated homolog 1 (Arabidopsis) |
| 1460174 | at | 1.36 | 0.001 | Dexi | dexamethasone-induced transcript |
| 1450885 | at | 1.38 | 0.000 | Dffa | DNA fragmentation factor, alpha subunit |
| 1422678 | at | -1.23 | 0.039 | Dgat2 | diacylglycerol O-acyltransferase 2 |
| 1421810 | at | -1.31 | 0.003 | Dgcr2 | DiGeorge syndrome critical region gene 2 |
| 1428753 | a_at | -1.15 | 0.046 | Dgcr6 | DiGeorge syndrome critical region gene 6 |
| 1455311 | at | 1.05 | 0.417 | Dgcr8 | DiGeorge syndrome critical region gene 8 |

Supplementary table S1

| | | | | | |
|---|---|---|---|---|---|
| 1418578 | at | -1.22 | 0.006 | Dgka | diacylglycerol kinase, alpha |
| 1433564 | at | 1.25 | 0.020 | Dgkd | diacylglycerol kinase, delta |
| 1436547 | at | -1.35 | 0.133 | Dgke | diacylglycerol kinase, epsilon |
| 1425228 | a_at | -1.21 | 0.003 | Dguok | deoxyguanosine kinase |
| 1448619 | at | 1.07 | 0.518 | Dhcr7 | 7-dehydrocholesterol reductase |
| 1453487 | at | -1.32 | 0.088 | Dhdh | dihydrodiol dehydrogenase (dimeric) |
| 1417581 | at | -1.13 | 0.135 | Dhodh | dihydroorotate dehydrogenase |
| 1415677 | at | 1.22 | 0.056 | Dhrs1 | dehydrogenase/reductase (SDR family) member 1 |
| 1448390 | a_at | -1.05 | 0.552 | Dhrs3 | dehydrogenase/reductase (SDR family) member 3 |
| 1419382 | a_at | -1.27 | 0.014 | Dhrs4 | dehydrogenase/reductase (SDR family) member 4 |
| 1426440 | at | -1.45 | 0.000 | Dhrs7 | dehydrogenase/reductase (SDR family) member 7 |
| 1434642 | at | -1.34 | 0.012 | Dhrs8 | dehydrogenase/reductase (SDR family) member 8 |
| 1423925 | at | 1.02 | 0.508 | Dhx16 | DEAH (Asp-Glu-Ala-His) box polypeptide 16 |
| 1434710 | at | 1.25 | 0.000 | Dhx29 | DEAH (Asp-Glu-Ala-His) box polypeptide 29 |
| 1453251 | at | -1.31 | 0.062 | Dhx30 | DEAH (Asp-Glu-Ala-His) box polypeptide 30 |
| 1420427 | a_at | 1.30 | 0.042 | Dhx32 | DEAH (Asp-Glu-Ala-His) box polypeptide 32 |
| 1429191 | at | 1.02 | 0.823 | Dhx33 | DEAH (Asp-Glu-Ala-His) box polypeptide 33 |
| 1452693 | at | 1.04 | 0.621 | Dhx35 | DEAH (Asp-Glu-Ala-His) box polypeptide 35 |
| 1437550 | at | -1.13 | 0.077 | Dhx36 | DEAH (Asp-Glu-Ala-His) box polypeptide 36 |
| 1428267 | at | -1.02 | 0.484 | Dhx40 | DEAH (Asp-Glu-Ala-His) box polypeptide 40 |
| 1425617 | at | 1.14 | 0.041 | Dhx9 | DEAH (Asp-Glu-Ala-His) box polypeptide 9 |
| 1422185 | a_at | 1.19 | 0.063 | Dia1 | diaphorase 1 (NADH) |
| 1417683 | at | 1.08 | 0.376 | Diablo | diablo homolog (Drosophila) |
| 1454979 | at | 1.13 | 0.206 | Diap1 | diaphanous homolog 1 (Drosophila) |
| 1427564 | at | 1.02 | 0.805 | Diap2 | diaphanous homolog 2 (Drosophila) |
| 1418844 | at | -1.00 | 0.984 | Dibd1 | disrupted in bipolar disorder 1 homolog (human) |
| 1452908 | at | -1.08 | 0.113 | Dip2 | disco interacting protein 2 homolog (Drosophila) |
| 1426743 | at | -1.29 | 0.003 | Dip3b | Dip3 beta |
| 1454654 | at | -1.36 | 0.014 | Dirc2 | disrupted in renal carcinoma 2 (human) |
| 1434795 | at | 1.02 | 0.893 | Disp1 | dispatched homolog 1 (Drosophila) |
| 1435207 | at | -1.96 | 0.006 | Dixdc1 | DIX domain containing 1 |
| 1460602 | at | -1.20 | 0.085 | Dlc1 | deleted in liver cancer 1 |
| 1423159 | at | -1.14 | 0.018 | Dld | dihydrolipoamide dehydrogenase |
| 1427410 | at | -1.25 | 0.034 | Dleu2 | deleted in lymphocytic leukemia, 2 |
| 1424198 | at | 1.31 | 0.011 | Dlg5 | discs, large homolog 5 (Drosophila) |
| 1455548 | at | 1.16 | 0.258 | Dlgap4 | discs, large homolog-associated protein 4 (Drosophila) |
| 1450768 | at | 1.05 | 0.602 | Dlgh1 | discs, large homolog 1 (Drosophila) |
| 1446585 | at | 1.14 | 0.461 | Dlgh2 | discs, large homolog 2 (Drosophila) |
| 1416918 | at | 1.10 | 0.127 | Dlgh3 | discs, large homolog 3 (Drosophila) |
| 1437775 | at | -1.10 | 0.398 | Dlst | dihydrolipoamide S-succinyltransferase (E2 component of 2-oxo-glutarate complex) |
| 1434944 | at | 1.02 | 0.700 | Dm15 | dystrophia myotonica kinase, B15 |

Supplementary table S1

| | | | | | |
|---|---|---|---|---|---|
| 1446156 | at | 1.23 | 0.139 | Dmd | dystrophin, muscular dystrophy |
| 1422636 | at | -1.17 | 0.334 | Dmtf1 | cyclin D binding myb-like transcription factor 1 |
| 1435505 | at | 1.07 | 0.375 | Dmwd | dystrophia myotonica-containing WD repeat motif |
| 1416288 | at | 1.15 | 0.376 | Dnaja1 | DnaJ (Hsp40) homolog, subfamily A, member 1 |
| 1417183 | at | 1.09 | 0.273 | Dnaja2 | DnaJ (Hsp40) homolog, subfamily A, member 2 |
| 1434196 | at | 1.61 | 0.002 | Dnaja4 | DnaJ (Hsp40) homolog, subfamily A, member 4 |
| 1416755 | at | 1.16 | 0.421 | Dnajb1 | DnaJ (Hsp40) homolog, subfamily B, member 1 |
| 1448657 | a_at | -1.09 | 0.037 | Dnajb10 | DnaJ (Hsp40) homolog, subfamily B, member 10 |
| 1423151 | at | 1.02 | 0.749 | Dnajb11 | DnaJ (Hsp40) homolog, subfamily B, member 11 |
| 1431734 | a_at | 1.27 | 0.029 | Dnajb4 | DnaJ (Hsp40) homolog, subfamily B, member 4 |
| 1421962 | at | -1.07 | 0.652 | Dnajb5 | DnaJ (Hsp40) homolog, subfamily B, member 5 |
| 1448234 | at | 1.12 | 0.022 | Dnajb6 | DnaJ (Hsp40) homolog, subfamily B, member 6 |
| 1417191 | at | 1.02 | 0.757 | Dnajb9 | DnaJ (Hsp40) homolog, subfamily B, member 9 |
| 1426905 | a_at | 1.23 | 0.001 | Dnajc10 | DnaJ (Hsp40) homolog, subfamily C, member 10 |
| 1433881 | at | 1.18 | 0.630 | Dnajc11 | DnaJ (Hsp40) homolog, subfamily C, member 11 |
| 1437546 | at | 1.27 | 0.001 | Dnajc14 | DnaJ (Hsp40) homolog, subfamily C, member 14 |
| 1417934 | at | -1.30 | 0.001 | Dnajc4 | DnaJ (Hsp40) homolog, subfamily C, member 4 |
| 1452683 | at | -1.01 | 0.869 | Dnajc8 | DnaJ (Hsp40) homolog, subfamily C, member 8 |
| 1426473 | at | -1.13 | 0.050 | Dnajc9 | DnaJ (Hsp40) homolog, subfamily C, member 9 |
| 1416910 | at | -1.09 | 0.008 | Dnajd1 | DnaJ (Hsp40) homolog, subfamily D, member 1 |
| 1416870 | at | -1.26 | 0.002 | Dnalc4 | dynein, axonemal, light chain 4 |
| 1429173 | at | 1.30 | 0.011 | Dnase1l1 | deoxyribonuclease 1-like 1 |
| 1416648 | at | 1.20 | 0.008 | Dnchc1 | dynein, cytoplasmic, heavy chain 1 |
| 1415841 | at | 1.14 | 0.012 | Dncic2 | dynein, cytoplasmic, intermediate chain 2 |
| 1440278 | at | 1.12 | 0.414 | Dnclc1 | dynein, cytoplasmic, light chain 1 |
| 1424947 | at | 1.31 | 0.107 | Dnclic1 | dynein, cytoplasmic, light intermediate chain 1 |
| 1456346 | at | 1.70 | 0.000 | Dnm1 | dynamin 1 |
| 1428087 | at | 1.02 | 0.696 | Dnm1l | dynamin 1-like |
| 1425135 | a_at | -1.02 | 0.740 | Dnm2 | dynamin 2 |
| 1427033 | at | -1.00 | 0.972 | Dnmbp | dynamin binding protein |
| 1419750 | at | -1.14 | 0.116 | Dnmt2 | DNA methyltransferase 2 |
| 1416127 | a_at | 1.05 | 0.369 | Dnpep | aspartyl aminopeptidase |
| 1417666 | at | 1.10 | 0.041 | Dnttip1 | deoxynucleotidyltransferase, terminal, interacting protein 1 |
| 1450707 | at | -1.20 | 0.037 | Doc2g | double C2, gamma |
| 1429028 | at | 1.24 | 0.008 | Dock11 | dedicator of cytokinesis 11 |
| 1431114 | at | 1.05 | 0.434 | Dock4 | dedicator of cytokinesis 4 |
| 1430291 | at | -1.05 | 0.336 | Dock5 | dedicator of cytokinesis 5 |
| 1449419 | at | -1.01 | 0.917 | Dock8 | dedicator of cytokinesis 8 |
| 1450931 | at | 1.23 | 0.054 | Dock9 | dedicator of cytokinesis 9 |
| 1426739 | at | 1.14 | 0.043 | Donson | downstream neighbor of SON |
| 1426376 | at | -1.13 | 0.170 | Dp1 | deleted in polyposis 1 |

Supplementary table S1

| | | | | | |
|---|---|---|---|---|---|
| 1430128 | a_at | -1.14 | 0.361 | Dp1l1 | deleted in polyposis 1-like 1 |
| 1448549 | a_at | 1.05 | 0.157 | Dpagt1 | dolichyl-phosphate (UDP-N-acetylglucosamine) acetylglucosaminephosphotransferase 1 (GlcNAc-1-P transferase) |
| 1435943 | at | -1.46 | 0.001 | Dpep1 | dipeptidase 1 (renal) |
| 1429903 | at | -1.04 | 0.698 | Dpf2 | D4, zinc and double PHD fingers family 2 |
| 1449438 | at | -1.06 | 0.034 | Dpm1 | dolichol-phosphate (beta-D) mannosyltransferase 1 |
| 1415675 | at | 1.07 | 0.145 | Dpm2 | dolichol-phosphate (beta-D) mannosyltransferase 2 |
| 1416697 | at | -1.04 | 0.672 | Dpp4 | dipeptidylpeptidase 4 |
| 1428697 | at | 1.07 | 0.051 | Dpp8 | dipeptidylpeptidase 8 |
| 1455884 | at | 1.04 | 0.652 | Dpp9 | dipeptidylpeptidase 9 |
| 1418511 | at | -1.02 | 0.690 | Dpt | dermatopontin |
| 1433770 | at | 1.02 | 0.582 | Dpysl2 | dihydropyrimidinase-like 2 |
| 1454613 | at | 1.71 | 0.000 | Dpysl3 | dihydropyrimidinase-like 3 |
| 1416018 | at | 1.07 | 0.284 | Dr1 | down-regulator of transcription 1 |
| 1451676 | at | -1.13 | 0.002 | Drap1 | Dr1 associated protein 1 (negative cofactor 2 alpha) |
| 1450724 | at | -1.08 | 0.297 | Drctnnb1a | down-regulated by Ctnnb1, a |
| 1448336 | at | 1.21 | 0.001 | Drg1 | developmentally regulated GTP binding protein 1 |
| 1417692 | at | 1.15 | 0.352 | Drg2 | developmentally regulated GTP binding protein 2 |
| 1421149 | a_at | -1.00 | 0.975 | Drpla | dentatorubral pallidoluysian atrophy |
| 1426911 | at | -1.09 | 0.392 | Dsc2 | desmocollin 2 |
| 1416600 | a_at | 3.51 | 0.000 | Dscr1 | Down syndrome critical region homolog 1 (human) |
| 1450243 | a_at | 1.00 | 0.971 | Dscr1l1 | Down syndrome critical region gene 1-like 1 |
| 1434027 | at | 1.16 | 0.186 | Dscr1l2 | Down syndrome critical region gene 1-like 2 |
| 1448307 | at | -1.13 | 0.018 | Dscr2 | Down syndrome critical region homolog 2 (human) |
| 1415745 | a_at | 1.25 | 0.038 | Dscr3 | Down syndrome critical region gene 3 |
| 1436038 | a_at | -1.04 | 0.368 | Dscr5 | Down syndrome critical region homolog 5 (human) |
| 1420459 | at | 1.07 | 0.326 | Dscr6 | Down syndrome critical region homolog 6 (human) |
| 1439476 | at | -1.51 | 0.052 | Dsg2 | desmoglein 2 |
| 1435493 | at | 1.06 | 0.025 | Dsp | desmoplakin |
| 1417124 | at | 1.07 | 0.046 | Dstn | destrin |
| 1456069 | at | 1.24 | 0.079 | Dtna | dystrobrevin alpha |
| 1431619 | a_at | -1.13 | 0.181 | Dtnbp1 | dystrobrevin binding protein 1 |
| 1420752 | at | 1.19 | 0.220 | Dtx3 | deltex 3 homolog (Drosophila) |
| 1436545 | at | 1.08 | 0.242 | Dtx4 | deltex 4 homolog (Drosophila) |
| 1452681 | at | -1.16 | 0.015 | Dtymk | deoxythymidylate kinase |
| 1452100 | at | -1.11 | 0.153 | Dullard | Dullard homolog (Xenopus laevis) |
| 1448830 | at | -1.13 | 0.111 | Dusp1 | dual specificity phosphatase 1 |
| 1417163 | at | -1.21 | 0.057 | Dusp10 | dual specificity phosphatase 10 |
| 1452594 | at | -1.05 | 0.551 | Dusp11 | dual specificity phosphatase 11 (RNA/RNP complex 1-interacting) |
| 1420636 | a_at | -1.05 | 0.537 | Dusp12 | dual specificity phosphatase 12 |
| 1442344 | at | 1.01 | 0.842 | Dusp14 | dual specificity phosphatase 14 |
| 1451270 | at | -1.77 | 0.007 | Dusp18 | dual specificity phosphatase 18 |

Supplementary table S1

| | | | | | |
|---|---|---|---|---|---|
| 1448922 | at | -1.10 | 0.274 | Dusp19 | dual specificity phosphatase 19 |
| 1448985 | at | 1.07 | 0.104 | Dusp22 | dual specificity phosphatase 22 |
| 1434472 | at | 1.13 | 0.164 | Dusp3 | dual specificity phosphatase 3 (vaccinia virus phosphatase VH1-related) |
| 1415834 | at | 1.18 | 0.014 | Dusp6 | dual specificity phosphatase 6 |
| 1452097 | a_at | -1.69 | 0.001 | Dusp7 | dual specificity phosphatase 7 |
| 1418714 | at | 1.27 | 0.102 | Dusp8 | dual specificity phosphatase 8 |
| 1419270 | a_at | -1.31 | 0.001 | Dut | deoxyuridine triphosphatase |
| 1450978 | at | -1.02 | 0.714 | Dvl1 | dishevelled, dsh homolog 1 (Drosophila) |
| 1442006 | at | 1.01 | 0.905 | Dvl3 | dishevelled 3, dsh homolog (Drosophila) |
| 1423736 | a_at | -1.07 | 0.047 | Dym | dymeclin |
| 1435591 | at | -1.21 | 0.017 | Dyrk1b | dual-specificity tyrosine-(Y)-phosphorylation regulated kinase 1b |
| 1451891 | a_at | 1.02 | 0.813 | Dysf | dysferlin |
| 1434564 | at | 1.16 | 0.057 | E2f3 | E2F transcription factor 3 |
| 1437914 | at | 1.06 | 0.298 | E2f6 | E2F transcription factor 6 |
| 1433555 | at | -1.02 | 0.781 | Eaf1 | ELL associated factor 1 |
| 1422669 | at | 1.10 | 0.046 | Ebag9 | estrogen receptor-binding fragment-associated gene 9 |
| 1418494 | at | -1.20 | 0.105 | Ebf2 | early B-cell factor 2 |
| 1428315 | at | 1.06 | 0.255 | Ebna1bp2 | EBNA1 binding protein 2 |
| 1416667 | at | 1.00 | 0.985 | Ebp | phenylalkylamine Ca2+ antagonist (emopamil) binding protein |
| 1455741 | a_at | -1.07 | 0.248 | Ece1 | endothelin converting enzyme 1 |
| 1448491 | at | -1.22 | 0.000 | Ech1 | enoyl coenzyme A hydratase 1, peroxisomal |
| 1425788 | a_at | -1.33 | 0.000 | Echdc2 | enoyl Coenzyme A hydratase domain containing 2 |
| 1418862 | at | -1.40 | 0.000 | Echdc3 | enoyl Coenzyme A hydratase domain containing 3 |
| 1452341 | at | -1.23 | 0.001 | Echs1 | enoyl Coenzyme A hydratase, short chain, 1, mitochondrial |
| 1448613 | at | 1.52 | 0.006 | Ecm1 | extracellular matrix protein 1 |
| 1448266 | at | -1.06 | 0.208 | Edf1 | endothelial differentiation-related factor 1 |
| 1423571 | at | -1.04 | 0.735 | Edg1 | endothelial differentiation sphingolipid G-protein-coupled receptor 1 |
| 1438658 | a_at | -1.33 | 0.027 | Edg3 | endothelial differentiation, sphingolipid G-protein-coupled receptor, 3 |
| 1433525 | at | -1.12 | 0.029 | Ednra | endothelin receptor type A |
| 1437347 | at | 1.44 | 0.000 | Ednrb | endothelin receptor type B |
| 1448653 | at | -1.00 | 0.942 | Eed | embryonic ectoderm development |
| 1424635 | at | 1.11 | 0.001 | Eef1a1 | eukaryotic translation elongation factor 1 alpha 1 |
| 1418062 | at | -1.18 | 0.023 | Eef1a2 | eukaryotic translation elongation factor 1 alpha 2 |
| 1448252 | a_at | 1.06 | 0.077 | Eef1b2 | eukaryotic translation elongation factor 1 beta 2 |
| 1449506 | a_at | -1.08 | 0.045 | Eef1d | eukaryotic translation elongation factor 1 delta (guanine nucleotide exchange protein) |
| 1449044 | at | 1.09 | 0.283 | Eef1e1 | eukaryotic translation elongation factor 1 epsilon 1 |
| 1417364 | at | 1.03 | 0.374 | Eef1g | eukaryotic translation elongation factor 1 gamma |
| 1424736 | at | -1.02 | 0.573 | Eef2 | eukaryotic translation elongation factor 2 |
| 1427183 | at | -2.24 | 0.001 | Efemp1 | epidermal growth factor-containing fibulin-like extracellular matrix protein 1 |
| 1417018 | at | 1.42 | 0.006 | Efemp2 | epidermal growth factor-containing fibulin-like extracellular matrix protein 2 |
| 1454808 | at | 1.19 | 0.001 | Efha1 | EF hand domain family A1 |

Supplementary table S1

| | | | | | |
|---|---|---|---|---|---|
| 1448507 | at | 1.27 | 0.013 | Efhd1 | EF hand domain containing 1 |
| 1416895 | at | -1.04 | 0.455 | Efna1 | ephrin A1 |
| 1444606 | at | -1.34 | 0.000 | Efna2 | ephrin A2 |
| 1418285 | at | 1.04 | 0.625 | Efnb1 | ephrin B1 |
| 1419639 | at | -1.10 | 0.274 | Efnb2 | ephrin B2 |
| 1423085 | at | -2.14 | 0.000 | Efnb3 | ephrin B3 |
| 1460562 | at | 1.15 | 0.233 | Eftud1 | elongation factor Tu GTP binding domain containing 1 |
| 1418093 | a_at | -1.09 | 0.405 | Egf | epidermal growth factor |
| 1433968 | a_at | 1.17 | 0.092 | Egfl5 | EGF-like-domain, multiple 5 |
| 1421335 | a_at | 1.11 | 0.189 | Egfl7 | EGF-like domain 7 |
| 1424932 | at | 1.05 | 0.607 | Egfr | epidermal growth factor receptor |
| 1451110 | at | -1.27 | 0.301 | Egln1 | EGL nine homolog 1 (C. elegans) |
| 1416533 | at | 1.05 | 0.379 | Egln2 | EGL nine homolog 2 (C. elegans) |
| 1418648 | at | 1.17 | 0.057 | Egln3 | EGL nine homolog 3 (C. elegans) |
| 1417065 | at | -1.07 | 0.574 | Egr1 | early growth response 1 |
| 1427683 | at | 2.85 | 0.002 | Egr2 | early growth response 2 |
| 1424586 | at | -1.13 | 0.051 | Ehbp1 | EH domain binding protein 1 |
| 1424157 | at | -1.03 | 0.492 | Ehd2 | EH-domain containing 2 |
| 1449852 | a_at | 1.24 | 0.000 | Ehd4 | EH-domain containing 4 |
| 1448382 | at | -1.33 | 0.017 | Ehhadh | enoyl-Coenzyme A, hydratase/3-hydroxyacyl Coenzyme A dehydrogenase |
| 1454776 | at | -1.08 | 0.208 | Ehmt1 | euchromatic histone methyltransferase 1 |
| 1416555 | at | 1.02 | 0.600 | Ei24 | etoposide induced 2.4 mRNA |
| 1424343 | a_at | 1.52 | 0.001 | Eif1a | eukaryotic translation initiation factor 1A |
| 1421901 | at | 1.01 | 0.885 | Eif2ak1 | eukaryotic translation initiation factor 2 alpha kinase 1 |
| 1421389 | a_at | -1.04 | 0.522 | Eif2ak4 | eukaryotic translation initiation factor 2 alpha kinase 4 |
| 1460730 | at | -1.18 | 0.023 | Eif2b1 | eukaryotic translation initiation factor 2B, subunit 1 (alpha) |
| 1434524 | at | 1.02 | 0.680 | Eif2b3 | eukaryotic translation initiation factor 2B, subunit 3 |
| 1449940 | a_at | -1.05 | 0.343 | Eif2b4 | eukaryotic translation initiation factor 2B, subunit 4 delta |
| 1433886 | at | -1.01 | 0.877 | Eif2b5 | eukaryotic translation initiation factor 2B, subunit 5 epsilon |
| 1434331 | at | -1.08 | 0.140 | Eif2c1 | eukaryotic translation initiation factor 2C, 1 |
| 1435904 | at | 1.01 | 0.759 | Eif2c3 | eukaryotic translation initiation factor 2C, 3 |
| 1448819 | at | 1.11 | 0.981 | Eif2s2 | eukaryotic translation initiation factor 2, subunit 2 (beta) |
| 1451090 | a_at | 1.05 | 0.230 | Eif2s3x | eukaryotic translation initiation factor 2, subunit 3, structural gene X-linked |
| 1417210 | at | 1.12 | 0.009 | Eif2s3y | eukaryotic translation initiation factor 2, subunit 3, structural gene Y-linked |
| 1426394 | at | 1.18 | 0.012 | Eif3s1 | eukaryotic translation initiation factor 3, subunit 1 alpha |
| 1416660 | at | 1.14 | 0.533 | Eif3s10 | eukaryotic translation initiation factor 3, subunit 10 (theta) |
| 1452674 | a_at | -1.09 | 0.039 | Eif3s12 | eukaryotic translation initiation factor 3, subunit 12 |
| 1415962 | at | 1.03 | 0.350 | Eif3s3 | eukaryotic translation initiation factor 3, subunit 3 (gamma) |
| 1417718 | at | -1.04 | 0.519 | Eif3s4 | eukaryotic translation initiation factor 3, subunit 4 (delta) |
| 1452285 | a_at | -1.07 | 0.012 | Eif3s5 | eukaryotic translation initiation factor 3, subunit 5 (epsilon) |
| 1460432 | a_at | 1.04 | 0.177 | Eif3s6 | eukaryotic translation initiation factor 3, subunit 6 |

Supplementary table S1

| | | | | | |
|---|---|---|---|---|---|
| 1423728 | at | 1.04 | 0.213 | Eif3s6ip | eukaryotic translation initiation factor 3, subunit 6 interacting protein |
| 1416100 | at | -1.02 | 0.808 | Eif3s7 | eukaryotic translation initiation factor 3, subunit 7 (zeta) |
| 1415858 | at | -1.05 | 0.893 | Eif3s8 | eukaryotic translation initiation factor 3, subunit 8 |
| 1426674 | at | 1.15 | 0.011 | Eif3s9 | eukaryotic translation initiation factor 3, subunit 9 (eta) |
| 1427058 | at | 1.18 | 0.219 | Eif4a1 | eukaryotic translation initiation factor 4A1 |
| 1450934 | at | -1.07 | 0.036 | Eif4a2 | eukaryotic translation initiation factor 4A2 |
| 1426380 | at | 1.08 | 0.349 | Eif4b | eukaryotic translation initiation factor 4B |
| 1450908 | at | 1.09 | 0.362 | Eif4e | eukaryotic translation initiation factor 4E |
| 1455108 | at | 1.36 | 0.002 | Eif4e2 | eukaryotic translation initiation factor 4E member 2 |
| 1417977 | at | 1.08 | 0.917 | Eif4e3 | eukaryotic translation initiation factor 4E member 3 |
| 1453023 | at | 1.16 | 0.132 | Eif4ebp3 | eukaryotic translation initiation factor 4E binding protein 3 |
| 1427037 | at | -1.09 | 0.848 | Eif4g1 | eukaryotic translation initiation factor 4, gamma 1 |
| 1415863 | at | -1.03 | 0.235 | Eif4g2 | eukaryotic translation initiation factor 4, gamma 2 |
| 1426833 | at | 1.15 | 0.009 | Eif4g3 | eukaryotic translation initiation factor 4 gamma, 3 |
| 1456256 | at | 1.08 | 0.593 | Eif5 | eukaryotic translation initiation factor 5 |
| 1419228 | at | -1.11 | 0.094 | Elac1 | elaC homolog 1 (E. coli) |
| 1428477 | at | 1.15 | 0.001 | Elac2 | elaC homolog 2 (E. coli) |
| 1448151 | at | -1.14 | 0.419 | Elavl1 | ELAV (embryonic lethal, abnormal vision, Drosophila)-like 1 (Hu antigen R) |
| 1417540 | at | -1.25 | 0.958 | Elf1 | E74-like factor 1 |
| 1421897 | at | -1.04 | 0.669 | Elk1 | ELK1, member of ETS oncogene family |
| 1417662 | at | 1.16 | 0.846 | Elk3 | ELK3, member of ETS oncogene family |
| 1427162 | a_at | 1.04 | 0.170 | Elk4 | ELK4, member of ETS oncogene family |
| 1460643 | at | 1.31 | 0.011 | Ell | elongation factor RNA polymerase II |
| 1450744 | at | 1.56 | 0.000 | Ell2 | elongation factor RNA polymerase II 2 |
| 1426035 | at | 1.21 | 0.132 | Ell3 | elongation factor RNA polymerase II-like 3 |
| 1433730 | at | 1.02 | 0.831 | Elmod2 | ELMO domain containing 2 |
| 1420855 | at | 1.59 | 0.005 | Eln | elastin |
| 1423820 | at | -1.09 | 0.148 | Elof1 | elongation factor 1 homolog (ELF1, S. cerevisiae) |
| 1415840 | at | -1.07 | 0.223 | Elovl5 | ELOVL family member 5, elongation of long chain fatty acids (yeast) |
| 1426643 | at | 1.02 | 0.734 | Elp3 | elongation protein 3 homolog (S. cerevisiae) |
| 1448535 | at | 1.02 | 0.716 | Elp4 | elongation protein 4 homolog (S. cerevisiae) |
| 1451459 | at | 1.08 | 0.938 | Elys | embryonic large molecule derived from yolk sac |
| 1425582 | a_at | -1.03 | 0.672 | Emcn | endomucin |
| 1417357 | at | 1.06 | 0.293 | Emd | emerin |
| 1435264 | at | -2.01 | 0.000 | Emilin2 | elastin microfibril interfacer 2 |
| 1428321 | at | 1.08 | 0.161 | Eml1 | echinoderm microtubule associated protein like 1 |
| 1428173 | at | -1.08 | 0.199 | Eml2 | echinoderm microtubule associated protein like 2 |
| 1416529 | at | 2.56 | 0.000 | Emp1 | epithelial membrane protein 1 |
| 1433670 | at | -1.04 | 0.414 | Emp2 | epithelial membrane protein 2 |
| 1417104 | at | 1.33 | 0.092 | Emp3 | epithelial membrane protein 3 |
| 1451161 | a_at | 1.56 | 0.000 | Emr1 | EGF-like module containing, mucin-like, hormone receptor-like sequence 1 |

Supplementary table S1

| | | | | | |
|---|---|---|---|---|---|
| 1458876 | at | 2.01 | 0.005 | Enah | enabled homolog (Drosophila) |
| 1450061 | at | 1.32 | 0.020 | Enc1 | ectodermal-neural cortex 1 |
| 1421097 | at | -1.22 | 0.053 | Endog | endonuclease G |
| 1419022 | a_at | 1.10 | 0.302 | Eno1 | enolase 1, alpha non-neuron |
| 1417951 | at | -1.09 | 0.001 | Eno3 | enolase 3, beta muscle |
| 1448649 | at | -1.01 | 0.942 | Enpep | glutamyl aminopeptidase |
| 1415894 | at | -2.11 | 0.131 | Enpp2 | ectonucleotide pyrophosphatase/phosphodiesterase 2 |
| 1439260 | a_at | -1.12 | 0.422 | Enpp3 | ectonucleotide pyrophosphatase/phosphodiesterase 3 |
| 1434580 | at | 1.15 | 0.046 | Enpp4 | ectonucleotide pyrophosphatase/phosphodiesterase 4 |
| 1428620 | at | 1.26 | 0.013 | Ensa | endosulfine alpha |
| 1418259 | a_at | 1.17 | 0.032 | Entpd2 | ectonucleoside triphosphate diphosphohydrolase 2 |
| 1449190 | a_at | -1.10 | 0.020 | Entpd4 | ectonucleoside triphosphate diphosphohydrolase 4 |
| 1417383 | at | -1.58 | 0.001 | Entpd5 | ectonucleoside triphosphate diphosphohydrolase 5 |
| 1460433 | at | -1.42 | 0.000 | Entpd6 | ectonucleoside triphosphate diphosphohydrolase 6 |
| 1429411 | a_at | 1.01 | 0.744 | Eny2 | enhancer of yellow 2 homolog (Drosophila) |
| 1434765 | at | 1.06 | 0.082 | Ep300 | E1A binding protein p300 |
| 1426860 | at | -1.16 | 0.301 | Ep400 | E1A binding protein p400 |
| 1435436 | at | 1.07 | 0.139 | Epas1 | endothelial PAS domain protein 1 |
| 1424092 | at | -1.41 | 0.030 | Epb4.1 | erythrocyte protein band 4.1 |
| 1434575 | at | -1.06 | 0.395 | Epb4.1l1 | erythrocyte protein band 4.1-like 1 |
| 1449167 | at | -1.08 | 0.275 | Epb4.1l4a | erythrocyte protein band 4.1-like 4a |
| 1418850 | at | -1.05 | 0.411 | Epc1 | enhancer of polycomb homolog 1 (Drosophila) |
| 1455317 | at | 1.07 | 0.182 | Epc2 | enhancer of polycomb homolog 2 (Drosophila) |
| 1450380 | at | 1.17 | 0.004 | Epdr2 | ependymin related protein 2 (zebrafish) |
| 1429021 | at | -2.22 | 0.000 | Epha4 | Eph receptor A4 |
| 1449845 | a_at | 1.07 | 0.189 | Ephb4 | Eph receptor B4 |
| 1422438 | at | 1.23 | 0.089 | Ephx1 | epoxide hydrolase 1, microsomal |
| 1448499 | a_at | -1.19 | 0.000 | Ephx2 | epoxide hydrolase 2, cytoplasmic |
| 1434106 | at | -1.36 | 0.071 | Epm2aip1 | EPM2A (laforin) interacting protein 1 |
| 1427039 | at | 1.03 | 0.638 | Epn1 | epsin 1 |
| 1427226 | at | -1.16 | 0.067 | Epn2 | epsin 2 |
| 1452158 | at | 1.19 | 0.123 | Eprs | glutamyl-prolyl-tRNA synthetase |
| 1419252 | at | 1.15 | 0.474 | Eps15 | epidermal growth factor receptor pathway substrate 15 |
| 1422823 | at | 1.02 | 0.859 | Eps8 | epidermal growth factor receptor pathway substrate 8 |
| 1420519 | a_at | -1.27 | 0.028 | Eral1 | Era (G-protein)-like 1 (E. coli) |
| 1434606 | at | -1.13 | 0.009 | Erbb3 | v-erb-b2 erythroblastic leukemia viral oncogene homolog 3 (avian) |
| 1447043 | at | -1.33 | 0.061 | Erbb4 | v-erb-a erythroblastic leukemia viral oncogene homolog 4 (avian) |
| 1417328 | at | -1.25 | 0.004 | Ercc1 | excision repair cross-complementing rodent repair deficiency, complementation group 1 |
| 1448497 | at | -1.09 | 0.249 | Ercc3 | excision repair cross-complementing rodent repair deficiency, complementation group 3 |
| 1450935 | at | -1.12 | 0.021 | Ercc5 | excision repair cross-complementing rodent repair deficiency, complementation group 5 |
| 1435561 | at | 1.04 | 0.670 | Erf | Ets2 repressor factor |

Supplementary table S1

| | | | | | |
|---|---|---|---|---|---|
| 1430536 | a_at | 1.06 | 0.112 | Erh | enhancer of rudimentary homolog (Drosophila) |
| 1437119 | at | 1.19 | 0.044 | Ern1 | endoplasmic reticulum (ER) to nucleus signalling 1 |
| 1449324 | at | 1.24 | 0.034 | Ero1l | ERO1-like (S. cerevisiae) |
| 1453634 | a_at | -1.03 | 0.693 | Erp29 | endoplasmic reticulum protein 29 |
| 1460356 | at | 1.06 | 0.411 | Esam1 | endothelial cell-specific adhesion molecule |
| 1424324 | at | 1.04 | 0.565 | Esco1 | establishment of cohesion 1 homolog 1 (S. cerevisiae) |
| 1417825 | at | 1.07 | 0.164 | Esd | esterase D/formylglutathione hydrolase |
| 1460652 | at | -1.27 | 0.001 | Esrra | estrogen related receptor, alpha |
| 1436926 | at | -1.02 | 0.617 | Esrrb | estrogen related receptor, beta |
| 1418929 | at | 1.06 | 0.303 | Esrrbl1 | estrogen-related receptor beta like 1 |
| 1455267 | at | -1.07 | 0.486 | Esrrg | estrogen-related receptor gamma |
| 1451208 | at | 1.25 | 0.014 | Etf1 | eukaryotic translation termination factor 1 |
| 1423972 | at | -1.08 | 0.007 | Etfa | electron transferring flavoprotein, alpha polypeptide |
| 1428181 | at | -1.19 | 0.000 | Etfb | electron transferring flavoprotein, beta polypeptide |
| 1451084 | at | -1.08 | 0.026 | Etfdh | electron transferring flavoprotein, dehydrogenase |
| 1417203 | at | -1.08 | 0.287 | Ethe1 | ethylmalonic encephalopathy 1 |
| 1433514 | at | -1.09 | 0.066 | Etnk1 | ethanolamine kinase 1 |
| 1429712 | at | 1.20 | 0.024 | Etohi1 | ethanol induced 1 |
| 1428656 | at | -1.13 | 0.029 | Etohi2 | ethanol induced 2 |
| 1416268 | at | 1.04 | 0.012 | Ets2 | E26 avian leukemia oncogene 2, 3' domain |
| 1428142 | at | 1.56 | 0.003 | Etv5 | ets variant gene 5 |
| 1434880 | at | 1.13 | 0.614 | Etv6 | ets variant gene 6 (TEL oncogene) |
| 1438325 | at | -1.10 | 0.010 | Evi1 | ecotropic viral integration site 1 |
| 1417513 | at | 1.07 | 0.449 | Evi5 | ecotropic viral integration site 5 |
| 1450106 | a_at | 1.14 | 0.060 | Evl | Ena-vasodilator stimulated phosphoprotein |
| 1434865 | a_at | 1.10 | 0.001 | Exoc7 | exocyst complex component 7 |
| 1458839 | at | -1.45 | 0.004 | Exoc8 | exocyst complex component 8 |
| 1452012 | a_at | 1.07 | 0.302 | Exosc1 | exosome component 1 |
| 1431502 | a_at | -1.04 | 0.674 | Exosc3 | exosome component 3 |
| 1424685 | at | -1.03 | 0.614 | Exosc4 | exosome component 4 |
| 1448365 | at | 1.08 | 0.179 | Exosc7 | exosome component 7 |
| 1428291 | at | 1.02 | 0.637 | Exosc8 | exosome component 8 |
| 1418462 | at | 1.00 | 0.968 | Exosc9 | exosome component 9 |
| 1423305 | at | 1.08 | 0.347 | Extl1 | exostoses (multiple)-like 1 |
| 1434703 | at | -1.09 | 0.120 | Extl3 | exostoses (multiple)-like 3 |
| 1418290 | a_at | -1.12 | 0.150 | Ezh1 | enhancer of zeste homolog 1 (Drosophila) |
| 1416544 | at | -1.05 | 0.245 | Ezh2 | enhancer of zeste homolog 2 (Drosophila) |
| 1424595 | at | -1.12 | 0.221 | F11r | F11 receptor |
| 1448929 | at | 1.30 | 0.004 | F13a1 | coagulation factor XIII, A1 subunit |
| 1417408 | at | -1.16 | 0.186 | F3 | coagulation factor III |
| 1416023 | at | -1.01 | 0.658 | Fabp3 | fatty acid binding protein 3, muscle and heart |

Supplementary table S1

| | | | | | |
|---|---|---|---|---|---|
| 1423680 | at | 1.31 | 0.003 | Fads1 | fatty acid desaturase 1 |
| 1435910 | at | 1.43 | 0.035 | Fads3 | fatty acid desaturase 3 |
| 1418242 | at | 1.04 | 0.130 | Faf1 | Fas-associated factor 1 |
| 1417220 | at | -1.37 | 0.004 | Fah | fumarylacetoacetate hydrolase |
| 1419547 | at | -1.09 | 0.192 | Fahd1 | fumarylacetoacetate hydrolase domain containing 1 |
| 1418029 | at | 1.25 | 0.082 | Faim | Fas apoptotic inhibitory molecule |
| 1423624 | at | -1.08 | 0.201 | Fancl | Fanconi anemia, complementation group L |
| 1417552 | at | 1.17 | 0.017 | Fap | fibroblast activation protein |
| 1452280 | at | 1.42 | 0.000 | Farp1 | FERM, RhoGEF (Arhgef) and pleckstrin domain protein 1 (chondrocyte-derived) |
| 1431354 | a_at | -1.18 | 0.095 | Fars2 | phenylalanine-tRNA synthetase 2 (mitochondrial) |
| 1424907 | a_at | -1.07 | 0.678 | Farsla | phenylalanine-tRNA synthetase-like, alpha subunit |
| 1423708 | a_at | -1.07 | 0.139 | Farslb | phenylalanine-tRNA synthetase-like, beta subunit |
| 1460251 | at | -1.30 | 0.008 | Fas | Fas (TNF receptor superfamily member) |
| 1460635 | at | -1.19 | 0.007 | Fastk | Fas-activated serine/threonine kinase |
| 1433857 | at | 1.32 | 0.001 | Fath | fat tumor suppressor homolog (Drosophila) |
| 1417452 | a_at | 1.04 | 0.123 | Fau | Finkel-Biskis-Reilly murine sarcoma virus (FBR-MuSV) ubiquitously expressed (fox derived) |
| 1425979 | a_at | 1.01 | 0.896 | Fbf1 | Fas (TNFRSF6) binding factor 1 |
| 1416684 | at | 1.18 | 0.048 | Fbl | fibrillarin |
| 1416164 | at | 1.83 | 0.000 | Fbln5 | fibulin 5 |
| 1460208 | at | 2.42 | 0.000 | Fbn1 | fibrillin 1 |
| 1449088 | at | -2.83 | 0.000 | Fbp2 | fructose bisphosphatase 2 |
| 1452198 | at | -1.04 | 0.441 | Fbxl10 | F-box and leucine-rich repeat protein 10 |
| 1435329 | at | -1.04 | 0.435 | Fbxl11 | F-box and leucine-rich repeat protein 11 |
| 1448420 | a_at | -1.07 | 0.483 | Fbxl12 | F-box and leucine-rich repeat protein 12 |
| 1417407 | at | -1.00 | 0.987 | Fbxl14 | F-box and leucine-rich repeat protein 14 |
| 1452860 | at | -1.04 | 0.483 | Fbxl17 | F-box and leucine-rich repeat protein 17 |
| 1454635 | at | -1.01 | 0.861 | Fbxl3 | F-box and leucine-rich repeat protein 3 |
| 1456038 | at | 1.13 | 0.181 | Fbxl4 | F-box and leucine-rich repeat protein 4 |
| 1433441 | at | -1.11 | 0.266 | Fbxl5 | F-box and leucine-rich repeat protein 5 |
| 1440441 | at | -1.22 | 0.027 | Fbxl7 | F-box and leucine-rich repeat protein 7 |
| 1452153 | at | -1.10 | 0.078 | Fbxo18 | F-box protein 18 |
| 1452828 | at | -1.37 | 0.004 | Fbxo21 | F-box only protein 21 |
| 1426592 | a_at | 1.13 | 0.939 | Fbxo22 | F-box only protein 22 |
| 1417535 | at | -1.03 | 0.569 | Fbxo25 | F-box only protein 25 |
| 1431811 | a_at | -1.22 | 0.036 | Fbxo34 | F-box only protein 34 |
| 1452255 | at | 1.06 | 0.647 | Fbxo38 | F-box protein 38 |
| 1427121 | at | -1.11 | 0.177 | Fbxo4 | F-box only protein 4 |
| 1435356 | at | -1.05 | 0.573 | Fbxo42 | F-box protein 42 |
| 1434455 | at | -1.05 | 0.439 | Fbxo44 | : F-box protein 44 |
| 1452895 | at | 1.02 | 0.775 | Fbxo45 | F-box protein 45 |
| 1417501 | at | 1.02 | 0.709 | Fbxo6b | F-box only protein 6b |

Supplementary table S1

| | | | | | |
|---|---|---|---|---|---|
| 1452739 | at | -1.04 | 0.519 | Fbxo7 | F-box only protein 7 |
| 1417480 | at | -1.06 | 0.399 | Fbxo9 | f-box only protein 9 |
| 1425462 | at | 1.08 | 0.101 | Fbxw11 | F-box and WD-40 domain protein 11 |
| 1451524 | at | 1.11 | 0.156 | Fbxw2 | F-box and WD-40 domain protein 2 |
| 1417226 | at | 1.01 | 0.912 | Fbxw4 | F-box and WD-40 domain protein 4 |
| 1417476 | at | 1.13 | 0.023 | Fbxw5 | F-box and WD-40 domain protein 5 |
| 1451558 | at | -1.36 | 0.000 | Fbxw7 | F-box and WD-40 domain protein 7, archipelago homolog (Drosophila) |
| 1426944 | at | -1.16 | 0.208 | Fbxw8 | F-box and WD-40 domain protein 8 |
| 1418340 | at | 1.29 | 0.003 | Fcer1g | Fc receptor, IgE, high affinity I, gamma polypeptide |
| 1448620 | at | 1.38 | 0.003 | Fcgr3 | Fc receptor, IgG, low affinity III |
| 1416978 | at | -1.09 | 0.011 | Fcgrt | Fc receptor, IgG, alpha chain transporter |
| 1437200 | at | -1.06 | 0.413 | Fcho2 | FCH domain only 2 |
| 1434260 | at | 1.27 | 0.001 | Fchsd2 | FCH and double SH3 domains 2 |
| 1435801 | at | -1.28 | 0.030 | Fcmd | Fukuyama type congenital muscular dystrophy homolog (human) |
| 1435258 | at | -1.15 | 0.202 | Fcna | ficolin A |
| 1448130 | at | -1.49 | 0.001 | Fdft1 | farnesyl diphosphate farnesyl transferase 1 |
| 1449108 | at | -1.10 | 0.089 | Fdx1 | ferredoxin 1 |
| 1449181 | at | 1.04 | 0.687 | Fech | ferrochelatase |
| 1423583 | at | 1.06 | 0.415 | Fem1a | feminization 1 homolog a (C. elegans) |
| 1418324 | at | -1.01 | 0.704 | Fem1b | feminization 1 homolog b (C. elegans) |
| 1421729 | a_at | -1.07 | 0.161 | Fert2 | fer (fms/fps related) protein kinase, testis specific 2 |
| 1434348 | at | 1.25 | 0.000 | Fez2 | fasciculation and elongation protein zeta 2 (zygin II) |
| 1455337 | at | -1.40 | 0.002 | Fgd4 | FYVE, RhoGEF and PH domain containing 4 |
| 1460578 | at | 1.09 | 0.175 | Fgd5 | FYVE, RhoGEF and PH domain containing 5 |
| 1435467 | at | -1.18 | 0.024 | Fgd6 | FYVE, RhoGEF and PH domain containing 6 |
| 1444344 | at | 1.10 | 0.627 | Fgf10 | fibroblast growth factor 10 |
| 1418497 | at | -1.31 | 0.000 | Fgf13 | fibroblast growth factor 13 |
| 1449545 | at | 1.02 | 0.867 | Fgf18 | fibroblast growth factor 18 |
| 1456945 | at | -1.01 | 0.933 | Fgf2 | fibroblast growth factor 2 |
| 1420795 | at | -1.10 | 0.470 | Fgf9 | fibroblast growth factor 9 |
| 1428919 | at | 1.01 | 0.669 | Fgfr1op | Fgfr1 oncogene partner |
| 1423591 | at | -1.06 | 0.377 | Fgfr1op2 | FGFR1 oncogene partner 2 |
| 1451912 | a_at | 1.04 | 0.439 | Fgfrl1 | fibroblast growth factor receptor-like 1 |
| 1421855 | at | 3.17 | 0.000 | Fgl2 | fibrinogen-like protein 2 |
| 1424828 | a_at | -1.07 | 0.038 | Fh1 | fumarate hydratase 1 |
| 1417872 | at | 1.59 | 0.000 | Fhl1 | four and a half LIM domains 1 |
| 1419184 | a_at | 1.09 | 0.007 | Fhl2 | four and a half LIM domains 2 |
| 1435557 | at | -1.01 | 0.921 | Fhod1 | formin homology 2 domain containing 1 |
| 1435551 | at | -1.09 | 0.079 | FHOS2 | formin-family protein FHOS2 |
| 1422516 | a_at | 1.14 | 0.102 | Fibp | fibroblast growth factor (acidic) intracellular binding protein |
| 1438953 | at | 1.12 | 0.122 | Figf | c-fos induced growth factor |

Supplementary table S1

| | | | | | |
|---|---|---|---|---|---|
| 1450268 | at | -1.84 | 0.008 | Fign | fidgetin |
| 1423042 | at | 1.07 | 0.107 | Fin14 | fibroblast growth factor inducible 14 |
| 1421534 | at | 1.12 | 0.125 | Fin15 | fibroblast growth factor inducible 15 |
| 1428280 | at | -1.02 | 0.425 | Fip1l1 | FIP1 like 1 (S. cerevisiae) |
| 1423733 | a_at | -1.16 | 0.111 | Fiz1 | Flt3 interacting zinc finger protein 1 |
| 1433481 | at | -1.00 | 0.975 | Fkbp14 | FK506 binding protein 14 |
| 1416036 | at | 1.24 | 0.010 | Fkbp1a | FK506 binding protein 1a |
| 1450694 | at | -1.28 | 0.001 | Fkbp2 | FK506 binding protein 2 |
| 1416362 | a_at | -1.30 | 0.005 | Fkbp4 | FK506 binding protein 4 |
| 1416125 | at | 1.39 | 0.013 | Fkbp5 | FK506 binding protein 5 |
| 1416803 | at | -1.01 | 0.707 | Fkbp7 | FK506 binding protein 7 |
| 1416113 | at | 1.12 | 0.111 | Fkbp8 | FK506 binding protein 8 |
| 1423677 | at | 1.09 | 0.012 | Fkbp9 | FK506 binding protein 9 |
| 1436812 | at | -1.18 | 0.005 | Fkrp | fukutin related protein |
| 1423663 | at | -1.56 | 0.000 | Flcn | folliculin |
| 1433512 | at | 1.03 | 0.725 | Fli1 | Friend leukemia integration 1 |
| 1416044 | at | 1.02 | 0.667 | Fliih | flightless I homolog (Drosophila) |
| 1428346 | at | 1.02 | 0.623 | Fln29 | FLN29 gene product |
| 1426677 | at | 1.12 | 0.208 | Flna | filamin, alpha |
| 1426750 | at | 1.25 | 0.042 | Flnb | filamin, beta |
| 1449073 | at | 1.97 | 0.000 | Flnc | filamin C, gamma (actin binding protein 280) |
| 1448559 | at | 1.04 | 0.412 | Flot1 | flotillin 1 |
| 1417544 | a_at | -1.13 | 0.185 | Flot2 | flotillin 2 |
| 1440926 | at | -1.13 | 0.190 | Flt1 | FMS-like tyrosine kinase 1 |
| 1421442 | at | 1.11 | 0.287 | Flt4 | FMS-like tyrosine kinase 4 |
| 1426982 | at | 1.09 | 0.294 | Flywch1 | FLYWCH-type zinc finger 1 |
| 1452672 | at | -1.02 | 0.801 | Fmip | Fms interacting protein |
| 1428579 | at | 1.08 | 0.066 | Fmnl2 | formin-like 2 |
| 1426825 | at | 1.29 | 0.093 | Fmnl3 | formin-like 3 |
| 1417429 | at | -1.08 | 0.322 | Fmo1 | flavin containing monooxygenase 1 |
| 1422904 | at | -1.02 | 0.811 | Fmo2 | flavin containing monooxygenase 2 |
| 1440899 | at | -1.04 | 0.332 | Fmo5 | flavin containing monooxygenase 5 |
| 1423369 | at | 1.09 | 0.112 | Fmr1 | fragile X mental retardation syndrome 1 homolog |
| 1426642 | at | 2.00 | 0.001 | Fn1 | fibronectin 1 |
| 1418311 | at | -1.51 | 0.000 | Fn3k | fructosamine 3 kinase |
| 1453746 | at | 1.12 | 0.333 | Fnbp1 | formin binding protein 1 |
| 1434339 | at | -1.29 | 0.014 | Fnbp1l | formin binding protein 1-like |
| 1438222 | at | 1.02 | 0.846 | Fnbp3 | formin binding protein 3 |
| 1423191 | at | -1.05 | 0.448 | Fnbp4 | formin binding protein 4 |
| 1453321 | at | 1.96 | 0.000 | Fndc1 | fibronectin type III domain containing 1 |
| 1435115 | at | -1.29 | 0.001 | Fndc5 | fibronectin type III domain containing 5 |

Supplementary table S1

| | | | | | |
|---|---|---|---|---|---|
| 1417465 | at | -1.05 | 0.329 | Fnta | farnesyltransferase, CAAX box, alpha |
| 1434309 | at | 1.03 | 0.581 | Fntb | farnesyltransferase, CAAX box, beta |
| 1451648 | a_at | 1.04 | 0.607 | Folr2 | folate receptor 2 (fetal) |
| 1423100 | at | -1.03 | 0.898 | Fos | FBJ osteosarcoma oncogene |
| 1437247 | at | 1.40 | 0.035 | Fosl2 | fos-like antigen 2 |
| 1419485 | at | -1.08 | 0.289 | Foxc1 | forkhead box C1 |
| 1416693 | at | 1.97 | 0.001 | Foxc2 | forkhead box C2 |
| 1420374 | at | -1.13 | 0.170 | Foxj2 | forkhead box J2 |
| 1435362 | at | 1.00 | 0.958 | Foxj3 | forkhead box J3 |
| 1434655 | at | -1.06 | 0.450 | Foxk1 | forkhead box K1 |
| 1454831 | at | 1.04 | 0.575 | Foxn2 | forkhead box N2 |
| 1416982 | at | 1.06 | 0.457 | Foxo1 | forkhead box O1 |
| 1434832 | at | -1.30 | 0.002 | Foxo3a | forkhead box O3a |
| 1438231 | at | 1.01 | 0.952 | Foxp2 | forkhead box P2 |
| 1429519 | at | 1.01 | 0.858 | Fpgt | fucose-1-phosphate guanylyltransferase |
| 1424614 | at | 1.33 | 0.379 | Frag1 | FGF receptor activating protein 1 |
| 1417592 | at | 1.18 | 0.038 | Frap1 | FK506 binding protein 12-rapamycin associated protein 1 |
| 1417253 | at | -1.14 | 0.015 | Frg1 | FSHD region gene 1 |
| 1426569 | a_at | -1.08 | 0.539 | Frk | fyn-related kinase |
| 1434285 | at | 1.12 | 0.001 | Frmd4a | FERM domain containing 4A |
| 1438169 | a_at | 1.26 | 0.037 | Frmd4b | FERM domain containing 4B |
| 1437891 | at | -1.09 | 0.186 | Frs2 | fibroblast growth factor receptor substrate 2 |
| 1448378 | at | 1.41 | 0.150 | Fscn1 | fascin homolog 1, actin bundling protein (Strongylocentrotus) purpuratus) |
| 1416221 | at | 2.31 | 0.000 | Fstl1 | follistatin-like 1 |
| 1448771 | a_at | 1.03 | 0.337 | Fth1 | ferritin heavy chain 1 |
| 1418364 | a_at | 1.10 | 0.171 | Ftl1 | ferritin light chain 1 |
| 1448223 | at | -1.13 | 0.189 | Fto | fatso |
| 1423364 | a_at | 1.02 | 0.832 | Fts | fused toes |
| 1417099 | at | -1.02 | 0.815 | Ftsj1 | Ftsj homolog 1 (E. coli) |
| 1448969 | at | -1.08 | 0.298 | Ftsj2 | FtsJ homolog 2 (E. coli) |
| 1451026 | at | 1.27 | 0.045 | Ftsj3 | FtsJ homolog 3 (E. coli) |
| 1437543 | at | 1.17 | 0.415 | Fubp1 | far upstream element (FUSE) binding protein 1 |
| 1416109 | at | 1.05 | 0.270 | Fuca1 | fucosidase, alpha-L- 1, tissue |
| 1416046 | a_at | 1.40 | 0.000 | Fuca2 | fucosidase, alpha-L- 2, plasma |
| 1418518 | at | -1.06 | 0.244 | Furin | furin (paired basic amino acid cleaving enzyme) |
| 1434123 | at | -1.02 | 0.775 | Fut11 | fucosyltransferase 11 |
| 1460319 | at | -1.06 | 0.230 | Fut8 | fucosyltransferase 8 |
| 1428559 | at | 1.26 | 0.004 | Fvt1 | follicular lymphoma variant translocation 1 |
| 1427282 | a_at | -1.19 | 0.001 | Fxn | frataxin |
| 1422522 | at | 1.12 | 0.184 | Fxr2h | fragile X mental retardation gene 2, autosomal homolog |
| 1421374 | a_at | -1.10 | 0.001 | Fxyd1 | FXYD domain-containing ion transport regulator 1 |

Supplementary table S1

| | | | | | |
|---|---|---|---|---|---|
| 1418296 | at | 1.75 | 0.004 | Fxyd5 | FXYD domain-containing ion transport regulator 5 |
| 1417343 | at | 2.49 | 0.000 | Fxyd6 | FXYD domain-containing ion transport regulator 6 |
| 1435144 | at | 1.22 | 0.023 | Fyb | FYN binding protein |
| 1453424 | at | -1.12 | 0.370 | Fyco1 | FYVE and coiled-coil domain containing 1 |
| 1448765 | at | -1.01 | 0.801 | Fyn | Fyn proto-oncogene |
| 1451325 | at | 1.04 | 0.707 | Fyttd1 | forty-two-three domain containing 1 |
| 1437284 | at | 1.64 | 0.001 | Fzd1 | frizzled homolog 1 (Drosophila) |
| 1418534 | at | 1.53 | 0.002 | Fzd2 | frizzled homolog 2 (Drosophila) |
| 1419301 | at | -1.06 | 0.360 | Fzd4 | frizzled homolog 4 (Drosophila) |
| 1417301 | at | -1.17 | 0.032 | Fzd6 | frizzled homolog 6 (Drosophila) |
| 1450044 | at | 1.16 | 0.151 | Fzd7 | frizzled homolog 7 (Drosophila) |
| 1419451 | at | -1.06 | 0.555 | Fzr1 | fizzy/cell division cycle 20 related 1 (Drosophila) |
| 1448700 | at | -1.50 | 0.016 | G0s2 | G0/G1 switch gene 2 |
| 1428245 | at | 1.03 | 0.598 | G6pc3 | glucose 6 phosphatase, catalytic, 3 |
| 1448354 | at | 1.39 | 0.007 | G6pdx | glucose-6-phosphate dehydrogenase X-linked |
| 1419428 | a_at | -1.22 | 0.050 | Gaa | glucosidase, alpha, acid |
| 1417694 | at | -1.33 | 0.020 | Gab1 | growth factor receptor bound protein 2-associated protein 1 |
| 1446860 | at | 1.47 | 0.074 | Gab2 | growth factor receptor bound protein 2-associated protein 2 |
| 1416937 | at | -1.01 | 0.847 | Gabarap | gamma-aminobutyric acid receptor associated protein |
| 1416418 | at | 1.11 | 0.077 | Gabarapl1 | gamma-aminobutyric acid (GABA(A)) receptor-associated protein-like 1 |
| 1423187 | at | 1.04 | 0.336 | Gabarapl2 | gamma-aminobutyric acid (GABA-A) receptor-associated protein-like 2 |
| 1455021 | at | -1.02 | 0.744 | Gabbr1 | gamma-aminobutyric acid (GABA-B) receptor, 1 |
| 1450665 | at | -1.10 | 0.187 | Gabpa | GA repeat binding protein, alpha |
| 1455110 | at | 1.03 | 0.500 | Gabpb2 | GA repeat binding protein, beta 2 |
| 1449519 | at | -1.25 | 0.191 | Gadd45a | growth arrest and DNA-damage-inducible 45 alpha |
| 1453851 | a_at | 1.66 | 0.042 | Gadd45g | growth arrest and DNA-damage-inducible 45 gamma |
| 1417619 | at | -1.28 | 0.001 | Gadd45gip1 | growth arrest and DNA-damage-inducible, gamma interacting protein 1 |
| 1426703 | at | -1.05 | 0.072 | Gak | cyclin G associated kinase |
| 1452907 | at | 1.32 | 0.005 | Galc | galactosylceramidase |
| 1455798 | at | 1.23 | 0.005 | Galk2 | galactokinase 2 |
| 1452582 | at | -1.41 | 0.007 | Galm | galactose mutarotase |
| 1424431 | at | 1.03 | 0.589 | Galnact2 | chondroitin sulfate GalNAcT-2 |
| 1423237 | at | 1.08 | 0.097 | Galnt1 | UDP-N-acetyl-alpha-D-galactosamine:polypeptide N-acetylgalactosaminyltransferase 1 |
| 1424748 | at | -1.08 | 0.180 | Galnt11 | UDP-N-acetyl-alpha-D-galactosamine:polypeptide N-acetylgalactosaminyltransferase 11 |
| 1452182 | at | -1.06 | 0.411 | Galnt2 | UDP-N-acetyl-alpha-D-galactosamine:polypeptide N-acetylgalactosaminyltransferase 2 |
| 1452232 | at | 1.01 | 0.917 | Galnt7 | UDP-N-acetyl-alpha-D-galactosamine: polypeptide N-acetylgalactosaminyltransferase 7 |
| 1429235 | at | -1.18 | 0.187 | Galntl2 | UDP-N-acetyl-alpha-D-galactosamine:polypeptide N-acetylgalactosaminyltransferase-like 2 |
| 1450187 | a_at | -1.14 | 0.062 | Galt | galactose-1-phosphate uridyl transferase |
| 1415787 | at | -1.09 | 0.059 | Ganab | alpha glucosidase 2 alpha neutral subunit |
| 1429454 | at | -1.05 | 0.425 | Gapvd1 | GTPase activating protein and VPS9 domains 1 |
| 1458685 | at | -1.18 | 0.347 | Garnl1 | GTPase activating RANGAP domain-like 1 |

Supplementary table S1

| | | | | | |
|---|---|---|---|---|---|
| 1423784 | at | 1.23 | 0.077 | Gars | glycyl-tRNA synthetase |
| 1448494 | at | -1.23 | 0.041 | Gas1 | growth arrest specific 1 |
| 1434928 | at | 1.06 | 0.414 | Gas2l1 | growth arrest-specific 2 like 1 |
| 1453416 | at | 2.28 | 0.000 | Gas2l3 | growth arrest-specific 2 like 3 |
| 1436222 | at | 1.35 | 0.012 | Gas5 | growth arrest specific 5 |
| 1417399 | at | -1.01 | 0.892 | Gas6 | growth arrest specific 6 |
| 1450333 | a_at | -1.12 | 0.327 | Gata2 | GATA binding protein 2 |
| 1418863 | at | 1.12 | 0.276 | Gata4 | GATA binding protein 4 |
| 1425464 | at | -1.57 | 0.012 | Gata6 | GATA binding protein 6 |
| 1452085 | at | -1.02 | 0.833 | Gatad1 | GATA zinc finger domain containing 1 |
| 1423992 | at | 1.04 | 0.445 | Gatad2a | GATA zinc finger domain containing 2A |
| 1425075 | at | -1.01 | 0.860 | Gatad2b | GATA zinc finger domain containing 2B |
| 1437252 | at | -1.18 | 0.106 | Gats | opposite strand transcription unit to Stag3 |
| 1437044 | a_at | 1.37 | 0.005 | Gba | glucosidase, beta, acid |
| 1419484 | a_at | -1.07 | 0.036 | Gbas | glioblastoma amplified sequence |
| 1420654 | a_at | 1.27 | 0.013 | Gbe1 | glucan (1,4-alpha-), branching enzyme 1 |
| 1423421 | at | -1.03 | 0.738 | Gbif | globin inducing factor, fetal |
| 1418240 | at | 1.06 | 0.739 | Gbp2 | guanylate nucleotide binding protein 2 |
| 1418392 | a_at | 1.20 | 0.315 | Gbp4 | guanylate nucleotide binding protein 4 |
| 1451451 | at | -1.32 | 0.023 | Gca | grancalcin |
| 1453240 | a_at | 1.12 | 0.110 | Gcap14 | granule cell antiserum positive 14 |
| 1417823 | at | -1.99 | 0.000 | Gcat | glycine C-acetyltransferase (2-amino-3-ketobutyrate-coenzyme A ligase) |
| 1429033 | at | 1.22 | 0.014 | Gcc1 | golgi coiled coil 1 |
| 1448717 | at | -1.34 | 0.000 | Gcdh | glutaryl-Coenzyme A dehydrogenase |
| 1420499 | at | -1.02 | 0.835 | Gch1 | GTP cyclohydrolase 1 |
| 1417113 | at | -1.05 | 0.530 | Gcl | germ cell-less homolog (Drosophila) |
| 1439050 | at | 1.09 | 0.518 | Gclm | glutamate-cysteine ligase , modifier subunit |
| 1433713 | at | -1.16 | 0.233 | Gcn1l1 | GCN1 general control of amino-acid synthesis 1-like 1 (yeast) |
| 1426783 | at | 1.08 | 0.435 | Gcn5l2 | GCN5 general control of amino acid synthesis-like 2 (yeast) |
| 1460431 | at | 1.32 | 0.057 | Gcnt1 | glucosaminyl (N-acetyl) transferase 1, core 2 |
| 1425503 | at | 1.26 | 0.015 | Gcnt2 | glucosaminyl (N-acetyl) transferase 2, I-branching enzyme |
| 1422489 | at | -1.07 | 0.320 | Gcs1 | glucosidase 1 |
| 1432164 | a_at | 1.10 | 0.028 | Gcsh | glycine cleavage system protein H (aminomethyl carrier) |
| 1435749 | at | 1.08 | 0.462 | Gda | guanine deaminase |
| 1420342 | at | -2.07 | 0.001 | Gdap10 | ganglioside-induced differentiation-associated-protein 10 |
| 1424057 | at | 1.13 | 0.018 | Gdap2 | ganglioside-induced differentiation-associated-protein 2 |
| 1451070 | at | 1.06 | 0.337 | Gdi1 | guanosine diphosphate (GDP) dissociation inhibitor 1 |
| 1424076 | at | -1.39 | 0.000 | Gdpd1 | glycerophosphodiester phosphodiesterase domain containing 1 |
| 1424150 | at | -1.12 | 0.329 | Gdpd5 | glycerophosphodiester phosphodiesterase domain containing 5 |
| 1455453 | at | -1.06 | 0.632 | Gemin5 | gem (nuclear organelle) associated protein 5 |
| 1435764 | a_at | -1.02 | 0.617 | Gemin7 | gem (nuclear organelle) associated protein 7 |

Supplementary table S1

| | | | | | |
|---|---|---|---|---|---|
| 1452272 | a_at | 1.13 | 0.118 | Gfer | growth factor, erv1 (S. cerevisiae)-like (augmenter of liver regeneration) |
| 1448381 | at | -1.20 | 0.179 | Gfm1 | G elongation factor 1 |
| 1426590 | at | -1.01 | 0.786 | Gfm2 | G elongation factor, mitochondrial 2 |
| 1418753 | at | 1.25 | 0.133 | Gfpt2 | glutamine fructose-6-phosphate transaminase 2 |
| 1428141 | at | -1.02 | 0.710 | Gga2 | golgi associated, gamma adaptin ear containing, ARF binding protein 2 |
| 1438008 | at | 1.08 | 0.178 | Gga3 | golgi associated, gamma adaptin ear containing, ARF binding protein 3 |
| 1419595 | a_at | 1.19 | 0.001 | Ggh | gamma-glutamyl hydrolase |
| 1429125 | at | 1.11 | 0.109 | Ggnbp1 | gametogenetin binding protein 1 |
| 1426011 | a_at | 1.06 | 0.140 | Ggnbp2 | gametogenetin binding protein 2 |
| 1419506 | at | -1.07 | 0.409 | Ggps1 | geranylgeranyl diphosphate synthase 1 |
| 1418483 | a_at | 1.18 | 0.031 | Ggta1 | glycoprotein galactosyltransferase alpha 1, 3 |
| 1425918 | at | -1.42 | 0.155 | Ghr | growth hormone receptor |
| 1417848 | at | 1.25 | 0.000 | Gig1 | glucocorticoid induced gene 1 |
| 1424374 | at | 1.19 | 0.114 | Gimap4 | GTPase, IMAP family member 4 |
| 1454759 | at | -1.05 | 0.545 | Git1 | G protein-coupled receptor kinase-interactor 1 |
| 1415800 | at | 1.07 | 0.367 | Gja1 | gap junction membrane channel protein alpha 1 |
| 1439793 | at | -1.35 | 0.003 | Gja3 | gap junction membrane channel protein alpha 3 |
| 1416255 | at | 1.16 | 0.064 | Gja4 | gap junction membrane channel protein alpha 4 |
| 1449094 | at | 1.12 | 0.051 | Gja7 | gap junction membrane channel protein alpha 7 |
| 1417594 | at | -1.03 | 0.647 | Gkap1 | G kinase anchoring protein 1 |
| 1416205 | at | 1.08 | 0.427 | Glb1 | galactosidase, beta 1 |
| 1428603 | at | 1.09 | 0.253 | Glcci1 | glucocorticoid induced transcript 1 |
| 1428374 | at | -1.10 | 0.005 | Glce | glucuronyl C5-epimerase |
| 1426344 | at | 1.01 | 0.884 | Gle1l | GLE1 RNA export mediator-like (yeast |
| 1455154 | at | -1.01 | 0.914 | Gli3 | GLI-Kruppel family member GLI3 |
| 1416598 | at | 1.45 | 0.012 | Glis2 | GLIS family zinc finger 2 |
| 1436070 | at | -1.01 | 0.845 | Glo1 | glyoxalase 1 |
| 1416592 | at | -1.02 | 0.725 | Glrx1 | glutaredoxin 1 (thioltransferase) |
| 1437679 | a_at | -1.00 | 0.951 | Glrx2 | glutaredoxin 2 (thioltransferase) |
| 1433495 | at | 1.16 | 0.246 | Glt25d1 | glycosyltransferase 25 domain containing 1 |
| 1428270 | at | 1.02 | 0.739 | Glt8d1 | glycosyltransferase 8 domain containing 1 |
| 1451121 | a_at | -1.22 | 0.000 | Gltscr2 | glioma tumor suppressor candidate region gene 2 |
| 1416209 | at | -1.06 | 0.133 | Glud1 | glutamate dehydrogenase 1 |
| 1426236 | a_at | 1.18 | 0.611 | Glul | glutamate-ammonia ligase (glutamine synthase) |
| 1424519 | at | -1.16 | 0.021 | Gm169 | gene model 169, (NCBI) |
| 1455293 | at | -1.12 | 0.088 | Gm185 | gene model 185, (NCBI) |
| 1448241 | at | 1.05 | 0.481 | Gm2a | GM2 ganglioside activator protein |
| 1460558 | at | 1.07 | 0.255 | Gm631 | gene model 631, (NCBI) |
| 1436047 | at | 1.10 | 0.046 | Gm672 | gene model 672, (NCBI) |
| 1453070 | at | 1.21 | 0.198 | Gm78 | gene model 78, (NCBI) |
| 1458778 | at | 1.04 | 0.851 | Gm83 | gene model 83, (NCBI) |

Supplementary table S1

| | | | | | |
|---|---|---|---|---|---|
| 1429942 | at | 1.09 | 0.242 | Gmeb1 | glucocorticoid modulatory element binding protein 1 |
| 1454818 | at | -1.02 | 0.684 | Gmeb2 | glucocorticoid modulatory element binding protein 2 |
| 1419193 | a_at | 1.10 | 0.313 | Gmfg | glia maturation factor, gamma |
| 1417506 | at | -1.09 | 0.149 | Gmnn | geminin |
| 1417578 | a_at | -1.01 | 0.898 | Gmppa | GDP-mannose pyrophosphorylase A |
| 1439030 | at | 1.18 | 0.018 | Gmppb | GDP-mannose pyrophosphorylase B |
| 1448530 | at | -1.00 | 0.982 | Gmpr | guanosine monophosphate reductase |
| 1416356 | at | 1.25 | 0.005 | Gmpr2 | guanosine monophosphate reductase 2 |
| 1434254 | at | -1.06 | 0.488 | Gna11 | guanine nucleotide binding protein, alpha 11 |
| 1455008 | at | -1.07 | 0.223 | Gna12 | guanine nucleotide binding protein, alpha 12 |
| 1450656 | at | 1.05 | 0.671 | Gna13 | guanine nucleotide binding protein, alpha 13 |
| 1419449 | a_at | 1.02 | 0.788 | Gnai2 | guanine nucleotide binding protein, alpha inhibiting 2 |
| 1428645 | at | 1.09 | 0.281 | Gnai3 | guanine nucleotide binding protein, alpha inhibiting 3 |
| 1421152 | a_at | 1.47 | 0.000 | Gnao1 | guanine nucleotide binding protein, alpha o |
| 1429559 | at | 1.04 | 0.562 | Gnaq | guanine nucleotide binding protein, alpha q polypeptide |
| 1418646 | at | 1.17 | 0.223 | Gna-rs1 | guanine nucleotide binding protein, related sequence 1 |
| 1454696 | at | 1.01 | 0.852 | Gnb1 | guanine nucleotide binding protein, beta 1 |
| 1450623 | at | -1.22 | 0.081 | Gnb2 | guanine nucleotide binding protein, beta 2 |
| 1455168 | a_at | 1.02 | 0.569 | Gnb2-rs1 | guanine nucleotide binding protein, beta 2, related sequence 1 |
| 1449159 | at | 1.46 | 0.056 | Gnb3 | guanine nucleotide binding protein, beta 3 |
| 1419470 | at | 1.04 | 0.624 | Gnb4 | guanine nucleotide binding protein, beta 4 |
| 1422208 | a_at | 1.03 | 0.641 | Gnb5 | guanine nucleotide binding protein, beta 5 |
| 1448810 | at | -1.07 | 0.145 | Gne | glucosamine |
| 1450649 | at | 1.29 | 0.000 | Gng10 | guanine nucleotide binding protein (G protein), gamma 10 |
| 1448942 | at | -1.02 | 0.760 | Gng11 | guanine nucleotide binding protein (G protein), gamma 11 |
| 1421947 | at | 1.14 | 0.167 | Gng12 | guanine nucleotide binding protein (G protein), gamma 12 |
| 1428156 | at | 1.03 | 0.644 | Gng2 | guanine nucleotide binding protein (G protein), gamma 2 subunit |
| 1433656 | a_at | 1.31 | 0.015 | Gnl3 | guanine nucleotide binding protein-like 3 (nucleolar) |
| 1439009 | at | -1.14 | 0.143 | Gnl3l | guanine nucleotide binding protein-like 3 (nucleolar)-like |
| 1417456 | at | -1.11 | 0.003 | Gnpat | glyceronephosphate O-acyltransferase |
| 1426524 | at | -1.09 | 0.044 | Gnpda2 | glucosamine-6-phosphate deaminase 2 |
| 1423158 | at | 1.24 | 0.026 | Gnpnat1 | glucosamine-phosphate N-acetyltransferase 1 |
| 1436526 | at | -1.12 | 0.116 | Gnptg | N-acetylglucosamine-1-phosphotransferase, gamma subunit |
| 1433546 | at | -1.01 | 0.783 | Gns | glucosamine (N-acetyl)-6-sulfatase |
| 1448625 | at | 1.09 | 0.083 | Golga2 | golgi autoantigen, golgin subfamily a, 2 |
| 1419159 | at | 1.27 | 0.133 | Golga3 | golgi autoantigen, golgin subfamily a, 3 |
| 1448803 | at | 1.22 | 0.231 | Golga4 | golgi autoantigen, golgin subfamily a, 4 |
| 1418447 | at | 1.10 | 0.077 | Golga5 | golgi autoantigen, golgin subfamily a, 5 |
| 1415672 | at | 1.31 | 0.000 | Golga7 | golgi autoantigen, golgin subfamily a, 7 |
| 1415698 | at | 1.52 | 0.000 | Golph2 | golgi phosphoprotein 2 |
| 1420117 | at | -1.04 | 0.191 | Golph3 | golgi phosphoprotein 3 |

Supplementary table S1

| | | | | | |
|---|---|---|---|---|---|
| 1428875 | at | 1.45 | 0.000 | Golph4 | golgi phosphoprotein 4 |
| 1449859 | at | 1.02 | 0.806 | Golt1b | golgi transport 1 homolog B (S. cerevisiae) |
| 1453221 | at | 1.37 | 0.008 | Gopc | golgi associated PDZ and coiled-coil motif containing |
| 1451671 | at | -1.02 | 0.830 | Gorasp1 | golgi reassembly stacking protein 1 |
| 1460460 | a_at | 1.12 | 0.975 | Gorasp2 | golgi reassembly stacking protein 2 |
| 1416215 | at | -1.09 | 0.128 | Gosr1 | golgi SNAP receptor complex member 1 |
| 1450970 | at | -1.11 | 0.153 | Got1 | glutamate oxaloacetate transaminase 1, soluble |
| 1417716 | at | -1.32 | 0.100 | Got2 | glutamate oxaloacetate transaminase 2, mitochondrial |
| 1419309 | at | 2.01 | 0.000 | Gp38 | glycoprotein 38 |
| 1419499 | at | 1.19 | 0.000 | Gpam | glycerol-3-phosphate acyltransferase, mitochondrial |
| 1419013 | at | 1.02 | 0.898 | Gpatc1 | G patch domain containing 1 |
| 1458566 | at | -1.18 | 0.172 | Gpatc2 | G patch domain containing 2 |
| 1449390 | at | 1.50 | 0.017 | Gpatc4 | G patch domain containing 4 |
| 1417389 | at | 1.05 | 0.655 | Gpc1 | glypican 1 |
| 1428774 | at | 1.56 | 0.000 | Gpc6 | glypican 6 |
| 1448249 | at | 1.33 | 0.026 | Gpd1 | glycerol-3-phosphate dehydrogenase 1 (soluble) |
| 1438195 | at | -1.06 | 0.331 | Gpd1l | glycerol-3-phosphate dehydrogenase 1-like |
| 1420997 | a_at | -1.01 | 0.708 | Gpi1 | glucose phosphate isomerase 1 |
| 1453022 | at | -1.05 | 0.589 | Gpihbp1 | GPI-anchored HDL-binding protein 1 |
| 1434344 | at | 1.13 | 0.164 | Gpkow | G patch domain and KOW motifs |
| 1418050 | at | -1.14 | 0.103 | Gpld1 | glycosylphosphatidylinositol specific phospholipase D1 |
| 1423091 | a_at | 1.32 | 0.001 | Gpm6b | glycoprotein m6b |
| 1426078 | a_at | -1.12 | 0.265 | Gpr108 | G protein-coupled receptor 108 |
| 1426782 | at | 1.05 | 0.433 | Gpr125 | G protein-coupled receptor 125 |
| 1437408 | at | -1.25 | 0.161 | Gpr126 | G protein-coupled receptor 126 |
| 1454685 | at | -1.06 | 0.432 | Gpr146 | G protein-coupled receptor 146 |
| 1426973 | at | 1.46 | 0.003 | Gpr153 | G protein-coupled receptor 153 |
| 1452353 | at | -1.37 | 0.005 | Gpr155 | G protein-coupled receptor 155 |
| 1460276 | a_at | 1.09 | 0.242 | Gpr175 | G protein-coupled receptor 175 |
| 1434673 | at | -3.22 | 0.000 | Gpr22 | G protein-coupled receptor 22 |
| 1434848 | at | -1.15 | 0.109 | Gpr27 | G protein-coupled receptor 27 |
| 1433891 | at | 1.07 | 0.397 | Gpr48 | G protein-coupled receptor 48 |
| 1421118 | a_at | 1.12 | 0.135 | Gpr56 | G protein-coupled receptor 56 |
| 1424696 | at | 1.11 | 0.055 | Gpr89 | G protein-coupled receptor 89 |
| 1424455 | at | -1.09 | 0.115 | Gprasp1 | G protein-coupled receptor associated sorting protein 1 |
| 1424613 | at | 1.41 | 0.020 | Gprc5b | G protein-coupled receptor, family C, group 5, member B |
| 1436672 | at | 1.33 | 0.037 | Gprk5 | G protein-coupled receptor kinase 5 |
| 1451672 | at | -1.03 | 0.658 | Gprk6 | G protein-coupled receptor kinase 6 |
| 1415699 | a_at | 1.02 | 0.866 | Gps1 | G protein pathway suppressor 1 |
| 1417796 | at | -1.00 | 0.991 | Gps2 | G protein pathway suppressor 2 |
| 1423689 | a_at | -1.15 | 0.021 | Gpsm1 | G-protein signalling modulator 1 (AGS3-like, C. elegans) |

Supplementary table S1

| | | | | | |
|---|---|---|---|---|---|
| 1429681 | a_at | -1.09 | 0.015 | Gpsn2 | glycoprotein, synaptic 2 |
| 1434542 | at | -1.78 | 0.000 | Gpt2 | glutamic pyruvate transaminase (alanine aminotransferase) 2 |
| 1460671 | at | 1.47 | 0.003 | Gpx1 | glutathione peroxidase 1 |
| 1449106 | at | 1.48 | 0.000 | Gpx3 | glutathione peroxidase 3 |
| 1451695 | a_at | 1.01 | 0.860 | Gpx4 | glutathione peroxidase 4 |
| 1417836 | at | 1.08 | 0.143 | Gpx7 | glutathione peroxidase 7 |
| 1425458 | a_at | 1.14 | 0.001 | Grb10 | growth factor receptor bound protein 10 |
| 1417673 | at | -1.30 | 0.000 | Grb14 | growth factor receptor bound protein 14 |
| 1449111 | a_at | -1.21 | 0.025 | Grb2 | growth factor receptor bound protein 2 |
| 1416522 | a_at | -1.43 | 0.000 | Grcc10 | gene rich cluster, C10 gene |
| 1416393 | at | -1.05 | 0.186 | Grcc2f | gene rich cluster, C2f gene |
| 1423960 | at | -1.22 | 0.030 | Grcc3f | gene rich cluster, C3f gene |
| 1417772 | at | 1.10 | 0.023 | Grhpr | glyoxylate reductase/hydroxypyruvate reductase |
| 1453098 | at | -1.09 | 0.682 | Gria2 | glutamate receptor, ionotropic, AMPA2 (alpha 2) |
| 1438223 | at | 1.01 | 0.981 | Grid2 | glutamate receptor, ionotropic, delta 2 |
| 1445090 | at | 1.27 | 0.433 | Grik2 | glutamate receptor, ionotropic, kainate 2 (beta 2) |
| 1436297 | a_at | -1.00 | 0.965 | Grina | glutamate receptor, ionotropic, N-methyl D-asparate-associated protein 1 (glutamate binding) |
| 1431390 | a_at | -1.00 | 0.917 | Grinl1a | glutamate receptor, ionotropic, N-methyl D-aspartate-like 1A |
| 1417315 | at | -1.03 | 0.573 | Gripap1 | GRIP1 associated protein 1 |
| 1440545 | at | -1.11 | 0.361 | Grm1 | glutamate receptor, metabotropic 1 |
| 1448148 | at | 1.29 | 0.000 | Grn | granulin |
| 1423423 | at | -1.13 | 0.109 | Grp58 | glucose regulated protein |
| 1417320 | at | -1.07 | 0.084 | Grpel1 | GrpE-like 1, mitochondrial |
| 1436256 | at | 1.03 | 0.353 | Grpel2 | GrpE-like 2, mitochondrial |
| 1452171 | at | 1.47 | 0.000 | Grwd1 | glutamate-rich WD repeat containing 1 |
| 1428767 | at | 1.12 | 0.471 | Gsdmdc1 | gasdermin domain containing 1 |
| 1415812 | at | -1.06 | 0.059 | Gsn | gelsolin |
| 1435000 | at | 1.04 | 0.551 | Gspt1 | G1 to S phase transition 1 |
| 1421817 | at | -1.04 | 0.567 | Gsr | glutathione reductase 1 |
| 1423436 | at | -1.23 | 0.073 | Gsta3 | glutathione S-transferase, alpha 3 |
| 1416368 | at | -1.05 | 0.276 | Gsta4 | glutathione S-transferase, alpha 4 |
| 1452823 | at | -1.88 | 0.000 | Gstk1 | glutathione S-transferase kappa 1 |
| 1448330 | at | -1.34 | 0.000 | Gstm1 | glutathione S-transferase, mu 1 |
| 1416411 | at | -1.37 | 0.000 | Gstm2 | glutathione S-transferase, mu 2 |
| 1416842 | at | -1.24 | 0.000 | Gstm5 | glutathione S-transferase, mu 5 |
| 1419072 | at | -1.48 | 0.000 | Gstm7 | glutathione S-transferase, mu 7 |
| 1416531 | at | 1.16 | 0.005 | Gsto1 | glutathione S-transferase omega 1 |
| 1449575 | a_at | -1.23 | 0.000 | Gstp1 | glutathione S-transferase, pi 1 |
| 1418186 | at | -1.64 | 0.000 | Gstt1 | glutathione S-transferase, theta 1 |
| 1427552 | a_at | -1.13 | 0.007 | Gstz1 | glutathione transferase zeta 1 (maleylacetoacetate isomerase) |
| 1452239 | at | 1.24 | 0.051 | Gt(ROSA)26So | gene trap ROSA 26, Philippe Soriano |

Supplementary table S1

| | | | | | |
|---|---|---|---|---|---|
| 1434290 | at | 1.08 | 0.322 | Gtdc1 | glycosyltransferase-like domain containing 1 |
| 1433511 | at | 1.00 | 0.961 | Gtf2a1 | general transcription factor II A, 1 |
| 1433528 | at | -1.06 | 0.194 | Gtf2a2 | general transcription factor II A, 2 |
| 1451135 | at | 1.00 | 0.998 | Gtf2b | general transcription factor IIB |
| 1428124 | at | 1.15 | 0.280 | Gtf2e1 | general transcription factor II E, polypeptide 1 (alpha subunit) |
| 1448884 | at | 1.27 | 0.004 | Gtf2e2 | general transcription factor II E, polypeptide 2 (beta subunit) |
| 1417698 | at | 1.08 | 0.497 | Gtf2f1 | general transcription factor IIF, polypeptide 1 |
| 1426626 | at | -1.00 | 0.989 | Gtf2f2 | general transcription factor IIF, polypeptide 2 |
| 1453169 | a_at | 1.15 | 0.001 | Gtf2h1 | general transcription factor II H, polypeptide 1 |
| 1450701 | a_at | 1.16 | 0.018 | Gtf2h2 | general transcription factor II H, polypeptide 2 |
| 1425628 | a_at | -1.15 | 0.008 | Gtf2i | general transcription factor II I |
| 1451899 | a_at | -1.35 | 0.017 | Gtf2ird1 | general transcription factor II I repeat domain-containing 1 |
| 1418343 | at | -1.11 | 0.137 | Gtf2ird2 | GTF2I repeat domain containing 2 |
| 1427027 | a_at | -1.03 | 0.599 | Gtf3a | general transcription factor III A |
| 1434066 | at | -1.07 | 0.508 | Gtf3c1 | general transcription factor III C 1 |
| 1428449 | at | -1.42 | 0.003 | Gtf3c2 | general transcription factor IIIC, polypeptide 2, beta |
| 1434320 | at | 1.00 | 0.941 | Gtf3c4 | general transcription factor IIIC, polypeptide 4 |
| 1452896 | at | -1.23 | 0.002 | Gtl3 | gene trap locus 3 |
| 1429047 | at | -1.13 | 0.090 | Gtl7 | gene trap locus 7 |
| 1419203 | at | 1.10 | 0.149 | Gtlf3a | gene trap locus F3a |
| 1417477 | at | 1.10 | 0.317 | Gtlf3b | gene trap locus F3b |
| 1450023 | at | -1.11 | 0.262 | Gtpbp1 | GTP binding protein 1 |
| 1450873 | at | 1.32 | 0.001 | Gtpbp4 | GTP binding protein 4 |
| 1434141 | at | -1.46 | 0.002 | Gucy1a3 | guanylate cyclase 1, soluble, alpha 3 |
| 1420872 | at | -1.07 | 0.325 | Gucy1b3 | guanylate cyclase 1, soluble, beta 3 |
| 1416395 | at | 1.01 | 0.946 | Guk1 | guanylate kinase 1 |
| 1448124 | at | 1.26 | 0.038 | Gusb | glucuronidase, beta |
| 1429184 | at | 1.07 | 0.675 | Gvin1 | GTPase, very large interferon inducible 1 |
| 1448429 | at | 1.26 | 0.000 | Gyg1 | glycogenin 1 |
| 1422704 | at | 1.23 | 0.009 | Gyk | glycerol kinase |
| 1423878 | at | -1.35 | 0.000 | Gypc | glycophorin C |
| 1416737 | at | -1.08 | 0.362 | Gys3 | glycogen synthase 3, brain |
| 1449501 | a_at | -2.17 | 0.001 | Gzmm | granzyme M (lymphocyte met-ase 1) |
| 1417287 | at | 1.07 | 0.428 | H13 | histocompatibility 13 |
| 1448194 | a_at | -2.02 | 0.000 | H19 | H19 fetal liver mRNA |
| 1423702 | at | 1.01 | 0.950 | H1f0 | H1 histone family, member 0 |
| 1451721 | a_at | 1.24 | 0.221 | H2-Ab1 | histocompatibility 2, class II antigen A, beta 1 |
| 1436596 | at | -1.16 | 0.035 | H2afv | H2A histone family, member V |
| 1424572 | a_at | -1.04 | 0.589 | H2afy | H2A histone family, member Y |
| 1421612 | a_at | 1.08 | 0.478 | H2afy2 | H2A histone family, member Y2 |
| 1417025 | at | -1.01 | 0.918 | H2-Eb1 | histocompatibility 2, class II antigen E beta |

Supplementary table S1

| | | | | | |
|---|---|---|---|---|---|
| 1415744 | at | -1.32 | 0.000 | H2-Ke2 | H2-K region expressed gene 2 |
| 1454987 | a_at | -1.27 | 0.000 | H2-Ke6 | H2-K region expressed gene 6 |
| 1426324 | at | 1.10 | 0.263 | H2-L | histocompatibility 2, D region |
| 1421358 | at | 1.10 | 0.224 | H2-M3 | histocompatibility 2, M region locus 3 |
| 1418536 | at | 1.18 | 0.351 | H2-Q7 | histocompatibility 2, Q region locus 7 |
| 1449556 | at | 1.15 | 0.076 | H2-T23 | histocompatibility 2, T region locus 23 |
| 1422160 | at | -1.09 | 0.199 | H2-T24 | histocompatibility 2, T region locus 24 |
| 1423263 | at | -1.18 | 0.005 | H3f3a | H3 histone, family 3A |
| 1455725 | a_at | 1.01 | 0.940 | H3f3b | H3 histone, family 3B |
| 1452095 | a_at | -1.14 | 0.227 | H47 | histocompatibility 47 |
| 1452145 | at | -1.47 | 0.016 | H6pd | hexose-6-phosphate dehydrogenase (glucose 1-dehydrogenase) |
| 1444908 | at | 1.42 | 0.050 | Habp4 | hyaluronic acid binding protein 4 |
| 1452278 | a_at | 1.05 | 0.600 | Hace1 | HECT domain and ankyrin repeat containing, E3 ubiquitin protein ligase 1 |
| 1448286 | at | -1.33 | 0.000 | Hadh2 | hydroxyacyl-Coenzyme A dehydrogenase type II |
| 1452173 | at | -1.45 | 0.000 | Hadha | hydroxyacyl-COA dehydrogenase/3-ketoacyl-COA thiolase/enoyl-COA hydratase, alpha subunit |
| 1426522 | at | -1.04 | 0.231 | Hadhb | hydroxyacyl-COA dehydrogenase/3-ketoacyl-COA thiolase/enoyl-COA hydratase, beta subunit |
| 1460184 | at | -1.08 | 0.039 | Hadhsc | L-3-hydroxyacyl-Coenzyme A dehydrogenase, short chain |
| 1424172 | at | 1.14 | 0.062 | Hagh | hydroxyacyl glutathione hydrolase |
| 1436041 | at | -1.21 | 0.082 | Hand2 | heart and neural crest derivatives expressed transcript 2 |
| 1417024 | at | 1.06 | 0.340 | Hars | histidyl-tRNA synthetase |
| 1451040 | at | -1.12 | 0.176 | Hars2 | histidyl tRNA synthetase 2 |
| 1449371 | at | -1.14 | 0.168 | Harsl | histidyl-tRNA synthetase-like |
| 1428061 | at | -1.00 | 0.995 | Hat1 | histone aminotransferase 1 |
| 1418350 | at | 2.97 | 0.000 | Hbegf | heparin-binding EGF-like growth factor |
| 1424154 | a_at | -1.06 | 0.069 | Hbld1 | HESB like domain containing 1 |
| 1423652 | at | -1.12 | 0.309 | Hbld2 | HESB like domain containing 2 |
| 1460367 | at | -1.19 | 0.010 | Hbp1 | high mobility group box transcription factor 1 |
| 1420890 | at | -1.09 | 0.151 | Hccs | holocytochrome c synthetase |
| 1433736 | at | -1.01 | 0.911 | Hcfc1 | host cell factor C1 |
| 1428405 | at | 1.06 | 0.128 | Hcfc1r1 | host cell factor C1 regulator 1 (XPO1-dependent) |
| 1451229 | at | -1.83 | 0.000 | Hdac11 | histone deacetylase 11 |
| 1448246 | at | 1.18 | 0.088 | Hdac1-ps | histone deacetylase 1 pseudogene |
| 1449080 | at | -1.12 | 0.100 | Hdac2 | histone deacetylase 2 |
| 1454693 | at | -1.28 | 0.013 | Hdac4 | histone deacetylase 4 |
| 1415743 | at | -1.38 | 0.000 | Hdac5 | histone deacetylase 5 |
| 1448928 | at | 1.27 | 0.007 | Hdac6 | histone deacetylase 6 |
| 1420813 | at | -1.22 | 0.026 | Hdac7a | histone deacetylase 7A |
| 1434572 | at | -1.29 | 0.012 | Hdac9 | histone deacetylase 9 |
| 1415888 | at | 1.11 | 0.114 | Hdgf | hepatoma-derived growth factor |
| 1450924 | at | 1.13 | 0.111 | Hdgfrp3 | hepatoma-derived growth factor, related protein 3 |
| 1425969 | a_at | 1.32 | 0.053 | Hdh | Huntington disease gene homolog |

Supplementary table S1

| | | | | | |
|---|---|---|---|---|---|
| 1428507 | at | 1.07 | 0.169 | Hdhd2 | haloacid dehalogenase-like hydrolase domain containing 2 |
| 1415988 | at | -1.19 | 0.071 | Hdlbp | high density lipoprotein (HDL) binding protein |
| 1418172 | at | 1.13 | 0.240 | Hebp1 | heme binding protein 1 |
| 1424141 | at | -1.03 | 0.563 | Hectd1 | HECT domain containing 1 |
| 1433944 | at | 1.59 | 0.001 | Hectd2 | HECT domain containing 2 |
| 1427469 | at | -1.27 | 0.001 | Helz | helicase with zinc finger domain |
| 1424703 | at | -1.04 | 0.451 | Hemk1 | HemK methyltransferase family member 1 |
| 1434060 | at | 1.05 | 0.443 | Herc1 | hect (homologous to the E6-AP (UBE3A) carboxyl terminus) domain and RCC1 (CHC1)-like domain (RLD) 1 |
| 1426491 | at | 1.14 | 0.004 | Herc2 | hect (homologous to the E6-AP (UBE3A) carboxyl terminus) domain and RCC1 (CHC1)-like domain (RLD) 2 |
| 1433590 | at | 1.16 | 0.277 | Herc3 | hect domain and RLD 3 |
| 1427032 | at | 1.09 | 0.012 | Herc4 | hect domain and RLD 4 |
| 1449024 | a_at | 1.01 | 0.852 | Hexa | hexosaminidase A |
| 1460180 | at | 1.18 | 0.017 | Hexb | hexosaminidase B |
| 1415999 | at | -1.14 | 0.166 | Hey1 | hairy/enhancer-of-split related with YRPW motif 1 |
| 1418106 | at | -1.17 | 0.233 | Hey2 | hairy/enhancer-of-split related with YRPW motif 2 |
| 1438886 | at | 1.27 | 0.009 | Heyl | hairy/enhancer-of-split related with YRPW motif-like |
| 1429223 | a_at | -1.17 | 0.006 | Hfe2 | hemochromatosis type 2 (juvenile) (human homolog) |
| 1416540 | at | 1.08 | 0.364 | Hgs | HGF-regulated tyrosine kinase substrate |
| 1423319 | at | 1.08 | 0.517 | Hhex | hematopoietically expressed homeobox |
| 1423195 | at | 1.02 | 0.636 | Hiat1 | hippocampus abundant gene transcript 1 |
| 1423780 | at | -1.21 | 0.012 | Hibadh | 3-hydroxyisobutyrate dehydrogenase |
| 1451511 | at | 1.09 | 0.042 | Hibch | 3-hydroxyisobutyryl-Coenzyme A hydrolase |
| 1457231 | at | 1.20 | 0.034 | Hif1a | hypoxia inducible factor 1, alpha subunit |
| 1428576 | at | 1.36 | 0.000 | Hif1an | hypoxia-inducible factor 1, alpha subunit inhibitor |
| 1425196 | a_at | -1.29 | 0.003 | Hint2 | histidine triad nucleotide binding protein 2 |
| 1455014 | at | -1.16 | 0.859 | Hint3 | histidine triad nucleotide binding protein 3 |
| 1424755 | at | 1.16 | 0.212 | Hip1 | huntingtin interacting protein 1 |
| 1425552 | at | 1.06 | 0.536 | Hip1r | huntingtin interacting protein 1 related |
| 1442671 | at | -1.11 | 0.495 | Hip2 | huntingtin interacting protein 2 |
| 1424540 | at | -1.14 | 0.022 | Hipk1 | homeodomain interacting protein kinase 1 |
| 1426819 | at | 1.05 | 0.432 | Hipk3 | homeodomain interacting protein kinase 3 |
| 1418228 | at | -1.00 | 0.986 | Hirip5 | histone cell cycle regulation defective interacting protein 5 |
| 1452418 | at | 1.17 | 0.283 | Hist1h1b | histone 1, H1b |
| 1416101 | a_at | 1.07 | 0.360 | Hist1h1c | histone 1, H1c |
| 1445565 | at | 1.16 | 0.189 | Hist1h1e | histone 1, H1e |
| 1418072 | at | -1.02 | 0.566 | Hist1h2bc | histone 1, H2bc |
| 1428014 | at | -1.35 | 0.000 | Hist1h4h | histone 1, H4h |
| 1455095 | at | -1.43 | 0.042 | Hist2h2bb | histone 2, H2bb |
| 1422155 | at | 1.03 | 0.700 | Hist2h3c2 | histone 2, H3c2 |
| 1455712 | at | -1.14 | 0.072 | Hist3h2a | histone 3, H2a |
| 1435865 | at | -1.27 | 0.058 | Hist3h2bb | histone 3, H2bb |

Supplementary table S1

| | | | | | |
|---|---|---|---|---|---|
| 1422742 | at | 1.31 | 0.009 | Hivep1 | human immunodeficiency virus type I enhancer binding protein 1 |
| 1422018 | at | -1.03 | 0.746 | Hivep2 | human immunodeficiency virus type I enhancer binding protein 2 |
| 1458802 | at | 1.44 | 0.004 | Hivep3 | human immunodeficiency virus type I enhancer binding protein 3 |
| 1437974 | a_at | 1.09 | 0.228 | Hk1 | hexokinase 1 |
| 1422612 | at | 1.00 | 0.902 | Hk2 | hexokinase 2 |
| 1434736 | at | -1.64 | 0.008 | Hlf | hepatic leukemia factor |
| 1426475 | at | -1.12 | 0.150 | Hmbs | hydroxymethylbilane synthase |
| 1416722 | at | -1.08 | 0.148 | Hmg20a | high mobility group 20A |
| 1452534 | a_at | -1.08 | 0.337 | Hmgb2 | high mobility group box 2 |
| 1426953 | at | 1.12 | 0.083 | Hmgb2l1 | high mobility group box 2-like 1 |
| 1416155 | at | 1.11 | 0.018 | Hmgb3 | high mobility group box 3 |
| 1424639 | a_at | -1.04 | 0.276 | Hmgcl | 3-hydroxy-3-methylglutaryl-Coenzyme A lyase |
| 1422495 | a_at | 1.02 | 0.733 | Hmgn1 | high mobility group nucleosomal binding domain 1 |
| 1433507 | a_at | -1.02 | 0.640 | Hmgn2 | high mobility group nucleosomal binding domain 2 |
| 1431777 | a_at | 1.44 | 0.002 | Hmgn3 | high mobility group nucleosomal binding domain 3 |
| 1416399 | a_at | -1.05 | 0.399 | Hmox2 | heme oxygenase (decycling) 2 |
| 1448180 | a_at | 1.39 | 0.002 | Hn1 | hematological and neurological expressed sequence 1 |
| 1417702 | a_at | -1.69 | 0.000 | Hnmt | histamine N-methyltransferase |
| 1428407 | at | -1.11 | 0.073 | Hnrpa0 | heterogeneous nuclear ribonucleoprotein A0 |
| 1452774 | at | 1.06 | 0.174 | Hnrpa3 | heterogeneous nuclear ribonucleoprotein A3 |
| 1448144 | at | 1.28 | 0.006 | Hnrpab | heterogeneous nuclear ribonucleoprotein A/B |
| 1418693 | at | 1.01 | 0.879 | Hnrpc | heterogeneous nuclear ribonucleoprotein C |
| 1457568 | at | -1.10 | 0.149 | Hnrpd | heterogeneous nuclear ribonucleoprotein D |
| 1415872 | at | -1.07 | 0.028 | Hnrph1 | heterogeneous nuclear ribonucleoprotein H1 |
| 1415963 | at | 1.04 | 0.551 | Hnrph2 | heterogeneous nuclear ribonucleoprotein H2 |
| 1423684 | at | 1.11 | 0.201 | Hnrpk | heterogeneous nuclear ribonucleoprotein K |
| 1427144 | at | -1.00 | 0.963 | Hnrpll | heterogeneous nuclear ribonucleoprotein L-like |
| 1426698 | a_at | 1.09 | 0.192 | Hnrpm | heterogeneous nuclear ribonucleoprotein M |
| 1452030 | a_at | 1.01 | 0.952 | Hnrpr | heterogeneous nuclear ribonucleoprotein R |
| 1423051 | at | -1.04 | 0.326 | Hnrpu | heterogeneous nuclear ribonucleoprotein U |
| 1451984 | at | -1.08 | 0.152 | Hnrpul1 | heterogeneous nuclear ribonucleoprotein U-like 1 |
| 1428662 | a_at | -1.72 | 0.000 | Hod | homeobox only domain |
| 1436387 | at | -1.24 | 0.031 | Homer1 | homer homolog 1 (Drosophila) |
| 1424367 | a_at | 1.27 | 0.011 | Homer2 | homer homolog 2 (Drosophila) |
| 1455563 | at | -1.01 | 0.894 | Homer3 | homer homolog 3 (Drosophila) |
| 1438018 | at | -1.18 | 0.018 | Hook1 | hook homolog 1 (Drosophila) |
| 1439196 | at | -1.06 | 0.634 | Hook3 | hook homolog 3 (Drosophila) |
| 1452412 | at | -1.55 | 0.061 | Hoxc8 | homeo box C8 |
| 1415751 | at | -1.07 | 0.055 | Hp1bp3 | heterochromatin protein 1, binding protein 3 |
| 1448812 | at | 1.26 | 0.078 | Hpcal1 | hippocalcin-like 1 |
| 1419906 | at | 1.13 | 0.155 | Hpgd | hydroxyprostaglandin dehydrogenase 15 (NAD) |

Supplementary table S1

| | | | | | |
|---|---|---|---|---|---|
| 1448736 | a_at | 1.03 | 0.574 | Hprt1 | hypoxanthine guanine phosphoribosyl transferase 1 |
| 1450647 | at | 1.03 | 0.329 | Hps3 | Hermansky-Pudlak syndrome 3 homolog (human) |
| 1448629 | at | -1.16 | 0.088 | Hps4 | Hermansky-Pudlak syndrome 4 homolog (human) |
| 1424132 | at | 1.03 | 0.642 | Hras1 | Harvey rat sarcoma virus oncogene 1 |
| 1428991 | at | 1.04 | 0.772 | Hrasls | HRAS-like suppressor |
| 1451611 | at | 1.15 | 0.011 | Hrasls3 | HRAS like suppressor 3 |
| 1434840 | at | 1.23 | 0.198 | Hrb | HIV-1 Rev binding protein |
| 1428084 | at | 1.08 | 0.030 | Hrb2 | HIV-1 Rev binding protein 2 |
| 1419109 | at | -1.44 | 0.000 | Hrc | histidine rich calcium binding protein |
| 1452787 | a_at | -1.02 | 0.851 | Hrmt1l2 | heterogeneous nuclear ribonucleoproteins methyltransferase-like 2 (S. cerevisiae) |
| 1426749 | at | 1.15 | 0.019 | Hrmt1l3 | heterogeneous nuclear ribonucleoprotein methyltransferase-like 3 (S. cerevisiae) |
| 1443986 | at | 1.06 | 0.524 | Hrpt2 | hyperparathyroidism 2 homolog (human) |
| 1426187 | a_at | 1.02 | 0.771 | Hs1bp1 | HS1 binding protein |
| 1433843 | at | -1.33 | 0.003 | Hs1bp3 | HS1 binding protein 3 |
| 1450729 | at | 1.07 | 0.409 | Hs2st1 | heparan sulfate 2-O-sulfotransferase 1 |
| 1417293 | at | 1.09 | 0.332 | Hs6st1 | heparan sulfate 6-O-sulfotransferase 1 |
| 1449038 | at | -1.54 | 0.000 | Hsd11b1 | hydroxysteroid 11-beta dehydrogenase 1 |
| 1450010 | at | 1.03 | 0.829 | Hsd17b12 | hydroxysteroid (17-beta) dehydrogenase 12 |
| 1417369 | at | -1.11 | 0.015 | Hsd17b4 | hydroxysteroid (17-beta) dehydrogenase 4 |
| 1416968 | a_at | -1.10 | 0.194 | Hsd3b7 | hydroxy-delta-5-steroid dehydrogenase, 3 beta- and steroid delta-isomerase 7 |
| 1454996 | at | -1.14 | 0.030 | Hsdl1 | hydroxysteroid dehydrogenase like 1 |
| 1426856 | at | -1.03 | 0.544 | Hsdl2 | hydroxysteroid dehydrogenase like 2 |
| 1424622 | at | -1.11 | 0.041 | Hsf1 | heat shock factor 1 |
| 1425993 | a_at | 1.07 | 0.399 | Hsp105 | heat shock protein 105 |
| 1460371 | at | -1.01 | 0.831 | Hspa12b | heat shock protein 12B |
| 1448586 | at | 1.02 | 0.736 | Hspa14 | heat shock 70kDa protein 14 |
| 1452318 | a_at | 2.64 | 0.038 | Hspa1b | heat shock protein 1B |
| 1417101 | at | 1.53 | 0.028 | Hspa2 | heat shock protein 2 |
| 1416146 | at | 1.21 | 0.371 | Hspa4 | heat shock protein 4 |
| 1416064 | a_at | -1.18 | 0.013 | Hspa5 | heat shock 70kD protein 5 (glucose-regulated protein) |
| 1420622 | a_at | 1.07 | 0.033 | Hspa8 | heat shock protein 8 |
| 1418504 | at | -1.01 | 0.738 | Hspa9a | heat shock protein 9A |
| 1422943 | a_at | 1.35 | 0.062 | Hspb1 | heat shock protein 1 |
| 1429888 | a_at | -1.03 | 0.352 | Hspb2 | heat shock protein 2 |
| 1449872 | at | 1.03 | 0.636 | Hspb3 | heat shock protein 3 |
| 1436332 | at | 1.13 | 0.000 | Hspb6 | heat shock protein, alpha-crystallin-related, B6 |
| 1421289 | at | 1.26 | 0.087 | Hspb7 | heat shock protein family, member 7 (cardiovascular) |
| 1417014 | at | 1.35 | 0.007 | Hspb8 | heat shock 27kDa protein 8 |
| 1419287 | at | -1.03 | 0.301 | Hspc171 | HSPC171 protein |
| 1426645 | at | 1.26 | 0.098 | Hspca | heat shock protein 1, alpha |
| 1416364 | at | -1.03 | 0.597 | Hspcb | heat shock protein 1, beta |

Supplementary table S1

| | | | | | |
|---|---|---|---|---|---|
| 1426351 | at | -1.05 | 0.179 | Hspd1 | heat shock protein 1 (chaperonin) |
| 1422579 | at | -1.10 | 0.146 | Hspe1 | heat shock protein 1 (chaperonin 10) |
| 1418669 | at | 1.23 | 0.072 | Hspg2 | perlecan (heparan sulfate proteoglycan 2) |
| 1433980 | at | -1.07 | 0.145 | Htatip | HIV-1 tat interactive protein, homolog (human) |
| 1451814 | a_at | 1.11 | 0.203 | Htatip2 | HIV-1 tat interactive protein 2, homolog (human) |
| 1454760 | at | 1.22 | 0.002 | Htatsf1 | HIV TAT specific factor 1 |
| 1448115 | at | 1.05 | 0.505 | Htf9c | HpaII tiny fragments locus 9c |
| 1419292 | at | -1.32 | 0.001 | Htra3 | serine protease HTRA3 |
| 1418308 | at | -1.11 | 0.048 | Hus1 | Hus1 homolog (S. pombe) |
| 1424949 | at | 1.01 | 0.879 | Huwe1 | HECT, UBA and WWE domain containing 1 |
| 1448679 | at | 1.23 | 0.052 | Hyal2 | hyaluronidase 2 |
| 1452154 | at | 1.12 | 0.006 | Iars | isoleucine-tRNA synthetase |
| 1439153 | at | -1.03 | 0.714 | Ibrdc2 | IBR domain containing 2 |
| 1435226 | at | 1.32 | 0.000 | Ibrdc3 | IBR domain containing 3 |
| 1454905 | at | -1.10 | 0.107 | Ibtk | inhibitor of Bruton agammaglobulinemia tyrosine kinase |
| 1417901 | a_at | -1.01 | 0.912 | Ica1 | islet cell autoantigen 1 |
| 1424067 | at | 1.58 | 0.005 | Icam1 | intercellular adhesion molecule |
| 1448862 | at | -1.10 | 0.292 | Icam2 | intercellular adhesion molecule 2 |
| 1426500 | at | 1.04 | 0.395 | Icmt | isoprenylcysteine carboxyl methyltransferase |
| 1419212 | at | 1.13 | 0.005 | Icosl | icos ligand |
| 1460308 | a_at | -1.20 | 0.001 | Ict1 | immature colon carcinoma transcript 1 |
| 1425895 | a_at | -1.19 | 0.127 | Id1 | inhibitor of DNA binding 1 |
| 1435176 | a_at | 1.16 | 0.161 | Id2 | inhibitor of DNA binding 2 |
| 1416630 | at | 1.09 | 0.210 | Id3 | inhibitor of DNA binding 3 |
| 1423121 | at | -1.26 | 0.244 | Ide | insulin degrading enzyme |
| 1458383 | at | -1.41 | 0.065 | Idh2 | isocitrate dehydrogenase 2 (NADP+), mitochondrial |
| 1422500 | at | -1.21 | 0.001 | Idh3a | isocitrate dehydrogenase 3 (NAD+) alpha |
| 1418885 | a_at | -1.05 | 0.482 | Idh3b | isocitrate dehydrogenase 3 (NAD+) beta |
| 1416789 | at | -1.10 | 0.000 | Idh3g | isocitrate dehydrogenase 3 (NAD+), gamma |
| 1451122 | at | 1.40 | 0.117 | Idi1 | isopentenyl-diphosphate delta isomerase |
| 1455481 | at | 1.01 | 0.816 | Ids | iduronate 2-sulfatase |
| 1439251 | at | -1.10 | 0.394 | Idua | iduronidase, alpha-L- |
| 1416442 | at | -1.05 | 0.753 | Ier2 | immediate early response 2 |
| 1419647 | a_at | 1.56 | 0.003 | Ier3 | immediate early response 3 |
| 1448702 | at | 1.10 | 0.223 | Ier3ip1 | immediate early response 3 interacting protein 1 |
| 1417612 | at | 2.48 | 0.000 | Ier5 | immediate early response 5 |
| 1418825 | at | 1.12 | 0.103 | Ifi1 | interferon inducible protein 1 |
| 1448775 | at | 1.07 | 0.254 | Ifi203 | interferon activated gene 203 |
| 1419858 | at | 1.25 | 0.056 | Ifi204 | interferon activated gene 204 |
| 1426906 | at | 1.01 | 0.852 | Ifi205 | interferon activated gene 205 |
| 1426278 | at | 1.40 | 0.006 | Ifi27 | interferon, alpha-inducible protein 27 |

Supplementary table S1

| | | | | | |
|---|---|---|---|---|---|
| 1422476 | at | 1.08 | 0.071 | Ifi30 | interferon gamma inducible protein 30 |
| 1424617 | at | 1.11 | 0.324 | Ifi35 | interferon-induced protein 35 |
| 1423555 | a_at | 1.48 | 0.001 | Ifi44 | interferon-induced protein 44 |
| 1417292 | at | 1.54 | 0.039 | Ifi47 | interferon gamma inducible protein 47 |
| 1426276 | at | 1.31 | 0.045 | Ifih1 | interferon induced with helicase C domain 1 |
| 1450783 | at | 1.48 | 0.019 | Ifit1 | interferon-induced protein with tetratricopeptide repeats 1 |
| 1418293 | at | 1.64 | 0.000 | Ifit2 | interferon-induced protein with tetratricopeptide repeats 2 |
| 1449025 | at | 1.63 | 0.031 | Ifit3 | interferon-induced protein with tetratricopeptide repeats 3 |
| 1424254 | at | 1.08 | 0.077 | Ifitm1 | interferon induced transmembrane protein 1 |
| 1417460 | at | 1.15 | 0.108 | Ifitm2 | interferon induced transmembrane protein 2 |
| 1423754 | at | 1.17 | 0.051 | Ifitm3 | interferon induced transmembrane protein 3 |
| 1449026 | at | 1.01 | 0.781 | Ifnar1 | interferon (alpha and beta) receptor 1 |
| 1451462 | a_at | -1.03 | 0.638 | Ifnar2 | interferon (alpha and beta) receptor 2 |
| 1443602 | at | 1.21 | 0.359 | Ifngr1 | interferon gamma receptor 1 |
| 1451016 | at | 1.00 | 0.994 | Ifrd2 | interferon-related developmental regulator 2 |
| 1448958 | at | -1.00 | 0.974 | Ifrg15 | interferon alpha responsive gene |
| 1437401 | at | 1.56 | 0.011 | Igf1 | insulin-like growth factor 1 |
| 1448152 | at | -1.57 | 0.000 | Igf2 | insulin-like growth factor 2 |
| 1423062 | at | 1.35 | 0.044 | Igfbp3 | insulin-like growth factor binding protein 3 |
| 1437405 | a_at | 1.08 | 0.267 | Igfbp4 | insulin-like growth factor binding protein 4 |
| 1422313 | a_at | 1.10 | 0.650 | Igfbp5 | insulin-like growth factor binding protein 5 |
| 1437804 | at | 1.97 | 0.002 | Igfbp7 | insulin-like growth factor binding protein 7 |
| 1425385 | a_at | -1.21 | 0.106 | Igh-1a | immunoglobulin heavy chain 1a (serum IgG2a) |
| 1452557 | a_at | -1.08 | 0.531 | Igk-C | immunoglobulin kappa chain, constant region |
| 1427108 | at | 1.01 | 0.938 | Igk-V4 | immunoglobulin kappa chain variable 4 (V4) |
| 1427855 | at | 1.05 | 0.707 | Igk-V8 | immunoglobulin kappa chain variable 8 (V8) |
| 1433652 | at | 1.85 | 0.000 | Igsf1 | immunoglobulin superfamily, member 1 |
| 1452864 | at | -1.33 | 0.004 | Igsf10 | immunoglobulin superfamily, member 10 |
| 1455048 | at | 1.01 | 0.871 | Igsf3 | immunoglobulin superfamily, member 3 |
| 1417376 | a_at | -1.08 | 0.655 | Igsf4a | immunoglobulin superfamily, member 4A |
| 1439048 | at | 1.08 | 0.593 | Igsf4d | immunoglobulin superfamily, member 4 |
| 1417141 | at | -1.13 | 0.486 | Igtp | interferon gamma induced GTPase |
| 1452694 | at | -1.13 | 0.057 | Ihpk1 | inositol hexaphosphate kinase 1 |
| 1425519 | a_at | 1.08 | 0.478 | Ii | Ia-associated invariant chain |
| 1419042 | at | -1.22 | 0.118 | Iigp1 | interferon inducible GTPase 1 |
| 1417793 | at | 1.00 | 0.999 | Iigp2 | interferon inducible GTPase 2 |
| 1423791 | at | -1.05 | 0.323 | Ik | IK cytokine |
| 1451254 | at | 1.12 | 0.005 | Ikbkap | inhibitor of kappa light polypeptide enhancer in B-cells, kinase complex-associated protein |
| 1426207 | at | 1.15 | 0.127 | Ikbkb | inhibitor of kappaB kinase beta |
| 1454690 | at | 1.05 | 0.459 | Ikbkg | inhibitor of kappaB kinase gamma |
| 1419455 | at | -1.21 | 0.001 | Il10rb | interleukin 10 receptor, beta |

Supplementary table S1

| | | | | | |
|---|---|---|---|---|---|
| 1427164 | at | 1.34 | 0.028 | Il13ra1 | interleukin 13 receptor, alpha 1 |
| 1418219 | at | -2.08 | 0.000 | Il15 | interleukin 15 |
| 1422397 | a_at | -1.54 | 0.001 | Il15ra | interleukin 15 receptor, alpha chain |
| 1450424 | a_at | 1.60 | 0.001 | Il18bp | interleukin 18 binding protein |
| 1448950 | at | 1.10 | 0.191 | Il1r1 | interleukin 1 receptor, type I |
| 1416295 | a_at | 1.35 | 0.057 | Il2rg | interleukin 2 receptor, gamma chain |
| 1452843 | at | 1.05 | 0.240 | Il6st | interleukin 6 signal transducer |
| 1417949 | at | -1.11 | 0.236 | Ilf2 | interleukin enhancer binding factor 2 |
| 1449942 | a_at | 1.08 | 0.369 | Ilk | integrin linked kinase |
| 1452506 | a_at | -1.08 | 0.201 | Ilkap | integrin-linked kinase-associated serine/threonine phosphatase 2C |
| 1454658 | at | -1.15 | 0.009 | Ilvbl | ilvB (bacterial acetolactate synthase)-like |
| 1419495 | at | -1.13 | 0.034 | Immp2l | inner mitochondrial membrane peptidase 2-like (S. cerevisiae) |
| 1429533 | at | -1.14 | 0.204 | Immt | inner membrane protein, mitochondrial |
| 1426213 | at | 1.29 | 0.000 | Imp4 | IMP4, U3 small nucleolar ribonucleoprotein, homolog (yeast) |
| 1423127 | at | 1.15 | 0.026 | Impa1 | inositol (myo)-1(or 4)-monophosphatase 1 |
| 1418665 | at | -2.09 | 0.001 | Impa2 | inositol (myo)-1(or 4)-monophosphatase 2 |
| 1418984 | at | -1.09 | 0.827 | Inadl | InaD-like (Drosophila) |
| 1436069 | at | -1.26 | 0.280 | Ing5 | inhibitor of growth family, member 5 |
| 1418697 | at | -3.29 | 0.001 | Inmt | indolethylamine N-methyltransferase |
| 1418045 | at | 1.12 | 0.228 | Inpp1 | inositol polyphosphate-1-phosphatase |
| 1436616 | at | -1.04 | 0.470 | Inpp4a | inositol polyphosphate-4-phosphatase, type I |
| 1433605 | at | 1.14 | 0.004 | Inpp5a | inositol polyphosphate-5-phosphatase A |
| 1451330 | a_at | 1.06 | 0.440 | Inpp5b | inositol polyphosphate-5-phosphatase B |
| 1418110 | a_at | 1.43 | 0.012 | Inpp5d | inositol polyphosphate-5-phosphatase D |
| 1423230 | at | -1.10 | 0.214 | Inpp5e | inositol polyphosphate-5-phosphatase E |
| 1433542 | at | 1.07 | 0.379 | Inpp5f | inositol polyphosphate-5-phosphatase F |
| 1460394 | a_at | -1.14 | 0.108 | Inppl1 | inositol polyphosphate phosphatase-like 1 |
| 1454671 | at | -1.02 | 0.794 | Insig1 | insulin induced gene 1 |
| 1417982 | at | 1.20 | 0.334 | Insig2 | insulin induced gene 2 |
| 1456200 | at | 1.12 | 0.049 | Ipmk | inositol polyphosphate multikinase |
| 1428096 | at | -1.03 | 0.427 | Ipo11 | importin 11 |
| 1433464 | at | -1.20 | 0.024 | Ipo13 | importin 13 |
| 1450814 | a_at | 1.36 | 0.182 | Ipo4 | importin 4 |
| 1454955 | at | 1.35 | 0.002 | Ipo7 | importin 7 |
| 1426760 | at | -1.01 | 0.884 | Ipo8 | importin 8 |
| 1424466 | at | 1.08 | 0.210 | Ipo9 | importin 9 |
| 1454756 | at | 1.12 | 0.053 | Iqcg | IQ motif containing G |
| 1434998 | at | 1.29 | 0.066 | Iqgap1 | IQ motif containing GTPase activating protein 1 |
| 1427145 | at | -1.20 | 0.378 | Iqsec1 | IQ motif and Sec7 domain 1 |
| 1448063 | at | -1.13 | 0.803 | Iqsec2 | IQ motif and Sec7 domain 2 |
| 1453312 | at | 1.09 | 0.056 | Iqwd1 | IQ motif and WD repeats 1 |

Supplementary table S1

| | | | | | |
|---|---|---|---|---|---|
| 1460649 | at | -1.01 | 0.820 | Irak1 | interleukin-1 receptor-associated kinase 1 |
| 1431771 | a_at | -1.05 | 0.448 | Irak1bp1 | interleukin-1 receptor-associated kinase 1 binding protein 1 |
| 1435040 | at | 1.44 | 0.009 | Irak3 | interleukin-1 receptor-associated kinase 3 |
| 1433527 | at | -1.03 | 0.433 | Ireb2 | iron responsive element binding protein 2 |
| 1448436 | a_at | -1.03 | 0.754 | Irf1 | interferon regulatory factor 1 |
| 1449678 | at | 1.03 | 0.566 | Irf5 | interferon regulatory factor 5 |
| 1435540 | at | 1.05 | 0.312 | Irgq1 | immunity-related GTPase family, Q1 |
| 1423104 | at | -1.05 | 0.508 | Irs1 | insulin receptor substrate 1 |
| 1418517 | at | -1.00 | 0.964 | Irx3 | Iroquois related homeobox 3 (Drosophila) |
| 1419539 | at | -1.11 | 0.223 | Irx4 | Iroquois related homeobox 4 (Drosophila) |
| 1421322 | a_at | 1.19 | 0.049 | Isgf3g | interferon dependent positive acting transcription factor 3 gamma |
| 1418450 | at | 1.58 | 0.004 | Islr | immunoglobulin superfamily containing leucine-rich repeat |
| 1425053 | at | -1.30 | 0.105 | Isoc1 | isochorismatase domain containing 1 |
| 1415769 | at | -1.04 | 0.349 | Itch | itchy |
| 1418393 | a_at | 1.03 | 0.599 | Itga7 | integrin alpha 7 |
| 1452784 | at | 1.21 | 0.000 | Itgav | integrin alpha V |
| 1422157 | a_at | 1.13 | 0.003 | Itgb1bp1 | integrin beta 1 binding protein 1 |
| 1423238 | at | -1.07 | 0.122 | Itgb1bp2 | integrin beta 1 binding protein 2 |
| 1453898 | at | 7.20 | 0.001 | Itgb1bp3 | integrin beta 1 binding protein 3 |
| 1456097 | a_at | 1.35 | 0.009 | Itgb3bp | integrin beta 3 binding protein (beta3-endonexin) |
| 1428442 | at | 1.24 | 0.001 | Itgb4bp | integrin beta 4 binding protein |
| 1417534 | at | 1.33 | 0.007 | Itgb5 | integrin beta 5 |
| 1425039 | at | 4.37 | 0.000 | Itgbl1 | integrin, beta-like 1 |
| 1457120 | at | -1.61 | 0.001 | Itk | IL2-inducible T-cell kinase |
| 1450841 | at | 1.13 | 0.060 | Itm1 | intergral membrane protein 1 |
| 1423608 | at | -1.05 | 0.330 | Itm2a | integral membrane protein 2A |
| 1415961 | at | 1.07 | 0.207 | Itm2c | integral membrane protein 2C |
| 1441058 | at | -1.06 | 0.375 | Itpkb | inositol 1,4,5-trisphosphate 3-kinase B |
| 1460203 | at | 1.05 | 0.269 | Itpr1 | inositol 1,4,5-triphosphate receptor 1 |
| 1435884 | at | 1.04 | 0.669 | Itsn1 | intersectin 1 (SH3 domain protein 1A) |
| 1418238 | at | -1.26 | 0.001 | Ivd | isovaleryl coenzyme A dehydrogenase |
| 1421903 | at | -1.15 | 0.440 | Ixl | intersex-like (Drosophila) |
| 1434070 | at | -1.11 | 0.055 | Jag1 | jagged 1 |
| 1426430 | at | -1.09 | 0.586 | Jag2 | jagged 2 |
| 1417064 | at | -1.09 | 0.007 | Jagn1 | jagunal homolog 1 (Drosophila) |
| 1421066 | at | 1.09 | 0.101 | Jak2 | Janus kinase 2 |
| 1436568 | at | -1.13 | 0.123 | Jam2 | junction adhesion molecule 2 |
| 1435348 | at | -1.05 | 0.344 | Jarid1c | jumonji, AT rich interactive domain 1C (Rbp2 like) |
| 1424903 | at | 1.03 | 0.788 | Jarid1d | jumonji, AT rich interactive domain 1D (Rbp2 like) |
| 1456661 | at | -1.28 | 0.043 | Jarid2 | jumonji, AT rich interactive domain 2 |
| 1426810 | at | 1.24 | 0.088 | Jmjd1a | jumonji domain containing 1A |

Supplementary table S1

| | | | | | |
|---|---|---|---|---|---|
| 1428320 | at | -1.07 | 0.460 | Jmjd1b | jumonji domain containing 1B |
| 1448049 | at | 1.01 | 0.929 | Jmjd1c | jumonji domain containing 1C |
| 1435340 | at | -1.27 | 0.005 | Jmjd2a | jumonji domain containing 2A |
| 1424044 | at | 1.07 | 0.414 | Jmjd2b | jumonji domain containing 2B |
| 1424458 | at | -1.08 | 0.299 | Jmjd2c | jumonji domain containing 2C |
| 1420640 | at | -1.17 | 0.094 | Jmy | junction-mediating and regulatory protein |
| 1438428 | at | 1.02 | 0.819 | Jph1 | junctophilin 1 |
| 1455404 | at | -1.01 | 0.869 | Jph2 | junctophilin 2 |
| 1424283 | at | 1.01 | 0.765 | Jtb | jumping translocation breakpoint |
| 1424151 | at | 1.09 | 0.334 | Jtv1 | JTV1 gene |
| 1421344 | a_at | -1.26 | 0.033 | Jub | ajuba |
| 1417409 | at | 1.29 | 0.024 | Jun | Jun oncogene |
| 1449117 | at | 1.10 | 0.029 | Jund1 | Jun proto-oncogene related gene d1 |
| 1450350 | a_at | -1.01 | 0.951 | Jundm2 | Jun dimerization protein 2 |
| 1416401 | at | -1.24 | 0.013 | Kai1 | kangai 1 (suppression of tumorigenicity 6, prostate) |
| 1429796 | at | 1.46 | 0.044 | Kalrn | kalirin, RhoGEF kinase |
| 1416068 | at | -1.17 | 0.010 | Kars | lysyl-tRNA synthetase |
| 1450949 | at | -1.07 | 0.086 | Katna1 | katanin p60 (ATPase-containing) subunit A1 |
| 1442348 | at | 1.05 | 0.264 | Katnal1 | katanin p60 subunit A-like 1 |
| 1451214 | at | -1.16 | 0.018 | Kbtbd2 | kelch repeat and BTB (POZ) domain containing 2 |
| 1439062 | at | -1.01 | 0.721 | Kbtbd3 | kelch repeat and BTB (POZ) domain containing 3 |
| 1417922 | at | -1.22 | 0.000 | Kbtbd4 | kelch repeat and BTB (POZ) domain containing 4 |
| 1431043 | at | 1.35 | 0.031 | Kbtbd5 | kelch repeat and BTB (POZ) domain containing 5 |
| 1418694 | at | -1.04 | 0.318 | Kcmf1 | potassium channel modulatory factor 1 |
| 1438613 | at | -1.19 | 0.103 | Kcna4 | potassium voltage-gated channel, shaker-related subfamily, member 4 |
| 1417680 | at | -1.02 | 0.824 | Kcna5 | potassium voltage-gated channel, shaker-related subfamily, member 5 |
| 1440838 | at | -1.38 | 0.005 | Kcnb1 | potassium voltage gated channel, Shab-related subfamily, member 1 |
| 1458167 | at | -1.80 | 0.026 | Kcnd2 | potassium voltage-gated channel, Shal-related family, member 2 |
| 1435720 | at | -1.29 | 0.001 | Kcnd3 | potassium voltage-gated channel, Shal-related family, member 3 |
| 1418156 | at | 1.61 | 0.003 | Kcne4 | potassium voltage-gated channel, Isk-related subfamily, gene 4 |
| 1449544 | a_at | -1.53 | 0.004 | Kcnh2 | potassium voltage-gated channel, subfamily H (eag-related), member 2 |
| 1425870 | a_at | -1.67 | 0.056 | Kcnip2 | Kv channel-interacting protein 2 |
| 1455417 | at | -1.19 | 0.001 | Kcnj11 | potassium inwardly rectifying channel, subfamily J, member 11 |
| 1444025 | at | -2.15 | 0.003 | Kcnj3 | potassium inwardly-rectifying channel, subfamily J, member 3 |
| 1441590 | at | -1.72 | 0.001 | Kcnj5 | potassium inwardly-rectifying channel, subfamily J, member 5 |
| 1418142 | at | -1.15 | 0.014 | Kcnj8 | potassium inwardly-rectifying channel, subfamily J, member 8 |
| 1425341 | at | -1.33 | 0.000 | Kcnk3 | potassium channel, subfamily K, member 3 |
| 1445676 | at | -1.64 | 0.051 | Kcnn2 | potassium intermediate/small conductance calcium-activated channel, subfamily N, member 2 |
| 1422293 | a_at | -1.03 | 0.647 | Kctd1 | potassium channel tetramerisation domain containing 1 |
| 1423694 | at | 1.20 | 0.017 | Kctd10 | potassium channel tetramerisation domain containing 10 |
| 1442368 | at | 1.29 | 0.401 | Kctd12b | potassium channel tetramerisation domain containing 12b |

Supplementary table S1

| | | | | | |
|---|---|---|---|---|---|
| 1435525 | at | 1.62 | 0.002 | Kctd17 | potassium channel tetramerisation domain containing 17 |
| 1452625 | at | -1.27 | 0.099 | Kctd2 | potassium channel tetramerisation domain containing 2 |
| 1455816 | a_at | 1.37 | 0.005 | Kctd3 | potassium channel tetramerisation domain containing 3 |
| 1428186 | at | 1.02 | 0.779 | Kctd6 | potassium channel tetramerisation domain containing 6 |
| 1418399 | at | 1.15 | 0.013 | Kctd9 | potassium channel tetramerisation domain containing 9 |
| 1460211 | a_at | 1.01 | 0.886 | Kdelr1 | KDEL (Lys-Asp-Glu-Leu) endoplasmic reticulum protein retention receptor 1 |
| 1417204 | at | -1.09 | 0.033 | Kdelr2 | KDEL (Lys-Asp-Glu-Leu) endoplasmic reticulum protein retention receptor 2 |
| 1418538 | at | 1.70 | 0.002 | Kdelr3 | KDEL (Lys-Asp-Glu-Leu) endoplasmic reticulum protein retention receptor 3 |
| 1449379 | at | -1.25 | 0.010 | Kdr | kinase insert domain protein receptor |
| 1450746 | at | -1.09 | 0.136 | Keap1 | kelch-like ECH-associated protein 1 |
| 1426481 | at | -1.01 | 0.714 | Kelchl | kelch-like (D. melanogaster) |
| 1418629 | a_at | -1.02 | 0.567 | Khdrbs1 | KH domain containing, RNA binding, signal transduction associated 1 |
| 1453317 | a_at | -1.26 | 0.012 | Khdrbs3 | KH domain containing, RNA binding, signal transduction associated 3 |
| 1429062 | at | 1.10 | 0.494 | Kif16b | kinesin family member 16B |
| 1450052 | at | -1.12 | 0.358 | Kif2a | kinesin family member 2A |
| 1449841 | at | 1.36 | 0.012 | Kif3a | kinesin family member 3A |
| 1450074 | at | 1.22 | 0.016 | Kif3b | kinesin family member 3B |
| 1418429 | at | 1.63 | 0.000 | Kif5b | kinesin family member 5B |
| 1442238 | a_at | -1.99 | 0.000 | Kif6 | kinesin family member 6 |
| 1417181 | a_at | 1.06 | 0.516 | Kifap3 | kinesin-associated protein 3 |
| 1416199 | at | -1.03 | 0.510 | Kifc3 | kinesin family member C3 |
| 1427317 | at | 1.01 | 0.864 | Kin | antigenic determinant of rec-A protein |
| 1418214 | at | -1.01 | 0.936 | Klc2 | kinesin light chain 2 |
| 1455521 | at | -1.13 | 0.113 | Klf12 | Kruppel-like factor 12 |
| 1437916 | at | -1.30 | 0.011 | Klf13 | Kruppel-like factor 13 |
| 1456551 | at | -1.81 | 0.006 | Klf15 | Kruppel-like factor 15 |
| 1448890 | at | -1.02 | 0.853 | Klf2 | Kruppel-like factor 2 (lung) |
| 1443932 | at | -1.27 | 0.059 | Klhdc1 | kelch domain containing 1 |
| 1448164 | at | -1.18 | 0.199 | Klhdc3 | kelch domain containing 3 |
| 1426988 | at | -1.30 | 0.012 | Klhdc5 | kelch domain containing 5 |
| 1435786 | at | 1.21 | 0.010 | Klhl12 | kelch-like 12 (Drosophila) |
| 1416242 | at | -1.13 | 0.082 | Klhl13 | kelch-like 13 (Drosophila) |
| 1435818 | at | -1.10 | 0.325 | Klhl15 | kelch-like 15 (Drosophila) |
| 1435257 | at | -1.07 | 0.170 | Klhl17 | kelch-like 17 (Drosophila) |
| 1455543 | at | 1.20 | 0.082 | Klhl18 | kelch-like 18 (Drosophila) |
| 1426978 | at | 1.10 | 0.257 | Klhl2 | kelch-like 2, Mayven (Drosophila) |
| 1454994 | at | 1.04 | 0.360 | Klhl20 | kelch-like 20 (Drosophila) |
| 1428081 | at | -1.23 | 0.022 | Klhl21 | kelch-like 21 (Drosophila) |
| 1439078 | at | 1.17 | 0.146 | Klhl4 | kelch-like 4 (Drosophila) |
| 1437886 | at | 1.68 | 0.001 | Klhl6 | kelch-like 6 (Drosophila) |
| 1428091 | at | 1.06 | 0.288 | Klhl7 | kelch-like 7 (Drosophila) |

Supplementary table S1

| | | | | | |
|---|---|---|---|---|---|
| 1454637 | at | -1.09 | 0.244 | Klhl8 | kelch-like 8 (Drosophila) |
| 1433442 | at | -1.05 | 0.596 | Klhl9 | kelch-like 9 (Drosophila) |
| 1428881 | at | 1.00 | 0.975 | Kns2 | kinesin 2 |
| 1415860 | at | 1.04 | 0.410 | Kpna2 | karyopherin (importin) alpha 2 |
| 1421828 | at | 1.06 | 0.715 | Kpna3 | karyopherin (importin) alpha 3 |
| 1448526 | at | 1.04 | 0.447 | Kpnb1 | karyopherin (importin) beta 1 |
| 1423438 | at | -1.46 | 0.003 | Kptn | kaptin |
| 1451979 | at | 1.06 | 0.358 | Kras | v-Ki-ras2 Kirsten rat sarcoma viral oncogene homolog |
| 1435472 | at | 1.29 | 0.000 | Kremen1 | kringle containing transmembrane protein 1 |
| 1452166 | a_at | -1.23 | 0.044 | Krt1-10 | keratin complex 1, acidic, gene 10 |
| 1417059 | at | -1.04 | 0.338 | Krtcap2 | keratinocyte associated protein 2 |
| 1455434 | a_at | -1.02 | 0.497 | Ktn1 | kinectin 1 |
| 1440435 | at | -2.39 | 0.001 | Ky | kyphoscoliosis |
| 1433777 | at | -1.01 | 0.749 | L3mbtl2 | l(3)mbt-like 2 (Drosophila) |
| 1434246 | at | -1.08 | 0.342 | L3mbtl3 | l(3)mbt-like 3 (Drosophila) |
| 1427073 | at | -1.10 | 0.242 | Lace1 | lactation elevated 1 |
| 1449014 | at | 1.03 | 0.621 | Lactb | lactamase, beta |
| 1460151 | at | -1.01 | 0.936 | Lactb2 | lactamase, beta 2 |
| 1440424 | at | -2.28 | 0.000 | Lad1 | ladinin |
| 1450031 | at | 1.32 | 0.000 | Laf4l | lymphoid nuclear protein related to AF4-like |
| 1426285 | at | 1.16 | 0.073 | Lama2 | laminin, alpha 2 |
| 1424807 | at | 1.07 | 0.372 | Lama4 | laminin, alpha 4 |
| 1427009 | at | -1.50 | 0.000 | Lama5 | laminin, alpha 5 |
| 1416513 | at | -1.19 | 0.010 | Lamb2 | laminin, beta 2 |
| 1415880 | a_at | 1.09 | 0.000 | Lamp1 | lysosomal membrane glycoprotein 1 |
| 1428094 | at | 1.05 | 0.232 | Lamp2 | lysosomal membrane glycoprotein 2 |
| 1448245 | at | -1.01 | 0.845 | Lamr1 | laminin receptor 1 (ribosomal protein SA) |
| 1427011 | a_at | 1.04 | 0.472 | Lancl1 | LanC (bacterial lantibiotic synthetase component C)-like 1 |
| 1458828 | at | -1.34 | 0.213 | Lancl2 | LanC (bacterial lantibiotic synthetase component C)-like 2 |
| 1450860 | at | 1.10 | 0.120 | Lap3 | leucine aminopeptidase 3 |
| 1423368 | at | -1.09 | 0.105 | Laptm4a | lysosomal-associated protein transmembrane 4A |
| 1416148 | at | -1.13 | 0.039 | Laptm4b | lysosomal-associated protein transmembrane 4B |
| 1448403 | at | 1.15 | 0.004 | Lars | leucyl-tRNA synthetase |
| 1435682 | at | 1.01 | 0.909 | Lars2 | leucyl-tRNA synthetase, mitochondrial |
| 1460173 | at | 1.08 | 0.268 | Lasp1 | LIM and SH3 protein 1 |
| 1423955 | a_at | 1.18 | 0.000 | Lass2 | longevity assurance homolog 2 (S. cerevisiae) |
| 1417782 | at | -1.10 | 0.300 | Lass4 | longevity assurance homolog 4 (S. cerevisiae) |
| 1451209 | at | 1.15 | 0.047 | Lass5 | longevity assurance homolog 5 (S. cerevisiae) |
| 1434418 | at | 1.18 | 0.057 | Lass6 | longevity assurance homolog 6 (S. cerevisiae) |
| 1435739 | at | 1.03 | 0.693 | Lats1 | large tumor supressor |
| 1437101 | at | 1.01 | 0.959 | Lats2 | large tumor suppressor 2 |

Supplementary table S1

| | | | | | |
|---|---|---|---|---|---|
| 1415829 | at | -1.01 | 0.859 | Lbr | lamin B receptor |
| 1417044 | at | -1.04 | 0.646 | Lcmt1 | leucine carboxyl methyltransferase 1 |
| 1433518 | at | -1.25 | 0.000 | Lcmt2 | leucine carboxyl methyltransferase 2 |
| 1415983 | at | 1.43 | 0.003 | Lcp1 | lymphocyte cytosolic protein 1 |
| 1452024 | a_at | -1.27 | 0.102 | Ldb1 | LIM domain binding 1 |
| 1451999 | at | 1.06 | 0.458 | Ldb3 | LIM domain binding 3 |
| 1419737 | a_at | 1.08 | 0.052 | Ldh1 | lactate dehydrogenase 1, A chain |
| 1434499 | a_at | -1.07 | 0.026 | Ldh2 | lactate dehydrogenase 2, B chain |
| 1428613 | at | -1.37 | 0.004 | Ldhd | lactate dehydrogenase D |
| 1421821 | at | 1.38 | 0.034 | Ldlr | low density lipoprotein receptor |
| 1424326 | at | 1.00 | 0.980 | Lemd2 | LEM domain containing 2 |
| 1425280 | at | -1.09 | 0.364 | Leng1 | leukocyte receptor cluster (LRC) member 1 |
| 1451066 | at | 1.03 | 0.632 | Leng4 | leukocyte receptor cluster (LRC) member 4 |
| 1431593 | a_at | 1.48 | 0.002 | Leng5 | leukocyte receptor cluster (LRC) member 5 |
| 1435152 | at | -1.23 | 0.005 | Leng8 | leukocyte receptor cluster (LRC) member 8 |
| 1452752 | at | 1.34 | 0.006 | Lepre1 | leprecan 1 |
| 1451350 | a_at | 1.18 | 0.162 | Leprot | leptin receptor overlapping transcript |
| 1435300 | at | -1.02 | 0.828 | Letm1 | leucine zipper-EF-hand containing transmembrane protein 1 |
| 1418860 | a_at | 1.05 | 0.209 | Letmd1 | LETM1 domain containing 1 |
| 1448380 | at | 1.11 | 0.196 | Lgals3bp | lectin, galactoside-binding, soluble, 3 binding protein |
| 1451336 | at | -2.65 | 0.000 | Lgals4 | lectin, galactose binding, soluble 4 |
| 1422662 | at | 1.05 | 0.202 | Lgals8 | lectin, galactose binding, soluble 8 |
| 1421217 | a_at | 1.12 | 0.194 | Lgals9 | lectin, galactose binding, soluble 9 |
| 1448883 | at | -1.01 | 0.914 | Lgmn | legumain |
| 1451211 | a_at | 1.01 | 0.882 | Lgtn | ligatin |
| 1418700 | at | -1.04 | 0.194 | Lias | lipoic acid synthetase |
| 1454984 | at | -1.39 | 0.008 | Lifr | leukemia inhibitory factor receptor |
| 1437198 | at | -1.07 | 0.225 | Lig3 | ligase III, DNA, ATP-dependent |
| 1439487 | at | -1.05 | 0.362 | Lig4 | ligase IV, DNA, ATP-dependent |
| 1422730 | at | 1.05 | 0.198 | Limd1 | LIM domains containing 1 |
| 1448500 | a_at | -1.16 | 0.207 | Lime1 | Lck interacting transmembrane adaptor 1 |
| 1417627 | a_at | 1.17 | 0.135 | Limk1 | LIM-domain containing, protein kinase |
| 1452060 | a_at | 1.21 | 0.025 | Limk2 | LIM motif-containing protein kinase 2 |
| 1443947 | at | -1.02 | 0.894 | Lims1 | LIM and senescent cell antigen-like domains 1 |
| 1424408 | at | -1.17 | 0.073 | Lims2 | LIM and senescent cell antigen like domains 2 |
| 1450937 | at | -1.01 | 0.939 | Lin7c | lin 7 homolog c (C. elegans) |
| 1423140 | at | -1.00 | 0.992 | Lip1 | lysosomal acid lipase 1 |
| 1416303 | at | 1.29 | 0.017 | Litaf | LPS-induced TN factor |
| 1416621 | at | 1.04 | 0.657 | Llglh | lethal giant larvae homolog |
| 1423075 | at | -1.08 | 0.181 | Lman2 | lectin, mannose-binding 2 |
| 1435475 | at | 1.02 | 0.643 | Lman2l | lectin, mannose-binding 2-like |

Supplementary table S1

| | | | | | |
|---|---|---|---|---|---|
| 1426272 | at | -1.15 | 0.009 | Lmbr1 | limb region 1 |
| 1421654 | a_at | 1.26 | 0.125 | Lmna | lamin A |
| 1448531 | at | 1.00 | 0.967 | Lmnb2 | lamin B2 |
| 1454086 | a_at | 1.41 | 0.007 | Lmo2 | LIM domain only 2 |
| 1420981 | a_at | -1.09 | 0.441 | Lmo4 | LIM domain only 4 |
| 1452345 | at | 1.44 | 0.001 | Lmod2 | leiomodin 2 (cardiac) |
| 1455018 | at | -1.38 | 0.006 | Lmtk2 | lemur tyrosine kinase 2 |
| 1433455 | at | 1.01 | 0.843 | Lnk | linker of T-cell receptor pathways |
| 1453035 | at | 1.12 | 0.161 | Lnp | limb and neural patterns |
| 1438432 | at | -1.09 | 0.512 | Lnpep | leucyl/cystinyl aminopeptidase |
| 1450251 | a_at | 1.26 | 0.000 | Lnx1 | ligand of numb-protein X 1 |
| 1460249 | at | -1.02 | 0.752 | Lnx2 | ligand of numb-protein X 2 |
| 1441075 | at | 1.26 | 0.037 | LOC329416 | nitric oxide synthase trafficker |
| 1426221 | at | 1.49 | 0.000 | Loh11cr2a | loss of heterozygosity, 11, chromosomal region 2, gene A homolog (human) |
| 1451918 | a_at | -1.13 | 0.147 | Loh12cr1 | loss of heterozygosity, 12, chromosomal region 1 homolog (human) |
| 1416121 | at | 3.82 | 0.000 | Lox | lysyl oxidase |
| 1451978 | at | 1.96 | 0.000 | Loxl1 | lysyl oxidase-like 1 |
| 1431004 | at | 2.26 | 0.001 | Loxl2 | lysyl oxidase-like 2 |
| 1435326 | at | 1.08 | 0.174 | Lpgat1 | lysophosphatidylglycerol acyltransferase 1 |
| 1428510 | at | -1.11 | 0.443 | Lphn1 | latrophilin 1 |
| 1458742 | at | 1.11 | 0.545 | Lphn3 | latrophilin 3 |
| 1418288 | at | -1.34 | 0.000 | Lpin1 | lipin 1 |
| 1452836 | at | 1.23 | 0.000 | Lpin2 | lipin 2 |
| 1415904 | at | -1.05 | 0.524 | Lpl | lipoprotein lipase |
| 1454899 | at | 1.19 | 0.025 | Lpp | LIM domain containing preferred translocation partner in lipoma |
| 1449099 | at | 1.01 | 0.909 | Lrba | LPS-responsive beige-like anchor |
| 1417290 | at | -1.53 | 0.016 | Lrg1 | leucine-rich alpha-2-glycoprotein 1 |
| 1449893 | a_at | -1.02 | 0.838 | Lrig1 | leucine-rich repeats and immunoglobulin-like domains 1 |
| 1448655 | at | 1.69 | 0.000 | Lrp1 | low density lipoprotein receptor-related protein 1 |
| 1416836 | at | 1.08 | 0.338 | Lrp10 | low-density lipoprotein receptor-related protein 10 |
| 1433536 | at | 2.17 | 0.000 | Lrp11 | low density lipoprotein receptor-related protein 11 |
| 1447478 | at | -1.03 | 0.901 | Lrp1b | low density lipoprotein-related protein 1B (deleted in tumors) |
| 1426288 | at | -1.56 | 0.008 | Lrp4 | low density lipoprotein receptor-related protein 4 |
| 1424353 | at | -1.23 | 0.001 | Lrpprc | leucine-rich PPR-motif containing |
| 1452411 | at | 1.12 | 0.123 | Lrrc1 | leucine rich repeat containing 1 |
| 1427388 | at | 1.22 | 0.000 | Lrrc2 | leucine rich repeat containing 2 |
| 1438422 | at | -1.02 | 0.674 | Lrrc20 | leucine rich repeat containing 20 |
| 1452865 | at | -1.08 | 0.125 | Lrrc27 | leucine rich repeat containing 27 |
| 1433858 | at | -1.05 | 0.633 | Lrrc28 | leucine rich repeat containing 28 |
| 1434330 | at | 1.18 | 0.374 | Lrrc35 | leucine rich repeat containing 35 |
| 1451245 | at | -1.41 | 0.011 | Lrrc3b | leucine rich repeat containing 3B |

Supplementary table S1

| | | | | | |
|---|---|---|---|---|---|
| 1447491 | at | 1.02 | 0.807 | Lrrc5 | leucine rich repeat containing 5 |
| 1434694 | at | -1.01 | 0.820 | Lrrc8 | leucine rich repeat containing 8 |
| 1433842 | at | 1.41 | 0.001 | Lrrfip1 | leucine rich repeat (in FLII) interacting protein 1 |
| 1436378 | at | -1.04 | 0.549 | Lrrfip2 | leucine rich repeat (in FLII) interacting protein 2 |
| 1456121 | at | -1.20 | 0.147 | Lrriq2 | leucine-rich repeats and IQ motif containing 2 |
| 1451985 | at | 1.43 | 0.011 | Lrrk1 | leucine-rich repeat kinase 1 |
| 1455489 | at | -1.33 | 0.001 | Lrrtm2 | leucine rich repeat transmembrane neuronal 2 |
| 1429176 | at | -1.01 | 0.916 | Lrsam1 | leucine rich repeat and sterile alpha motif containing 1 |
| 1460092 | at | -1.03 | 0.791 | Lsamp | limbic system-associated membrane protein |
| 1423873 | at | 1.06 | 0.585 | Lsm1 | LSM1 homolog, U6 small nuclear RNA associated (S. cerevisiae) |
| 1417515 | at | -1.04 | 0.484 | Lsm10 | U7 snRNP-specific Sm-like protein LSM10 |
| 1451884 | a_at | -1.16 | 0.116 | Lsm2 | LSM2 homolog, U6 small nuclear RNA associated (S. cerevisiae) |
| 1448536 | at | 1.02 | 0.801 | Lsm3 | LSM3 homolog, U6 small nuclear RNA associated (S. cerevisiae) |
| 1448622 | at | -1.10 | 0.055 | Lsm4 | LSM4 homolog, U6 small nuclear RNA associated (S. cerevisiae) |
| 1418656 | at | -1.04 | 0.528 | Lsm5 | LSM5 homolog, U6 small nuclear RNA associated (S. cerevisiae) |
| 1431054 | at | 1.01 | 0.905 | Lsm6 | LSM6 homolog, U6 small nuclear RNA associated (S. cerevisiae) |
| 1448703 | at | -1.11 | 0.126 | Lsm8 | LSM8 homolog, U6 small nuclear RNA associated (S. cerevisiae) |
| 1417756 | a_at | -1.02 | 0.877 | Lsp1 | lymphocyte specific 1 |
| 1454964 | at | -1.23 | 0.001 | Lt1 | lurcher transcript 1 |
| 1434790 | a_at | 1.01 | 0.857 | Lta4h | leukotriene A4 hydrolase |
| 1417777 | at | -1.03 | 0.765 | Ltb4dh | leukotriene B4 12-hydroxydehydrogenase |
| 1448870 | at | -1.36 | 0.003 | Ltbp1 | latent transforming growth factor beta binding protein 1 |
| 1418049 | at | 1.70 | 0.002 | Ltbp3 | latent transforming growth factor beta binding protein 3 |
| 1436665 | a_at | 1.32 | 0.005 | Ltbp4 | latent transforming growth factor beta binding protein 4 |
| 1416435 | at | 1.10 | 0.047 | Ltbr | lymphotoxin B receptor |
| 1424791 | a_at | -1.66 | 0.000 | Lu | Lutheran blood group (Auberger b antigen included) |
| 1455261 | at | 1.08 | 0.260 | Luc7l | Luc7 homolog (S. cerevisiae)-like |
| 1423607 | at | 2.05 | 0.001 | Lum | lumican |
| 1448352 | at | 1.04 | 0.758 | Luzp1 | leucine zipper protein 1 |
| 1416503 | at | 1.59 | 0.003 | Lxn | latexin |
| 1417185 | at | 1.12 | 0.004 | Ly6a | lymphocyte antigen 6 complex, locus A |
| 1421571 | a_at | 1.16 | 0.069 | Ly6c | lymphocyte antigen 6 complex, locus C |
| 1422903 | at | 1.42 | 0.001 | Ly86 | lymphocyte antigen 86 |
| 1449874 | at | -1.04 | 0.542 | Ly96 | lymphocyte antigen 96 |
| 1417511 | at | -1.02 | 0.823 | Lyar | Ly1 antibody reactive clone |
| 1451318 | a_at | -1.01 | 0.837 | Lyn | Yamaguchi sarcoma viral (v-yes-1) oncogene homolog |
| 1417283 | at | -1.12 | 0.211 | Lynx1 | Ly6/neurotoxin 1 |
| 1448244 | at | 1.01 | 0.877 | Lypla1 | lysophospholipase 1 |
| 1423547 | at | 1.25 | 0.002 | Lyzs | lysozyme |
| 1451421 | a_at | 1.09 | 0.499 | Lzf | leucine zipper domain protein |
| 1448095 | at | -1.33 | 0.031 | Lzic | leucine zipper and CTNNBIP1 domain containing |

Supplementary table S1

| | | | | | |
|---|---|---|---|---|---|
| 1417170 | at | 1.16 | 0.010 | Lztfl1 | leucine zipper transcription factor-like 1 |
| 1426409 | at | -1.20 | 0.042 | Lzts2 | leucine zipper, putative tumor suppressor 2 |
| 1416424 | at | -1.29 | 0.006 | M6prbp1 | mannose-6-phosphate receptor binding protein 1 |
| 1439582 | at | -1.02 | 0.707 | Macf1 | microtubule-actin crosslinking factor 1 |
| 1434830 | at | -1.03 | 0.676 | Mad | Max dimerization protein |
| 1423031 | at | 1.11 | 0.138 | Maea | macrophage erythroblast attacher |
| 1451716 | at | 1.11 | 0.538 | Mafb | v-maf musculoaponeurotic fibrosarcoma oncogene family, protein B (avian) |
| 1448916 | at | 1.10 | 0.257 | Mafg | v-maf musculoaponeurotic fibrosarcoma oncogene family, protein G (avian) |
| 1418616 | at | 1.70 | 0.000 | Mafk | v-maf musculoaponeurotic fibrosarcoma oncogene family, protein K (avian) |
| 1450062 | a_at | -1.04 | 0.628 | Maged1 | melanoma antigen, family D, 1 |
| 1426306 | a_at | 1.02 | 0.847 | Maged2 | melanoma antigen, family D, 2 |
| 1422498 | at | 1.14 | 0.088 | Mageh1 | melanoma antigen, family H, 1 |
| 1416212 | at | -1.01 | 0.801 | Magoh | mago-nashi homolog, proliferation-associated (Drosophila) |
| 1429720 | at | 1.05 | 0.476 | Mak10 | MAK10 homolog, amino-acid N-acetyltransferase subunit, (S. cerevisiae) |
| 1428410 | at | 1.22 | 0.007 | Mak3 | Mak3 homolog (S. cerevisiae) |
| 1417275 | at | -1.17 | 0.001 | Mal | myelin and lymphocyte protein, T-cell differentiation protein |
| 1456429 | at | -1.23 | 0.017 | Malt1 | mucosa associated lymphoid tissue lymphoma translocation gene 1 |
| 1436193 | at | -1.02 | 0.769 | Man1c1 | mannosidase, alpha, class 1C, member 1 |
| 1448647 | at | 1.04 | 0.468 | Man2a1 | mannosidase 2, alpha 1 |
| 1416340 | a_at | -1.07 | 0.312 | Man2b1 | mannosidase 2, alpha B1 |
| 1416584 | at | 1.14 | 0.334 | Man2b2 | mannosidase 2, alpha B2 |
| 1423687 | a_at | 1.09 | 0.108 | Man2c1 | mannosidase, alpha, class 2C, member 1 |
| 1450626 | at | -1.08 | 0.084 | Manba | mannosidase, beta A, lysosomal |
| 1455403 | at | -1.08 | 0.199 | Manea | mannosidase, endo-alpha |
| 1423284 | at | -1.14 | 0.206 | Mansc1 | MANSC domain containing 1 |
| 1434354 | at | -1.58 | 0.001 | Maob | monoamine oxidase B |
| 1451290 | at | -1.15 | 0.002 | Map1lc3a | microtubule-associated protein 1 light chain 3 alpha |
| 1416351 | at | 1.28 | 0.002 | Map2k1 | mitogen activated protein kinase kinase 1 |
| 1416272 | at | 1.24 | 0.001 | Map2k1ip1 | mitogen-activated protein kinase kinase 1 interacting protein 1 |
| 1415974 | at | 1.01 | 0.915 | Map2k2 | mitogen activated protein kinase kinase 2 |
| 1451714 | a_at | 1.09 | 0.071 | Map2k3 | mitogen activated protein kinase kinase 3 |
| 1417854 | at | -1.20 | 0.039 | Map2k5 | mitogen activated protein kinase kinase 5 |
| 1426850 | a_at | -1.54 | 0.007 | Map2k6 | mitogen activated protein kinase kinase 6 |
| 1424850 | at | 1.06 | 0.409 | Map3k1 | mitogen activated protein kinase kinase kinase 1 |
| 1450669 | at | -1.04 | 0.670 | Map3k11 | mitogen activated protein kinase kinase kinase 11 |
| 1455597 | at | -1.13 | 0.483 | Map3k2 | mitogen activated protein kinase kinase kinase 2 |
| 1436522 | at | 1.01 | 0.920 | Map3k3 | mitogen activated protein kinase kinase kinase 3 |
| 1439830 | at | -1.12 | 0.017 | Map3k5 | mitogen activated protein kinase kinase kinase 5 |
| 1434833 | at | -1.18 | 0.113 | Map4k2 | mitogen activated protein kinase kinase kinase kinase 2 |
| 1426759 | at | -1.19 | 0.002 | Map4k3 | mitogen-activated protein kinase kinase kinase kinase 3 |
| 1440609 | at | 1.15 | 0.945 | Map4k4 | mitogen-activated protein kinase kinase kinase kinase 4 |

Supplementary table S1

| | | | | | |
|---|---|---|---|---|---|
| 1427084 | a_at | -1.07 | 0.491 | Map4k5 | mitogen-activated protein kinase kinase kinase kinase 5 |
| 1422797 | at | -1.18 | 0.042 | Mapbpip | mitogen activated protein binding protein interacting protein |
| 1458002 | at | -1.19 | 0.201 | Mapk10 | mitogen activated protein kinase 10 |
| 1449283 | a_at | 1.18 | 0.030 | Mapk12 | mitogen-activated protein kinase 12 |
| 1416703 | at | -1.33 | 0.000 | Mapk14 | mitogen activated protein kinase 14 |
| 1427060 | at | -1.04 | 0.427 | Mapk3 | mitogen activated protein kinase 3 |
| 1419169 | at | 1.08 | 0.192 | Mapk6 | mitogen-activated protein kinase 6 |
| 1440856 | at | -1.09 | 0.073 | Mapk8 | mitogen activated protein kinase 8 |
| 1425975 | a_at | -1.12 | 0.313 | Mapk8ip3 | mitogen-activated protein kinase 8 interacting protein 3 |
| 1426648 | at | 1.08 | 0.116 | Mapkapk2 | MAP kinase-activated protein kinase 2 |
| 1437494 | at | -1.74 | 0.002 | Mapkapk3 | mitogen-activated protein kinase-activated protein kinase 3 |
| 1417016 | at | -1.26 | 0.001 | Mapkapk5 | MAP kinase-activated protein kinase 5 |
| 1423403 | at | 1.00 | 0.999 | Mapkbp1 | mitogen activated protein kinase binding protein 1 |
| 1422764 | at | 1.29 | 0.020 | Mapre1 | microtubule-associated protein, RP/EB family, member 1 |
| 1451990 | at | 1.16 | 0.360 | Mapre2 | microtubule-associated protein, RP/EB family, member 2 |
| 1427079 | at | 1.04 | 0.648 | Mapre3 | microtubule-associated protein, RP/EB family, member 3 |
| 1424888 | at | -1.25 | 0.160 | marf2 | membrane-associated ring finger (C3HC4) 2 |
| 1436614 | at | 1.63 | 0.001 | marf3 | membrane-associated ring finger (C3HC4) 3 |
| 1433834 | at | -1.02 | 0.650 | marf6 | membrane-associated ring finger (C3HC4) 6 |
| 1440966 | at | -1.14 | 0.219 | marf7 | membrane-associated ring finger (C3HC4) 7 |
| 1428385 | at | 1.04 | 0.275 | marf8 | membrane-associated ring finger (C3HC4) 8 |
| 1435889 | at | -1.22 | 0.107 | Mark2 | MAP/microtubule affinity-regulating kinase 2 |
| 1418316 | a_at | 1.01 | 0.799 | Mark3 | MAP/microtubule affinity-regulating kinase 3 |
| 1455951 | at | -1.08 | 0.271 | Mars | methionine-tRNA synthetase |
| 1436777 | at | -1.12 | 0.206 | Mars2 | methionine-tRNA synthetase 2 (mitochondrial) |
| 1423667 | at | 1.01 | 0.858 | Mat2a | methionine adenosyltransferase II, alpha |
| 1448196 | at | -1.04 | 0.199 | Mat2b | methionine adenosyltransferase II, beta |
| 1419442 | at | 1.44 | 0.082 | Matn2 | matrilin 2 |
| 1423501 | at | -1.26 | 0.001 | Max | Max protein |
| 1427099 | at | -1.06 | 0.380 | Maz | MYC-associated zinc finger protein (purine-binding transcription factor) |
| 1451203 | at | 1.03 | 0.050 | Mb | myoglobin |
| 1451099 | at | 1.12 | 0.268 | Mbc2 | membrane bound C2 domain containing protein |
| 1417728 | at | 1.09 | 0.417 | Mbd3 | methyl-CpG binding domain protein 3 |
| 1424405 | at | -1.26 | 0.003 | Mbip | MAP3K12 binding inhibitory protein 1 |
| 1440315 | at | -1.28 | 0.175 | Mbnl1 | muscleblind-like 1 (Drosophila) |
| 1448641 | at | -1.24 | 0.144 | Mbtd1 | mbt domain containing 1 |
| 1448240 | at | -1.08 | 0.092 | Mbtps1 | membrane-bound transcription factor protease, site 1 |
| 1416357 | a_at | 1.36 | 0.000 | Mcam | melanoma cell adhesion molecule |
| 1454712 | at | -1.01 | 0.830 | Mcart1 | mitochondrial carrier triple repeat 1 |
| 1417227 | at | -1.27 | 0.005 | Mccc1 | methylcrotonoyl-Coenzyme A carboxylase 1 (alpha) |
| 1454840 | at | -1.48 | 0.001 | Mccc2 | methylcrotonoyl-Coenzyme A carboxylase 2 (beta) |

Supplementary table S1

| | | | | | |
|---|---|---|---|---|---|
| 1438477 | a_at | -1.34 | 0.000 | Mcee | methylmalonyl CoA epimerase |
| 1434140 | at | -1.20 | 0.025 | Mcf2l | mcf.2 transforming sequence-like |
| 1424024 | at | 1.07 | 0.129 | Mcfd2 | multiple coagulation factor deficiency 2 |
| 1448503 | at | 1.20 | 0.067 | Mcl1 | myeloid cell leukemia sequence 1 |
| 1448711 | at | -1.05 | 0.453 | Mcm3ap | minichromosome maintenance deficient 3 (S. cerevisiae) associated protein |
| 1416214 | at | 1.16 | 0.071 | Mcm4 | minichromosome maintenance deficient 4 homolog (S. cerevisiae) |
| 1416251 | at | 1.39 | 0.073 | Mcm6 | minichromosome maintenance deficient 6 (MIS5 homolog, S. pombe) (S. cerevisiae) |
| 1416671 | a_at | -1.01 | 0.873 | Mcoln1 | mucolipin 1 |
| 1416535 | at | -1.28 | 0.011 | Mcrs1 | microspherule protein 1 |
| 1425018 | at | 1.10 | 0.019 | Mcts1 | malignant T cell amplified sequence 1 |
| 1434837 | at | -1.02 | 0.642 | Mdc1 | mediator of DNA damage checkpoint 1 |
| 1427040 | at | 1.18 | 0.003 | Mdfic | MyoD family inhibitor domain containing |
| 1448172 | at | -1.04 | 0.461 | Mdh1 | malate dehydrogenase 1, NAD (soluble) |
| 1423605 | a_at | -1.01 | 0.877 | Mdm2 | transformed mouse 3T3 cell double minute 2 |
| 1419558 | at | 1.07 | 0.279 | Mdm4 | transformed mouse 3T3 cell double minute 4 |
| 1417675 | a_at | 1.27 | 0.012 | Mdn1 | midasin homolog (yeast) |
| 1429071 | at | -1.31 | 0.006 | Me3 | malic enzyme 3, NADP(+)-dependent, mitochondrial |
| 1416810 | at | 1.15 | 0.007 | Mea1 | male enhanced antigen 1 |
| 1438538 | at | -1.36 | 0.002 | Mecp2 | methyl CpG binding protein 2 |
| 1453197 | at | -1.00 | 0.975 | Med25 | mediator of RNA polymerase II transcription, subunit 25 homolog (yeast) |
| 1416508 | at | 1.05 | 0.323 | Med28 | mediator of RNA polymerase II transcription, subunit 28 homolog (yeast) |
| 1417387 | at | -1.18 | 0.002 | Med31 | mediator of RNA polymerase II transcription, subunit 31 homolog (yeast) |
| 1437957 | at | -1.01 | 0.720 | Med6 | mediator of RNA polymerase II transcription, subunit 6 homolog (yeast) |
| 1431423 | a_at | 1.14 | 0.042 | Med8 | mediator of RNA polymerase II transcription, subunit 8 homolog (yeast) |
| 1449448 | at | 1.18 | 0.203 | Med9 | mediator of RNA polymerase II transcription, subunit 9 homolog (yeast) |
| 1452347 | at | -1.33 | 0.022 | Mef2a | myocyte enhancer factor 2A |
| 1456267 | at | -1.05 | 0.578 | Mef2b | myocyte enhancer factor 2B |
| 1440431 | at | -1.38 | 0.053 | Meis1 | myeloid ecotropic viral integration site 1 |
| 1416347 | at | -1.01 | 0.951 | Men1 | multiple endocrine neoplasia 1 |
| 1417595 | at | 3.36 | 0.000 | Meox1 | mesenchyme homeobox 1 |
| 1424233 | at | 1.14 | 0.121 | Meox2 | mesenchyme homeobox 2 |
| 1422869 | at | 1.10 | 0.206 | Mertk | c-mer proto-oncogene tyrosine kinase |
| 1416181 | at | -1.02 | 0.441 | Mesdc2 | mesoderm development candiate 2 |
| 1423294 | at | 1.10 | 0.230 | Mest | mesoderm specific transcript |
| 1426835 | at | 1.08 | 0.197 | Metap1 | methionyl aminopeptidase 1 |
| 1423610 | at | 1.18 | 0.022 | Metap2 | methionine aminopeptidase 2 |
| 1452464 | a_at | -1.26 | 0.001 | Metapl1 | methionine aminopeptidase-like 1 |
| 1427100 | at | 1.01 | 0.963 | Metrn | meteorin, glial cell differentiation regulator |
| 1424356 | a_at | 1.03 | 0.557 | Metrnl | meteorin, glial cell differentiation regulator-like |
| 1460519 | a_at | -1.18 | 0.005 | Mettl5 | methyltransferase like 5 |
| 1434562 | at | -1.08 | 0.197 | Mfap3 | microfibrillar-associated protein 3 |

Supplementary table S1

| | | | | | |
|---|---|---|---|---|---|
| 1428804 | at | -1.38 | 0.000 | Mfap3l | microfibrillar-associated protein 3-like |
| 1424010 | at | 4.51 | 0.000 | Mfap4 | microfibrillar-associated protein 4 |
| 1418454 | at | 3.35 | 0.000 | Mfap5 | microfibrillar associated protein 5 |
| 1420911 | a_at | 1.32 | 0.061 | Mfge8 | milk fat globule-EGF factor 8 protein |
| 1429005 | at | -1.17 | 0.005 | Mfhas1 | malignant fibrous histiocytoma amplified sequence 1 |
| 1448131 | at | 1.06 | 0.006 | Mfn2 | mitofusin 2 |
| 1451247 | at | 1.10 | 0.889 | Mfsd1 | major facilitator superfamily domain containing 1 |
| 1434746 | at | -1.17 | 0.007 | Mga | MAX gene associated |
| 1423609 | a_at | 1.28 | 0.002 | Mgat1 | mannoside acetylglucosaminyltransferase 1 |
| 1435641 | at | 1.21 | 0.190 | Mgat4a | mannoside acetylglucosaminyltransferase 4, isoenzyme A |
| 1444458 | at | -1.48 | 0.010 | Mgea6 | meningioma expressed antigen 6 (coiled-coil proline-rich) |
| 1448416 | at | 1.30 | 0.000 | Mglap | matrix gamma-carboxyglutamate (gla) protein |
| 1450391 | a_at | -1.14 | 0.103 | Mgll | monoglyceride lipase |
| 1421309 | at | -2.85 | 0.000 | Mgmt | O-6-methylguanine-DNA methyltransferase |
| 1454645 | at | -1.22 | 0.000 | Mgrn1 | mahogunin, ring finger 1 |
| 1415897 | a_at | 1.00 | 0.981 | Mgst1 | microsomal glutathione S-transferase 1 |
| 1427219 | at | -1.28 | 0.003 | Mgst2 | microsomal glutathione S-transferase 2 |
| 1448300 | at | -1.40 | 0.000 | Mgst3 | microsomal glutathione S-transferase 3 |
| 1459984 | at | -1.31 | 0.090 | Mia3 | melanoma inhibitory activity 3 |
| 1454740 | at | 1.02 | 0.711 | Mib1 | mindbomb homolog 1 (Drosophila) |
| 1455685 | at | 1.69 | 0.006 | Mical2 | microtubule associated monoxygenase, calponin and LIM domain containing 2 |
| 1416840 | at | -1.18 | 0.022 | Mid1ip1 | Mid1 interacting protein 1 (gastrulation specific G12-like (zebrafish)) |
| 1449188 | at | 1.15 | 0.012 | Midn | midnolin |
| 1416335 | at | 1.17 | 0.074 | Mif | macrophage migration inhibitory factor |
| 1449362 | a_at | 1.24 | 0.014 | Mink1 | misshapen-like kinase 1 (zebrafish) |
| 1428290 | at | -1.24 | 0.003 | Mipep | mitochondrial intermediate peptidase |
| 1433903 | at | -1.03 | 0.621 | Mipol1 | mirror-image polydactyly gene 1 homolog (human) |
| 1418444 | a_at | 1.02 | 0.838 | Mir16 | membrane interacting protein of RGS16 |
| 1424717 | at | 1.10 | 0.065 | Mis12 | MIS12 homolog (yeast) |
| 1433606 | at | -1.00 | 0.942 | Mitc1 | MAD homolog 4 interacting transcription coactivator 1 |
| 1422025 | at | -1.54 | 0.005 | Mitf | microphthalmia-associated transcription factor |
| 1428725 | at | 1.05 | 0.427 | Miz1 | Msx-interacting-zinc finger |
| 1434408 | at | -1.05 | 0.289 | Mjd | Machado-Joseph disease (spinocerebellar ataxia 3, olivopontocerebellar ataxia 3, ataxin 3) homolog (human) |
| 1426817 | at | 1.15 | 0.274 | Mki67 | antigen identified by monoclonal antibody Ki 67 |
| 1424001 | at | 1.34 | 0.000 | Mki67ip | Mki67 (FHA domain) interacting nucleolar phosphoprotein |
| 1422627 | a_at | -1.03 | 0.562 | Mkks | McKusick-Kaufman syndrome protein |
| 1434900 | at | 1.04 | 0.659 | Mkl1 | MKL (megakaryoblastic leukemia)/myocardin-like 1 |
| 1435547 | at | -1.04 | 0.364 | Mkl2 | MKL/myocardin-like 2 |
| 1417630 | at | 1.13 | 0.174 | Mknk1 | MAP kinase-interacting serine/threonine kinase 1 |
| 1449029 | at | 1.45 | 0.045 | Mknk2 | MAP kinase-interacting serine/threonine kinase 2 |
| 1418589 | a_at | -1.32 | 0.014 | Mlf1 | myeloid leukemia factor 1 |

Supplementary table S1

| | | | | | |
|---|---|---|---|---|---|
| 1417360 | at | 1.00 | 0.984 | Mlh1 | mutL homolog 1 (E. coli) |
| 1420870 | at | -1.29 | 0.064 | Mllt10 | myeloid/lymphoid or mixed lineage-leukemia translocation to 10 homolog (Drosophila) |
| 1418135 | at | -1.08 | 0.094 | Mllt2h | homolog of human MLLT2 unidentified gene |
| 1435017 | at | -1.13 | 0.001 | Mllt6 | myeloid/lymphoid or mixed lineage-leukemia translocation to 6 homolog (Drosophila) |
| 1456753 | at | -1.01 | 0.846 | Mllt7 | myeloid/lymphoid or mixed lineage-leukemia translocation to 7 homolog (Drosophila) |
| 1426371 | at | 1.12 | 0.306 | Mlstd2 | male sterility domain containing 2 |
| 1449964 | a_at | -1.54 | 0.000 | Mlycd | malonyl-CoA decarboxylase |
| 1417857 | at | -1.10 | 0.235 | Mmaa | methylmalonic aciduria (cobalamin deficiency) type A |
| 1435097 | at | -1.18 | 0.072 | Mmab | methylmalonic aciduria (cobalamin deficiency) type B homolog (human) |
| 1423488 | at | -1.15 | 0.300 | Mmd | monocyte to macrophage differentiation-associated |
| 1422597 | at | -1.41 | 0.026 | Mmp15 | matrix metalloproteinase 15 |
| 1416136 | at | 2.04 | 0.000 | Mmp2 | matrix metalloproteinase 2 |
| 1417281 | a_at | 2.45 | 0.000 | Mmp23 | matrix metalloproteinase 23 |
| 1448806 | at | 1.18 | 0.028 | Mmp24 | matrix metalloproteinase 24 |
| 1418945 | at | 1.63 | 0.005 | Mmp3 | matrix metalloproteinase 3 |
| 1437123 | at | 1.05 | 0.681 | Mmrn2 | multimerin 2 |
| 1417967 | at | -1.15 | 0.036 | Mms19l | MMS19 (MET18 S. cerevisiae)-like |
| 1426924 | at | 1.10 | 0.093 | Mnab | membrane associated DNA binding protein |
| 1432177 | a_at | 1.11 | 0.064 | Mnat1 | menage a trois 1 |
| 1418192 | at | 1.08 | 0.481 | Mnt | max binding protein |
| 1448787 | at | 1.13 | 0.014 | Moap1 | modulator of apoptosis 1 |
| 1423782 | at | 1.31 | 0.001 | Mobk1b | MOB1, Mps One Binder kinase activator-like 1B (yeast) |
| 1429352 | at | -1.03 | 0.408 | Mocos | molybdenum cofactor sulfurase |
| 1424309 | a_at | 1.01 | 0.894 | Mocs2 | molybdenum cofactor synthesis 2 |
| 1428712 | at | 1.03 | 0.707 | Mon1b | MON1 homolog b (yeast) |
| 1454800 | at | -1.12 | 0.071 | Mopt | protein containing single MORN motif in testis |
| 1415778 | at | 1.21 | 0.007 | Morf4l2 | mortality factor 4 like 2 |
| 1424280 | at | -1.06 | 0.340 | Mospd1 | motile sperm domain containing 1 |
| 1424124 | at | 1.22 | 0.005 | Mospd2 | motile sperm domain containing 2 |
| 1419340 | at | -1.42 | 0.000 | Mov10l1 | Moloney leukemia virus 10-like 1 |
| 1421104 | at | 1.05 | 0.658 | Mpa2 | macrophage activation 2 |
| 1416104 | at | 1.16 | 0.244 | Mpdu1 | mannose-P-dolichol utilization defect 1 |
| 1418664 | at | 1.10 | 0.023 | Mpdz | multiple PDZ domain protein |
| 1427076 | at | 1.80 | 0.004 | Mpeg1 | macrophage expressed gene 1 |
| 1417572 | at | 1.05 | 0.635 | Mpg | N-methylpurine-DNA glycosylase |
| 1429080 | at | 1.16 | 0.168 | Mphosph10 | M-phase phosphoprotein 10 (U3 small nucleolar ribonucleoprotein) |
| 1431053 | at | -1.13 | 0.279 | Mphosph9 | M-phase phosphoprotein 9 |
| 1451540 | at | -1.05 | 0.296 | Mpi1 | mannose phosphate isomerase 1 |
| 1450919 | at | 1.09 | 0.284 | Mpp1 | membrane protein, palmitoylated |
| 1419077 | at | -1.33 | 0.012 | Mpp3 | membrane protein, palmitoylated 3 (MAGUK p55 subfamily member 3) |
| 1435661 | at | 1.43 | 0.067 | Mpp4 | membrane protein, palmitoylated 4 (MAGUK p55 subfamily member 4) |

Supplementary table S1

| | | | | | |
|---|---|---|---|---|---|
| 1421064 | at | 1.17 | 0.215 | Mpp5 | membrane protein, palmitoylated 5 (MAGUK p55 subfamily member 5) |
| 1449348 | at | 1.08 | 0.272 | Mpp6 | membrane protein, palmitoylated 6 (MAGUK p55 subfamily member 6) |
| 1455179 | at | -1.02 | 0.810 | Mpp7 | membrane protein, palmitoylated 7 (MAGUK p55 subfamily member 7) |
| 1460674 | at | -1.01 | 0.911 | Mpra | membrane progestin receptor alpha |
| 1418356 | at | -1.60 | 0.000 | Mpst | mercaptopyruvate sulfurtransferase |
| 1420387 | at | -1.63 | 0.000 | Mpv17 | Mpv17 transgene, kidney disease mutant |
| 1428168 | at | 1.05 | 0.460 | Mpzl1 | myelin protein zero-like 1 |
| 1421899 | a_at | 1.08 | 0.104 | Mr1 | major histocompatibility complex, class I-related |
| 1451371 | at | 1.12 | 0.055 | Mrap | melanocortin 2 receptor accessory protein |
| 1450430 | at | 1.31 | 0.001 | Mrc1 | mannose receptor, C type 1 |
| 1421045 | at | 1.97 | 0.001 | Mrc2 | mannose receptor, C type 2 |
| 1416496 | at | 1.03 | 0.129 | Mrfap1 | Morf4 family associated protein 1 |
| 1440091 | at | -1.49 | 0.067 | Mrg1 | myeloid ecotropic viral integration site-related gene 1 |
| 1423351 | at | -1.11 | 0.098 | Mrpl1 | mitochondrial ribosomal protein L1 |
| 1418112 | at | -1.07 | 0.037 | Mrpl10 | mitochondrial ribosomal protein L10 |
| 1417918 | at | -1.04 | 0.414 | Mrpl11 | mitochondrial ribosomal protein L11 |
| 1452048 | at | -1.05 | 0.464 | Mrpl12 | mitochondrial ribosomal protein L12 |
| 1424204 | at | 1.02 | 0.563 | Mrpl13 | mitochondrial ribosomal protein L13 |
| 1451307 | at | -1.52 | 0.000 | Mrpl14 | mitochondrial ribosomal protein L14 |
| 1439988 | at | -1.29 | 0.094 | Mrpl15 | mitochondrial ribosomal protein L15 |
| 1450880 | at | -1.06 | 0.071 | Mrpl16 | mitochondrial ribosomal protein L16 |
| 1450866 | a_at | -1.08 | 0.160 | Mrpl17 | mitochondrial ribosomal protein L17 |
| 1448373 | at | -1.01 | 0.802 | Mrpl18 | mitochondrial ribosomal protein L18 |
| 1453571 | at | -1.17 | 0.259 | Mrpl19 | mitochondrial ribosomal protein L19 |
| 1448811 | at | -1.21 | 0.001 | Mrpl2 | mitochondrial ribosomal protein L2 |
| 1416093 | a_at | 1.06 | 0.369 | Mrpl20 | mitochondrial ribosomal protein L20 |
| 1435995 | at | -1.20 | 0.002 | Mrpl22 | mitochondrial ribosomal protein L22 |
| 1416948 | at | -1.21 | 0.000 | Mrpl23 | mitochondrial ribosomal protein L23 |
| 1428709 | a_at | -1.07 | 0.091 | Mrpl24 | mitochondrial ribosomal protein L24 |
| 1415690 | at | -1.04 | 0.263 | Mrpl27 | mitochondrial ribosomal protein L27 |
| 1416284 | at | -1.10 | 0.257 | Mrpl28 | mitochondrial ribosomal protein L28 |
| 1423857 | at | -1.02 | 0.457 | Mrpl30 | mitochondrial ribosomal protein L30 |
| 1424372 | at | -1.04 | 0.329 | Mrpl32 | mitochondrial ribosomal protein L32 |
| 1448190 | at | 1.05 | 0.724 | Mrpl33 | mitochondrial ribosomal protein L33 |
| 1438014 | at | -1.05 | 0.357 | Mrpl34 | mitochondrial ribosomal protein L34 |
| 1440989 | at | -1.07 | 0.384 | Mrpl35 | mitochondrial ribosomal protein L35 |
| 1422819 | at | -1.15 | 0.001 | Mrpl36 | mitochondrial ribosomal protein L36 |
| 1417966 | at | -1.47 | 0.000 | Mrpl39 | mitochondrial ribosomal protein L39 |
| 1416510 | at | -1.17 | 0.030 | Mrpl4 | mitochondrial ribosomal protein L4 |
| 1448849 | at | -1.01 | 0.795 | Mrpl40 | mitochondrial ribosomal protein L40 |
| 1415681 | at | 1.08 | 0.177 | Mrpl43 | mitochondrial ribosomal protein L43 |

Supplementary table S1

| | | | | | |
|---|---|---|---|---|---|
| 1426651 | at | 1.08 | 0.101 | Mrpl44 | mitochondrial ribosomal protein L44 |
| 1423492 | at | -1.23 | 0.043 | Mrpl45 | mitochondrial ribosomal protein L45 |
| 1449004 | at | -1.00 | 0.977 | Mrpl46 | mitochondrial ribosomal protein L46 |
| 1443915 | at | -1.38 | 0.042 | Mrpl47 | mitochondrial ribosomal protein L47 |
| 1428302 | at | -1.14 | 0.119 | Mrpl48 | mitochondrial ribosomal protein L48 |
| 1423219 | a_at | -1.13 | 0.473 | Mrpl49 | mitochondrial ribosomal protein L49 |
| 1451266 | at | -1.09 | 0.707 | Mrpl50 | mitochondrial ribosomal protein L50 |
| 1416878 | at | -1.17 | 0.031 | Mrpl51 | mitochondrial ribosomal protein L51 |
| 1460701 | a_at | 1.06 | 0.260 | Mrpl52 | mitochondrial ribosomal protein L52 |
| 1451178 | at | 1.17 | 0.005 | Mrpl53 | mitochondrial ribosomal protein L53 |
| 1448699 | at | 1.02 | 0.817 | Mrpl54 | mitochondrial ribosomal protein L54 |
| 1429453 | a_at | -1.10 | 0.125 | Mrpl55 | mitochondrial ribosomal protein L55 |
| 1427297 | at | -1.08 | 0.375 | Mrpl9 | mitochondrial ribosomal protein L9 |
| 1428153 | at | 1.03 | 0.529 | Mrps10 | mitochondrial ribosomal protein S10 |
| 1460195 | at | -1.15 | 0.028 | Mrps11 | mitochondrial ribosomal protein S11 |
| 1448869 | a_at | -1.06 | 0.184 | Mrps16 | mitochondrial ribosomal protein S16 |
| 1453728 | a_at | -1.02 | 0.542 | Mrps17 | mitochondrial ribosomal protein S17 |
| 1416984 | at | -1.11 | 0.070 | Mrps18a | mitochondrial ribosomal protein S18A |
| 1451164 | a_at | 1.11 | 0.193 | Mrps18b | mitochondrial ribosomal protein S18B |
| 1427901 | at | 1.00 | 0.932 | Mrps18c | mitochondrial ribosomal protein S18C |
| 1420846 | at | 1.03 | 0.488 | Mrps2 | mitochondrial ribosomal protein S2 |
| 1422451 | at | -1.16 | 0.000 | Mrps21 | mitochondrial ribosomal protein S21 |
| 1416595 | at | -1.15 | 0.002 | Mrps22 | mitochondrial ribosomal protein S22 |
| 1421874 | a_at | 1.25 | 0.015 | Mrps23 | mitochondrial ribosomal protein S23 |
| 1433878 | at | 1.01 | 0.810 | Mrps26 | mitochondrial ribosomal protein S26 |
| 1433646 | at | 1.07 | 0.232 | Mrps27 | mitochondrial ribosomal protein S27 |
| 1452585 | at | -1.21 | 0.000 | Mrps28 | mitochondrial ribosomal protein S28 |
| 1427158 | at | -1.11 | 0.034 | Mrps30 | mitochondrial ribosomal protein S30 |
| 1417737 | at | -1.18 | 0.001 | Mrps31 | mitochondrial ribosomal protein S31 |
| 1427173 | a_at | -1.09 | 0.001 | Mrps33 | mitochondrial ribosomal protein S33 |
| 1421971 | a_at | 1.01 | 0.899 | Mrps34 | mitochondrial ribosomal protein S34 |
| 1452111 | at | -1.27 | 0.006 | Mrps35 | mitochondrial ribosomal protein S35 |
| 1423242 | at | -1.24 | 0.000 | Mrps36 | mitochondrial ribosomal protein S36 |
| 1448488 | at | -1.11 | 0.248 | Mrps5 | mitochondrial ribosomal protein S5 |
| 1424440 | at | -1.01 | 0.729 | Mrps6 | mitochondrial ribosomal protein S6 |
| 1453725 | a_at | 1.02 | 0.611 | Mrps7 | mitchondrial ribosomal protein S7 |
| 1426364 | at | -1.21 | 0.000 | Mrrf | mitochondrial ribosome recycling factor |
| 1418990 | at | 1.19 | 0.018 | Ms4a4d | membrane-spanning 4-domains, subfamily A, member 4D |
| 1418826 | at | 1.39 | 0.008 | Ms4a6b | membrane-spanning 4-domains, subfamily A, member 6B |
| 1417750 | a_at | -1.10 | 0.341 | Mscp | mitochondrial solute carrier protein |
| 1416988 | at | -1.16 | 0.048 | Msh2 | mutS homolog 2 (E. coli) |

Supplementary table S1

| | | | | | |
|---|---|---|---|---|---|
| 1416915 | at | -1.03 | 0.684 | Msh6 | mutS homolog 6 (E. coli) |
| 1458542 | at | -1.31 | 0.053 | Msi2h | Musashi homolog 2 (Drosophila) |
| 1448645 | at | -1.13 | 0.010 | Msl31 | male-specific lethal-3 homolog 1 (Drosophila) |
| 1421814 | at | 1.34 | 0.000 | Msn | moesin |
| 1448856 | a_at | -1.03 | 0.711 | Msra | methionine sulfoxide reductase A |
| 1424433 | at | -1.23 | 0.001 | Msrb2 | methionine sulfoxide reductase B2 |
| 1454997 | at | 1.06 | 0.126 | Msrb3 | methionine sulfoxide reductase B3 |
| 1428942 | at | 2.10 | 0.000 | Mt2 | metallothionein 2 |
| 1417295 | at | 1.08 | 0.140 | Mta1 | metastasis associated 1 |
| 1449897 | a_at | -1.10 | 0.100 | Mtcp1 | mature T-cell proliferation 1 |
| 1455129 | at | 1.12 | 0.124 | Mtdh | Metadherin |
| 1439478 | at | -1.45 | 0.000 | Mte1 | mitochondrial acyl-CoA thioesterase 1 |
| 1429698 | at | -1.07 | 0.455 | MTERF | mitochondrial transcription termination factor 1 |
| 1428979 | at | -1.20 | 0.016 | Mtf1 | metal response element binding transcription factor 1 |
| 1418516 | at | -1.11 | 0.238 | Mtf2 | metal response element binding transcription factor 2 |
| 1434087 | at | 1.12 | 0.135 | Mthfr | 5,10-methylenetetrahydrofolate reductase |
| 1460257 | a_at | 1.28 | 0.007 | Mthfs | 5, 10-methenyltetrahydrofolate synthetase |
| 1415685 | at | -1.07 | 0.369 | Mtif2 | mitochondrial translational initiation factor 2 |
| 1416934 | at | -1.09 | 0.540 | Mtm1 | X-linked myotubular myopathy gene 1 |
| 1421880 | at | 1.04 | 0.765 | Mtmr1 | myotubularin related protein 1 |
| 1425460 | at | 1.03 | 0.607 | Mtmr2 | myotubularin related protein 2 |
| 1418150 | at | -1.50 | 0.002 | Mtmr4 | myotubularin related protein 4 |
| 1451726 | at | 1.40 | 0.017 | Mtmr6 | myotubularin related protein 6 |
| 1416991 | at | -1.16 | 0.044 | Mto1 | mitochondrial translation optimization 1 homolog (S. cerevisiae) |
| 1437457 | a_at | 1.34 | 0.003 | Mtpn | myotrophin |
| 1438025 | at | -1.16 | 0.069 | Mtrf1l | mitochondrial translational release factor 1-like |
| 1452110 | at | 1.00 | 0.961 | Mtrr | 5-methyltetrahydrofolate-homocysteine methyltransferase reductase |
| 1434036 | at | -1.25 | 0.001 | Mtss1 | metastasis suppressor 1 |
| 1419400 | at | 1.20 | 0.056 | Mttp | microsomal triglyceride transfer protein |
| 1436502 | at | 1.09 | 0.179 | Mtus1 | mitochondrial tumor suppressor 1 |
| 1418521 | a_at | -1.25 | 0.081 | Mtx1 | metaxin 1 |
| 1447871 | at | 1.15 | 0.463 | Mtx2 | metaxin 2 |
| 1434806 | at | -1.01 | 0.942 | Mtx3 | metaxin 3 |
| 1427201 | at | -1.03 | 0.698 | Mustn1 | musculoskeletal, embryonic nuclear protein 1 |
| 1416838 | at | -1.39 | 0.000 | Mut | methylmalonyl-Coenzyme A mutase |
| 1448979 | at | 1.08 | 0.153 | Muted | muted |
| 1448618 | at | 1.28 | 0.177 | Mvp | major vault protein |
| 1438847 | at | 1.19 | 0.180 | Mxd3 | Max dimerization protein 3 |
| 1434379 | at | -1.05 | 0.714 | Mxd4 | Max dimerization protein 4 |
| 1450376 | at | -1.06 | 0.250 | Mxi1 | Max interacting protein 1 |
| 1423321 | at | -1.07 | 0.148 | Myadm | myeloid-associated differentiation marker |

Supplementary table S1

| | | | | | |
|---|---|---|---|---|---|
| 1455736 | at | 2.55 | 0.009 | Mybpc2 | myosin binding protein C, fast-type |
| 1418551 | at | 1.07 | 0.026 | Mybpc3 | myosin binding protein C, cardiac |
| 1427938 | at | 1.02 | 0.628 | Mycbp | c-myc binding protein |
| 1438048 | at | -1.17 | 0.069 | Myct1 | myc target 1 |
| 1448325 | at | 1.34 | 0.003 | Myd116 | myeloid differentiation primary response gene 116 |
| 1419272 | at | 1.20 | 0.051 | Myd88 | myeloid differentiation primary response gene 88 |
| 1425350 | a_at | 1.03 | 0.603 | Myef2 | myelin basic protein expression factor 2, repressor |
| 1452740 | at | 1.56 | 0.000 | Myh10 | myosin, heavy polypeptide 10, non-muscle |
| 1448962 | at | -1.41 | 0.004 | Myh11 | myosin, heavy polypeptide 11, smooth muscle |
| 1428835 | at | 1.06 | 0.599 | Myh14 | myosin, heavy polypeptide 14 |
| 1448826 | at | -1.11 | 0.455 | Myh6 | myosin, heavy polypeptide 6, cardiac muscle, alpha |
| 1448553 | at | 4.87 | 0.000 | Myh7 | myosin, heavy polypeptide 7, cardiac muscle, beta |
| 1417472 | at | 1.14 | 0.156 | Myh9 | myosin, heavy polypeptide 9, non-muscle |
| 1452651 | a_at | 1.37 | 0.003 | Myl1 | myosin, light polypeptide 1 |
| 1448394 | at | -1.03 | 0.130 | Myl2 | myosin, light polypeptide 2, regulatory, cardiac, slow |
| 1428266 | at | 1.02 | 0.248 | Myl3 | myosin, light polypeptide 3 |
| 1422580 | at | 1.32 | 0.040 | Myl4 | myosin, light polypeptide 4 |
| 1424269 | a_at | 1.16 | 0.043 | Myl6 | myosin, light polypeptide 6, alkali, smooth muscle and non-muscle |
| 1428608 | at | 1.08 | 0.152 | Mylc2b | myosin light chain, regulatory B |
| 1424988 | at | -1.33 | 0.045 | Mylip | myosin regulatory light chain interacting protein |
| 1425505 | at | -1.22 | 0.101 | Mylk | myosin, light polypeptide kinase |
| 1450204 | a_at | 1.10 | 0.307 | Mynn | myoneurin |
| 1454731 | at | -1.08 | 0.044 | Myo10 | myosin X |
| 1419648 | at | 1.09 | 0.291 | Myo1c | myosin IC |
| 1450543 | at | 1.23 | 0.002 | Myo1h | myosin 1H |
| 1436051 | at | 1.36 | 0.004 | Myo5a | myosin Va |
| 1452298 | a_at | 1.16 | 0.062 | Myo5b | myosin Vb |
| 1424933 | at | -1.67 | 0.004 | Myo5c | myosin VC |
| 1421385 | a_at | 1.41 | 0.004 | Myo7a | myosin VIIa |
| 1436307 | at | 1.15 | 0.019 | Myo9a | myosin IXa |
| 1418031 | at | 1.63 | 0.000 | Myo9b | myosin IXb |
| 1425978 | at | -1.53 | 0.002 | Myocd | myocardin |
| 1452880 | at | -1.04 | 0.668 | Myohd1 | myosin head domain containing 1 |
| 1420693 | at | 1.14 | 0.000 | Myom1 | myomesin 1 |
| 1450917 | at | 1.10 | 0.043 | Myom2 | myomesin 2 |
| 1418769 | at | 1.10 | 0.000 | Myoz2 | myozenin 2 |
| 1460601 | at | -1.33 | 0.009 | Myrip | myosin VIIA and Rab interacting protein |
| 1433433 | at | -1.16 | 0.071 | Myst2 | MYST histone acetyltransferase 2 |
| 1423508 | at | -1.19 | 0.125 | Myst4 | MYST histone acetyltransferase monocytic leukemia 4 |
| 1417624 | at | 1.01 | 0.850 | Nab1 | Ngfi-A binding protein 1 |
| 1448430 | a_at | 1.06 | 0.038 | Naca | nascent polypeptide-associated complex alpha polypeptide |

Supplementary table S1

| | | | | | |
|---|---|---|---|---|---|
| 1449043 | at | 1.01 | 0.914 | Naga | N-acetyl galactosaminidase, alpha |
| 1419509 | a_at | 1.14 | 0.015 | Nagk | N-acetylglucosamine kinase |
| 1456211 | at | 1.25 | 0.007 | Nalp10 | NACHT, leucine rich repeat and PYD containing 10 |
| 1420476 | a_at | 1.14 | 0.259 | Nap1l1 | nucleosome assembly protein 1-like 1 |
| 1418500 | at | -1.25 | 0.061 | Nap1l3 | nucleosome assembly protein 1-like 3 |
| 1448476 | at | -1.02 | 0.399 | Nap1l4 | nucleosome assembly protein 1-like 4 |
| 1448243 | at | -1.08 | 0.002 | Napa | N-ethylmaleimide sensitive fusion protein attachment protein alpha |
| 1441076 | at | -1.13 | 0.335 | Napb | N-ethylmaleimide sensitive fusion protein attachment protein beta |
| 1424069 | at | 1.06 | 0.220 | Napg | N-ethylmaleimide sensitive fusion protein attachment protein gamma |
| 1450452 | a_at | -1.02 | 0.824 | Narfl | nuclear prelamin A recognition factor-like |
| 1429057 | at | -1.15 | 0.172 | Narg1l | NMDA receptor regulated 1-like |
| 1426536 | at | -1.13 | 0.032 | Narg2 | NMDA receptor-regulated gene 2 |
| 1428666 | at | 1.29 | 0.001 | Nars | asparaginyl-tRNA synthetase |
| 1442728 | at | -1.00 | 0.998 | Nasp | nuclear autoantigenic sperm protein (histone-binding) |
| 1418244 | at | -1.16 | 0.004 | Nat5 | N-acetyltransferase 5 (ARD1 homolog, S. cerevisiae) |
| 1419213 | at | -1.01 | 0.895 | Nat6 | N-acetyltransferase 6 |
| 1436907 | at | 1.26 | 0.003 | Nav1 | neuron navigator 1 |
| 1452251 | at | 1.17 | 0.039 | Nbea | neurobeachin |
| 1442027 | at | 1.10 | 0.111 | Nbeal1 | neurobeachin like 1 |
| 1448746 | at | -1.29 | 0.011 | Nbn | nibrin |
| 1451897 | a_at | -1.12 | 0.094 | Nbr1 | neighbor of Brca1 gene 1 |
| 1426865 | a_at | -1.41 | 0.055 | Ncam1 | neural cell adhesion molecule 1 |
| 1421592 | at | 1.62 | 0.029 | Ncam2 | neural cell adhesion molecule 2 |
| 1426307 | at | 1.22 | 0.001 | Ncb5or | NADPH cytochrome B5 oxidoreductase |
| 1450847 | at | -1.07 | 0.225 | Ncbp2 | nuclear cap binding protein subunit 2 |
| 1416853 | at | 1.13 | 0.052 | Ncdn | neurochondrin |
| 1448561 | at | 1.28 | 0.019 | Ncf2 | neutrophil cytosolic factor 2 |
| 1424543 | at | -1.09 | 0.371 | Nck1 | non-catalytic region of tyrosine kinase adaptor protein 1 |
| 1416796 | at | -1.06 | 0.170 | Nck2 | non-catalytic region of tyrosine kinase adaptor protein 2 |
| 1452196 | a_at | 1.16 | 0.045 | Nckap1 | NCK-associated protein 1 |
| 1415773 | at | 1.13 | 0.118 | Ncl | nucleolin |
| 1434515 | at | -1.26 | 0.016 | Ncoa1 | nuclear receptor coactivator 1 |
| 1424055 | at | -1.06 | 0.511 | Ncoa5 | nuclear receptor coactivator 5 |
| 1423374 | at | 1.05 | 0.392 | Ncoa6 | nuclear receptor coactivator 6 |
| 1454809 | at | 1.24 | 0.041 | Ncoa7 | nuclear receptor coactivator 7 |
| 1423201 | at | -1.07 | 0.283 | Ncor1 | nuclear receptor co-repressor 1 |
| 1448893 | at | 1.06 | 0.242 | Ncor2 | nuclear receptor co-repressor 2 |
| 1435737 | a_at | -1.06 | 0.307 | Nde1 | nuclear distribution gene E homolog 1 (A nidulans) |
| 1422568 | at | 1.10 | 0.050 | Ndel1 | nuclear distribution gene E-like homolog 1 (A. nidulans) |
| 1424821 | at | 1.03 | 0.289 | Ndfip1 | Nedd4 family interacting protein 1 |
| 1452066 | a_at | 1.18 | 0.023 | Ndfip2 | Nedd4 family interacting protein 2 |

Supplementary table S1

| | | | | | |
|---|---|---|---|---|---|
| 1435382 | at | 1.09 | 0.251 | Ndn | necdin |
| 1450976 | at | 1.10 | 0.367 | Ndrg1 | N-myc downstream regulated gene 1 |
| 1448154 | at | 1.03 | 0.118 | Ndrg2 | N-myc downstream regulated gene 2 |
| 1417663 | a_at | 1.05 | 0.171 | Ndrg3 | N-myc downstream regulated gene 3 |
| 1417931 | at | -1.02 | 0.784 | Ndst2 | N-deacetylase/N-sulfotransferase (heparan glucosaminyl) 2 |
| 1422241 | a_at | -1.03 | 0.225 | Ndufa1 | NADH dehydrogenase (ubiquinone) 1 alpha subcomplex, 1 |
| 1429708 | at | -1.00 | 0.918 | Ndufa11 | NADH dehydrogenase (ubiquinone) 1 alpha subcomplex 11 |
| 1428464 | at | -1.03 | 0.117 | Ndufa3 | NADH dehydrogenase (ubiquinone) 1 alpha subcomplex, 3 |
| 1424085 | at | -1.07 | 0.023 | Ndufa4 | NADH dehydrogenase (ubiquinone) 1 alpha subcomplex, 4 |
| 1417286 | at | -1.10 | 0.003 | Ndufa5 | NADH dehydrogenase (ubiquinone) 1 alpha subcomplex, 5 |
| 1448427 | at | -1.20 | 0.006 | Ndufa6 | NADH dehydrogenase (ubiquinone) 1 alpha subcomplex, 6 (B14) |
| 1450818 | a_at | -1.17 | 0.032 | Ndufa7 | NADH dehydrogenase (ubiquinone) 1 alpha subcomplex, 7 (B14.5a) |
| 1423692 | at | -1.18 | 0.003 | Ndufa8 | NADH dehydrogenase (ubiquinone) 1 alpha subcomplex, 8 |
| 1416663 | at | -1.14 | 0.202 | Ndufa9 | NADH dehydrogenase (ubiquinone) 1 alpha subcomplex, 9 |
| 1435934 | at | -1.30 | 0.030 | Ndufab1 | NADH dehydrogenase (ubiquinone) 1, alpha/beta subcomplex, 1 |
| 1423711 | at | 1.05 | 0.208 | Ndufaf1 | NADH dehydrogenase (ubiquinone) 1 alpha subcomplex, assembly factor 1 |
| 1428322 | a_at | -1.02 | 0.119 | Ndufb10 | NADH dehydrogenase (ubiquinone) 1 beta subcomplex, 10 |
| 1448483 | a_at | -1.18 | 0.000 | Ndufb2 | NADH dehydrogenase (ubiquinone) 1 beta subcomplex, 2 |
| 1416547 | at | 1.08 | 0.039 | Ndufb3 | NADH dehydrogenase (ubiquinone) 1 beta subcomplex 3 |
| 1428075 | at | -1.07 | 0.002 | Ndufb4 | NADH dehydrogenase (ubiquinone) 1 beta subcomplex 4 |
| 1448589 | at | -1.07 | 0.135 | Ndufb5 | NADH dehydrogenase (ubiquinone) 1 beta subcomplex, 5 |
| 1448331 | at | -1.13 | 0.046 | Ndufb7 | NADH dehydrogenase (ubiquinone) 1 beta subcomplex, 7 |
| 1458404 | at | -1.10 | 0.366 | Ndufb8 | NADH dehydrogenase (ubiquinone) 1 beta subcomplex 8 |
| 1452184 | at | -1.02 | 0.405 | Ndufb9 | NADH dehydrogenase (ubiquinone) 1 beta subcomplex, 9 |
| 1416285 | at | -1.02 | 0.631 | Ndufc1 | NADH dehydrogenase (ubiquinone) 1, subcomplex unknown, 1 |
| 1416366 | at | -1.06 | 0.023 | Ndufc2 | NADH dehydrogenase (ubiquinone) 1, subcomplex unknown, 2 |
| 1434856 | at | -1.05 | 0.658 | Ndufs1 | NADH dehydrogenase (ubiquinone) Fe-S protein 1 |
| 1423737 | at | -1.04 | 0.003 | Ndufs3 | NADH dehydrogenase (ubiquinone) Fe-S protein 3 |
| 1448959 | at | -1.30 | 0.000 | Ndufs4 | NADH dehydrogenase (ubiquinone) Fe-S protein 4 |
| 1433603 | at | -1.08 | 0.011 | Ndufs6 | NADH dehydrogenase (ubiquinone) Fe-S protein 6 |
| 1451312 | at | -1.20 | 0.000 | Ndufs7 | NADH dehydrogenase (ubiquinone) Fe-S protein 7 |
| 1423908 | at | -1.19 | 0.000 | Ndufs8 | NADH dehydrogenase (ubiquinone) Fe-S protein 8 |
| 1415967 | at | -1.15 | 0.000 | Ndufv1 | NADH dehydrogenase (ubiquinone) flavoprotein 1 |
| 1428179 | at | -1.10 | 0.009 | Ndufv2 | NADH dehydrogenase (ubiquinone) flavoprotein 2 |
| 1451846 | at | -1.00 | 0.986 | Nebl | nebulette |
| 1450899 | at | -1.10 | 0.794 | Nedd1 | neural precursor cell expressed, developmentally down-regulated gene 1 |
| 1423269 | a_at | -1.45 | 0.003 | Nedd4l | neural precursor cell expressed, developmentally down-regulated gene 4-like |
| 1423715 | a_at | -1.03 | 0.386 | Nedd8 | neural precursor cell expressed, developmentally down-regulated gene 8 |
| 1422818 | at | 1.39 | 0.001 | Nedd9 | neural precursor cell expressed, developmentally down-regulated gene 9 |
| 1434267 | at | -1.03 | 0.425 | Nek1 | NIMA (never in mitosis gene a)-related expressed kinase 1 |
| 1418947 | at | 1.15 | 0.017 | Nek3 | NIMA (never in mitosis gene a)-related expressed kinase 3 |

Supplementary table S1

| | | | | | |
|---|---|---|---|---|---|
| 1416816 | at | 1.06 | 0.121 | Nek7 | NIMA (never in mitosis gene a)-related expressed kinase 7 |
| 1424094 | at | 1.08 | 0.005 | Nek9 | NIMA (never in mitosis gene a)-related expressed kinase 9 |
| 1440789 | at | -1.35 | 0.070 | Neo1 | neogenin |
| 1418289 | at | 1.14 | 0.224 | Nes | nestin |
| 1419339 | at | -1.08 | 0.195 | Neu3 | neuraminidase 3 |
| 1435649 | at | 1.15 | 0.021 | Nexn | nexilin |
| 1421820 | a_at | -1.01 | 0.902 | Nf2 | neurofibromatosis 2 |
| 1438999 | a_at | 1.01 | 0.913 | Nfat5 | nuclear factor of activated T-cells 5 |
| 1417621 | at | -1.13 | 0.355 | Nfatc1 | nuclear factor of activated T-cells, cytoplasmic, calcineurin-dependent 1 |
| 1434836 | at | -1.04 | 0.648 | Nfatc2ip | nuclear factor of activated T-cells, cytoplasmic, calcineurin-dependent 2 interacting protein |
| 1416331 | a_at | 1.05 | 0.104 | Nfe2l1 | nuclear factor, erythroid derived 2,-like 1 |
| 1416543 | at | 1.19 | 0.004 | Nfe2l2 | nuclear factor, erythroid derived 2, like 2 |
| 1438236 | at | -1.42 | 0.013 | Nfia | nuclear factor I/A |
| 1438244 | at | -1.22 | 0.087 | Nfib | nuclear factor I/B |
| 1418932 | at | 1.20 | 0.423 | Nfil3 | nuclear factor, interleukin 3, regulated |
| 1427705 | a_at | -1.08 | 0.273 | Nfkb1 | nuclear factor of kappa light chain gene enhancer in B-cells 1, p105 |
| 1457404 | at | 1.84 | 0.011 | Nfkbiz | nuclear factor of kappa light polypeptide gene enhancer in B-cells inhibitor, zeta |
| 1434753 | at | -1.09 | 0.083 | Nfrkb | nuclear factor related to kappa B binding protein |
| 1416373 | at | -1.33 | 0.001 | Nfs1 | nitrogen fixation gene 1 (S. cerevisiae) |
| 1428248 | at | -1.05 | 0.361 | Nfx1 | nuclear transcription factor, X-box binding 1 |
| 1452560 | a_at | 1.06 | 0.441 | Nfya | nuclear transcription factor-Y alpha |
| 1419267 | at | -1.03 | 0.526 | Nfyb | nuclear transcription factor-Y beta |
| 1446288 | at | -1.45 | 0.015 | Nfyc | nuclear transcription factor-Y gamma |
| 1419675 | at | 1.02 | 0.815 | Ngfb | nerve growth factor, beta |
| 1417827 | at | 1.04 | 0.423 | Ngly1 | N-glycanase 1 |
| 1435213 | at | -1.06 | 0.545 | Nhlrc1 | NHL repeat containing 1 |
| 1433929 | at | 1.05 | 0.315 | Nhlrc2 | NHL repeat containing 2 |
| 1416973 | at | -1.01 | 0.730 | Nhp2l1 | NHP2 non-histone chromosome protein 2-like 1 (S. cerevisiae) |
| 1426934 | at | -1.72 | 0.001 | Nhsl1 | NHS-like 1 |
| 1454942 | at | 1.28 | 0.006 | Niban | niban protein |
| 1449376 | at | 1.01 | 0.774 | Nicn1 | nicolin 1 |
| 1423516 | a_at | 1.22 | 0.011 | Nid2 | nidogen 2 |
| 1438728 | at | -1.07 | 0.327 | Nif3l1 | Ngg1 interacting factor 3-like 1 (S. pombe) |
| 1452579 | at | -1.06 | 0.071 | Nifun | NifU-like N-terminal domain containing |
| 1434864 | at | 1.12 | 0.065 | Nipa1 | non imprinted in Prader-Willi/Angelman syndrome 1 homolog (human) |
| 1437158 | at | -1.14 | 0.186 | Nipbl | Nipped-B homolog (Drosophila) |
| 1451338 | at | 1.08 | 0.018 | Nisch | nischarin |
| 1418763 | at | -1.26 | 0.059 | Nit2 | nitrilase family, member 2 |
| 1428503 | a_at | 1.08 | 0.026 | Nkiras1 | NFKB inhibitor interacting Ras-like protein 1 |
| 1424416 | at | 1.09 | 0.165 | Nkiras2 | NFKB inhibitor interacting Ras-like protein 2 |
| 1455143 | at | 1.25 | 0.050 | Nlgn2 | neuroligin 2 |

Supplementary table S1

| | | | | | |
|---|---|---|---|---|---|
| 1419112 | at | -1.00 | 0.991 | Nlk | nemo like kinase |
| 1451555 | at | 1.31 | 0.002 | Nln | neurolysin (metallopeptidase M3 family) |
| 1419405 | at | -3.16 | 0.000 | Nmb | neuromedin B |
| 1448133 | at | -1.20 | 0.590 | Nmd3 | NMD3 homolog (S. cerevisiae) |
| 1424110 | a_at | 1.25 | 0.013 | Nme1 | expressed in non-metastatic cells 1, protein |
| 1448808 | a_at | 1.03 | 0.270 | Nme2 | expressed in non-metastatic cells 2, protein |
| 1448905 | at | -1.04 | 0.498 | Nme3 | expressed in non-metastatic cells 3 |
| 1422727 | at | -1.16 | 0.154 | Nme5 | expressed in non-metastatic cells 5 |
| 1448574 | at | -1.10 | 0.115 | Nme6 | expressed in non-metastatic cells 6, protein |
| 1418217 | at | 1.02 | 0.736 | Nme7 | non-metastatic cells 7, protein expressed in |
| 1425719 | a_at | -1.11 | 0.120 | Nmi | N-myc (and STAT) interactor |
| 1429819 | at | -1.27 | 0.000 | Nmnat1 | nicotinamide nucleotide adenylyltransferase 1 |
| 1424899 | at | -1.17 | 0.142 | Nmnat3 | nicotinamide nucleotide adenylyltransferase 3 |
| 1417155 | at | 1.15 | 0.598 | Nmyc1 | neuroblastoma myc-related oncogene 1 |
| 1427720 | a_at | -1.07 | 0.282 | Nnp1 | novel nuclear protein 1 |
| 1416105 | at | -1.07 | 0.024 | Nnt | nicotinamide nucleotide transhydrogenase |
| 1448677 | at | 1.08 | 0.381 | Noc4 | neighbor of Cox4 |
| 1424019 | at | 1.16 | 0.013 | Nol1 | nucleolar protein 1 |
| 1451503 | at | -1.08 | 0.278 | Nol3 | nucleolar protein 3 (apoptosis repressor with CARD domain) |
| 1450986 | at | 1.13 | 0.033 | Nol5 | nucleolar protein 5 |
| 1426533 | at | 1.06 | 0.494 | Nol5a | nucleolar protein 5A |
| 1424472 | at | -1.00 | 0.997 | Nol6 | nucleolar protein family 6 (RNA-associated) |
| 1420487 | at | -1.05 | 0.348 | Nol7 | nucleolar protein 7 |
| 1428951 | at | 1.17 | 0.082 | Nol8 | nucleolar protein 8 |
| 1423210 | a_at | 1.01 | 0.781 | Nola3 | nucleolar protein family A, member 3 |
| 1451169 | at | 1.13 | 0.062 | Nomo1 | nodal modulator 1 |
| 1422622 | at | 1.22 | 0.040 | Nos3 | nitric oxide synthase 3, endothelial cell |
| 1418634 | at | -1.31 | 0.058 | Notch1 | Notch gene homolog 1 (Drosophila) |
| 1455556 | at | -1.25 | 0.015 | Notch2 | Notch gene homolog 2 (Drosophila) |
| 1421964 | at | -1.28 | 0.001 | Notch3 | Notch gene homolog 3 (Drosophila) |
| 1449146 | at | 1.34 | 0.026 | Notch4 | Notch gene homolog 4 (Drosophila) |
| 1416057 | at | -1.12 | 0.001 | Np15 | nuclear protein 15.6 |
| 1421037 | at | 1.07 | 0.709 | Npas2 | neuronal PAS domain protein 2 |
| 1423086 | at | -1.46 | 0.000 | Npc1 | Niemann Pick type C1 |
| 1416901 | at | 1.25 | 0.002 | Npc2 | Niemann Pick type C2 |
| 1415919 | at | 1.40 | 0.001 | Npdc1 | neural proliferation, differentiation and control gene 1 |
| 1434507 | at | -1.00 | 0.954 | Npepl1 | aminopeptidase-like 1 |
| 1417385 | at | -1.31 | 0.002 | Npepps | aminopeptidase puromycin sensitive |
| 1432416 | a_at | 1.01 | 0.811 | Npm1 | nucleophosmin 1 |
| 1423522 | at | -1.05 | 0.686 | Npm3 | nucleoplasmin 3 |
| 1452382 | at | 1.15 | 0.239 | Npn1 | neoplastic progression 1 |

Supplementary table S1

| | | | | | |
|---|---|---|---|---|---|
| 1451680 | at | 1.50 | 0.001 | Npn3 | neoplastic progression 3 |
| 1456062 | at | 3.04 | 0.008 | Nppa | natriuretic peptide precursor type A |
| 1450791 | at | 2.05 | 0.018 | Nppb | natriuretic peptide precursor type B |
| 1427191 | at | -1.08 | 0.217 | Npr2 | natriuretic peptide receptor 2 |
| 1435184 | at | 1.49 | 0.000 | Npr3 | natriuretic peptide receptor 3 |
| 1429290 | at | -1.17 | 0.264 | Nptxr | neuronal pentraxin receptor |
| 1423627 | at | 1.20 | 0.062 | Nqo1 | NAD(P)H dehydrogenase, quinone 1 |
| 1436189 | at | -1.36 | 0.002 | Nqo2 | NAD(P)H dehydrogenase, quinone 2 |
| 1424048 | a_at | 1.09 | 0.303 | Nqo3a2 | NAD(P)H:quinone oxidoreductase type 3, polypeptide A2 |
| 1449854 | at | -1.37 | 0.004 | Nr0b2 | nuclear receptor subfamily 0, group B, member 2 |
| 1426464 | at | -1.20 | 0.327 | Nr1d1 | nuclear receptor subfamily 1, group D, member 1 |
| 1416958 | at | -1.26 | 0.048 | Nr1d2 | nuclear receptor subfamily 1, group D, member 2 |
| 1416353 | at | -1.01 | 0.906 | Nr1h2 | nuclear receptor subfamily 1, group H, member 2 |
| 1439696 | at | -1.17 | 0.147 | Nr2c2 | nuclear receptor subfamily 2, group C, member 2 |
| 1418157 | at | -1.03 | 0.748 | Nr2f1 | nuclear receptor subfamily 2, group F, member 1 |
| 1416158 | at | -1.24 | 0.012 | Nr2f2 | nuclear receptor subfamily 2, group F, member 2 |
| 1460648 | at | -1.13 | 0.306 | Nr2f6 | nuclear receptor subfamily 2, group F, member 6 |
| 1435991 | at | -1.68 | 0.010 | Nr3c2 | nuclear receptor subfamily 3, group C, member 2 |
| 1416505 | at | 1.45 | 0.212 | Nr4a1 | nuclear receptor subfamily 4, group A, member 1 |
| 1438796 | at | 1.80 | 0.004 | Nr4a3 | nuclear receptor subfamily 4, group A, member 3 |
| 1421253 | at | 1.40 | 0.004 | Nrap | nebulin-related anchoring protein |
| 1417985 | at | 1.27 | 0.012 | Nrarp | Notch-regulated ankyrin repeat protein |
| 1422687 | at | 1.08 | 0.333 | Nras | neuroblastoma ras oncogene |
| 1457760 | at | -1.03 | 0.892 | Nrbf1 | nuclear receptor binding factor 1 |
| 1448758 | at | 1.11 | 0.139 | Nrbf2 | nuclear receptor binding factor 2 |
| 1423864 | at | -1.02 | 0.756 | Nrbp | nuclear receptor binding protein |
| 1424544 | at | -1.10 | 0.292 | Nrbp2 | nuclear receptor binding protein 2 |
| 1434627 | at | -1.13 | 0.096 | Nrf1 | nuclear respiratory factor 1 |
| 1418469 | at | -1.06 | 0.316 | Nrip1 | nuclear receptor interacting protein 1 |
| 1428393 | at | 1.30 | 0.087 | Nrn1 | neuritin 1 |
| 1435349 | at | 1.20 | 0.287 | Nrp2 | neuropilin 2 |
| 1418152 | at | -1.11 | 0.265 | Nsbp1 | nucleosome binding protein 1 |
| 1418624 | at | -1.14 | 0.000 | Nsep1 | nuclease sensitive element binding protein 1 |
| 1422456 | at | -1.10 | 0.039 | Nsf | N-ethylmaleimide sensitive fusion protein |
| 1429328 | at | 1.01 | 0.860 | Nsfl1c | NSFL1 (p97) cofactor (p47) |
| 1423055 | at | 1.10 | 0.444 | Nsg1 | neuron specific gene family member 1 |
| 1416412 | at | -1.24 | 0.020 | Nsmaf | neutral sphingomyelinase (N-SMase) activation associated factor |
| 1436121 | a_at | -1.15 | 0.001 | Nsmce1 | non-SMC element 1 homolog (S. cerevisiae) |
| 1423850 | at | 1.28 | 0.000 | Nsun2 | NOL1/NOP2/Sun domain family 2 |
| 1436735 | at | 1.25 | 0.122 | Nsun3 | NOL1/NOP2/Sun domain family 3 |
| 1417252 | at | 1.34 | 0.009 | Nt5c | 5',3'-nucleotidase, cytosolic |

Supplementary table S1

| | | | | | |
|---|---|---|---|---|---|
| 1425933 | a_at | 1.34 | 0.003 | Nt5c2 | 5'-nucleotidase, cytosolic II |
| 1460615 | at | -1.08 | 0.571 | Nt5c2l1 | 5'-nucleotidase, cytosolic II-like 1 |
| 1451050 | at | -1.20 | 0.001 | Nt5c3 | 5'-nucleotidase, cytosolic III |
| 1428547 | at | -1.00 | 0.978 | Nt5e | 5' nucleotidase, ecto |
| 1453767 | a_at | -1.01 | 0.874 | Nt5m | 5',3'-nucleotidase, mitochondrial |
| 1423538 | at | -1.01 | 0.756 | Ntan1 | N-terminal Asn amidase |
| 1416517 | at | 1.00 | 0.996 | Nte | neuropathy target esterase |
| 1454974 | at | -1.18 | 0.281 | Ntn1 | netrin 1 |
| 1447723 | at | -1.17 | 0.519 | Ntrk3 | neurotrophic tyrosine kinase, receptor, type 3 |
| 1422860 | at | 1.57 | 0.000 | Nts | neurotensin |
| 1416512 | at | 1.09 | 0.045 | Nubp2 | nucleotide binding protein 2 |
| 1429278 | at | 1.02 | 0.735 | Nubpl | nucleotide binding protein-like |
| 1416903 | at | -1.07 | 0.272 | Nucb1 | nucleobindin 1 |
| 1418355 | at | 1.19 | 0.113 | Nucb2 | nucleobindin 2 |
| 1419471 | a_at | 1.03 | 0.671 | Nudc | nuclear distribution gene C homolog (Aspergillus) |
| 1449652 | at | 1.24 | 0.167 | Nudcd1 | NudC domain containing 1 |
| 1449140 | at | 1.29 | 0.003 | Nudcd2 | NudC domain containing 2 |
| 1460395 | at | -1.15 | 0.075 | Nudcd3 | NudC domain containing 3 |
| 1449349 | at | -1.06 | 0.423 | Nudt1 | nudix (nucleoside diphosphate linked moiety X)-type motif 1 |
| 1426887 | at | -1.31 | 0.131 | Nudt11 | nudix (nucleoside diphosphate linked moiety X)-type motif 11 |
| 1453139 | at | -1.10 | 0.193 | Nudt12 | nudix (nucleoside diphosphate linked moiety X)-type motif 12 |
| 1428943 | at | 1.17 | 0.031 | Nudt13 | nudix (nucleoside diphosphate linked moiety X)-type motif 13 |
| 1424753 | at | -1.16 | 0.152 | Nudt14 | nudix (nucleoside diphosphate linked moiety X)-type motif 14 |
| 1439884 | at | 1.51 | 0.000 | Nudt16 | nudix (nucleoside diphosphate linked moiety X)-type motif 16 |
| 1418737 | at | -1.09 | 0.171 | Nudt2 | nudix (nucleoside diphosphate linked moiety X)-type motif 2 |
| 1451575 | a_at | -1.12 | 0.135 | Nudt3 | nudix (nucleotide diphosphate linked moiety X)-type motif 3 |
| 1449107 | at | -1.06 | 0.429 | Nudt4 | nudix (nucleoside diphosphate linked moiety X)-type motif 4 |
| 1424510 | at | -1.28 | 0.010 | Nudt6 | nudix (nucleoside diphosphate linked moiety X)-type motif 6 |
| 1450111 | a_at | -1.16 | 0.016 | Nudt8 | nudix (nucleoside diphosphate linked moiety X)-type motif 8 |
| 1428164 | at | 1.00 | 0.971 | Nudt9 | nudix (nucleoside diphosphate linked moiety X)-type motif 9 |
| 1448802 | at | 1.11 | 0.060 | Nufip1 | nuclear fragile X mental retardation protein interacting protein |
| 1423974 | at | -1.30 | 0.023 | Numa1 | nuclear mitotic apparatus protein 1 |
| 1416891 | at | 1.09 | 0.294 | Numb | numb gene homolog (Drosophila) |
| 1423787 | at | -1.00 | 0.958 | Nup133 | nucleoporin 133 |
| 1452176 | at | -1.00 | 0.972 | Nup153 | nucleoporin 153 |
| 1449200 | at | 1.10 | 0.357 | Nup155 | nucleoporin 155 |
| 1418530 | at | 1.19 | 0.034 | Nup160 | nucleoporin 160 |
| 1427047 | at | 1.06 | 0.497 | Nup188 | nucleoporin 188 |
| 1426447 | at | 1.09 | 0.163 | Nup35 | nucleoporin 35 |
| 1423969 | at | -1.05 | 0.425 | Nup37 | nucleoporin 37 |
| 1433580 | at | -1.00 | 0.980 | Nup54 | nucleoporin 54 |

Supplementary table S1

| | | | | | |
|---|---|---|---|---|---|
| 1415926 | at | -1.12 | 0.183 | Nup62 | nucleoporin 62 |
| 1428073 | a_at | -1.12 | 0.092 | Nup88 | nucleoporin 88 |
| 1424291 | at | 1.24 | 0.062 | Nup93 | nucleoporin 93 |
| 1424390 | at | 1.21 | 0.494 | Nupl1 | nucleoporin like 1 |
| 1419665 | a_at | 2.57 | 0.000 | Nupr1 | nuclear protein 1 |
| 1418274 | at | 1.00 | 0.955 | Nutf2 | nuclear transport factor 2 |
| 1453019 | at | 1.06 | 0.529 | Nvl | nuclear VCP-like |
| 1416791 | a_at | 1.03 | 0.481 | Nxf1 | nuclear RNA export factor 1 homolog (S. cerevisiae) |
| 1456160 | at | -1.29 | 0.109 | Nxn | nucleoredoxin |
| 1422488 | at | -1.07 | 0.218 | Nxt1 | NTF2-related export protein 1 |
| 1416452 | at | 1.01 | 0.721 | Oat | ornithine aminotransferase |
| 1426764 | at | -1.07 | 0.337 | Oaz2 | ornithine decarboxylase antizyme 2 |
| 1422702 | at | 1.56 | 0.001 | Oazin | ornithine decarboxylase antizyme inhibitor |
| 1434737 | at | 1.19 | 0.131 | Obfc1 | oligonucleotide/oligosaccharide-binding fold containing 1 |
| 1443632 | at | -1.15 | 0.084 | Obscn | obscurin, cytoskeletal calmodulin and titin-interacting RhoGEF |
| 1449889 | a_at | -1.04 | 0.269 | Ociad1 | OCIA domain containing 1 |
| 1427364 | a_at | 1.27 | 0.005 | Odc1 | ornithine decarboxylase, structural 1 |
| 1454735 | at | -1.11 | 0.035 | Odf2 | outer dense fiber of sperm tails 2 |
| 1427172 | at | 1.13 | 0.125 | Ofd1 | oral-facial-digital syndrome 1 gene homolog (human) |
| 1422511 | a_at | -1.01 | 0.912 | Ogfr | opioid growth factor receptor |
| 1424412 | at | 1.10 | 0.460 | Ogfrl1 | opioid growth factor receptor-like 1 |
| 1419662 | at | -1.19 | 0.136 | Ogn | osteoglycin |
| 1448475 | at | 2.13 | 0.000 | Olfml3 | olfactomedin-like 3 |
| 1458543 | at | 1.04 | 0.860 | Olfr619 | olfactory receptor 619 |
| 1424299 | at | -1.31 | 0.000 | Oma1 | OMA1 homolog, zinc metallopeptidase (S. cerevisiae) |
| 1419108 | at | 1.33 | 0.000 | Ophn1 | oligophrenin 1 |
| 1424359 | at | -1.38 | 0.006 | Oplah | 5-oxoprolinase (ATP-hydrolysing) |
| 1429778 | at | 1.04 | 0.458 | Optn | optineurin |
| 1424319 | at | 1.14 | 0.067 | Oraov1 | oral cancer overexpressed 1 |
| 1418225 | at | -1.00 | 0.973 | Orc2l | origin recognition complex, subunit 2-like (S. cerevisiae) |
| 1423337 | at | -1.07 | 0.365 | Orc4l | origin recognition complex, subunit 4-like (S. cerevisiae) |
| 1415830 | at | 1.00 | 0.995 | Orc5l | origin recognition complex, subunit 5-like (S. cerevisiae) |
| 1417037 | at | -1.05 | 0.235 | Orc6l | origin recognition complex, subunit 6-like (S. cerevisiae) |
| 1424597 | at | -1.02 | 0.803 | ORF19 | open reading frame 19 |
| 1420543 | at | -1.33 | 0.000 | ORF28 | open reading frame 28 |
| 1419760 | a_at | 1.28 | 0.001 | ORF5 | open reading frame 5 |
| 1427109 | at | -1.44 | 0.015 | ORF61 | open reading frame 61 |
| 1451219 | at | -1.15 | 0.146 | Ormdl1 | ORM1-like 1 (S. cerevisiae) |
| 1424235 | at | 1.04 | 0.356 | Ormdl2 | ORM1-like 2 (S. cerevisiae) |
| 1419450 | at | 1.14 | 0.228 | Ormdl3 | ORM1-like 3 (S. cerevisiae) |
| 1460350 | at | 1.08 | 0.048 | Osbp | oxysterol binding protein |

Supplementary table S1

| | | | | | |
|---|---|---|---|---|---|
| 1428355 | at | 1.04 | 0.779 | Osbp2 | oxysterol binding protein 2 |
| 1436027 | at | -1.23 | 0.000 | Osbpl11 | oxysterol binding protein-like 11 |
| 1460192 | at | -1.17 | 0.099 | Osbpl1a | oxysterol binding protein-like 1A |
| 1451974 | at | -1.14 | 0.113 | Osbpl2 | oxysterol binding protein-like 2 |
| 1428484 | at | -1.26 | 0.263 | Osbpl3 | oxysterol binding protein-like 3 |
| 1457881 | at | -1.24 | 0.147 | Osbpl6 | oxysterol binding protein-like 6 |
| 1450964 | a_at | 1.22 | 0.017 | Osbpl9 | oxysterol binding protein-like 9 |
| 1418529 | at | -1.11 | 0.017 | Osgep | O-sialoglycoprotein endopeptidase |
| 1428557 | a_at | -1.18 | 0.080 | Osgepl1 | O-sialoglycoprotein endopeptidase-like 1 |
| 1418674 | at | 1.47 | 0.002 | Osmr | oncostatin M receptor |
| 1452246 | at | -1.02 | 0.687 | Ostf1 | osteoclast stimulating factor 1 |
| 1428334 | at | 1.15 | 0.033 | Ostm1 | osteopetrosis associated transmembrane protein 1 |
| 1427346 | at | 1.05 | 0.834 | Ott | ovary testis transcribed |
| 1448817 | at | -1.07 | 0.300 | Otub1 | OTU domain, ubiquitin aldehyde binding 1 |
| 1451088 | a_at | -1.22 | 0.008 | Oxa1l | oxidase assembly 1-like |
| 1428140 | at | 1.02 | 0.149 | Oxct1 | 3-oxoacid CoA transferase 1 |
| 1418502 | a_at | -1.14 | 0.221 | Oxr1 | oxidation resistance 1 |
| 1426933 | at | 1.03 | 0.383 | Oxsr1 | oxidative-stress responsive 1 |
| 1425525 | a_at | 1.18 | 0.038 | P2rx4 | purinergic receptor P2X, ligand-gated ion channel 4 |
| 1449433 | at | 1.41 | 0.021 | P2rx5 | purinergic receptor P2X, ligand-gated ion channel, 5 |
| 1424733 | at | -1.03 | 0.728 | P2ry14 | purinergic receptor P2Y, G-protein coupled, 14 |
| 1428615 | at | -1.10 | 0.101 | P2y5 | purinergic receptor (family A group 5) |
| 1452094 | at | 1.37 | 0.035 | P4ha1 | procollagen-proline, 2-oxoglutarate 4-dioxygenase (proline 4-hydroxylase), alpha 1 polypeptide |
| 1417149 | at | -1.00 | 0.997 | P4ha2 | procollagen-proline, 2-oxoglutarate 4-dioxygenase (proline 4-hydroxylase), alpha II polypeptide |
| 1437465 | a_at | 1.12 | 0.010 | P4hb | prolyl 4-hydroxylase, beta polypeptide |
| 1450854 | at | 1.02 | 0.853 | Pa2g4 | proliferation-associated 2G4 |
| 1418883 | a_at | 1.18 | 0.004 | Pabpc1 | poly A binding protein, cytoplasmic 1 |
| 1421046 | a_at | 1.01 | 0.772 | Pabpc4 | poly A binding protein, cytoplasmic 4 |
| 1458110 | at | -1.20 | 0.014 | Pacs1 | phosphofurin acidic cluster sorting protein 1 |
| 1434255 | at | 1.03 | 0.852 | Pacs1l | phosphofurin acidic cluster sorting protein 1-like |
| 1417810 | a_at | -1.01 | 0.810 | Pacsin2 | protein kinase C and casein kinase substrate in neurons 2 |
| 1416995 | at | 1.12 | 0.073 | Pacsin3 | protein kinase C and casein kinase substrate in neurons 3 |
| 1419058 | at | 1.06 | 0.353 | Paf53 | RNA polymerase I associated factor |
| 1450755 | at | 1.05 | 0.691 | Pafah1b2 | platelet-activating factor acetylhydrolase, isoform 1b, alpha2 subunit |
| 1448489 | at | 1.04 | 0.460 | Pafah2 | platelet-activating factor acetylhydrolase 2 |
| 1436570 | at | 1.14 | 0.100 | Pag1 | phosphoprotein associated with glycosphingolipid microdomains 1 |
| 1425521 | at | -1.13 | 0.033 | Paip1 | polyadenylate binding protein-interacting protein 1 |
| 1448493 | at | 1.03 | 0.449 | Paip2 | polyadenylate-binding protein-interacting protein 2 |
| 1434250 | at | -1.07 | 0.389 | Pak2 | p21 (CDKN1A)-activated kinase 2 |
| 1423967 | at | -1.09 | 0.387 | Palm | paralemmin |
| 1459377 | at | -1.31 | 0.112 | Palm2 | paralemmin 2 |

Supplementary table S1

| | | | | | |
|---|---|---|---|---|---|
| 1417251 | at | 1.03 | 0.679 | Palmd | palmdelphin |
| 1418908 | at | 1.38 | 0.001 | Pam | peptidylglycine alpha-amidating monooxygenase |
| 1441765 | at | -1.46 | 0.086 | Pank1 | pantothenate kinase 1 |
| 1431948 | a_at | 1.06 | 0.371 | Pank2 | pantothenate kinase 2 (Hallervorden-Spatz syndrome) |
| 1433613 | at | -1.03 | 0.579 | Pank3 | pantothenate kinase 3 |
| 1435004 | at | -1.37 | 0.003 | Pank4 | pantothenate kinase 4 |
| 1452749 | at | 1.01 | 0.882 | Papd1 | PAP associated domain containing 1 |
| 1433568 | at | 1.03 | 0.605 | Papd4 | PAP associated domain containing 4 |
| 1427241 | at | -1.04 | 0.648 | Papolg | poly(A) polymerase gamma |
| 1415948 | at | -1.35 | 0.003 | Pappa2 | pappalysin 2 |
| 1415890 | at | 1.15 | 0.065 | Papss1 | 3'-phosphoadenosine 5'-phosphosulfate synthase 1 |
| 1434510 | at | 1.04 | 0.429 | Papss2 | 3'-phosphoadenosine 5'-phosphosulfate synthase 2 |
| 1429216 | at | 1.12 | 0.071 | Paqr3 | progestin and adipoQ receptor family member III |
| 1423101 | at | -1.05 | 0.630 | Paqr4 | progestin and adipoQ receptor family member IV |
| 1436169 | at | -1.37 | 0.068 | Paqr9 | progestin and adipoQ receptor family member IX |
| 1436764 | at | -1.16 | 0.157 | Pard3 | par-3 (partitioning defective 3) homolog (C. elegans) |
| 1420851 | at | 1.05 | 0.600 | Pard6g | par-6 partitioning defective 6 homolog gamma (C. elegans) |
| 1448725 | at | 1.05 | 0.717 | Parg | poly (ADP-ribose) glycohydrolase |
| 1449975 | a_at | -1.03 | 0.738 | Park2 | parkin |
| 1422502 | at | -1.21 | 0.087 | Parp1 | poly (ADP-ribose) polymerase family, member 1 |
| 1434139 | at | 1.08 | 0.211 | Parp11 | poly (ADP-ribose) polymerase family, member 11 |
| 1451564 | at | 1.17 | 0.216 | Parp14 | poly (ADP-ribose) polymerase family, member 14 |
| 1417800 | at | -1.14 | 0.029 | Parp2 | poly (ADP-ribose) polymerase family, member 2 |
| 1428931 | a_at | -1.08 | 0.221 | Parp6 | poly (ADP-ribose) polymerase family, member 6 |
| 1440721 | at | -1.11 | 0.268 | Parp8 | poly (ADP-ribose) polymerase family, member 8 |
| 1416897 | at | 1.37 | 0.007 | Parp9 | poly (ADP-ribose) polymerase family, member 9 |
| 1438672 | at | 1.01 | 0.906 | Parvb | parvin, beta |
| 1426910 | at | -1.24 | 0.043 | Pawr | PRKC, apoptosis, WT1, regulator |
| 1423303 | at | -1.14 | 0.018 | Paxip1 | PAX interacting (with transcription-activation domain) protein 1 |
| 1455320 | at | -1.33 | 0.003 | Pbef1 | pre-B-cell colony-enhancing factor 1 |
| 1415950 | a_at | 1.10 | 0.001 | Pbp | phosphatidylethanolamine binding protein |
| 1428647 | at | -0.87 | 0.059 | Pbx1 | pre B-cell leukemia transcription factor 1 |
| 1418893 | at | -1.02 | 0.676 | Pbx2 | pre B-cell leukemia transcription factor 2 |
| 1442270 | at | -1.03 | 0.725 | Pbx3 | pre B-cell leukemia transcription factor 3 |
| 1451132 | at | 1.20 | 0.007 | Pbxip1 | pre-B-cell leukemia transcription factor interacting protein 1 |
| 1452621 | at | -1.08 | 0.049 | Pcbd2 | pterin 4 alpha carbinolamine dehydratase/dimerization cofactor of hepatocyte nuclear factor 1 alpha (TCF1) 2 |
| 1449054 | a_at | 1.28 | 0.256 | Pcbp4 | poly(rC) binding protein 4 |
| 1451405 | at | -1.23 | 0.004 | Pcca | propionyl-Coenzyme A carboxylase, alpha polypeptide |
| 1450969 | at | -1.25 | 0.000 | Pccb | propionyl Coenzyme A carboxylase, beta polypeptide |
| 1437928 | at | -1.03 | 0.752 | Pcdh12 | protocadherin 12 |
| 1422889 | at | 1.11 | 0.302 | Pcdh18 | protocadherin 18 |

Supplementary table S1

| | | | | | |
|---|---|---|---|---|---|
| 1449583 | at | -1.01 | 0.843 | Pcdhb20 | protocadherin beta 20 |
| 1458721 | at | -1.46 | 0.027 | Pcdhgb1 | protocadherin gamma subfamily B, 1 |
| 1427956 | at | 1.00 | 0.944 | Pcgf1 | polycomb group ring finger 1 |
| 1428423 | at | 1.08 | 0.227 | Pcgf3 | polycomb group ring finger 3 |
| 1425615 | a_at | 1.04 | 0.595 | Pck2 | phosphoenolpyruvate carboxykinase 2 (mitochondrial) |
| 1417947 | at | -1.03 | 0.356 | Pcna | proliferating cell nuclear antigen |
| 1454749 | at | -1.58 | 0.001 | Pcnt2 | pericentrin 2 |
| 1419048 | at | 1.17 | 0.013 | Pcnx | pecanex homolog (Drosophila) |
| 1437165 | a_at | 1.82 | 0.000 | Pcolce | procollagen C-proteinase enhancer protein |
| 1451527 | at | 1.67 | 0.000 | Pcolce2 | procollagen C-endopeptidase enhancer 2 |
| 1451098 | at | 1.16 | 0.003 | Pcoln3 | procollagen (type III) N-endopeptidase |
| 1452913 | at | -1.30 | 0.002 | Pcp4l1 | Purkinje cell protein 4-like 1 |
| 1448435 | at | 1.02 | 0.772 | Pcqap | positive cofactor 2, multiprotein complex, glutamine/Q-rich-associated protein |
| 1426981 | at | -1.13 | 0.001 | Pcsk6 | proprotein convertase subtilisin/kexin type 6 |
| 1460169 | a_at | -1.07 | 0.209 | Pctk1 | PCTAIRE-motif protein kinase 1 |
| 1446272 | at | -1.02 | 0.831 | Pctk2 | PCTAIRE-motif protein kinase 2 |
| 1449151 | at | -1.29 | 0.066 | Pctk3 | PCTAIRE-motif protein kinase 3 |
| 1416383 | a_at | -1.37 | 0.000 | Pcx | pyruvate carboxylase |
| 1423394 | at | -1.02 | 0.819 | Pcyox1 | prenylcysteine oxidase 1 |
| 1438011 | at | -1.10 | 0.036 | Pcyt1a | phosphate cytidylyltransferase 1, choline, alpha isoform |
| 1420493 | a_at | -1.31 | 0.005 | Pcyt2 | phosphate cytidylyltransferase 2, ethanolamine |
| 1448527 | at | 1.03 | 0.564 | Pdcd10 | programmed cell death 10 |
| 1428688 | at | 1.05 | 0.446 | Pdcd11 | programmed cell death protein 11 |
| 1423534 | at | -1.08 | 0.045 | Pdcd2 | programmed cell death 2 |
| 1418840 | at | -1.06 | 0.439 | Pdcd4 | programmed cell death 4 |
| 1450638 | at | -1.04 | 0.272 | Pdcd5 | programmed cell death 5 |
| 1416027 | at | 1.12 | 0.002 | Pdcd6 | programmed cell death 6 |
| 1448155 | at | 1.08 | 0.333 | Pdcd6ip | programmed cell death 6 interacting protein |
| 1416377 | at | 1.11 | 0.172 | Pdcd7 | programmed cell death protein 7 |
| 1418127 | a_at | -1.10 | 0.145 | Pdcd8 | programmed cell death 8 |
| 1423599 | a_at | -1.14 | 0.210 | Pdcl | phosducin-like |
| 1424002 | at | 1.18 | 0.036 | Pdcl3 | phosducin-like 3 |
| 1440374 | at | 1.03 | 0.645 | Pde1c | phosphodiesterase 1C |
| 1452202 | at | -1.05 | 0.633 | Pde2a | phosphodiesterase 2A, cGMP-stimulated |
| 1445373 | at | -1.79 | 0.099 | Pde3a | phosphodiesterase 3A, cGMP inhibited |
| 1433694 | at | -2.02 | 0.099 | Pde3b | phosphodiesterase 3B, cGMP-inhibited |
| 1421535 | a_at | -1.42 | 0.002 | Pde4a | phosphodiesterase 4A, cAMP specific |
| 1435280 | at | -1.77 | 0.000 | Pde4d | phosphodiesterase 4D, cAMP specific |
| 1416843 | at | -1.16 | 0.142 | Pde6d | phosphodiesterase 6D, cGMP-specific, rod, delta |
| 1423313 | at | -1.59 | 0.010 | Pde7a | phosphodiesterase 7A |
| 1418406 | at | 1.60 | 0.000 | Pde8a | phosphodiesterase 8A |

Supplementary table S1

| | | | | | |
|---|---|---|---|---|---|
| 1418711 | at | -1.33 | 0.006 | Pdgfa | platelet derived growth factor, alpha |
| 1450414 | at | -1.01 | 0.869 | Pdgfb | platelet derived growth factor, B polypeptide |
| 1419123 | a_at | 1.73 | 0.012 | Pdgfc | platelet-derived growth factor, C polypeptide |
| 1456532 | at | 1.00 | 0.996 | Pdgfd | platelet-derived growth factor, D polypeptide |
| 1421917 | at | 1.13 | 0.029 | Pdgfra | platelet derived growth factor receptor, alpha polypeptide |
| 1418560 | at | -1.13 | 0.043 | Pdha1 | pyruvate dehydrogenase E1 alpha 1 |
| 1456090 | at | 1.07 | 0.310 | Pdhx | pyruvate dehydrogenase complex, component X |
| 1424650 | at | -1.05 | 0.559 | Pdia5 | protein disulfide isomerase-associated 5 |
| 1437175 | at | -1.03 | 0.779 | Pdik1l | PDLIM1 interacting kinase 1 like |
| 1423748 | at | 1.14 | 0.255 | Pdk1 | pyruvate dehydrogenase kinase, isoenzyme 1 |
| 1448825 | at | -1.12 | 0.006 | Pdk2 | pyruvate dehydrogenase kinase, isoenzyme 2 |
| 1417273 | at | 1.04 | 0.474 | Pdk4 | pyruvate dehydrogenase kinase, isoenzyme 4 |
| 1416554 | at | 1.36 | 0.007 | Pdlim1 | PDZ and LIM domain 1 (elfin) |
| 1449178 | at | 1.33 | 0.006 | Pdlim3 | PDZ and LIM domain 3 |
| 1417928 | at | -1.75 | 0.000 | Pdlim4 | PDZ and LIM domain 4 |
| 1442710 | at | 1.24 | 0.081 | Pdlim5 | PDZ and LIM domain 5 |
| 1415729 | at | -1.07 | 0.222 | Pdpk1 | 3-phosphoinositide dependent protein kinase-1 |
| 1428308 | at | -1.04 | 0.085 | Pdrg1 | p53 and DNA damage regulated 1 |
| 1427929 | a_at | -1.06 | 0.261 | Pdxk | pyridoxal (pyridoxine, vitamin B6) kinase |
| 1451269 | at | 1.10 | 0.002 | Pdzk11 | PDZ domain containing 11 |
| 1435553 | at | -1.33 | 0.013 | Pdzk3 | PDZ domain containing 3 |
| 1416846 | a_at | 1.10 | 0.179 | Pdzrn3 | PDZ domain containing RING finger 3 |
| 1449834 | at | -1.36 | 0.001 | Pdzx | PDZ domain containing, X chromosome |
| 1416406 | at | 1.18 | 0.312 | Pea15 | phosphoprotein enriched in astrocytes 15 |
| 1447092 | at | -1.10 | 0.288 | Pecam1 | platelet/endothelial cell adhesion molecule 1 |
| 1431012 | a_at | -1.20 | 0.008 | Peci | peroxisomal delta3, delta2-enoyl-Coenzyme A isomerase |
| 1448910 | at | -1.05 | 0.445 | Pecr | peroxisomal trans-2-enoyl-CoA reductase |
| 1433924 | at | -1.03 | 0.712 | Peg3 | paternally expressed 3 |
| 1417371 | at | -1.05 | 0.301 | Peli1 | pellino 1 |
| 1437181 | at | -1.30 | 0.008 | Peli2 | pellino 2 |
| 1434071 | a_at | 1.07 | 0.523 | Pelo | pelota homolog (Drosophila) |
| 1427038 | at | -3.64 | 0.000 | Penk1 | preproenkephalin 1 |
| 1434168 | at | -1.15 | 0.037 | Peo1 | progressive external ophthalmoplegia 1 (human) |
| 1416712 | at | 1.16 | 0.110 | Pep4 | peptidase 4 |
| 1449851 | at | 1.01 | 0.947 | Per1 | period homolog 1 (Drosophila) |
| 1417602 | at | -1.20 | 0.533 | Per2 | period homolog 2 (Drosophila) |
| 1441445 | at | -1.30 | 0.151 | Per3 | period homolog 3 (Drosophila) |
| 1416271 | at | 1.17 | 0.000 | Perp | PERP, TP53 apoptosis effector |
| 1416172 | at | 1.24 | 0.203 | Pes1 | pescadillo homolog 1, containing BRCT domain (zebrafish) |
| 1424282 | at | -1.16 | 0.185 | Pet112l | PET112-like (yeast) |
| 1428716 | at | -1.32 | 0.009 | Pex1 | peroxisome biogenesis factor 1 |

Supplementary table S1

| | | | | | |
|---|---|---|---|---|---|
| 1419365 | at | -1.35 | 0.029 | Pex11a | peroxisomal biogenesis factor 11a |
| 1430856 | at | -1.18 | 0.089 | Pex11c | peroxisomal biogenesis factor 11c |
| 1416259 | at | 1.09 | 0.155 | Pex12 | peroxisomal biogenesis factor 12 |
| 1422471 | at | -1.03 | 0.529 | Pex13 | peroxisomal biogenesis factor 13 |
| 1425021 | a_at | -1.06 | 0.459 | Pex16 | peroxisome biogenesis factor 16 |
| 1451393 | at | -1.19 | 0.040 | Pex26 | peroxisome biogenesis factor 26 |
| 1417442 | a_at | 1.14 | 0.120 | Pex3 | peroxisomal biogenesis factor 3 |
| 1426770 | at | -1.01 | 0.865 | Pex5 | peroxisome biogenesis factor 5 |
| 1418988 | at | -1.22 | 0.000 | Pex7 | peroxisome biogenesis factor 7 |
| 1452279 | at | -1.31 | 0.012 | Pfc | properdin factor, complement |
| 1417026 | at | 1.04 | 0.405 | Pfdn1 | prefoldin 1 |
| 1421950 | at | -1.30 | 0.000 | Pfdn2 | prefoldin 2 |
| 1454888 | at | 1.00 | 0.964 | Pfdn4 | prefoldin 4 |
| 1415736 | at | -1.09 | 0.010 | Pfdn5 | prefoldin 5 |
| 1427213 | at | -3.29 | 0.000 | Pfkfb1 | 6-phosphofructo-2-kinase/fructose-2,6-biphosphatase 1 |
| 1429486 | at | -1.19 | 0.041 | Pfkfb2 | 6-phosphofructo-2-kinase/fructose-2,6-biphosphatase 2 |
| 1450269 | a_at | -1.00 | 0.974 | Pfkl | phosphofructokinase, liver, B-type |
| 1416780 | at | -1.21 | 0.000 | Pfkm | phosphofructokinase, muscle |
| 1416069 | at | 2.02 | 0.000 | Pfkp | phosphofructokinase, platelet |
| 1449018 | at | -1.05 | 0.613 | Pfn1 | profilin 1 |
| 1418210 | at | 1.16 | 0.045 | Pfn2 | profilin 2 |
| 1445984 | at | -1.14 | 0.387 | Pftk1 | PFTAIRE protein kinase 1 |
| 1426554 | a_at | 1.22 | 0.008 | Pgam1 | phosphoglycerate mutase 1 |
| 1418373 | at | 1.04 | 0.148 | Pgam2 | phosphoglycerate mutase 2 |
| 1416441 | at | 1.10 | 0.026 | Pgcp | plasma glutamate carboxypeptidase |
| 1423706 | a_at | 1.12 | 0.089 | Pgd | phosphogluconate dehydrogenase |
| 1451305 | at | -1.32 | 0.013 | Pgea1 | PKD2 interactor, golgi and endoplasmic reticulum associated 1 |
| 1418471 | at | -1.45 | 0.021 | Pgf | placental growth factor |
| 1438229 | at | 1.06 | 0.385 | Pggt1b | protein geranylgeranyltransferase type I, beta subunit |
| 1417864 | at | 1.14 | 0.000 | Pgk1 | phosphoglycerate kinase 1 |
| 1417722 | at | 1.10 | 0.335 | Pgls | 6-phosphogluconolactonase |
| 1437871 | at | -1.06 | 0.193 | Pgm5 | phosphoglucomutase 5 |
| 1460001 | at | -1.25 | 0.640 | Pgpep1 | pyroglutamyl-peptidase I |
| 1423451 | at | 1.16 | 0.081 | Pgrmc1 | progesterone receptor membrane component 1 |
| 1452882 | at | 1.20 | 0.035 | Pgrmc2 | progesterone receptor membrane component 2 |
| 1454832 | at | -1.03 | 0.789 | Phactr1 | phosphatase and actin regulator 1 |
| 1450778 | a_at | 1.05 | 0.293 | Phax | phosphorylated adaptor for RNA export |
| 1417053 | at | -1.14 | 0.240 | Phb | prohibitin |
| 1423212 | at | 1.10 | 0.083 | Phc1 | polyhomeotic-like 1 (Drosophila) |
| 1416048 | at | 1.08 | 0.380 | Phc2 | polyhomeotic-like 2 (Drosophila) |
| 1455312 | at | -1.05 | 0.263 | Phc3 | polyhomeotic-like 3 (Drosophila) |

Supplementary table S1

| | | | | | |
|---|---|---|---|---|---|
| 1438435 | at | -1.14 | 0.140 | Phca | phytoceramidase, alkaline |
| 1449085 | at | 1.10 | 0.015 | Phf10 | PHD finger protein 10 |
| 1434922 | at | -1.19 | 0.014 | Phf12 | PHD finger protein 12 |
| 1455175 | at | 1.02 | 0.762 | Phf13 | PHD finger protein 13 |
| 1455345 | at | -1.02 | 0.824 | Phf15 | PHD finger protein 15 |
| 1452179 | at | -1.07 | 0.452 | Phf17 | PHD finger protein 17 |
| 1434924 | at | -1.04 | 0.515 | Phf2 | PHD finger protein 2 |
| 1441145 | at | -1.12 | 0.296 | Phf21a | PHD finger protein 21A |
| 1424170 | at | -1.09 | 0.032 | Phf5a | PHD finger protein 5A |
| 1420260 | at | -1.10 | 0.309 | Phf7 | PHD finger protein 7 |
| 1460398 | at | 1.07 | 0.374 | Phf8 | PHD finger protein 8 |
| 1423928 | at | -1.12 | 0.104 | Phgdhl1 | phosphoglycerate dehydrogenase like 1 |
| 1422743 | at | 1.28 | 0.038 | Phka1 | phosphorylase kinase alpha 1 |
| 1434518 | at | -1.08 | 0.191 | Phka2 | phosphorylase kinase alpha 2 |
| 1434511 | at | -1.10 | 0.879 | Phkb | phosphorylase kinase beta |
| 1425164 | a_at | -3.73 | 0.000 | Phkg1 | phosphorylase kinase gamma 1 |
| 1428511 | at | 1.10 | 0.419 | Phkg2 | phosphorylase kinase, gamma 2 (testis) |
| 1418835 | at | 1.53 | 0.006 | Phlda1 | pleckstrin homology-like domain, family A, member 1 |
| 1424467 | at | 1.05 | 0.513 | Phldb1 | pleckstrin homology-like domain, family B, member 1 |
| 1454862 | at | 1.55 | 0.000 | Phldb2 | pleckstrin homology-like domain, family B, member 2 |
| 1459612 | at | -1.19 | 0.131 | Phlpp | PH domain and leucine rich repeat protein phosphatase |
| 1427903 | at | -1.42 | 0.000 | Phpt1 | phosphohistidine phosphatase 1 |
| 1434274 | at | 1.02 | 0.771 | Phr1 | pam, highwire, rpm 1 |
| 1449148 | a_at | 1.34 | 0.002 | Phtf1 | putative homeodomain transcription factor 1 |
| 1437637 | at | -1.06 | 0.429 | Phtf2 | putative homeodomain transcription factor 2 |
| 1422272 | at | 1.07 | 0.163 | Phxr4 | per-hexamer repeat gene 4 |
| 1460194 | at | -1.01 | 0.497 | Phyh | phytanoyl-CoA hydroxylase |
| 1455463 | at | -1.18 | 0.155 | Phyhip | phytanoyl-CoA hydroxylase interacting protein |
| 1453839 | a_at | 2.37 | 0.000 | Pi16 | protease inhibitor 16 |
| 1454605 | a_at | -1.09 | 0.461 | Pi4k2a | phosphatidylinositol 4-kinase type 2 alpha |
| 1416489 | at | 1.30 | 0.010 | Pi4k2b | phosphatidylinositol 4-kinase type 2 beta |
| 1455486 | at | -1.19 | 0.040 | Pias1 | protein inhibitor of activated STAT 1 |
| 1451115 | at | 1.13 | 0.124 | Pias3 | protein inhibitor of activated STAT 3 |
| 1421257 | at | -1.16 | 0.264 | Pigb | phosphatidylinositol glycan, class B |
| 1424189 | at | -1.11 | 0.170 | Pigc | phosphatidylinositol glycan, classC |
| 1448896 | at | 1.10 | 0.196 | Pigf | phosphatidylinositol glycan, class F |
| 1435262 | at | 1.27 | 0.003 | Pign | phosphatidylinositol glycan, class N |
| 1415992 | at | -1.21 | 0.028 | Pigo | phosphatidylinositol glycan, class O |
| 1415946 | at | 1.11 | 0.049 | Pigq | phosphatidylinositol glycan, class Q |
| 1433429 | at | 1.08 | 0.169 | Pigs | phosphatidylinositol glycan, class S |
| 1428740 | a_at | 1.18 | 0.024 | Pigt | phosphatidylinositol glycan, class T |

Supplementary table S1

| | | | | | |
|---|---|---|---|---|---|
| 1455075 | at | -1.26 | 0.007 | Pigv | phosphatidylinositol glycan, class V |
| 1425134 | a_at | -1.19 | 0.005 | Pigx | phosphatidylinositol glycan, class X |
| 1421023 | at | -1.05 | 0.323 | Pik3c2a | phosphatidylinositol 3-kinase, C2 domain containing, alpha polypeptide |
| 1425580 | a_at | -1.03 | 0.630 | Pik3c3 | phosphoinositide-3-kinase, class 3 |
| 1453069 | at | 1.11 | 0.064 | Pik3cb | phosphatidylinositol 3-kinase, catalytic, beta polypeptide |
| 1425514 | at | -1.47 | 0.814 | Pik3r1 | phosphatidylinositol 3-kinase, regulatory subunit, polypeptide 1 (p85 alpha) |
| 1456482 | at | 1.32 | 0.011 | Pik3r3 | phosphatidylinositol 3 kinase, regulatory subunit, polypeptide 3 (p55) |
| 1435003 | at | 1.01 | 0.896 | Pik4ca | phosphatidylinositol 4-kinase, catalytic, alpha polypeptide |
| 1435458 | at | -1.26 | 0.047 | Pim1 | proviral integration site 1 |
| 1451069 | at | -1.80 | 0.003 | Pim3 | proviral integration site 3 |
| 1416228 | at | -1.18 | 0.004 | Pin1 | protein (peptidyl-prolyl cis/trans isomerase) NIMA-interacting 1 |
| 1428658 | at | 1.08 | 0.105 | Pin4 | protein (peptidyl-prolyl cis/trans isomerase) NIMA-interacting, 4 (parvulin) |
| 1451148 | at | -1.22 | 0.000 | Pink1 | PTEN induced putative kinase 1 |
| 1434058 | at | 1.30 | 0.000 | Pip3ap | phosphatidylinositol-3-phosphatase associated protein |
| 1424954 | a_at | 1.10 | 0.275 | Pip5k1c | phosphatidylinositol-4-phosphate 5-kinase, type 1 gamma |
| 1441068 | at | 1.17 | 0.353 | Pip5k2a | phosphatidylinositol-4-phosphate 5-kinase, type II, alpha |
| 1435068 | at | -1.00 | 0.956 | Pip5k2b | phosphatidylinositol-4-phosphate 5-kinase, type II, beta |
| 1416387 | at | -1.08 | 0.010 | Pip5k2c | phosphatidylinositol-4-phosphate 5-kinase, type II, gamma |
| 1452994 | at | 1.11 | 0.098 | Pip5k3 | phosphatidylinositol-3-phosphate/phosphatidylinositol 5-kinase, type III |
| 1429001 | at | 1.18 | 0.002 | Pir | pirin |
| 1440335 | at | -1.32 | 0.012 | Pitpnc1 | phosphatidylinositol transfer protein, cytoplasmic 1 |
| 1425760 | a_at | -1.21 | 0.225 | Pitpnm1 | phosphatidylinositol membrane-associated 1 |
| 1419757 | at | -1.06 | 0.615 | Pitpnm2 | phosphatidylinositol transfer protein, membrane-associated 2 |
| 1426453 | at | -1.18 | 0.000 | Pitrm1 | pitrilysin metalloprotease 1 |
| 1452083 | a_at | -1.15 | 0.109 | Pja1 | praja1, RING-H2 motif containing |
| 1434383 | at | 1.25 | 0.012 | Pja2 | praja 2, RING-H2 motif containing |
| 1460210 | at | 1.24 | 0.011 | Pkd1 | polycystic kidney disease 1 homolog |
| 1417753 | at | 1.29 | 0.000 | Pkd2 | polycystic kidney disease 2 |
| 1419422 | at | -1.20 | 0.058 | Pkd2l2 | polycystic kidney disease 2-like 2 |
| 1420858 | at | -1.05 | 0.485 | Pkia | protein kinase inhibitor, alpha |
| 1423945 | a_at | -1.27 | 0.001 | Pkig | protein kinase inhibitor, gamma |
| 1417308 | at | -1.02 | 0.427 | Pkm2 | pyruvate kinase, muscle |
| 1434525 | at | 1.16 | 0.111 | Pkn3 | protein kinase N3 |
| 1450172 | at | -1.20 | 0.256 | Pknox1 | Pbx/knotted 1 homeobox |
| 1433907 | at | 1.00 | 0.940 | Pknox2 | Pbx/knotted 1 homeobox 2 |
| 1438677 | at | -1.10 | 0.073 | Pkp4 | plakophilin 4 |
| 1417785 | at | 1.48 | 0.005 | Pla1a | phospholipase A1 member A |
| 1452026 | a_at | 1.01 | 0.850 | Pla2g12a | phospholipase A2, group XIIA |
| 1417814 | at | -2.25 | 0.000 | Pla2g5 | phospholipase A2, group V |
| 1435001 | at | 1.23 | 0.087 | Plaa | phospholipase A2, activating protein |
| 1415806 | at | 1.68 | 0.000 | Plat | plasminogen activator, tissue |

Supplementary table S1

| | | | | | |
|---|---|---|---|---|---|
| 1435043 | at | 1.21 | 0.025 | Plcb1 | phospholipase C, beta 1 |
| 1448661 | at | 1.02 | 0.794 | Plcb3 | phospholipase C, beta 3 |
| 1448432 | at | -1.21 | 0.006 | Plcd1 | phospholipase C, delta 1 |
| 1452398 | at | 1.10 | 0.192 | Plce1 | phospholipase C, epsilon 1 |
| 1435149 | at | 1.01 | 0.770 | Plcg1 | phospholipase C, gamma 1 |
| 1426926 | at | 1.67 | 0.000 | Plcg2 | phospholipase C, gamma 2 |
| 1426450 | at | -1.18 | 0.060 | Plcl2 | phospholipase C-like 2 |
| 1416013 | at | 1.01 | 0.891 | Pld3 | phospholipase D3 |
| 1417349 | at | 1.44 | 0.000 | Pldn | pallidin |
| 1437554 | at | 1.04 | 0.833 | Plec1 | plectin 1 |
| 1417965 | at | 1.10 | 0.064 | Plekha1 | pleckstrin homology domain containing, family A (phosphoinositide binding specific) member 1 |
| 1436177 | at | 1.08 | 0.216 | Plekha2 | pleckstrin homology domain-containing, family A (phosphoinositide binding specific) member 2 |
| 1420840 | at | 1.08 | 0.184 | Plekha3 | pleckstrin homology domain-containing, family A (phosphoinositide binding specific) member 3 |
| 1427149 | at | 1.36 | 0.001 | Plekha6 | pleckstrin homology domain containing, family A member 6 |
| 1454819 | at | -1.10 | 0.223 | Plekha8 | pleckstrin homology domain containing, family A (phosphoinositide binding specific) member 8 |
| 1460341 | at | 1.08 | 0.445 | Plekhb2 | pleckstrin homology domain containing, family B (evectins) member 2 |
| 1424671 | at | -2.46 | 0.000 | Plekhf1 | pleckstrin homology domain containing, family F (with FYVE domain) member 1 |
| 1418201 | at | 1.28 | 0.014 | Plekhg2 | pleckstrin homology domain containing, family G (with RhoGef domain) member 2 |
| 1455860 | at | -1.23 | 0.097 | Plekhh1 | pleckstrin homology domain containing, family H (with MyTH4 domain) member 1 |
| 1424179 | at | -1.30 | 0.003 | Plekhj1 | pleckstrin homology domain containing, family J member 1 |
| 1454746 | at | -1.00 | 0.942 | Plekhm1 | pleckstrin homology domain containing, family M (with RUN domain) member 1 |
| 1427005 | at | 1.33 | 0.002 | Plk2 | polo-like kinase 2 (Drosophila) |
| 1423359 | at | -1.19 | 0.174 | Pln | phospholamban |
| 1416289 | at | -1.04 | 0.650 | Plod1 | procollagen-lysine, 2-oxoglutarate 5-dioxygenase 1 |
| 1416687 | at | 1.12 | 0.913 | Plod2 | procollagen lysine, 2-oxoglutarate 5-dioxygenase 2 |
| 1415901 | at | 1.13 | 0.536 | Plod3 | procollagen-lysine, 2-oxoglutarate 5-dioxygenase 3 |
| 1451718 | at | -1.41 | 0.002 | Plp1 | proteolipid protein (myelin) 1 |
| 1439296 | at | 1.12 | 0.473 | Plp2 | proteolipid protein 2 |
| 1448282 | at | -1.12 | 0.012 | Plrg1 | pleiotropic regulator 1, PRL1 homolog (Arabidopsis) |
| 1423725 | at | 1.25 | 0.002 | Pls3 | plastin 3 (T-isoform) |
| 1429527 | a_at | 1.39 | 0.002 | Plscr1 | phospholipid scramblase 1 |
| 1448961 | at | 1.02 | 0.894 | Plscr2 | phospholipid scramblase 2 |
| 1449020 | at | 1.15 | 0.023 | Plscr3 | phospholipid scramblase 3 |
| 1433626 | at | 1.10 | 0.162 | Plscr4 | phospholipid scramblase 4 |
| 1429141 | at | -1.06 | 0.866 | Pltp | phospholipid transfer protein |
| 1424902 | at | -1.19 | 0.046 | Plxdc1 | plexin domain containing 1 |
| 1418912 | at | 1.08 | 0.181 | Plxdc2 | plexin domain containing 2 |
| 1428623 | at | -1.11 | 0.033 | Plxna1 | plexin A1 |
| 1435254 | at | -2.31 | 0.000 | Plxnb1 | plexin B1 |
| 1416683 | at | 1.00 | 0.983 | Plxnb2 | plexin B2 |
| 1451475 | at | -1.03 | 0.801 | Plxnd1 | plexin D1 |

Supplementary table S1

| | | | | | |
|---|---|---|---|---|---|
| 1424167 | a_at | 1.22 | 0.009 | Pmm1 | phosphomannomutase 1 |
| 1431938 | a_at | 1.00 | 0.950 | Pmm2 | phosphomannomutase 2 |
| 1417133 | at | 1.33 | 0.000 | Pmp22 | peripheral myelin protein |
| 1424662 | at | -1.04 | 0.590 | Pmpca | peptidase (mitochondrial processing) alpha |
| 1429709 | at | 1.01 | 0.728 | Pmpcb | peptidase (mitochondrial processing) beta |
| 1427893 | a_at | -1.12 | 0.092 | Pmvk | phosphomevalonate kinase |
| 1422711 | a_at | -1.62 | 0.001 | Pnck | pregnancy upregulated non-ubiquitously expressed CaM kinase |
| 1416378 | at | -1.02 | 0.645 | Pnkp | polynucleotide kinase 3'- phosphatase |
| 1453299 | a_at | 1.35 | 0.003 | Pnp | purine-nucleoside phosphorylase |
| 1427367 | at | -1.18 | 0.401 | Pnpla2 | patatin-like phospholipase domain containing 2 |
| 1415793 | at | 1.03 | 0.485 | Pnpo | pyridoxine 5'-phosphate oxidase |
| 1452676 | a_at | -1.01 | 0.825 | Pnpt1 | polyribonucleotide nucleotidyltransferase 1 |
| 1416186 | at | -1.15 | 0.017 | Pnrc2 | proline-rich nuclear receptor coactivator 2 |
| 1448688 | at | 1.15 | 0.074 | Podxl | podocalyxin-like |
| 1443881 | at | 1.05 | 0.178 | Pofut1 | protein O-fucosyltransferase 1 |
| 1448384 | at | -1.05 | 0.709 | Pofut2 | protein O-fucosyltransferase 2 |
| 1434650 | at | -1.09 | 0.586 | Pogz | pogo transposable element with ZNF domain |
| 1434230 | at | 1.10 | 0.465 | Polb | polymerase (DNA directed), beta |
| 1448277 | at | -1.27 | 0.003 | Pold2 | polymerase (DNA directed), delta 2, regulatory subunit |
| 1426838 | at | 1.05 | 0.442 | Pold3 | polymerase (DNA-directed), delta 3, accessory subunit |
| 1427885 | at | 1.01 | 0.920 | Pold4 | polymerase (DNA-directed), delta 4 |
| 1422732 | at | -1.04 | 0.937 | Poldip2 | polymerase (DNA-directed), delta interacting protein 2 |
| 1452709 | at | 1.00 | 0.975 | Poldip3 | polymerase (DNA-directed), delta interacting protein 3 |
| 1452743 | at | 1.07 | 0.605 | Pole3 | polymerase (DNA directed), epsilon 3 (p17 subunit) |
| 1423371 | at | 1.04 | 0.428 | Pole4 | polymerase (DNA-directed), epsilon 4 (p12 subunit) |
| 1450816 | at | -1.19 | 0.039 | Polg2 | polymerase (DNA directed), gamma 2, accessory subunit |
| 1426242 | at | 1.24 | 0.002 | Polr2a | polymerase (RNA) II (DNA directed) polypeptide A |
| 1433552 | a_at | 1.09 | 0.008 | Polr2b | polymerase (RNA) II (DNA directed) polypeptide B |
| 1416341 | at | -1.12 | 0.053 | Polr2c | polymerase (RNA) II (DNA directed) polypeptide C |
| 1424258 | at | 1.21 | 0.047 | Polr2d | polymerase (RNA) II (DNA directed) polypeptide D |
| 1416270 | at | 1.19 | 0.002 | Polr2g | polymerase (RNA) II (DNA directed) polypeptide G |
| 1424473 | at | 1.00 | 0.942 | Polr2h | polymerase (RNA) II (DNA directed) polypeptide H |
| 1428494 | a_at | -1.15 | 0.025 | Polr2i | polymerase (RNA) II (DNA directed) polypeptide I |
| 1417720 | at | -1.24 | 0.000 | Polr2j | polymerase (RNA) II (DNA directed) polypeptide J |
| 1452596 | at | -1.20 | 0.006 | Polr2k | polymerase (RNA) II (DNA directed) polypeptide K |
| 1428296 | at | 1.16 | 0.108 | Polr2l | polymerase (RNA) II (DNA directed) polypeptide L |
| 1437525 | a_at | 1.19 | 0.016 | Polr3a | polymerase (RNA) III (DNA directed) polypeptide A |
| 1453256 | at | -1.13 | 0.341 | Polr3c | polymerase (RNA) III (DNA directed) polypeptide C |
| 1451364 | at | -1.34 | 0.002 | Polr3gl | polymerase (RNA) III (DNA directed) polypeptide G like |
| 1424227 | at | 1.28 | 0.012 | Polr3h | polymerase (RNA) III (DNA directed) polypeptide H |
| 1422752 | at | 1.06 | 0.452 | Polr3k | polymerase (RNA) III (DNA directed) polypeptide K |

Supplementary table S1

| | | | | | |
|---|---|---|---|---|---|
| 1437377 | a_at | -1.25 | 0.085 | Polrmt | polymerase (RNA) mitochondrial (DNA directed) |
| 1438004 | at | -1.02 | 0.864 | Pols | polymerase (DNA directed) sigma |
| 1450686 | at | 1.19 | 0.036 | Pon2 | paraoxonase 2 |
| 1419298 | at | 1.40 | 0.001 | Pon3 | paraoxonase 3 |
| 1448419 | at | 1.05 | 0.539 | Pop4 | processing of precursor 4, ribonuclease P/MRP family, (S. cerevisiae) |
| 1449162 | at | 1.11 | 0.207 | Pop7 | processing of precursor 7, ribonuclease P family, (S. cerevisiae) |
| 1417806 | at | 1.07 | 0.229 | Popdc2 | popeye domain containing 2 |
| 1423856 | at | 1.25 | 0.022 | Popdc3 | popeye domain containing 3 |
| 1416933 | at | 1.26 | 0.098 | Por | P450 (cytochrome) oxidoreductase |
| 1423606 | at | 5.40 | 0.000 | Postn | periostin, osteoblast specific factor |
| 1448379 | at | -1.10 | 0.199 | Pot1 | protection of telomeres 1 |
| 1427835 | at | -1.21 | 0.218 | Pou2f1 | POU domain, class 2, transcription factor 1 |
| 1452844 | at | -1.02 | 0.755 | Pou6f1 | POU domain, class 6, transcription factor 1 |
| 1449937 | at | 1.12 | 0.267 | Pp11r | placental protein 11 related |
| 1423703 | at | 1.11 | 0.060 | Ppan | peter pan homolog (Drosophila) |
| 1422619 | at | 1.26 | 0.000 | Ppap2a | phosphatidic acid phosphatase 2a |
| 1449051 | at | -1.62 | 0.007 | Ppara | peroxisome proliferator activated receptor alpha |
| 1420715 | a_at | -1.08 | 0.512 | Pparg | peroxisome proliferator activated receptor gamma |
| 1437751 | at | -1.40 | 0.065 | Ppargc1a | peroxisome proliferative activated receptor, gamma, coactivator 1 alpha |
| 1428543 | at | -1.18 | 0.010 | Ppat | phosphoribosyl pyrophosphate amidotransferase |
| 1448128 | at | 1.11 | 0.203 | Ppgb | protective protein for beta-galactosidase |
| 1450911 | at | 1.03 | 0.652 | Ppib | peptidylprolyl isomerase B |
| 1416498 | at | 1.61 | 0.000 | Ppic | peptidylprolyl isomerase C |
| 1417057 | a_at | 1.07 | 0.301 | Ppid | peptidylprolyl isomerase D (cyclophilin D) |
| 1416940 | at | -1.05 | 0.502 | Ppif | peptidylprolyl isomerase F (cyclophilin F) |
| 1424136 | a_at | 1.03 | 0.644 | Ppih | peptidyl prolyl isomerase H |
| 1428892 | at | -1.72 | 0.000 | Ppil1 | peptidylprolyl isomerase (cyclophilin)-like 1 |
| 1452920 | a_at | -1.09 | 0.224 | Ppil2 | peptidylprolyl isomerase (cyclophilin)-like 2 |
| 1429627 | at | -1.02 | 0.433 | Ppil3 | peptidylprolyl isomerase (cyclophilin)-like 3 |
| 1424043 | at | -1.04 | 0.311 | Ppil4 | peptidylprolyl isomerase (cyclophilin)-like 4 |
| 1415678 | at | 1.26 | 0.022 | Ppm1a | protein phosphatase 1A, magnesium dependent, alpha isoform |
| 1451669 | at | 1.31 | 0.000 | Ppm1b | protein phosphatase 1B, magnesium dependent, beta isoform |
| 1434142 | at | -1.05 | 0.453 | Ppm1d | protein phosphatase 1D magnesium-dependent, delta isoform |
| 1434990 | at | 2.18 | 0.000 | Ppm1e | protein phosphatase 1E (PP2C domain containing) |
| 1454934 | at | 1.12 | 0.043 | Ppm1f | protein phosphatase 1F (PP2C domain containing) |
| 1416792 | at | -1.06 | 0.160 | Ppm1g | protein phosphatase 1G (formerly 2C), magnesium-dependent, gamma isoform |
| 1435510 | at | -1.36 | 0.031 | Ppm1h | protein phosphatase 1H (PP2C domain containing) |
| 1453127 | at | 1.09 | 0.392 | Ppm1j | protein phosphatase 1J |
| 1452973 | at | -1.78 | 0.007 | Ppm1k | protein phosphatase 1K (PP2C domain containing) |
| 1435699 | at | -1.57 | 0.009 | Ppm1l | protein phosphatase 1 (formerly 2C)-like |
| 1428351 | at | -1.02 | 0.852 | Ppm1m | protein phosphatase 1M |

Supplementary table S1

| | | | | | |
|---|---|---|---|---|---|
| 1416618 | at | -1.02 | 0.702 | Ppox | protoporphyrinogen oxidase |
| 1460165 | at | -1.07 | 0.112 | Ppp1ca | protein phosphatase 1, catalytic subunit, alpha isoform |
| 1452046 | a_at | 1.05 | 0.333 | Ppp1cc | protein phosphatase 1, catalytic subunit, gamma isoform |
| 1437576 | at | -1.09 | 0.517 | Ppp1r10 | protein phosphatase 1, regulatory subunit 10 |
| 1448565 | at | -1.07 | 0.345 | Ppp1r11 | protein phosphatase 1, regulatory (inhibitor) subunit 11 |
| 1454848 | at | 1.05 | 0.614 | Ppp1r12c | protein phosphatase 1, regulatory (inhibitor) subunit 12C |
| 1439530 | a_at | -1.10 | 0.760 | Ppp1r13l | protein phosphatase 1, regulatory (inhibitor) subunit 13 like |
| 1450914 | at | 1.06 | 0.801 | Ppp1r14b | protein phosphatase 1, regulatory (inhibitor) subunit 14B |
| 1417701 | at | -1.27 | 0.542 | Ppp1r14c | protein phosphatase 1, regulatory (inhibitor) subunit 14c |
| 1426798 | a_at | -1.09 | 0.125 | Ppp1r15b | protein phosphatase 1, regulatory (inhibitor) subunit 15b |
| 1455080 | at | -1.34 | 0.014 | Ppp1r16b | protein phosphatase 1, regulatory (inhibitor) subunit 16B |
| 1417342 | at | 1.15 | 0.298 | Ppp1r2 | protein phosphatase 1, regulatory (inhibitor) subunit 2 |
| 1436590 | at | -1.25 | 0.222 | Ppp1r3b | protein phosphatase 1, regulatory (inhibitor) subunit 3B |
| 1433691 | at | 1.33 | 0.002 | Ppp1r3c | protein phosphatase 1, regulatory (inhibitor) subunit 3C |
| 1420734 | at | -1.32 | 0.002 | Ppp1r3f | protein phosphatase 1, regulatory (inhibitor) subunit 3F |
| 1417919 | at | 0.05 | 0.888 | Ppp1r7 | protein phosphatase 1, regulatory (inhibitor) subunit 7 |
| 1440285 | at | -1.51 | 0.002 | Ppp1r9a | protein phosphatase 1, regulatory (inhibitor) subunit 9A |
| 1417367 | at | 1.09 | 0.093 | Ppp2ca | protein phosphatase 2a, catalytic subunit, alpha isoform |
| 1421823 | a_at | -1.02 | 0.820 | Ppp2cb | protein phosphatase 2a, catalytic subunit, beta isoform |
| 1415819 | a_at | -1.12 | 0.067 | Ppp2r1a | protein phosphatase 2 (formerly 2A), regulatory subunit A (PR 65), alpha isoform |
| 1428265 | at | -1.03 | 0.658 | Ppp2r1b | protein phosphatase 2 (formerly 2A), regulatory subunit A (PR 65), beta isoform |
| 1429715 | at | -1.09 | 0.235 | Ppp2r2a | protein phosphatase 2 (formerly 2A), regulatory subunit B (PR 52), alpha isoform |
| 1448138 | at | -1.08 | 0.471 | Ppp2r4 | protein phosphatase 2A, regulatory subunit B (PR 53) |
| 1423911 | at | -1.14 | 0.048 | Ppp2r5a | protein phosphatase 2, regulatory subunit B (B56), alpha isoform |
| 1450560 | a_at | -1.03 | 0.737 | Ppp2r5d | protein phosphatase 2, regulatory subunit B (B56), delta isoform |
| 1420743 | a_at | 1.18 | 0.007 | Ppp3cc | protein phosphatase 3, catalytic subunit, gamma isoform |
| 1433591 | at | 1.05 | 0.352 | Ppp3r1 | protein phosphatase 3, regulatory subunit B, alpha isoform (calcineurin B, type I) |
| 1451242 | a_at | -1.15 | 0.764 | Ppp5c | protein phosphatase 5, catalytic subunit |
| 1426381 | at | 1.13 | 0.311 | Pprc1 | peroxisome proliferative activated receptor, gamma, coactivator-related 1 |
| 1416195 | at | -1.19 | 0.008 | Pps | putative phosphatase |
| 1418302 | at | -1.04 | 0.276 | Ppt2 | palmitoyl-protein thioesterase 2 |
| 1417731 | at | -1.17 | 0.064 | Pqbp1 | polyglutamine binding protein 1 |
| 1430747 | at | -1.11 | 0.229 | Pqlc1 | PQ loop repeat containing 1 |
| 1449250 | at | -1.30 | 0.033 | Prcc | papillary renal cell carcinoma (translocation-associated) |
| 1452190 | at | -1.03 | 0.812 | Prcp | prolylcarboxypeptidase (angiotensinase C) |
| 1420425 | at | 1.05 | 0.807 | Prdm1 | PR domain containing 1, with ZNF domain |
| 1455459 | at | -1.34 | 0.005 | Prdm15 | PR domain containing 15 |
| 1440745 | at | -1.47 | 0.024 | Prdm16 | PR domain containing 16 |
| 1453068 | at | 1.19 | 0.294 | Prdm2 | PR domain containing 2, with ZNF domain |
| 1452208 | at | 1.19 | 0.021 | Prdm4 | PR domain containing 4 |
| 1432057 | a_at | -1.15 | 0.066 | Prdm5 | PR domain containing 5 |

Supplementary table S1

| | | | | | |
|---|---|---|---|---|---|
| 1416000 | a_at | 1.02 | 0.198 | Prdx1 | peroxiredoxin 1 |
| 1418506 | a_at | 1.00 | 0.798 | Prdx2 | peroxiredoxin 2 |
| 1416292 | at | -1.09 | 0.034 | Prdx3 | peroxiredoxin 3 |
| 1416167 | at | 1.06 | 0.175 | Prdx4 | peroxiredoxin 4 |
| 1416381 | a_at | -1.10 | 0.001 | Prdx5 | peroxiredoxin 5 |
| 1423223 | a_at | -1.03 | 0.584 | Prdx6 | peroxiredoxin 6 |
| 1416502 | a_at | -1.09 | 0.272 | Preb | prolactin regulatory element binding |
| 1423167 | at | 1.04 | 0.278 | Prei3 | preimplantation protein 3 |
| 1437953 | at | -1.28 | 0.035 | Prei4 | preimplantation protein 4 |
| 1422796 | at | 1.25 | 0.111 | Prep | prolyl endopeptidase |
| 1426345 | at | 1.11 | 0.178 | Prepl | prolyl endopeptidase-like |
| 1417426 | at | 1.04 | 0.388 | Prg1 | proteoglycan 1, secretory granule |
| 1456112 | at | 1.34 | 0.023 | Prg4 | proteoglycan 4 (megakaryocyte stimulating factor, articular superficial zone protein) |
| 1452249 | at | -1.39 | 0.001 | Prickle1 | prickle like 1 (Drosophila) |
| 1428808 | at | 1.31 | 0.012 | Prickle2 | prickle-like 2 (Drosophila) |
| 1449061 | a_at | 1.08 | 0.255 | Prim1 | DNA primase, p49 subunit |
| 1429463 | at | -1.09 | 0.121 | Prkaa2 | protein kinase, AMP-activated, alpha 2 catalytic subunit |
| 1424119 | at | -1.15 | 0.105 | Prkab1 | protein kinase, AMP-activated, beta 1 non-catalytic subunit |
| 1435875 | at | 1.45 | 0.019 | Prkab2 | protein kinase, AMP-activated, beta 2 non-catalytic subunit |
| 1450519 | a_at | -1.10 | 0.082 | Prkaca | protein kinase, cAMP dependent, catalytic, alpha |
| 1420611 | at | -1.14 | 0.113 | Prkacb | protein kinase, cAMP dependent, catalytic, beta |
| 1417690 | at | -1.30 | 0.067 | Prkag1 | protein kinase, AMP-activated, gamma 1 non-catalytic subunit |
| 1423831 | at | 1.49 | 0.009 | Prkag2 | protein kinase, AMP-activated, gamma 2 non-catalytic subunit |
| 1444480 | at | 1.45 | 0.004 | Prkag3 | protein kinase, AMP-activated, gamma 3 non-catatlytic subunit |
| 1452032 | at | 1.23 | 0.037 | Prkar1a | protein kinase, cAMP dependent regulatory, type I, alpha |
| 1428783 | at | 1.09 | 0.414 | Prkar2a | protein kinase, cAMP dependent regulatory, type II alpha |
| 1446598 | at | 1.30 | 0.020 | Prkca | protein kinase C, alpha |
| 1419384 | at | -1.16 | 0.154 | Prkcabp | protein kinase C, alpha binding protein |
| 1456143 | at | -1.26 | 0.090 | Prkcbp1 | protein kinase C binding protein 1 |
| 1449956 | at | -1.24 | 0.044 | Prkce | protein kinase C, epsilon |
| 1434248 | at | 1.13 | 0.040 | Prkch | protein kinase C, eta |
| 1446596 | at | 1.02 | 0.879 | Prkcm | protein kinase C, mu |
| 1428229 | at | 1.05 | 0.236 | Prkcn | protein kinase C, nu |
| 1416339 | a_at | -1.07 | 0.306 | Prkcsh | protein kinase C substrate 80K-H |
| 1434333 | a_at | -1.03 | 0.695 | Prkd2 | protein kinase D2 |
| 1451576 | at | 1.02 | 0.860 | Prkdc | protein kinase, DNA activated, catalytic polypeptide |
| 1435460 | at | -1.32 | 0.149 | Prkg2 | protein kinase, cGMP-dependent, type II |
| 1422006 | at | 1.29 | 0.038 | Prkr | protein kinase, interferon-inducible double stranded RNA dependent |
| 1448923 | at | -1.00 | 0.934 | Prkra | protein kinase, interferon inducible double stranded RNA dependent activator |
| 1417425 | at | 1.04 | 0.738 | Prkrip1 | Prkr interacting protein 1 (IL11 inducible) |
| 1436746 | at | 1.08 | 0.351 | Prkwnk1 | protein kinase, lysine deficient 1 |

Supplementary table S1

| | | | | | |
|---|---|---|---|---|---|
| 1427196 | at | 1.09 | 0.249 | Prkwnk4 | protein kinase, lysine deficient 4 |
| 1424286 | at | -1.12 | 0.089 | Prkx | protein kinase, X-linked |
| 1448556 | at | -1.55 | 0.011 | Prlr | prolactin receptor |
| 1416130 | at | 1.52 | 0.003 | Prnp | prion protein |
| 1425265 | a_at | -1.06 | 0.130 | Prnpip1 | prion protein interacting protein 1 |
| 1417629 | at | -1.37 | 0.036 | Prodh | proline dehydrogenase |
| 1426246 | at | -1.00 | 0.916 | Pros1 | protein S (alpha) |
| 1423044 | at | -1.05 | 0.269 | Prosc | proline synthetase co-transcribed |
| 1437894 | at | -1.40 | 0.083 | Prox1 | prospero-related homeobox 1 |
| 1449635 | at | -1.09 | 0.465 | Prp19 | PRP19/PSO4 homolog (S. cerevisiae) |
| 1429614 | at | -1.01 | 0.872 | Prpf18 | PRP18 pre-mRNA processing factor 18 homolog (yeast) |
| 1424314 | at | -1.08 | 0.250 | Prpf3 | PRP3 pre-mRNA processing factor 3 homolog (yeast) |
| 1429724 | at | -1.12 | 0.015 | Prpf4 | PRP4 pre-mRNA processing factor 4 homolog (yeast) |
| 1422453 | at | -1.21 | 0.002 | Prpf8 | pre-mRNA processing factor 8 |
| 1416052 | at | 1.10 | 0.267 | Prps1 | phosphoribosyl pyrophosphate synthetase 1 |
| 1454843 | at | -1.09 | 0.206 | Prps2 | phosphoribosyl pyrophosphate synthetase 2 |
| 1428182 | at | 1.15 | 0.166 | Prpsap1 | phosphoribosyl pyrophosphate synthetase-associated protein 1 |
| 1428706 | at | -1.47 | 0.000 | Prr6 | proline-rich polypeptide 6 |
| 1416749 | at | -1.46 | 0.002 | Prss11 | protease, serine, 11 (Igf binding) |
| 1428365 | a_at | 1.22 | 0.049 | Prss15 | protease, serine, 15 |
| 1419722 | at | 1.02 | 0.821 | Prss19 | protease, serine, 19 (neuropsin) |
| 1434138 | at | 1.11 | 0.049 | Prune | prune homolog (Drosophila) |
| 1421813 | a_at | -1.18 | 0.013 | Psap | prosaposin |
| 1433478 | at | -1.02 | 0.752 | Psarl | presenilin associated, rhomboid-like |
| 1451064 | a_at | 1.16 | 0.009 | Psat1 | phosphoserine aminotransferase 1 |
| 1418183 | a_at | -1.16 | 0.116 | Pscd1 | pleckstrin homology, Sec7 and coiled-coil domains 1 |
| 1450103 | a_at | -1.01 | 0.897 | Pscd2 | pleckstrin homology, Sec7 and coiled-coil domains 2 |
| 1415679 | at | -1.11 | 0.018 | Psenen | presenilin enhancer 2 homolog (C. elegans) |
| 1417166 | at | -1.08 | 0.366 | Psip1 | PC4 and SFRS1 interacting protein 1 |
| 1433522 | at | -1.09 | 0.152 | Pskh1 | protein serine kinase H1 |
| 1415695 | at | 1.07 | 0.047 | Psma1 | proteasome (prosome, macropain) subunit, alpha type 1 |
| 1448206 | at | -1.11 | 0.008 | Psma2 | proteasome (prosome, macropain) subunit, alpha type 2 |
| 1448442 | a_at | 1.11 | 0.004 | Psma3 | proteasome (prosome, macropain) subunit, alpha type 3 |
| 1460339 | at | -1.04 | 0.353 | Psma4 | proteasome (prosome, macropain) subunit, alpha type 4 |
| 1424681 | a_at | 1.17 | 0.000 | Psma5 | proteasome (prosome, macropain) subunit, alpha type 5 |
| 1416506 | at | 1.01 | 0.660 | Psma6 | proteasome (prosome, macropain) subunit, alpha type 6 |
| 1420053 | at | 1.33 | 0.002 | Psmb1 | proteasome (prosome, macropain) subunit, beta type 1 |
| 1448632 | at | 1.27 | 0.078 | Psmb10 | proteasome (prosome, macropain) subunit, beta type 10 |
| 1448262 | at | -1.18 | 0.014 | Psmb2 | proteasome (prosome, macropain) subunit, beta type 2 |
| 1460198 | a_at | -1.02 | 0.621 | Psmb3 | proteasome (prosome, macropain) subunit, beta type 3 |
| 1451205 | at | 1.03 | 0.422 | Psmb4 | proteasome (prosome, macropain) subunit, beta type 4 |

Supplementary table S1

| | | | | | |
|---|---|---|---|---|---|
| 1415676 | a_at | -1.04 | 0.394 | Psmb5 | proteasome (prosome, macropain) subunit, beta type 5 |
| 1448822 | at | -1.10 | 0.013 | Psmb6 | proteasome (prosome, macropain) subunit, beta type 6 |
| 1416240 | at | 1.07 | 0.256 | Psmb7 | proteasome (prosome, macropain) subunit, beta type 7 |
| 1422962 | a_at | 1.01 | 0.932 | Psmb8 | proteosome (prosome, macropain) subunit, beta type 8 (large multifunctional protease 7) |
| 1450696 | at | -1.05 | 0.594 | Psmb9 | proteosome (prosome, macropain) subunit, beta type 9 (large multifunctional protease 2) |
| 1416005 | at | 1.07 | 0.108 | Psmc1 | protease (prosome, macropain) 26S subunit, ATPase 1 |
| 1426611 | at | -1.03 | 0.359 | Psmc2 | proteasome (prosome, macropain) 26S subunit, ATPase 2 |
| 1416282 | at | -1.05 | 0.261 | Psmc3 | proteasome (prosome, macropain) 26S subunit, ATPase 3 |
| 1415740 | at | -1.05 | 0.387 | Psmc5 | protease (prosome, macropain) 26S subunit, ATPase 5 |
| 1417771 | a_at | 1.04 | 0.174 | Psmc6 | proteasome (prosome, macropain) 26S subunit, ATPase, 6 |
| 1429291 | at | -1.01 | 0.914 | Psmd1 | proteasome (prosome, macropain) 26S subunit, non-ATPase, 1 |
| 1436559 | a_at | 1.00 | 0.999 | Psmd10 | proteosome (prosome, macropain) 26S subunit, non-ATPase, 10 |
| 1456104 | at | 1.21 | 0.162 | Psmd11 | proteasome (prosome, macropain) 26S subunit, non-ATPase, 11 |
| 1448492 | a_at | -1.03 | 0.455 | Psmd12 | proteasome (prosome, macropain) 26S subunit, non-ATPase, 12 |
| 1422459 | a_at | 1.03 | 0.698 | Psmd13 | proteasome (prosome, macropain) 26S subunit, non-ATPase, 13 |
| 1446521 | at | 1.19 | 0.078 | Psmd14 | proteasome (prosome, macropain) 26S subunit, non-ATPase, 14 |
| 1415831 | at | 1.04 | 0.354 | Psmd2 | proteasome (prosome, macropain) 26S subunit, non-ATPase, 2 |
| 1448479 | at | 1.15 | 0.033 | Psmd3 | proteasome (prosome, macropain) 26S subunit, non-ATPase, 3 |
| 1423234 | at | 1.09 | 0.172 | Psmd5 | proteasome (prosome, macropain) 26S subunit, non-ATPase, 5 |
| 1451056 | at | 1.11 | 0.367 | Psmd7 | proteasome (prosome, macropain) 26S subunit, non-ATPase, 7 |
| 1423296 | at | 1.31 | 0.000 | Psmd8 | proteasome (prosome, macropain) 26S subunit, non-ATPase, 8 |
| 1423386 | at | 1.13 | 0.031 | Psmd9 | proteasome (prosome, macropain) 26S subunit, non-ATPase, 9 |
| 1417056 | at | -1.24 | 0.005 | Psme1 | proteasome (prosome, macropain) 28 subunit, alpha |
| 1417189 | at | -1.13 | 0.035 | Psme2 | proteasome (prosome, macropain) 28 subunit, beta |
| 1418078 | at | 1.19 | 0.229 | Psme3 | proteaseome (prosome, macropain) 28 subunit, 3 |
| 1415673 | at | -1.05 | 0.428 | Psph | phosphoserine phosphatase |
| 1434516 | at | 1.08 | 0.043 | Pstk | phosphoseryl-tRNA kinase |
| 1421410 | a_at | 1.08 | 0.401 | Pstpip2 | proline-serine-threonine phosphatase-interacting protein 2 |
| 1424874 | a_at | 1.21 | 0.070 | Ptbp1 | polypyrimidine tract binding protein 1 |
| 1423470 | at | -1.01 | 0.945 | Ptbp2 | polypyrimidine tract binding protein 2 |
| 1428853 | at | -1.08 | 0.201 | Ptch1 | patched homolog 1 |
| 1420057 | at | 1.05 | 0.603 | Ptdsr | phosphatidylserine receptor |
| 1449739 | at | 1.14 | 0.020 | Ptdss1 | phosphatidylserine synthase 1 |
| 1434586 | a_at | -1.06 | 0.338 | Ptdss2 | phosphatidylserine synthase 2 |
| 1417449 | at | -1.02 | 0.775 | Pte1 | peroxisomal acyl-CoA thioesterase 1 |
| 1417667 | a_at | 1.59 | 0.004 | Pter | phosphotriesterase related |
| 1423860 | at | 1.05 | 0.452 | Ptgds | prostaglandin D2 synthase (brain) |
| 1436542 | at | 1.17 | 0.158 | Ptger1 | prostaglandin E receptor 1 (subtype EP1) |
| 1424208 | at | 1.87 | 0.000 | Ptger4 | prostaglandin E receptor 4 (subtype EP4) |
| 1417591 | at | -1.01 | 0.818 | Ptges2 | prostaglandin E synthase 2 |
| 1434891 | at | 1.65 | 0.000 | Ptgfrn | prostaglandin F2 receptor negative regulator |

Supplementary table S1

| | | | | | |
|---|---|---|---|---|---|
| 1427313 | at | 1.55 | 0.001 | Ptgir | prostaglandin I receptor (IP) |
| 1448816 | at | 2.04 | 0.000 | Ptgis | prostaglandin I2 (prostacyclin) synthase |
| 1420875 | at | 1.14 | 0.045 | Ptk9 | protein tyrosine kinase 9 |
| 1431292 | a_at | 1.22 | 0.000 | Ptk9l | protein tyrosine kinase 9-like (A6-related protein) |
| 1423455 | at | 1.05 | 0.116 | Ptma | prothymosin alpha |
| 1448254 | at | 1.66 | 0.038 | Ptn | pleiotrophin |
| 1416945 | at | -1.00 | 0.977 | Ptov1 | prostate tumor over expressed gene 1 |
| 1449322 | at | 1.17 | 0.019 | Ptp4a1 | protein tyrosine phosphatase 4a1 |
| 1418181 | at | -1.26 | 0.000 | Ptp4a3 | protein tyrosine phosphatase 4a3 |
| 1456315 | a_at | 1.28 | 0.002 | Ptpla | protein tyrosine phosphatase-like (proline instead of catalytic arginine), member a |
| 1449342 | at | -1.05 | 0.178 | Ptplb | protein tyrosine phosphatase-like (proline instead of catalytic arginine), member b |
| 1451225 | at | 1.18 | 0.000 | Ptpn11 | protein tyrosine phosphatase, non-receptor type 11 |
| 1455359 | at | 1.01 | 0.919 | Ptpn14 | protein tyrosine phosphatase, non-receptor type 14 |
| 1417140 | a_at | 1.03 | 0.685 | Ptpn2 | protein tyrosine phosphatase, non-receptor type 2 |
| 1419054 | a_at | 1.13 | 0.079 | Ptpn21 | protein tyrosine phosphatase, non-receptor type 21 |
| 1427128 | at | 1.10 | 0.287 | Ptpn23 | protein tyrosine phosphatase, non-receptor type 23 |
| 1455450 | at | -1.18 | 0.014 | Ptpn3 | protein tyrosine phosphatase, non-receptor type 3 |
| 1451037 | at | 1.24 | 0.020 | Ptpn9 | protein tyrosine phosphatase, non-receptor type 9 |
| 1416985 | at | 1.12 | 0.039 | Ptpns1 | protein tyrosine phosphatase, non-receptor type substrate 1 |
| 1425340 | a_at | 1.11 | 0.470 | Ptpra | protein tyrosine phosphatase, receptor type, A |
| 1422124 | a_at | 1.69 | 0.002 | Ptprc | protein tyrosine phosphatase, receptor type, C |
| 1435537 | at | -1.20 | 0.148 | Ptprd | protein tyrosine phosphatase, receptor type, D |
| 1420843 | at | 1.78 | 0.000 | Ptprf | protein tyrosine phosphatase, receptor type, F |
| 1444273 | at | -1.29 | 0.384 | Ptprj | protein tyrosine phosphatase, receptor type, J |
| 1444763 | at | 1.09 | 0.427 | Ptprk | protein tyrosine phosphatase, receptor type, K |
| 1422541 | at | -1.02 | 0.600 | Ptprm | protein tyrosine phosphatase, receptor type, M |
| 1416588 | at | 1.54 | 0.003 | Ptprn | protein tyrosine phosphatase, receptor type, N |
| 1460121 | at | -1.10 | 0.507 | Ptprs | protein tyrosine phosphatase, receptor type, S |
| 1450660 | at | -1.30 | 0.004 | Pts | 6-pyruvoyl-tetrahydropterin synthase |
| 1419620 | at | -1.19 | 0.559 | Pttg1 | pituitary tumor-transforming 1 |
| 1460325 | at | 1.06 | 0.280 | Pum1 | pumilio 1 (Drosophila) |
| 1419642 | at | 1.09 | 0.144 | Purb | purine rich element binding protein B |
| 1424970 | at | -1.06 | 0.572 | Purg | purine-rich element binding protein G |
| 1418491 | a_at | -1.04 | 0.663 | Pus3 | pseudouridine synthase 3 |
| 1423903 | at | 1.15 | 0.751 | Pvr | poliovirus receptor |
| 1424456 | at | -1.41 | 0.001 | Pvrl2 | poliovirus receptor-related 2 |
| 1424193 | at | 1.10 | 0.232 | Pwp2h | PWP2 (periodic tryptophan protein) homolog, yeast |
| 1451253 | at | 1.03 | 0.579 | Pxk | PX domain containing serine/threonine kinase |
| 1417841 | at | -1.39 | 0.001 | Pxmp2 | peroxisomal membrane protein 2 |
| 1420711 | a_at | -1.06 | 0.304 | Pxmp3 | peroxisomal membrane protein 3 |
| 1422780 | at | -1.23 | 0.074 | Pxmp4 | peroxisomal membrane protein 4 |

Supplementary table S1

| | | | | | |
|---|---|---|---|---|---|
| 1424027 | at | 1.06 | 0.303 | Pxn | paxillin |
| 1417346 | at | -1.16 | 0.019 | Pycard | PYD and CARD domain containing |
| 1448315 | a_at | 1.17 | 0.031 | Pycr2 | pyrroline-5-carboxylate reductase family, member 2 |
| 1448602 | at | -1.21 | 0.001 | Pygm | muscle glycogen phosphorylase |
| 1428249 | at | 1.10 | 0.017 | Pygo2 | pygopus 2 |
| 1416939 | at | -1.01 | 0.917 | Pyp | pyrophosphatase |
| 1423664 | at | -1.14 | 0.111 | Qdpr | quininoid dihydropteridine reductase |
| 1429317 | at | -1.22 | 0.084 | Qrsl1 | glutaminyl-tRNA synthase (glutamine-hydrolyzing)-like 1 |
| 1420832 | at | 2.17 | 0.000 | Qscn6 | quiescin Q6 |
| 1435469 | at | -1.62 | 0.000 | Qscn6l1 | quiescin Q6-like 1 |
| 1451199 | at | -1.02 | 0.806 | Qtrtd1 | queuine tRNA-ribosyltransferase domain containing 1 |
| 1434549 | at | 1.04 | 0.273 | Rab11a | RAB11a, member RAS oncogene family |
| 1423448 | at | -1.08 | 0.181 | Rab11b | RAB11B, member RAS oncogene family |
| 1455169 | at | -1.09 | 0.283 | Rab11fip2 | RAB11 family interacting protein 2 (class I) |
| 1438022 | at | -1.07 | 0.501 | Rab11fip3 | RAB11 family interacting protein 3 (class II) |
| 1427992 | a_at | -1.19 | 0.023 | Rab12 | RAB12, member RAS oncogene family |
| 1424894 | at | 1.35 | 0.005 | Rab13 | RAB13, member RAS oncogene family |
| 1420900 | a_at | 1.05 | 0.581 | Rab18 | RAB18, member RAS oncogene family |
| 1424064 | at | -1.08 | 0.127 | Rab1b | RAB1B, member RAS oncogene family |
| 1438097 | at | 1.12 | 0.398 | Rab20 | RAB20, member RAS oncogene family |
| 1451379 | at | -1.01 | 0.854 | Rab22a | RAB22A, member RAS oncogene family |
| 1421872 | at | 1.11 | 0.170 | Rab24 | RAB24, member RAS oncogene family |
| 1423990 | at | 1.09 | 0.192 | Rab28 | RAB28, member RAS oncogene family |
| 1452667 | at | 1.19 | 0.024 | Rab2b | RAB2B, member RAS oncogene family |
| 1423083 | at | 1.14 | 0.017 | Rab33b | RAB33B, member of RAS oncogene family |
| 1433922 | at | 1.20 | 0.000 | Rab35 | RAB35, member RAS oncogene family |
| 1422589 | at | -1.36 | 0.001 | Rab3a | RAB3A, member RAS oncogene family |
| 1434306 | at | 1.18 | 0.132 | Rab3ip | RAB3A interacting protein |
| 1416426 | at | 1.11 | 0.077 | Rab5a | RAB5A, member RAS oncogene family |
| 1459315 | at | 1.10 | 0.312 | Rab5c | RAB5C, member RAS oncogene family |
| 1448304 | a_at | 1.12 | 0.667 | Rab6 | RAB6, member RAS oncogene family |
| 1434914 | at | -1.13 | 0.144 | Rab6b | RAB6B, member RAS oncogene family |
| 1424015 | at | 1.18 | 0.023 | Rab6ip1 | Rab6 interacting protein 1 |
| 1424247 | at | -1.61 | 0.006 | Rab6ip2 | Rab6 interacting protein 2 |
| 1451362 | at | 1.20 | 0.004 | Rab7l1 | RAB7, member RAS oncogene family-like 1 |
| 1418692 | at | 1.04 | 0.572 | Rab8a | RAB8A, member RAS oncogene family |
| 1426799 | at | 1.23 | 0.013 | Rab8b | RAB8B, member RAS oncogene family |
| 1448391 | at | -1.27 | 0.002 | Rab9 | RAB9, member RAS oncogene family |
| 1427773 | a_at | -1.39 | 0.000 | Rabac1 | Rab acceptor 1 (prenylated) |
| 1443856 | at | 1.08 | 0.257 | Rabep1 | rabaptin, RAB GTPase binding effector protein 1 |
| 1424188 | at | 1.01 | 0.787 | Rabgap1 | RAB GTPase activating protein 1 |

Supplementary table S1

| | | | | | |
|---|---|---|---|---|---|
| 1426046 | a_at | -1.12 | 0.154 | Rabggta | Rab geranylgeranyl transferase, a subunit |
| 1419553 | a_at | 1.16 | 0.107 | Rabggtb | RAB geranylgeranyl transferase, b subunit |
| 1449669 | at | 1.13 | 0.011 | Rabif | RAB interacting factor |
| 1418039 | at | -1.11 | 0.278 | Rabl3 | RAB, member of RAS oncogene family-like 3 |
| 1424648 | at | -1.01 | 0.930 | Rabl4 | RAB, member of RAS oncogene family-like 4 |
| 1424575 | at | 1.12 | 0.349 | Rabl5 | RAB, member of RAS oncogene family-like 5 |
| 1423734 | at | -1.01 | 0.785 | Rac1 | RAS-related C3 botulinum substrate 1 |
| 1448414 | at | -1.10 | 0.372 | Rad1 | RAD1 homolog (S. pombe) |
| 1448762 | at | 1.02 | 0.830 | Rad17 | RAD17 homolog (S. pombe) |
| 1416161 | at | -1.02 | 0.738 | Rad21 | RAD21 homolog (S. pombe) |
| 1455391 | at | -1.13 | 0.156 | Rad23a | RAD23a homolog (S. cerevisiae) |
| 1455420 | at | 1.06 | 0.285 | Rad23b | RAD23b homolog (S. cerevisiae) |
| 1422770 | at | -1.14 | 0.214 | Rad51l3 | RAD51-like 3 (S. cerevisiae) |
| 1416602 | a_at | 1.06 | 0.358 | Rad52 | RAD52 homolog (S. cerevisiae) |
| 1417661 | at | -1.37 | 0.002 | Rad52b | RAD52 homolog B (S. cerevisiae) |
| 1429528 | at | -1.07 | 0.366 | Rae1 | RAE1 RNA export 1 homolog (S. pombe) |
| 1420090 | at | -1.27 | 0.016 | Raf1 | v-raf-1 leukemia viral oncogene 1 |
| 1431411 | a_at | 1.05 | 0.064 | Rai12 | retinoic acid induced 12 |
| 1417400 | at | 1.48 | 0.001 | Rai14 | retinoic acid induced 14 |
| 1435740 | at | 1.02 | 0.790 | Rai17 | retinoic acid induced 17 |
| 1452358 | at | 1.12 | 0.101 | Rai2 | retinoic acid induced 2 |
| 1450870 | at | 1.05 | 0.687 | Rala | v-ral simian leukemia viral oncogene homolog A (ras related) |
| 1417744 | a_at | 1.02 | 0.732 | Ralb | v-ral simian leukemia viral oncogene homolog B (ras related) |
| 1417248 | at | -1.09 | 0.969 | Ralbp1 | ralA binding protein 1 |
| 1460634 | at | 1.13 | 0.112 | Ralgds | ral guanine nucleotide dissociation stimulator |
| 1428789 | at | -1.34 | 0.008 | Ralgps2 | Ral GEF with PH domain and SH3 binding motif 2 |
| 1418573 | a_at | -1.07 | 0.670 | Raly | hnRNP-associated with lethal yellow |
| 1418187 | at | -1.11 | 0.356 | Ramp2 | receptor (calcitonin) activity modifying protein 2 |
| 1460551 | at | 1.10 | 0.124 | Ran | RAN, member RAS oncogene family |
| 1422547 | at | 1.01 | 0.898 | Ranbp1 | RAN binding protein 1 |
| 1424584 | a_at | 1.14 | 0.320 | Ranbp10 | RAN binding protein 10 |
| 1452673 | at | 1.10 | 0.247 | Ranbp3 | RAN binding protein 3 |
| 1426945 | at | 1.27 | 0.002 | Ranbp5 | RAN binding protein 5 |
| 1435167 | at | -1.11 | 0.105 | Ranbp6 | RAN binding protein 6 |
| 1422736 | at | -1.08 | 0.294 | Ranbp9 | RAN binding protein 9 |
| 1448020 | at | 1.07 | 0.035 | Rap1a | RAS-related protein-1a |
| 1435519 | at | 1.27 | 0.002 | Rap1b | RAS related protein 1b |
| 1425266 | a_at | 1.02 | 0.839 | Rap1gds1 | RAP1, GTP-GDP dissociation stimulator 1 |
| 1417914 | at | 1.05 | 0.571 | Rap2b | RAP2B, member of RAS oncogene family |
| 1460430 | at | 1.17 | 0.247 | Rap2c | RAP2C, member of RAS oncogene family |
| 1452833 | at | -1.10 | 0.129 | Rapgef2 | Rap guanine nucleotide exchange factor (GEF) 2 |

Supplementary table S1

| | | | | | |
|---|---|---|---|---|---|
| 1424470 | a_at | 1.08 | 0.422 | Rapgef3 | Rap guanine nucleotide exchange factor (GEF) 3 |
| 1425518 | at | 1.20 | 0.007 | Rapgef4 | Rap guanine nucleotide exchange factor (GEF) 4 |
| 1454906 | at | -1.32 | 0.041 | Rarb | retinoic acid receptor, beta |
| 1419415 | a_at | 1.22 | 0.053 | Rarg | retinoic acid receptor, gamma |
| 1416312 | at | 1.16 | 0.018 | Rars | arginyl-tRNA synthetase |
| 1424209 | at | 1.01 | 0.915 | Rarsl | arginyl-tRNA synthetase-like |
| 1455181 | at | -1.11 | 0.332 | Rasa2 | RAS p21 protein activator 2 |
| 1415850 | at | 1.16 | 0.025 | Rasa3 | RAS p21 protein activator 3 |
| 1440849 | at | -1.02 | 0.735 | Rasgrf2 | RAS protein-specific guanine nucleotide-releasing factor 2 |
| 1417804 | at | -1.43 | 0.047 | Rasgrp2 | RAS, guanyl releasing protein 2 |
| 1438030 | at | -1.15 | 0.468 | Rasgrp3 | RAS, guanyl releasing protein 3 |
| 1428016 | a_at | 1.08 | 0.475 | Rasip1 | Ras interacting protein 1 |
| 1423854 | a_at | 2.00 | 0.000 | Rasl11b | RAS-like, family 11, member B |
| 1422656 | at | -1.10 | 0.272 | Rasl2-9 | RAS-like, family 2, locus 9 |
| 1448855 | at | 1.34 | 0.010 | Rassf1 | Ras association (RalGDS/AF-6) domain family 1 |
| 1448546 | at | 1.13 | 0.196 | Rassf3 | Ras association (RalGDS/AF-6) domain family 3 |
| 1439622 | at | 1.14 | 0.107 | Rassf4 | Ras association (RalGDS/AF-6) domain family 4 |
| 1417850 | at | -1.16 | 0.499 | Rb1 | retinoblastoma 1 |
| 1449292 | at | -1.20 | 0.110 | Rb1cc1 | RB1-inducible coiled-coil 1 |
| 1455323 | at | -1.23 | 0.010 | Rbak | RB-associated KRAB repressor |
| 1428886 | at | -1.19 | 0.044 | Rbbp5 | retinoblastoma binding protein 5 |
| 1415775 | at | 1.08 | 0.018 | Rbbp7 | retinoblastoma binding protein 7 |
| 1427061 | at | 1.05 | 0.572 | Rbbp8 | retinoblastoma binding protein 8 |
| 1416174 | at | 1.27 | 0.001 | Rbbp9 | retinoblastoma binding protein 9 |
| 1451392 | at | -1.05 | 0.681 | Rbed1 | RNA binding motif and ELMO domain 1 |
| 1425981 | a_at | 1.04 | 0.605 | Rbl2 | retinoblastoma-like 2 |
| 1423740 | a_at | -1.23 | 0.050 | Rbm10 | RNA binding motif protein 10 |
| 1438741 | at | 1.39 | 0.001 | Rbm13 | RNA binding motif protein 13 |
| 1456514 | at | -1.15 | 0.067 | Rbm16 | RNA binding motif protein 16 |
| 1452691 | at | 1.16 | 0.000 | Rbm17 | RNA binding motif protein 17 |
| 1420608 | at | 1.08 | 0.118 | Rbm18 | RNA binding motif protein 18 |
| 1451106 | at | 1.03 | 0.797 | Rbm21 | RNA binding motif protein 21 |
| 1452675 | at | 1.22 | 0.009 | Rbm22 | RNA binding motif protein 22 |
| 1456058 | at | -1.05 | 0.449 | Rbm27 | RNA binding motif protein 27 |
| 1426513 | at | 1.02 | 0.859 | Rbm28 | RNA binding motif protein 28 |
| 1422660 | at | 1.12 | 0.020 | Rbm3 | RNA binding motif protein 3 |
| 1436641 | at | 1.07 | 0.369 | Rbm4 | RNA binding motif protein 4 |
| 1441319 | at | -1.25 | 0.151 | Rbm5 | RNA binding motif protein 5 |
| 1458912 | at | -1.16 | 0.556 | Rbm6 | RNA binding motif protein 6 |
| 1418119 | at | 1.02 | 0.569 | Rbm8a | RNA binding motif protein 8a |
| 1434938 | at | 1.02 | 0.830 | Rbm9 | RNA binding motif protein 9 |

Supplementary table S1

| | | | | | |
|---|---|---|---|---|---|
| 1423540 | at | -1.04 | 0.391 | Rbms2 | RNA binding motif, single stranded interacting protein 2 |
| 1455635 | at | -1.12 | 0.293 | Rbmx | RNA binding motif protein, X chromosome |
| 1416177 | at | -1.15 | 0.036 | Rbmxrt | RNA binding motif protein, X chromosome retrogene |
| 1448754 | at | 2.43 | 0.000 | Rbp1 | retinol binding protein 1, cellular |
| 1449461 | at | 1.11 | 0.206 | Rbp7 | retinol binding protein 7, cellular |
| 1455936 | a_at | 1.03 | 0.785 | Rbpms | RNA binding protein gene with multiple splicing |
| 1424008 | a_at | 1.05 | 0.274 | Rbpms2 | RNA binding protein with multiple splicing 2 |
| 1448957 | at | 1.14 | 0.243 | Rbpsuh | recombining binding protein suppressor of hairless (Drosophila) |
| 1418779 | at | 1.09 | 0.075 | Rce1 | Ras and a-factor-converting enzyme 1 homolog (S. cerevisiae) |
| 1432144 | a_at | 1.05 | 0.417 | Rchy1 | ring finger and CHY zinc finger domain containing 1 |
| 1417090 | at | 1.23 | 0.000 | Rcn1 | reticulocalbin 1 |
| 1424382 | at | 1.27 | 0.017 | Rcn3 | reticulocalbin 3, EF-hand calcium binding domain |
| 1437545 | at | -1.08 | 0.212 | Rcor1 | REST corepressor 1 |
| 1428342 | at | -1.19 | 0.154 | Rcor3 | REST corepressor 3 |
| 1450705 | at | 1.26 | 0.017 | Rdbp | RD RNA-binding protein |
| 1426968 | a_at | 1.03 | 0.841 | Rdh10 | retinol dehydrogenase 10 (all-trans) |
| 1433799 | at | -1.17 | 0.005 | Rdh13 | retinol dehydrogenase 13 (all-trans and 9-cis) |
| 1417438 | at | -1.18 | 0.052 | Rdh14 | retinol dehydrogenase 14 (all-trans and 9-cis) |
| 1451920 | a_at | -1.04 | 0.712 | Recc1 | replication factor C 1 |
| 1450784 | at | 1.55 | 0.000 | Reck | reversion-inducing-cysteine-rich protein with kazal motifs |
| 1418339 | at | 1.34 | 0.010 | Recql | RecQ protein-like |
| 1436196 | at | 1.08 | 0.553 | Rel | reticuloendotheliosis oncogene |
| 1419536 | a_at | 1.17 | 0.007 | Rela | v-rel reticuloendotheliosis viral oncogene homolog A (avian) |
| 1419685 | at | -1.06 | 0.377 | Rent1 | regulator of nonsense transcripts 1 |
| 1434043 | a_at | -1.04 | 0.607 | Repin1 | replication initiator 1 |
| 1416216 | at | -1.05 | 0.343 | Reps1 | RalBP1 associated Eps domain containing protein |
| 1454670 | at | 1.03 | 0.483 | Rere | arginine glutamic acid dipeptide (RE) repeats |
| 1451236 | at | -1.12 | 0.241 | Rerg | RAS-like, estrogen-regulated, growth-inhibitor |
| 1428227 | at | -1.24 | 0.029 | Rest | RE1-silencing transcription factor |
| 1436359 | at | -2.06 | 0.000 | Ret | ret proto-oncogene |
| 1449015 | at | -3.71 | 0.000 | Retnla | resistin like alpha |
| 1422624 | at | -1.16 | 0.019 | Rev1l | REV1-like (S. cerevisiae) |
| 1424632 | a_at | 1.09 | 0.341 | Rev3l | REV3-like, catalytic subunit of DNA polymerase zeta RAD54 like (S. cerevisiae) |
| 1448595 | a_at | 1.91 | 0.027 | Rex3 | reduced expression 3 |
| 1417503 | at | 1.01 | 0.917 | Rfc2 | replication factor C (activator 1) 2 |
| 1424321 | at | 1.00 | 0.948 | Rfc4 | replication factor C (activator 1) 4 |
| 1460704 | at | -1.16 | 0.035 | Rfng | radical fringe gene homolog (Drosophila) |
| 1426912 | at | -1.06 | 0.071 | Rfwd2 | ring finger and WD repeat domain 2 |
| 1423882 | at | -1.15 | 0.093 | Rfwd3 | ring finger and WD repeat domain 3 |
| 1436059 | at | 1.03 | 0.678 | Rfx1 | regulatory factor X, 1 (influences HLA class II expression) |
| 1437375 | at | -1.06 | 0.686 | Rfx3 | regulatory factor X, 3 (influences HLA class II expression) |

Supplementary table S1

| | | | | | |
|---|---|---|---|---|---|
| 1423103 | at | 1.22 | 0.017 | Rfx5 | regulatory factor X, 5 (influences HLA class II expression) |
| 1428534 | at | 1.21 | 0.008 | Rfxank | regulatory factor X-associated ankyrin-containing protein |
| 1424333 | at | 1.00 | 0.969 | Rg9mtd1 | RNA (guanine-9-) methyltransferase domain containing 1 |
| 1448948 | at | 1.21 | 0.002 | Rga | recombination activating gene 1 gene activation |
| 1449124 | at | -1.15 | 0.384 | Rgl1 | ral guanine nucleotide dissociation stimulator,-like 1 |
| 1450688 | at | -1.04 | 0.579 | Rgl2 | ral guanine nucleotide dissociation stimulator-like 2 |
| 1433587 | at | -1.04 | 0.994 | Rgmb | RGM domain family, member B |
| 1416882 | at | 1.22 | 0.003 | Rgs10 | regulator of G-protein signalling 10 |
| 1417786 | a_at | 1.52 | 0.000 | Rgs19 | regulator of G-protein signaling 19 |
| 1422763 | at | 1.03 | 0.666 | Rgs19ip1 | regulator of G-protein signaling 19 interacting protein 1 |
| 1419247 | at | -2.77 | 0.000 | Rgs2 | regulator of G-protein signaling 2 |
| 1425296 | a_at | 1.23 | 0.043 | Rgs3 | regulator of G-protein signaling 3 |
| 1416286 | at | 1.07 | 0.334 | Rgs4 | regulator of G-protein signaling 4 |
| 1424138 | at | 1.30 | 0.019 | Rhbdf1 | rhomboid family 1 (Drosophila) |
| 1435941 | at | -1.67 | 0.000 | Rhbdl4 | rhomboid, veinlet-like 4 (Drosophila) |
| 1417049 | at | -1.07 | 0.323 | Rhced | Rhesus blood group CE and D |
| 1416636 | at | -1.07 | 0.046 | Rheb | RAS-homolog enriched in brain |
| 1450632 | at | 1.21 | 0.031 | Rhoa | ras homolog gene family, member A |
| 1449110 | at | 1.02 | 0.697 | Rhob | ras homolog gene family, member B |
| 1448605 | at | 1.63 | 0.004 | Rhoc | ras homolog gene family, member C |
| 1416700 | at | 1.14 | 0.159 | Rhoe | ras homolog gene family, member E |
| 1418892 | at | -1.10 | 0.108 | Rhoj | ras homolog gene family, member J |
| 1440853 | at | -1.16 | 0.047 | Rhot1 | ras homolog gene family, member T1 |
| 1426822 | at | -1.31 | 0.000 | Rhot2 | ras homolog gene family, member T2 |
| 1449027 | at | 1.87 | 0.000 | Rhou | ras homolog gene family, member U |
| 1434628 | a_at | -1.12 | 0.346 | Rhpn2 | rhophilin, Rho GTPase binding protein 2 |
| 1442939 | at | -1.18 | 0.367 | Rif1 | Rap1 interacting factor 1 homolog (yeast) |
| 1426368 | at | -1.16 | 0.252 | Rin2 | Ras and Rab interactor 2 |
| 1422647 | at | -1.06 | 0.304 | Ring1 | ring finger protein 1 |
| 1423481 | at | 1.14 | 0.335 | Riok2 | RIO kinase 2 (yeast) |
| 1422650 | a_at | 1.12 | 0.758 | Riok3 | RIO kinase 3 (yeast) |
| 1436301 | at | -1.09 | 0.444 | Ripk5 | receptor interacting protein kinase 5 |
| 1420540 | a_at | -1.33 | 0.005 | Rit1 | Ras-like without CAAX 1 |
| 1427171 | at | 1.13 | 0.151 | Rlf | rearranged L-myc fusion sequence |
| 1422603 | at | 1.21 | 0.002 | Rnase4 | ribonuclease, RNase A family 4 |
| 1453833 | a_at | 1.03 | 0.626 | Rnaseh1 | ribonuclease H1 |
| 1452725 | a_at | -1.12 | 0.046 | Rnaseh2a | ribonuclease H2, large subunit |
| 1452734 | at | -1.05 | 0.225 | Rnaset2 | ribonuclease T2 |
| 1423102 | a_at | -1.07 | 0.106 | Rnf10 | ring finger protein 10 |
| 1448434 | at | 1.27 | 0.000 | Rnf103 | ring finger protein 103 |
| 1423304 | a_at | -1.06 | 0.296 | Rnf111 | ring finger 111 |

Supplementary table S1

| | | | | | |
|---|---|---|---|---|---|
| 1417250 | at | -1.13 | 0.161 | Rnf12 | ring finger protein 12 |
| 1426504 | a_at | 1.01 | 0.797 | Rnf121 | ring finger protein 121 |
| 1434611 | at | -1.19 | 0.029 | Rnf123 | ring finger protein 123 |
| 1424514 | at | -1.01 | 0.943 | Rnf126 | ring finger protein 126 |
| 1418318 | at | -1.15 | 0.011 | Rnf128 | ring finger protein 128 |
| 1420620 | a_at | 1.17 | 0.894 | Rnf13 | ring finger protein 13 |
| 1452289 | a_at | 1.22 | 0.000 | Rnf135 | ring finger protein 135 |
| 1429425 | at | -1.12 | 0.249 | Rnf139 | ring finger protein 139 |
| 1425929 | a_at | 1.10 | 0.543 | Rnf14 | ring finger protein 14 |
| 1449086 | at | 1.07 | 0.913 | Rnf141 | ring finger protein 141 |
| 1438404 | at | -1.03 | 0.597 | Rnf144 | ring finger protein 144 |
| 1429321 | at | 1.55 | 0.000 | Rnf149 | ring finger protein 149 |
| 1436906 | at | -1.15 | 0.136 | Rnf166 | ring finger protein 166 |
| 1430527 | a_at | -1.15 | 0.153 | Rnf167 | ring finger protein 167 |
| 1455585 | at | 1.04 | 0.321 | Rnf168 | ring fnger protein 168 |
| 1437207 | at | 1.09 | 0.922 | Rnf170 | ring finger protein 170 |
| 1451108 | at | 1.07 | 0.315 | Rnf185 | ring finger protein 185 |
| 1417508 | at | 1.42 | 0.001 | Rnf19 | ring finger protein (C3HC4 type) 19 |
| 1419916 | at | 1.18 | 0.003 | Rnf20 | ring finger protein 20 |
| 1452733 | at | 1.03 | 0.621 | Rnf24 | ring finger protein 24 |
| 1448674 | at | -1.12 | 0.194 | Rnf25 | ring finger protein 25 |
| 1423650 | at | 1.30 | 0.001 | Rnf26 | ring finger protein 26 |
| 1455051 | at | -1.17 | 0.006 | Rnf31 | ring finger protein 31 |
| 1415791 | at | 1.02 | 0.639 | Rnf34 | ring finger protein 34 |
| 1428101 | at | -1.10 | 0.054 | Rnf38 | ring finger protein 38 |
| 1423654 | a_at | 1.11 | 0.052 | Rnf4 | ring finger protein 4 |
| 1438191 | a_at | -1.05 | 0.618 | Rnf40 | ring finger protein 40 |
| 1423532 | at | -1.02 | 0.670 | Rnf44 | ring finger protein 44 |
| 1422888 | at | -1.14 | 0.110 | Rnf5 | ring finger protein 5 |
| 1427898 | at | -1.10 | 0.001 | Rnf6 | ring finger protein (C3H2C3 type) 6 |
| 1424120 | at | -1.03 | 0.681 | Rnf8 | ring finger protein 8 |
| 1453106 | a_at | 1.15 | 0.038 | Rnmt | RNA (guanine-7-) methyltransferase |
| 1428766 | at | -1.16 | 0.122 | Rnmtl1 | RNA methyltransferase like 1 |
| 1421265 | a_at | -1.11 | 0.372 | Rnpc1 | RNA-binding region (RNP1, RRM) containing 1 |
| 1456386 | at | -1.20 | 0.246 | Rnpc2 | RNA-binding region (RNP1, RRM) containing 2 |
| 1457367 | at | -1.28 | 0.030 | Rnpc3 | RNA-binding region (RNP1, RRM) containing 3 |
| 1451243 | at | 1.25 | 0.002 | Rnpep | arginyl aminopeptidase (aminopeptidase B) |
| 1454753 | at | 1.03 | 0.682 | Rnpepl1 | arginyl aminopeptidase (aminopeptidase B)-like 1 |
| 1448324 | at | -1.04 | 0.330 | Rnps1 | ribonucleic acid binding protein S1 |
| 1452808 | at | 1.02 | 0.624 | Rnut1 | RNA, U transporter 1 |
| 1427231 | at | 1.42 | 0.081 | Robo1 | roundabout homolog 1 (Drosophila) |

Supplementary table S1

| | | | | | |
|---|---|---|---|---|---|
| 1426206 | at | 1.02 | 0.817 | Robo4 | roundabout homolog 4 (Drosophila) |
| 1450994 | at | -1.14 | 0.084 | Rock1 | Rho-associated coiled-coil forming kinase 1 |
| 1423592 | at | 1.16 | 0.011 | Rock2 | Rho-associated coiled-coil forming kinase 2 |
| 1441407 | at | 1.15 | 0.172 | Rod1 | ROD1 regulator of differentiation 1 (S. pombe) |
| 1448996 | at | 1.18 | 0.099 | Rom1 | rod outer segment membrane protein 1 |
| 1429313 | at | 1.08 | 0.600 | Ror1 | receptor tyrosine kinase-like orphan receptor 1 |
| 1436326 | at | -1.07 | 0.542 | Rora | RAR-related orphan receptor alpha |
| 1425792 | a_at | -1.36 | 0.103 | Rorc | RAR-related orphan receptor gamma |
| 1418277 | at | -1.07 | 0.374 | Rp9h | retinitis pigmentosa 9 homolog (human) |
| 1423293 | at | -1.14 | 0.013 | Rpa1 | replication protein A1 |
| 1416433 | at | 1.19 | 0.044 | Rpa2 | replication protein A2 |
| 1448938 | at | -1.34 | 0.001 | Rpa3 | replication protein A3 |
| 1416705 | at | 1.10 | 0.031 | Rpe | ribulose-5-phosphate-3-epimerase |
| 1427467 | a_at | -1.02 | 0.748 | Rpgr | retinitis pigmentosa GTPase regulator |
| 1421144 | at | -1.02 | 0.644 | Rpgrip1 | retinitis pigmentosa GTPase regulator interacting protein 1 |
| 1444409 | at | -1.13 | 0.194 | Rph3al | rabphilin 3A-like (without C2 domains) |
| 1418337 | at | -1.11 | 0.036 | Rpia | ribose 5-phosphate isomerase A |
| 1415942 | at | -1.03 | 0.111 | Rpl10 | ribosomal protein 10 |
| 1431177 | a_at | 1.11 | 0.027 | Rpl10a | ribosomal protein L10A |
| 1417615 | a_at | 1.06 | 0.004 | Rpl11 | ribosomal protein L11 |
| 1416026 | a_at | 1.02 | 0.630 | Rpl12 | ribosomal protein L12 |
| 1449255 | a_at | 1.02 | 0.075 | Rpl15 | ribosomal protein L15 |
| 1453752 | at | 1.31 | 0.014 | Rpl17 | ribosomal protein L17 |
| 1437005 | a_at | -1.09 | 0.090 | Rpl18 | ribosomal protein L18 |
| 1428152 | a_at | 1.00 | 0.974 | Rpl18a | Ribosomal protein L18A |
| 1416219 | at | 1.01 | 0.868 | Rpl19 | ribosomal protein L19 |
| 1416603 | at | 1.03 | 0.593 | Rpl22 | ribosomal protein L22 |
| 1422859 | a_at | 1.04 | 0.560 | Rpl23 | ribosomal protein L23 |
| 1437975 | a_at | -1.04 | 0.380 | Rpl23a | ribosomal protein L23a |
| 1460201 | a_at | 1.02 | 0.021 | Rpl24 | ribosomal protein L24 |
| 1448109 | a_at | -1.03 | 0.256 | Rpl26 | ribosomal protein L26 |
| 1416099 | at | -1.07 | 0.178 | Rpl27 | ribosomal protein L27 |
| 1416074 | a_at | -1.02 | 0.414 | Rpl28 | ribosomal protein L28 |
| 1454627 | a_at | -1.04 | 0.426 | Rpl29 | ribosomal protein L29 |
| 1449323 | a_at | 1.21 | 0.001 | Rpl3 | ribosomal protein L3 |
| 1418273 | a_at | 1.09 | 0.001 | Rpl30 | ribosomal protein L30 |
| 1420381 | a_at | 1.19 | 0.015 | Rpl31 | ribosomal protein L31 |
| 1435897 | at | -1.08 | 0.009 | Rpl32 | ribosomal protein L32 |
| 1416243 | a_at | 1.04 | 0.037 | Rpl35 | ribosomal protein L35 |
| 1416519 | at | -1.03 | 0.226 | Rpl36 | ribosomal protein L36 |
| 1416807 | at | -1.13 | 0.031 | Rpl36a | ribosomal protein L36a |

Supplementary table S1

| | | | | | |
|---|---|---|---|---|---|
| 1453729 | a_at | -1.00 | 0.913 | Rpl37 | ribosomal protein L37 |
| 1416217 | a_at | -1.15 | 0.000 | Rpl37a | ribosomal protein L37a |
| 1450840 | a_at | 1.07 | 0.018 | Rpl39 | ribosomal protein L39 |
| 1449398 | at | -1.65 | 0.000 | Rpl3l | ribosomal protein L3-like |
| 1425183 | a_at | 1.06 | 0.002 | Rpl4 | ribosomal protein L4 |
| 1423665 | a_at | -1.05 | 0.157 | Rpl5 | ribosomal protein L5 |
| 1416546 | a_at | 1.03 | 0.259 | Rpl6 | ribosomal protein L6 |
| 1426162 | a_at | 1.01 | 0.775 | Rpl7 | ribosomal protein L7 |
| 1422613 | a_at | -1.03 | 0.093 | Rpl7a | ribosomal protein L7a |
| 1439780 | at | -1.09 | 0.139 | Rpl7l1 | ribosomal protein L7-like 1 |
| 1417762 | a_at | -1.04 | 0.169 | Rpl8 | ribosomal protein L8 |
| 1416420 | a_at | 1.03 | 0.407 | Rpl9 | ribosomal protein L9 |
| 1416277 | a_at | 1.00 | 0.991 | Rplp1 | ribosomal protein, large, P1 |
| 1415879 | a_at | -1.02 | 0.561 | Rplp2 | ribosomal protein, large P2 |
| 1416470 | a_at | 1.02 | 0.850 | Rpn1 | ribophorin I |
| 1418896 | a_at | 1.07 | 0.395 | Rpn2 | ribophorin II |
| 1417041 | at | -1.05 | 0.199 | Rpo1-1 | RNA polymerase 1-1 |
| 1451120 | at | 1.16 | 0.012 | Rpo1-3 | RNA polymerase 1-3 |
| 1460215 | at | -1.18 | 0.221 | Rpo1-4 | RNA polymerase 1-4 |
| 1422693 | a_at | 1.06 | 0.739 | Rpo2tc1 | RNA polymerase II transcriptional coactivator |
| 1419461 | at | 1.05 | 0.124 | Rpp14 | ribonuclease P 14 subunit (human) |
| 1449021 | at | -1.18 | 0.051 | Rpp21 | ribonuclease P 21 subunit (human) |
| 1423373 | at | -1.21 | 0.003 | Rpp30 | ribonuclease P/MRP 30 subunit (human) |
| 1424792 | at | 1.01 | 0.897 | Rpp40 | ribonuclease P 40 subunit (human) |
| 1424000 | a_at | -1.02 | 0.663 | Rps11 | ribosomal protein S11 |
| 1448344 | at | 1.09 | 0.025 | Rps12 | ribosomal protein S12 |
| 1415912 | a_at | -1.03 | 0.417 | Rps13 | ribosomal protein S13 |
| 1459986 | a_at | -1.01 | 0.824 | Rps17 | ribosomal protein S17 |
| 1435712 | a_at | -1.08 | 0.077 | Rps18 | ribosomal protein S18 |
| 1449243 | a_at | -1.02 | 0.670 | Rps19 | ribosomal protein S19 |
| 1431765 | a_at | -1.11 | 0.015 | Rps2 | ribosomal protein S2 |
| 1460175 | at | 1.02 | 0.323 | Rps23 | ribosomal protein S23 |
| 1455195 | at | 1.34 | 0.040 | Rps24 | ribosomal protein S24 |
| 1415876 | a_at | 1.01 | 0.871 | Rps26 | ribosomal protein S26 |
| 1415716 | a_at | 1.01 | 0.323 | Rps27 | ribosomal protein S27 |
| 1449196 | a_at | 1.12 | 0.039 | Rps27a | ribosomal protein S27a |
| 1450925 | a_at | 1.04 | 0.146 | Rps27l | ribosomal protein S27-like |
| 1451101 | a_at | 1.00 | 0.875 | Rps28 | ribosomal protein S28 |
| 1422475 | a_at | -1.02 | 0.486 | Rps3a | ribosomal protein S3a |
| 1416276 | a_at | 1.06 | 0.068 | Rps4x | ribosomal protein S4, X-linked |
| 1416054 | at | -1.02 | 0.463 | Rps5 | ribosomal protein S5 |

Supplementary table S1

| | | | | | |
|---|---|---|---|---|---|
| 1435816 | at | 1.14 | 0.245 | Rps6 | ribosomal protein S6 |
| 1417542 | at | -1.08 | 0.266 | Rps6ka2 | ribosomal protein S6 kinase, polypeptide 2 |
| 1438243 | at | 1.13 | 0.071 | Rps6ka4 | ribosomal protein S6 kinase, polypeptide 4 |
| 1431050 | at | -1.51 | 0.009 | Rps6ka5 | ribosomal protein S6 kinase, polypeptide 5 |
| 1434563 | at | 1.09 | 0.091 | Rps6kc1 | ribosomal protein S6 kinase polypeptide 1 |
| 1419364 | a_at | -1.04 | 0.605 | Rps7 | ribosomal protein S7 |
| 1436760 | a_at | -1.01 | 0.732 | Rps8 | ribosomal protein S8 |
| 1452710 | at | -1.04 | 0.482 | Rpusd4 | RNA pseudouridylate synthase domain containing 4 |
| 1450887 | at | -1.11 | 0.277 | Rqcd1 | rcd1 (required for cell differentiation) homolog 1 (S. pombe) |
| 1422562 | at | 1.10 | 0.413 | Rrad | Ras-related associated with diabetes |
| 1428905 | at | -1.06 | 0.413 | Rraga | Ras-related GTP binding A |
| 1415749 | a_at | 1.06 | 0.084 | Rragc | Ras-related GTP binding C |
| 1434909 | at | -1.04 | 0.222 | Rragd | Ras-related GTP binding D |
| 1418448 | at | 1.30 | 0.092 | Rras | Harvey rat sarcoma oncogene, subgroup R |
| 1448689 | at | 1.49 | 0.001 | Rras2 | related RAS viral (r-ras) oncogene homolog 2 |
| 1449221 | a_at | 1.23 | 0.294 | Rrbp1 | ribosome binding protein 1 |
| 1415878 | at | -1.00 | 0.963 | Rrm1 | ribonucleotide reductase M1 |
| 1416998 | at | 1.12 | 0.170 | Rrs1 | RRS1 ribosome biogenesis regulator homolog (S. cerevisiae) |
| 1436058 | at | 1.06 | 0.713 | Rsad2 | radical S-adenosyl methionine domain containing 2 |
| 1440325 | at | -1.06 | 0.362 | Rsb30 | RAB30, member RAS oncogene family |
| 1435174 | at | -1.26 | 0.054 | Rsbn1 | rosbin, round spermatid basic protein 1 |
| 1423119 | at | 1.12 | 0.114 | Rshl2 | radial spokehead-like 2 |
| 1456383 | at | 1.16 | 0.077 | Rsl1d1 | ribosomal L1 domain containing 1 |
| 1444750 | at | 1.23 | 0.745 | Rsn | restin (Reed-Steinberg cell-expressed intermediate filament-associated protein) |
| 1427278 | at | 1.17 | 0.139 | Rsnl2 | restin-like 2 |
| 1448584 | at | 1.00 | 0.999 | Rsrc1 | arginine/serine-rich coiled-coil 1 |
| 1419259 | at | 1.07 | 0.169 | Rsu1 | Ras suppressor protein 1 |
| 1424095 | at | 1.08 | 0.149 | Rtcd1 | RNA terminal phosphate cyclase domain 1 |
| 1419056 | at | -1.78 | 0.000 | Rtn2 | reticulon 2 (Z-band associated protein) |
| 1418101 | a_at | 1.09 | 0.299 | Rtn3 | reticulon 3 |
| 1437224 | at | 1.93 | 0.000 | Rtn4 | reticulon 4 |
| 1448800 | at | 1.05 | 0.683 | Rtn4ip1 | reticulon 4 interacting protein 1 |
| 1436868 | at | -1.43 | 0.050 | Rtn4rl1 | reticulon 4 receptor-like 1 |
| 1436588 | at | -1.22 | 0.152 | Rttn | rotatin |
| 1434131 | at | -1.05 | 0.483 | Rufy1 | RUN and FYVE domain containing 1 |
| 1454829 | at | 1.01 | 0.925 | Rundc1 | RUN domain containing 1 |
| 1436014 | a_at | -1.08 | 0.116 | Rusc1 | RUN and SH3 domain containing 1 |
| 1416585 | at | 1.21 | 0.022 | Ruvbl1 | RuvB-like protein 1 |
| 1422482 | at | -1.20 | 0.079 | Ruvbl2 | RuvB-like protein 2 |
| 1427913 | at | 1.01 | 0.892 | Rwdd1 | RWD domain containing 1 |
| 1454773 | at | -1.33 | 0.000 | Rxra | retinoid X receptor alpha |

Supplementary table S1

| | | | | | |
|---|---|---|---|---|---|
| 1416990 | at | -1.06 | 0.628 | Rxrb | retinoid X receptor beta |
| 1418782 | at | -1.64 | 0.000 | Rxrg | retinoid X receptor gamma |
| 1426829 | at | -1.03 | 0.596 | Rxrip110 | retinoid X receptor interacting protein 110 |
| 1428219 | at | 1.06 | 0.402 | Rybp | RING1 and YY1 binding protein |
| 1451789 | a_at | -1.02 | 0.859 | Ryk | receptor-like tyrosine kinase |
| 1445498 | at | -1.48 | 0.013 | Ryr2 | ryanodine receptor 2, cardiac |
| 1417421 | at | -1.05 | 0.330 | S100a1 | S100 calcium binding protein A1 |
| 1416762 | at | 1.11 | 0.361 | S100a10 | S100 calcium binding protein A10 (calpactin) |
| 1460351 | at | 1.17 | 0.025 | S100a11 | S100 calcium binding protein A11 (calizzarin) |
| 1418704 | at | 1.07 | 0.229 | S100a13 | S100 calcium binding protein A13 |
| 1425560 | a_at | 1.22 | 0.005 | S100a16 | S100 calcium binding protein A16 |
| 1424542 | at | 1.57 | 0.002 | S100a4 | S100 calcium binding protein A4 |
| 1421375 | a_at | 1.32 | 0.002 | S100a6 | S100 calcium binding protein A6 (calcyclin) |
| 1448756 | at | -1.35 | 0.134 | S100a9 | S100 calcium binding protein A9 (calgranulin B) |
| 1418595 | at | -1.26 | 0.005 | S3-12 | plasma membrane associated protein, S3-12 |
| 1416500 | at | 1.01 | 0.733 | Sacm1l | SAC1 (supressor of actin mutations 1, homolog)-like (S. cerevisiae) |
| 1434958 | at | 1.03 | 0.645 | Sacs | sacsin |
| 1454928 | at | 1.03 | 0.791 | Safb | scaffold attachment factor B |
| 1442254 | at | -1.13 | 0.239 | Sag | retinal S-antigen |
| 1424152 | at | -1.27 | 0.004 | Sall4 | sal-like 4 (Drosophila) |
| 1436356 | at | 1.32 | 0.004 | Samd4 | sterile alpha motif domain containing 4 |
| 1434438 | at | -1.09 | 0.664 | Samhd1 | SAM domain and HD domain, 1 |
| 1449480 | at | -1.20 | 0.051 | Sap18 | Sin3-associated polypeptide 18 |
| 1417719 | at | 1.33 | 0.004 | Sap30 | sin3 associated polypeptide |
| 1428163 | at | 1.05 | 0.130 | Sara2 | SAR1a gene homolog 2 (S. cerevisiae) |
| 1426257 | a_at | 1.08 | 0.301 | Sars1 | seryl-aminoacyl-tRNA synthetase 1 |
| 1448658 | at | -1.02 | 0.772 | Sart1 | squamous cell carcinoma antigen recognized by T-cells 1 |
| 1455795 | at | 1.36 | 0.012 | Sart2 | squamous cell carcinoma antigen recognized by T cells 2 |
| 1417547 | at | -1.27 | 0.096 | Sart3 | squamous cell carcinoma antigen recognized by T-cells 3 |
| 1416556 | at | -1.12 | 0.288 | Sas | sarcoma amplified sequence |
| 1448005 | at | -1.07 | 0.346 | Sash1 | SAM and SH3 domain containing 1 |
| 1420502 | at | 1.03 | 0.655 | Sat1 | spermidine/spermine N1-acetyl transferase 1 |
| 1430318 | at | -1.23 | 0.034 | Sat2 | spermidine/spermine N1-acetyl transferase 2 |
| 1416008 | at | 1.28 | 0.015 | Satb1 | special AT-rich sequence binding protein 1 |
| 1416075 | at | 1.07 | 0.732 | Sav1 | salvador homolog 1 (Drosophila) |
| 1426480 | at | -1.27 | 0.002 | Sbds | Shwachman-Bodian-Diamond syndrome homolog (human) |
| 1434707 | at | 1.11 | 0.322 | Sbf1 | SET binding factor 1 |
| 1426559 | at | -1.03 | 0.517 | Sbno1 | sno, strawberry notch homolog 1 (Drosophila) |
| 1451457 | at | -1.10 | 0.193 | Sc5d | sterol-C5-desaturase (fungal ERG3, delta-5-desaturase) homolog (S. cerevisae) |
| 1438294 | at | -1.25 | 0.064 | Sca1 | spinocerebellar ataxia 1 homolog (human) |
| 1422576 | at | 1.18 | 0.000 | Sca10 | spinocerebellar ataxia 10 homolog (human) |

Supplementary table S1

| | | | | | |
|---|---|---|---|---|---|
| 1426287 | at | 1.09 | 0.391 | Sca7 | spinocerebellar ataxia 7 homolog (human) |
| 1416611 | at | 1.05 | 0.239 | Scamp2 | secretory carrier membrane protein 2 |
| 1416294 | at | 1.01 | 0.791 | Scamp3 | secretory carrier membrane protein 3 |
| 1450393 | a_at | -1.01 | 0.930 | Scamp4 | secretory carrier membrane protein 4 |
| 1433520 | at | 1.07 | 0.211 | Scap | SREBP cleavage activating protein |
| 1418895 | at | 1.39 | 0.000 | Scap2 | src family associated phosphoprotein 2 |
| 1416050 | a_at | 1.21 | 0.005 | Scarb1 | scavenger receptor class B, member 1 |
| 1454704 | at | -1.06 | 0.411 | Scarb2 | scavenger receptor class B, member 2 |
| 1415964 | at | 1.39 | 0.000 | Scd1 | stearoyl-Coenzyme A desaturase 1 |
| 1415822 | at | 1.16 | 0.125 | Scd2 | stearoyl-Coenzyme A desaturase 2 |
| 1428335 | a_at | 1.13 | 0.039 | Scfd1 | sec1 family domain containing 1 |
| 1433608 | at | 1.23 | 0.031 | Scfd2 | sec1 family domain containing 2 |
| 1423025 | a_at | -1.45 | 0.053 | Schip1 | schwannomin interacting protein 1 |
| 1426241 | a_at | -1.29 | 0.172 | Scmh1 | sex comb on midleg homolog 1 |
| 1439204 | at | -1.16 | 0.330 | Scn3a | sodium channel, voltage-gated, type III, alpha |
| 1434008 | at | -1.40 | 0.058 | Scn4b | sodium channel, type IV, beta |
| 1458813 | at | 1.34 | 0.004 | Scn5a | sodium channel, voltage-gated, type V, alpha |
| 1436043 | at | 1.13 | 0.437 | Scn7a | sodium channel, voltage-gated, type VII, alpha |
| 1428364 | at | -1.13 | 0.012 | Scnm1 | sodium channel modifier 1 |
| 1423986 | a_at | 1.33 | 0.085 | Scotin | scotin gene |
| 1426555 | at | 1.31 | 0.010 | Scpep1 | serine carboxypeptidase 1 |
| 1455547 | at | -1.00 | 0.970 | Scrg3 | scrapie responsive gene 3 |
| 1436573 | at | 1.06 | 0.489 | Scrn3 | secernin 3 |
| 1453486 | a_at | -1.20 | 0.009 | Scube2 | signal peptide, CUB domain, EGF-like 2 |
| 1416486 | at | 1.05 | 0.393 | Scye1 | small inducible cytokine subfamily E, member 1 |
| 1451051 | a_at | -1.06 | 0.029 | Scyl1 | SCY1-like 1 (S. cerevisiae) |
| 1436314 | at | -1.09 | 0.470 | Scyl2 | SCY1-like 2 (S. cerevisiae) |
| 1452960 | at | 1.17 | 0.027 | Scyl3 | SCY1-like 3 (S. cerevisiae) |
| 1437088 | at | 1.06 | 0.406 | Sdad1 | SDA1 domain containing 1 |
| 1451017 | at | 1.02 | 0.658 | Sdbcag84 | serologically defined breast cancer antigen 84 |
| 1415943 | at | 1.19 | 0.031 | Sdc1 | syndecan 1 |
| 1448545 | at | 1.13 | 0.129 | Sdc2 | syndecan 2 |
| 1450027 | at | 1.07 | 0.606 | Sdc3 | syndecan 3 |
| 1417654 | at | -1.06 | 0.628 | Sdc4 | syndecan 4 |
| 1450941 | at | 1.26 | 0.003 | Sdcbp | syndecan binding protein |
| 1426455 | at | 1.30 | 0.000 | Sdccag10 | serologically defined colon cancer antigen 10 |
| 1427232 | at | -1.23 | 0.048 | Sdccag33 | serologically defined colon cancer antigen 33 |
| 1435385 | at | 1.19 | 0.075 | Sdccag33l | serologically defined colon cancer antigen 33 like |
| 1456538 | at | -1.00 | 0.976 | Sdccag8 | serologically defined colon cancer antigen 8 |
| 1416857 | at | -1.09 | 0.001 | Sdf2 | stromal cell derived factor 2 |
| 1423465 | at | 1.27 | 0.146 | Sdfr2 | stromal cell derived factor receptor 2 |

Supplementary table S1

| | | | | | |
|---|---|---|---|---|---|
| 1426584 | a_at | -1.43 | 0.001 | Sdh1 | sorbitol dehydrogenase 1 |
| 1426688 | at | -1.08 | 0.000 | Sdha | succinate dehydrogenase complex, subunit A, flavoprotein (Fp) |
| 1418005 | at | -1.07 | 0.006 | Sdhb | succinate dehydrogenase complex, subunit B, iron sulfur (Ip) |
| 1448630 | a_at | -1.43 | 0.000 | Sdhc | succinate dehydrogenase complex, subunit C, integral membrane protein |
| 1428235 | at | 1.01 | 0.771 | Sdhd | succinate dehydrogenase complex, subunit D, integral membrane protein |
| 1416778 | at | -1.10 | 0.332 | Sdpr | serum deprivation response |
| 1436817 | at | 1.03 | 0.582 | Sec10l1 | SEC10-like 1 (S. cerevisiae) |
| 1418223 | at | -1.13 | 0.005 | Sec11l1 | Sec11-like 1 (S. cerevisiae) |
| 1460698 | a_at | 1.15 | 0.002 | Sec11l3 | Sec11-like 3 (S. cerevisiae) |
| 1416241 | at | 1.03 | 0.497 | Sec13l1 | SEC13-like 1 (S. cerevisiae) |
| 1453412 | a_at | -1.18 | 0.043 | Sec14l1 | SEC14-like 1 (S. cerevisiae) |
| 1424230 | at | 1.11 | 0.447 | Sec15l1 | SEC15-like 1 (S. cerevisiae) |
| 1434804 | at | 1.07 | 0.151 | Sec15l2 | SEC15-like 2 (S. cerevisiae) |
| 1449063 | at | 1.03 | 0.680 | Sec22l1 | SEC22 vesicle trafficking protein-like 1 (S. cerevisiae) |
| 1441696 | at | -1.22 | 0.005 | Sec22l2 | SEC22 vesicle trafficking protein-like 2 (S. cerevisiae) |
| 1436102 | at | -1.22 | 0.068 | Sec22l3 | SEC22 vesicle trafficking protein-like 3 (S. cerevisiae) |
| 1423347 | at | 1.04 | 0.566 | Sec23a | SEC23A (S. cerevisiae) |
| 1416059 | at | -1.06 | 0.821 | Sec23b | SEC23B (S. cerevisiae) |
| 1426848 | at | -1.10 | 0.111 | Sec24b | SEC24 related gene family, member B (S. cerevisiae) |
| 1452147 | at | -1.03 | 0.564 | Sec24c | SEC24 related gene family, member C (S. cerevisiae) |
| 1426972 | at | 1.02 | 0.695 | Sec24d | SEC24 related gene family, member D (S. cerevisiae) |
| 1453014 | a_at | -1.02 | 0.823 | Sec31l1 | SEC31-like 1 (S. cerevisiae) |
| 1451142 | at | 1.23 | 0.028 | Sec3l1 | SEC3-like 1 (S. cerevisiae) |
| 1428470 | at | 1.05 | 0.491 | Sec5l1 | SEC5-like 1 (S. cerevisiae) |
| 1449944 | a_at | -1.23 | 0.047 | Sec61a2 | Sec61, alpha subunit 2 (S. cerevisiae) |
| 1417083 | at | 1.00 | 0.955 | Sec61b | Sec61 beta subunit |
| 1434750 | at | -1.07 | 0.343 | Sec6l1 | SEC6-like 1 (S. cerevisiae) |
| 1422684 | a_at | 1.09 | 0.234 | Sec8l1 | SEC8-like 1 (S. cerevisiae) |
| 1428496 | at | -1.21 | 0.206 | Secisbp2 | SECIS binding protein 2 |
| 1424201 | a_at | 1.10 | 0.204 | Seh1l | SEH1-like (S. cerevisiae |
| 1450699 | at | -1.54 | 0.000 | Selenbp1 | selenium binding protein 1 |
| 1448903 | at | 1.15 | 0.003 | selenoproteir | selenoprotein |
| 1423225 | at | -1.03 | 0.500 | Selk | selenoprotein K |
| 1448415 | a_at | 1.34 | 0.012 | Sema3b | sema domain, immunoglobulin domain (Ig), short basic domain, secreted, (semaphorin) 3B |
| 1429348 | at | 1.16 | 0.130 | Sema3c | sema domain, immunoglobulin domain (Ig), short basic domain, secreted, (semaphorin) 3C |
| 1429459 | at | 1.69 | 0.003 | Sema3d | sema domain, immunoglobulin domain (Ig), short basic domain, secreted, (semaphorin) 3D |
| 1455678 | at | -1.42 | 0.051 | Sema4b | sema domain, Ig domain, transmembrane domain (TM) and short cytoplasmic domain, (semaphorin) 4B |
| 1433920 | at | 1.66 | 0.000 | Sema4c | sema domain, Ig domain, transmembrane domain (TM) and short cytoplasmic domain, (semaphorin) 4C |
| 1434596 | at | -1.15 | 0.448 | Sema4d | sema domain, Ig domain, transmembrane domain (TM) and short cytoplasmic domain, (semaphorin) 4D |
| 1436458 | at | 1.15 | 0.138 | Sema6a | sema domain, transmembrane domain (TM), and cytoplasmic domain, (semaphorin) 6A |
| 1453055 | at | 1.17 | 0.004 | Sema6d | sema domain, transmembrane domain (TM), and cytoplasmic domain, (semaphorin) 6D |

Supplementary table S1

| | | | | | |
|---|---|---|---|---|---|
| 1459903 | at | 1.02 | 0.881 | Sema7a | sema domain, immunoglobulin domain (Ig), and GPI membrane anchor, (semaphorin) 7A |
| 1425466 | at | 1.11 | 0.561 | Senp2 | SUMO/sentrin specific protease 2 |
| 1448866 | at | -1.00 | 0.962 | Senp3 | SUMO/sentrin specific protease 3 |
| 1452627 | at | -1.05 | 0.564 | Senp6 | SUMO/sentrin specific protease 6 |
| 1454084 | a_at | 1.02 | 0.766 | Senp8 | SUMO/sentrin specific protease 8 |
| 1433973 | at | 1.15 | 0.026 | Sephs1 | selenophosphate synthetase 1 |
| 1418325 | at | 1.13 | 0.110 | Sephs2 | selenophosphate synthetase 2 |
| 1424394 | at | 1.06 | 0.342 | Sepm | selenoprotein M |
| 1426680 | at | -1.01 | 0.802 | Sepn1 | selenoprotein N, 1 |
| 1452141 | a_at | 1.03 | 0.113 | Sepp1 | selenoprotein P, plasma, 1 |
| 1423472 | at | 1.18 | 0.013 | septin 2 | septin 2 |
| 1448729 | a_at | -1.23 | 0.002 | septin 4 | septin 4 |
| 1436819 | at | 1.11 | 0.081 | septin 6 | septin 6 |
| 1454610 | at | 1.11 | 0.130 | septin 7 | septin 7 |
| 1417038 | at | 1.09 | 0.298 | septin 9 | septin 9 |
| 1416521 | at | 1.02 | 0.654 | Sepw1 | selenoprotein W, muscle 1 |
| 1418888 | a_at | 1.40 | 0.001 | Sepx1 | selenoprotein X 1 |
| 1440936 | at | -1.35 | 0.000 | Serac1 | serine active site containing 1 |
| 1434291 | a_at | 1.49 | 0.002 | Serf1 | small EDRK-rich factor 1 |
| 1421938 | at | 1.25 | 0.000 | Serhl | serine hydrolase-like |
| 1450138 | a_at | 1.14 | 0.028 | Serpinb6a | serine (or cysteine) proteinase inhibitor, clade B, member 6a |
| 1422804 | at | 1.44 | 0.000 | Serpinb6b | serine (or cysteine) proteinase inhibitor, clade B, member 6b |
| 1439790 | at | 1.24 | 0.030 | Serpinb9 | serine (or cysteine) proteinase inhibitor, clade B, member 9 |
| 1416666 | at | 1.57 | 0.001 | Serpine2 | serine (or cysteine) proteinase inhibitor, clade E, member 2 |
| 1416168 | at | 2.14 | 0.000 | Serpinf1 | serine (or cysteine) proteinase inhibitor, clade F, member 1 |
| 1416625 | at | 1.35 | 0.000 | Serping1 | serine (or cysteine) proteinase inhibitor, clade G, member 1 |
| 1450843 | a_at | 1.09 | 0.463 | Serpinh1 | serine (or cysteine) proteinase inhibitor, clade H, member 1 |
| 1416702 | at | -1.06 | 0.440 | Serpini1 | serine (or cysteine) proteinase inhibitor, clade I, member 1 |
| 1417406 | at | 1.86 | 0.006 | Sertad1 | SERTA domain containing 1 |
| 1444645 | at | -1.06 | 0.711 | Sertad2 | SERTA domain containing 2 |
| 1454877 | at | 1.61 | 0.000 | Sertad4 | SERTA domain containing 4 |
| 1454699 | at | -1.13 | 0.114 | Sesn1 | sestrin 1 |
| 1449303 | at | 1.11 | 0.003 | Sesn3 | sestrin 3 |
| 1455549 | at | -1.01 | 0.904 | Sestd1 | SEC14 and spectrin domains 1 |
| 1426853 | at | 1.09 | 0.410 | Set | SET translocation |
| 1435437 | at | 1.37 | 0.000 | Set7 | SET domain-containing protein 7 |
| 1451833 | a_at | 1.04 | 0.584 | Setdb1 | SET domain, bifurcated 1 |
| 1449333 | at | -1.02 | 0.520 | Sf3a1 | splicing factor 3a, subunit 1 |
| 1423811 | at | 1.06 | 0.490 | Sf3a3 | splicing factor 3a, subunit 3 |
| 1415985 | at | 1.03 | 0.564 | Sf3b3 | splicing factor 3b, subunit 3 |
| 1451185 | at | -1.15 | 0.029 | Sf3b5 | splicing factor 3b, subunit 5 |

Supplementary table S1

| | | | | | |
|---|---|---|---|---|---|
| 1424381 | at | 1.07 | 0.296 | Sf4 | splicing factor 4 |
| 1423796 | at | -1.15 | 0.132 | Sfpq | splicing factor proline/glutamine rich (polypyrimidine tract binding protein associated) |
| 1448201 | at | 3.74 | 0.005 | Sfrp2 | secreted frizzled-related sequence protein 2 |
| 1452396 | at | -1.32 | 0.000 | Sfrs15 | splicing factor, arginine/serine-rich 15 |
| 1428961 | a_at | 1.01 | 0.946 | Sfrs16 | splicing factor, arginine/serine-rich 16 (suppressor-of-white-apricot homolog, Drosophila) |
| 1452884 | at | -1.17 | 0.157 | Sfrs2ip | splicing factor, arginine/serine-rich 2, interacting protein |
| 1448778 | at | -1.15 | 0.118 | Sfrs4 | splicing factor, arginine/serine-rich 4 (SRp75) |
| 1443547 | at | -1.00 | 0.975 | Sfrs5 | splicing factor, arginine/serine-rich 5 (SRp40, HRS) |
| 1448454 | at | -1.09 | 0.067 | Sfrs6 | splicing factor, arginine/serine-rich 6 |
| 1424033 | at | -1.00 | 0.998 | Sfrs7 | splicing factor, arginine/serine-rich 7 |
| 1433677 | at | -1.15 | 0.789 | Sfrs8 | splicing factor, arginine/serine-rich 8 |
| 1417727 | at | -1.01 | 0.892 | Sfrs9 | splicing factor, arginine/serine rich 9 |
| 1417559 | at | -1.07 | 0.335 | Sfxn1 | sideroflexin 1 |
| 1417893 | at | 1.53 | 0.002 | Sfxn3 | sideroflexin 3 |
| 1418873 | at | -1.13 | 0.318 | Sfxn4 | sideroflexin 4 |
| 1436618 | at | -1.30 | 0.032 | Sfxn5 | sideroflexin 5 |
| 1422654 | at | -1.22 | 0.001 | Sgca | sarcoglycan, alpha (dystrophin-associated glycoprotein) |
| 1420688 | a_at | 1.21 | 0.042 | Sgce | sarcoglycan, epsilon |
| 1421254 | a_at | -1.11 | 0.210 | Sgcg | sarcoglycan, gamma (dystrophin-associated glycoprotein) |
| 1416041 | at | -1.03 | 0.638 | Sgk | serum/glucocorticoid regulated kinase |
| 1437370 | at | -1.24 | 0.158 | Sgol2 | shugoshin-like 2 (S. pombe) |
| 1415892 | at | 1.08 | 0.269 | Sgpl1 | sphingosine phosphate lyase 1 |
| 1420821 | at | 1.33 | 0.057 | Sgpp1 | sphingosine-1-phosphate phosphatase 1 |
| 1451067 | at | -1.01 | 0.883 | Sgta | small glutamine-rich tetratricopeptide repeat (TPR)-containing, alpha |
| 1437977 | at | 1.25 | 0.092 | Sgtb | small glutamine-rich tetratricopeptide repeat (TPR)-containing, beta |
| 1448874 | a_at | -1.07 | 0.599 | Sh2bpsm1 | SH2-B PH domain containing signaling mediator 1 |
| 1415886 | at | -1.22 | 0.113 | Sh2d3c | SH2 domain containing 3C |
| 1453377 | at | 1.59 | 0.001 | Sh2d4a | SH2 domain containing 4A |
| 1422644 | at | 1.33 | 0.000 | Sh3bgr | SH3-binding domain glutamic acid-rich protein |
| 1421871 | at | 1.29 | 0.003 | Sh3bgrl | SH3-binding domain glutamic acid-rich protein like |
| 1416528 | at | 1.02 | 0.664 | Sh3bgrl3 | SH3 domain binding glutamic acid-rich protein-like 3 |
| 1455250 | at | -1.02 | 0.702 | Sh3bp4 | SH3-domain binding protein 4 |
| 1421923 | at | 1.18 | 0.050 | Sh3bp5 | SH3-domain binding protein 5 (BTK-associated) |
| 1445781 | at | -1.27 | 0.085 | Sh3d19 | SH3 domain protein D19 |
| 1419329 | at | 1.33 | 0.003 | Sh3d4 | SH3 domain protein 4 |
| 1455385 | at | 1.09 | 0.194 | Sh3glb1 | SH3-domain GRB2-like B1 (endophilin) |
| 1426499 | at | -1.31 | 0.000 | Sh3glb2 | SH3-domain GRB2-like endophilin B2 |
| 1456261 | at | -1.31 | 0.003 | Sh3kbp1 | SH3-domain kinase binding protein 1 |
| 1455149 | at | 1.02 | 0.782 | Sh3md2 | SH3 multiple domains 2 |
| 1419137 | at | -1.06 | 0.273 | Shank3 | SH3/ankyrin domain gene 3 |
| 1422854 | at | 1.05 | 0.587 | Shc1 | src homology 2 domain-containing transforming protein C1 |

Supplementary table S1

| | | | | | |
|---|---|---|---|---|---|
| 1418575 | at | 1.02 | 0.197 | Shfdg1 | split hand/foot deleted gene 1 |
| 1424174 | at | 1.02 | 0.905 | Shkbp1 | Sh3kbp1 binding protein 1 |
| 1425177 | at | 1.17 | 0.424 | Shmt1 | serine hydroxymethyl transferase 1 (soluble) |
| 1426423 | at | 1.09 | 0.416 | Shmt2 | serine hydroxymethyl transferase 2 (mitochondrial) |
| 1423129 | at | 1.07 | 0.173 | Shoc2 | soc-2 (suppressor of clear) homolog (C. elegans) |
| 1451854 | a_at | -1.16 | 0.130 | Shrm | shroom |
| 1423390 | at | -1.12 | 0.521 | Siah1a | seven in absentia 1A |
| 1448170 | at | -1.21 | 0.187 | Siah2 | seven in absentia 2 |
| 1426940 | at | -1.00 | 0.878 | Sidt2 | SID1 transmembrane family, member 2 |
| 1419900 | at | -1.16 | 0.086 | Sin3a | transcriptional regulator, SIN3A (yeast) |
| 1451044 | at | -1.10 | 0.123 | Sip1 | survivor of motor neuron protein interacting protein 1 |
| 1416206 | at | 1.05 | 0.794 | Sipa1 | signal-induced proliferation associated gene 1 |
| 1439755 | at | -1.04 | 0.539 | Sipa1l1 | signal-induced proliferation-associated 1 like 1 |
| 1418640 | at | -1.17 | 0.169 | Sirt1 | sirtuin 1 ((silent mating type information regulation 2, homolog) 1 (S. cerevisiae) |
| 1423507 | a_at | 1.08 | 0.010 | Sirt2 | sirtuin 2 (silent mating type information regulation 2, homolog) 2 (S. cerevisiae) |
| 1417892 | a_at | -1.25 | 0.039 | Sirt3 | sirtuin 3 (silent mating type information regulation 2, homolog) 3 (S. cerevisiae) |
| 1426847 | at | -1.16 | 0.009 | Sirt4 | sirtuin 4 (silent mating type information regulation 2 homolog) 4 (S. cerevisiae) |
| 1428915 | at | -1.04 | 0.457 | Sirt5 | sirtuin 5 (silent mating type information regulation 2 homolog) 5 (S. cerevisiae) |
| 1429631 | at | -1.21 | 0.133 | Sirt6 | sirtuin 6 (silent mating type information regulation 2, homolog) 6 (S. cerevisiae) |
| 1424238 | at | 1.05 | 0.723 | Sirt7 | sirtuin 7 (silent mating type information regulation 2, homolog) 7 (S. cerevisiae) |
| 1417080 | a_at | -1.14 | 0.012 | Sitpec | signaling intermediate in Toll pathway-evolutionarily conserved |
| 1452020 | a_at | -1.33 | 0.007 | Siva | Cd27 binding protein (Hindu God of destruction) |
| 1456862 | at | -1.13 | 0.238 | Six4 | sine oculis-related homeobox 4 homolog (Drosophila) |
| 1427560 | at | -1.28 | 0.011 | Six5 | sine oculis-related homeobox 5 homolog (Drosophila) |
| 1416541 | at | -1.15 | 0.262 | Skd3 | suppressor of K+ transport defect 3 |
| 1444785 | at | 1.06 | 0.473 | Ski | Sloan-Kettering viral oncogene homolog |
| 1452214 | at | 1.31 | 0.002 | Skil | SKI-like |
| 1426718 | at | -1.03 | 0.499 | Skiv2l2 | superkiller viralicidic activity 2-like 2 (S. cerevisiae) |
| 1423149 | at | -1.10 | 0.695 | Skp1a | S-phase kinase-associated protein 1A |
| 1419315 | at | 1.58 | 0.004 | Slamf9 | SLAM family member 9 |
| 1460168 | at | -1.06 | 0.117 | Slbp | stem-loop binding protein |
| 1420361 | at | 1.18 | 0.049 | Slc11a1 | solute carrier family 11 (proton-coupled divalent metal ion transporters), member 1 |
| 1417622 | at | -1.04 | 0.457 | Slc12a2 | solute carrier family 12, member 2 |
| 1417446 | at | 1.50 | 0.002 | Slc12a4 | solute carrier family 12, member 4 |
| 1418257 | at | 1.08 | 0.182 | Slc12a7 | solute carrier family 12, member 7 |
| 1417600 | at | 1.28 | 0.662 | Slc15a2 | solute carrier family 15 (H+/peptide transporter), member 2 |
| 1448385 | at | -1.12 | 0.135 | Slc15a4 | solute carrier family 15, member 4 |
| 1415802 | at | 1.07 | 0.143 | Slc16a1 | solute carrier family 16 (monocarboxylic acid transporters), member 1 |
| 1434592 | at | -1.53 | 0.006 | Slc16a10 | solute carrier family 16 (monocarboxylic acid transporters), member 10 |
| 1418445 | at | -1.07 | 0.626 | Slc16a2 | solute carrier family 16 (monocarboxylic acid transporters), member 2 |
| 1456914 | at | -1.18 | 0.031 | Slc16a4 | solute carrier family 16 (monocarboxylic acid transporters), member 4 |

Supplementary table S1

| | | | | | |
|---|---|---|---|---|---|
| 1417884 | at | 1.45 | 0.005 | Slc16a6 | solute carrier family 16 (monocarboxylic acid transporters), member 6 |
| 1448502 | at | -2.12 | 0.000 | Slc16a7 | solute carrier family 16 (monocarboxylic acid transporters), member 7 |
| 1429727 | at | -1.52 | 0.017 | Slc16a9 | solute carrier family 16 (monocarboxylic acid transporters), member 9 |
| 1417902 | at | 1.01 | 0.835 | Slc19a2 | solute carrier family 19 (thiamine transporter), member 2 |
| 1448299 | at | 1.27 | 0.028 | Slc1a1 | solute carrier family 1 (neuronal/epithelial high affinity glutamate transporter, system Xag), member 1 |
| 1456003 | a_at | 1.64 | 0.006 | Slc1a4 | solute carrier family 1 (glutamate/neutral amino acid transporter), member 4 |
| 1416629 | at | 1.28 | 0.002 | Slc1a5 | solute carrier family 1 (neutral amino acid transporter), member 5 |
| 1448568 | a_at | 1.54 | 0.005 | Slc20a1 | solute carrier family 20, member 1 |
| 1457302 | at | -1.21 | 0.151 | Slc20a2 | solute carrier family 20, member 2 |
| 1420444 | at | -2.51 | 0.000 | Slc22a3 | solute carrier family 22 (organic cation transporter), member 3 |
| 1450395 | at | -1.48 | 0.312 | Slc22a5 | solute carrier family 22 (organic cation transporter), member 5 |
| 1426586 | at | -1.14 | 0.002 | Slc25a11 | solute carrier family 25 (mitochondrial carrier oxoglutarate carrier), member 11 |
| 1428440 | at | -1.03 | 0.523 | Slc25a12 | solute carrier family 25 (mitochondrial carrier, Aralar), member 12 |
| 1449481 | at | 1.06 | 0.177 | Slc25a13 | solute carrier family 25 (mitochondrial carrier, adenine nucleotide translocator), member 13 |
| 1417154 | at | 1.16 | 0.020 | Slc25a14 | solute carrier family 25 (mitochondrial carrier, brain), member 14 |
| 1420967 | at | -1.09 | 0.257 | Slc25a15 | solute carrier family 25 (mitochondrial carrier ornithine transporter), member 15 |
| 1434996 | at | 1.02 | 0.731 | Slc25a16 | solute carrier family 25 (mitochondrial carrier, Graves disease autoantigen), member 16 |
| 1420038 | at | -1.03 | 0.548 | Slc25a18 | solute carrier family 25 (mitochondrial carrier), member 18 |
| 1424316 | at | -1.22 | 0.016 | Slc25a19 | solute carrier family 25 (mitochondrial deoxynucleotide carrier), member 19 |
| 1423108 | at | -1.06 | 0.040 | Slc25a20 | solute carrier family 25 (mitochondrial carnitine/acylcarnitine translocase), member 20 |
| 1452653 | at | -1.68 | 0.000 | Slc25a22 | solute carrier family 25 (mitochondrial carrier, glutamate), member 22 |
| 1452717 | at | 1.29 | 0.011 | Slc25a24 | solute carrier family 25 (mitochondrial carrier, phosphate carrier), member 24 |
| 1419928 | at | -1.39 | 0.001 | Slc25a26 | solute carrier family 25 (mitochondrial carrier, phosphate carrier), member 26 |
| 1424776 | a_at | -1.22 | 0.005 | Slc25a28 | solute carrier family 25, member 28 |
| 1423979 | a_at | -1.28 | 0.068 | Slc25a29 | solute carrier family 25 (mitochondrial carrier, palmitoylcarnitine transporter), member 29 |
| 1440302 | at | -1.17 | 0.022 | Slc25a3 | solute carrier family 25 (mitochondrial carrier, phosphate carrier), member 3 |
| 1433898 | at | -1.05 | 0.505 | Slc25a30 | solute carrier family 25, member 30 |
| 1430542 | a_at | 1.03 | 0.271 | Slc25a5 | solute carrier family 25 (mitochondrial carrier, adenine nucleotide translocator), member 5 |
| 1427195 | at | 1.05 | 0.378 | Slc26a2 | solute carrier family 26 (sulfate transporter), member 2 |
| 1416275 | at | -2.20 | 0.000 | Slc26a6 | solute carrier family 26, member 6 |
| 1422811 | at | -1.80 | 0.003 | Slc27a1 | solute carrier family 27 (fatty acid transporter), member 1 |
| 1424441 | at | 1.09 | 0.418 | Slc27a4 | solute carrier family 27 (fatty acid transporter), member 4 |
| 1450639 | at | 1.01 | 0.882 | Slc28a2 | solute carrier family 28 (sodium-coupled nucleoside transporter), member 2 |
| 1455731 | at | 1.13 | 0.039 | Slc29a3 | solute carrier family 29 (nucleoside transporters), member 3 |
| 1426600 | at | 1.34 | 0.025 | Slc2a1 | solute carrier family 2 (facilitated glucose transporter), member 1 |
| 1415958 | at | -1.56 | 0.003 | Slc2a4 | solute carrier family 2 (facilitated glucose transporter), member 4 |
| 1436164 | at | 1.07 | 0.331 | Slc30a1 | solute carrier family 30 (zinc transporter), member 1 |
| 1418843 | at | 1.51 | 0.001 | Slc30a4 | solute carrier family 30 (zinc transporter), member 4 |
| 1422497 | at | -1.13 | 0.012 | Slc30a5 | solute carrier family 30 (zinc transporter), member 5 |
| 1424241 | at | -1.03 | 0.691 | Slc30a6 | solute carrier family 30 (zinc transporter), member 6 |
| 1426668 | at | 1.14 | 0.007 | Slc30a9 | solute carrier family 30 (zinc transporter), member 9 |

Supplementary table S1

| | | | | | |
|---|---|---|---|---|---|
| 1416654 | at | -1.20 | 0.014 | Slc31a2 | solute carrier family 31, member 2 |
| 1423621 | a_at | 1.09 | 0.230 | Slc33a1 | solute carrier family 33 (acetyl-CoA transporter), member 1 |
| 1417538 | at | -1.01 | 0.730 | Slc35a1 | solute carrier family 35 (CMP-sialic acid transporter), member 1 |
| 1439433 | a_at | 1.00 | 0.964 | Slc35a2 | solute carrier family 35 (UDP-galactose transporter), member 2 |
| 1429648 | at | 1.05 | 0.960 | Slc35a3 | solute carrier family 35 (UDP-N-acetylglucosamine (UDP-GlcNAc) transporter), member 3 |
| 1416110 | at | 1.16 | 0.265 | Slc35a4 | solute carrier family 35, member A4 |
| 1419972 | at | 1.01 | 0.932 | Slc35a5 | solute carrier family 35, member A5 |
| 1448769 | at | -1.04 | 0.426 | Slc35b1 | solute carrier family 35, member B1 |
| 1423927 | at | -1.03 | 0.656 | Slc35b2 | solute carrier family 35, member B2 |
| 1448937 | at | 1.06 | 0.264 | Slc35b3 | solute carrier family 35, member B3 |
| 1433595 | at | 1.13 | 0.071 | Slc35d1 | solute carrier family 35 (UDP-glucuronic acid/UDP-N-acetylgalactosamine dual transporter), member D1 |
| 1428805 | at | 1.03 | 0.590 | Slc35e3 | solute carrier family 35, member E3 |
| 1452059 | at | 1.11 | 0.002 | Slc35f5 | solute carrier family 35, member F5 |
| 1436521 | at | -1.87 | 0.000 | Slc36a2 | solute carrier family 36 (proton/amino acid symporter), member 2 |
| 1455249 | at | 1.02 | 0.780 | Slc36a4 | solute carrier family 36 (proton/amino acid symporter), member 4 |
| 1424920 | at | 1.21 | 0.635 | Slc37a3 | solute carrier family 37 (glycerol-3-phosphate transporter), member 3 |
| 1429593 | at | 1.27 | 0.127 | Slc38a2 | solute carrier family 38, member 2 |
| 1418706 | at | -1.46 | 0.000 | Slc38a3 | solute carrier family 38, member 3 |
| 1428111 | at | 1.36 | 0.000 | Slc38a4 | solute carrier family 38, member 4 |
| 1424424 | at | 1.12 | 0.053 | Slc39a1 | solute carrier family 39 (zinc transporter), member 1 |
| 1433751 | at | 1.12 | 0.194 | Slc39a10 | solute carrier family 39 (zinc transporter), member 10 |
| 1423926 | at | -1.07 | 0.311 | Slc39a13 | solute carrier family 39 (metal ion transporter), member 13 |
| 1427035 | at | 1.13 | 0.047 | Slc39a14 | solute carrier family 39 (zinc transporter), member 14 |
| 1424674 | at | 1.66 | 0.000 | Slc39a6 | solute carrier family 39 (metal ion transporter), member 6 |
| 1452765 | at | -1.01 | 0.745 | Slc39a9 | solute carrier family 39 (zinc transporter), member 9 |
| 1448741 | at | -1.06 | 0.753 | Slc3a1 | solute carrier family 3, member 1 |
| 1425364 | a_at | 1.26 | 0.007 | Slc3a2 | solute carrier family 3 (activators of dibasic and neutral amino acid transport), member 2 |
| 1417061 | at | -2.02 | 0.000 | Slc40a1 | solute carrier family 40 (iron-regulated transporter), member 1 |
| 1460565 | at | -1.11 | 0.064 | Slc41a1 | solute carrier family 41, member 1 |
| 1425439 | a_at | -1.57 | 0.009 | Slc41a3 | solute carrier family 41, member 3 |
| 1439965 | at | 1.32 | 0.017 | Slc43a2 | solute carrier family 43, member 2 |
| 1422788 | at | -1.11 | 0.077 | Slc43a3 | solute carrier family 43, member 3 |
| 1437758 | a_at | 1.13 | 0.085 | Slc4a1ap | solute carrier family 4 (anion exchanger), member 1, adaptor protein |
| 1416637 | at | 1.06 | 0.575 | Slc4a2 | solute carrier family 4 (anion exchanger), member 2 |
| 1418485 | at | -1.56 | 0.000 | Slc4a3 | solute carrier family 4 (anion exchanger), member 3 |
| 1434096 | at | 1.04 | 0.719 | Slc4a4 | solute carrier family 4 (anion exchanger), member 4 |
| 1435484 | at | 1.14 | 0.442 | Slc5a3 | solute carrier family 5 (inositol transporters), member 3 |
| 1436137 | at | -2.10 | 0.000 | Slc6a17 | solute carrier family 6 (neurotransmitter transporter), member 17 |
| 1421346 | a_at | 1.23 | 0.118 | Slc6a6 | solute carrier family 6 (neurotransmitter transporter, taurine), member 6 |
| 1448596 | at | 1.19 | 0.053 | Slc6a8 | solute carrier family 6 (neurotransmitter transporter, creatine), member 8 |
| 1454992 | at | -1.33 | 0.001 | Slc7a1 | solute carrier family 7 (cationic amino acid transporter, y+ system), member 1 |

Supplementary table S1

| | | | | | |
|---|---|---|---|---|---|
| 1440506 | at | -1.20 | 0.446 | Slc7a2 | solute carrier family 7 (cationic amino acid transporter, y+ system), member 2 |
| 1433467 | at | 1.52 | 0.002 | Slc7a6 | solute carrier family 7 (cationic amino acid transporter, y+ system), member 6 |
| 1438115 | a_at | -1.10 | 0.381 | Slc9a3r1 | solute carrier family 9 (sodium/hydrogen exchanger), isoform 3 regulator 1 |
| 1435008 | at | -1.03 | 0.500 | Slc9a6 | solute carrier family 9 (sodium/hydrogen exchanger), isoform 6 |
| 1454777 | at | -1.20 | 0.103 | Slco2b1 | solute carrier organic anion transporter family, member 2b1 |
| 1440874 | at | -1.06 | 0.382 | Slco5a1 | solute carrier organic anion transporter family, member 5A1 |
| 1450165 | at | 1.48 | 0.018 | Slfn2 | schlafen 2 |
| 1421215 | a_at | 1.47 | 0.003 | Slmap | sarcolemma associated protein |
| 1448208 | at | 1.25 | 0.010 | Smad1 | MAD homolog 1 (Drosophila) |
| 1420634 | a_at | 1.02 | 0.750 | Smad2 | MAD homolog 2 (Drosophila) |
| 1454960 | at | -1.19 | 0.001 | Smad3 | MAD homolog 3 (Drosophila) |
| 1421047 | at | -1.10 | 0.438 | Smad5 | MAD homolog 5 (Drosophila) |
| 1423389 | at | -1.14 | 0.211 | Smad7 | MAD homolog 7 (Drosophila) |
| 1423956 | at | 1.36 | 0.000 | Smap1 | stromal membrane-associated protein 1 |
| 1450889 | at | 1.06 | 0.371 | Smarca3 | SWI/SNF related, matrix associated, actin dependent regulator of chromatin, subfamily a, member 3 |
| 1416620 | at | 1.13 | 0.238 | Smarcal1 | Swi/SNF related matrix associated, actin dependent regulator of chromatin, subfamily a-like 1 |
| 1416045 | a_at | 1.01 | 0.933 | Smarcb1 | SWI/SNF related, matrix associated, actin dependent regulator of chromatin, subfamily b, member 1 |
| 1448401 | at | 1.12 | 0.046 | Smarcd2 | SWI/SNF related, matrix associated, actin dependent regulator of chromatin, subfamily d, member 2 |
| 1418467 | at | 1.10 | 0.245 | Smarcd3 | SWI/SNF related, matrix associated, actin dependent regulator of chromatin, subfamily d, member 3 |
| 1422676 | at | -1.06 | 0.330 | Smarce1 | SWI/SNF related, matrix associated, actin dependent regulator of chromatin, subfamily e, member 1 |
| 1417831 | at | -1.12 | 0.244 | Smc1l1 | SMC (structural maintenance of chromosomes 1)-like 1 (S. cerevisiae) |
| 1458479 | at | 1.13 | 0.542 | Smc2l1 | SMC2 structural maintenance of chromosomes 2-like 1 (yeast) |
| 1427276 | at | 1.30 | 0.023 | Smc4l1 | SMC4 structural maintenance of chromosomes 4-like 1 (yeast) |
| 1428251 | at | 1.07 | 0.011 | Smchd1 | SMC hinge domain containing 1 |
| 1434072 | at | -1.05 | 0.280 | Smcr7 | Smith-Magenis syndrome chromosome region, candidate 7 homolog (human) |
| 1429017 | at | -1.02 | 0.779 | Smcr8 | Smith-Magenis syndrome chromosome region, candidate 8 homolog (human) |
| 1451259 | at | 1.17 | 0.070 | Smfn | small fragment nuclease |
| 1429043 | at | 1.10 | 0.358 | Smndc1 | survival motor neuron domain containing 1 |
| 1415935 | at | 1.47 | 0.000 | Smoc2 | SPARC related modular calcium binding 2 |
| 1446509 | at | -1.08 | 0.358 | Smox | spermine oxidase |
| 1448621 | a_at | -1.01 | 0.838 | Smpd1 | sphingomyelin phosphodiesterase 1, acid lysosomal |
| 1416999 | at | -1.24 | 0.091 | Smpd2 | sphingomyelin phosphodiesterase 2, neutral |
| 1416635 | at | -1.12 | 0.014 | Smpdl3a | sphingomyelin phosphodiesterase, acid-like 3A |
| 1418095 | at | 1.08 | 0.007 | Smpx | small muscle protein, X-linked |
| 1434190 | at | 1.14 | 0.354 | Sms | spermine synthase |
| 1452469 | a_at | -1.29 | 0.003 | Smtn | smoothelin |
| 1428763 | at | -1.08 | 0.030 | Smug1 | single-strand selective monofunctional uracil DNA glycosylase |
| 1428395 | at | 1.21 | 0.054 | Smurf1 | SMAD specific E3 ubiquitin protein ligase 1 |
| 1429045 | at | 1.21 | 0.091 | Smurf2 | SMAD specific E3 ubiquitin protein ligase 2 |
| 1450203 | at | -1.06 | 0.660 | Smyd1 | SET and MYND domain containing 1 |
| 1424760 | a_at | 1.12 | 0.035 | Smyd2 | SET and MYND domain containing 2 |

Supplementary table S1

| | | | | | |
|---|---|---|---|---|---|
| 1444670 | at | -1.36 | 0.084 | Smyd3 | SET and MYND domain containing 3 |
| 1423356 | at | -1.07 | 0.196 | Snap29 | synaptosomal-associated protein |
| 1416688 | at | 1.78 | 0.000 | Snap91 | synaptosomal-associated protein 91 |
| 1415756 | a_at | 1.07 | 0.076 | Snapap | SNAP-associated protein |
| 1429335 | at | 1.10 | 0.511 | Snapc1 | small nuclear RNA activating complex, polypeptide 1 |
| 1448726 | at | 1.07 | 0.363 | Snapc2 | small nuclear RNA activating complex, polypeptide 2 |
| 1436853 | a_at | -1.01 | 0.972 | Snca | synuclein, alpha |
| 1417788 | at | -1.16 | 0.022 | Sncg | synuclein, gamma |
| 1435603 | at | -1.76 | 0.001 | Sned1 | sushi, nidogen and EGF-like domains 1 |
| 1436338 | at | -1.30 | 0.004 | Snf1lk2 | SNF1-like kinase 2 |
| 1452789 | at | -1.24 | 0.016 | Snn | stannin |
| 1448864 | at | -1.17 | 0.174 | Snrk | SNF related kinase |
| 1416197 | at | 1.03 | 0.349 | Snrp1c | U1 small nuclear ribonucleoprotein 1C |
| 1429009 | at | -1.08 | 0.566 | Snrp70 | U1 small nuclear ribonucleoprotein polypeptide A |
| 1417274 | at | 1.08 | 0.419 | Snrpa | small nuclear ribonucleoprotein polypeptide A |
| 1417351 | a_at | 1.03 | 0.572 | Snrpa1 | small nuclear ribonucleoprotein polypeptide A' |
| 1419260 | a_at | -1.02 | 0.846 | Snrpb | small nuclear ribonucleoprotein B |
| 1426613 | a_at | 1.02 | 0.880 | Snrpb2 | U2 small nuclear ribonucleoprotein B |
| 1452680 | at | -1.25 | 0.000 | Snrpd2 | small nuclear ribonucleoprotein D2 |
| 1422885 | at | -1.18 | 0.066 | Snrpd3 | small nuclear ribonucleoprotein D3 |
| 1428672 | at | -1.23 | 0.106 | Snrpf | small nuclear ribonucleoprotein polypeptide F |
| 1448357 | at | -1.15 | 0.000 | Snrpg | small nuclear ribonucleoprotein polypeptide G |
| 1455087 | at | -1.42 | 0.142 | Snrpn | small nuclear ribonucleoprotein N |
| 1416825 | at | 1.02 | 0.835 | Snta1 | syntrophin, acidic 1 |
| 1416260 | a_at | 1.04 | 0.636 | Snx1 | sorting nexin 1 |
| 1431055 | a_at | 1.47 | 0.000 | Snx10 | sorting nexin 10 |
| 1454807 | a_at | 1.19 | 0.197 | Snx12 | sorting nexin 12 |
| 1456039 | at | -1.05 | 0.395 | Snx14 | sorting nexin 14 |
| 1424337 | at | 1.13 | 0.130 | Snx15 | sorting nexin 15 |
| 1456041 | at | 1.02 | 0.847 | Snx16 | sorting nexin 16 |
| 1427078 | at | -1.13 | 0.007 | Snx19 | sorting nexin 19 |
| 1460224 | at | 1.01 | 0.896 | Snx2 | sorting nexin 2 |
| 1428822 | a_at | 1.18 | 0.001 | Snx24 | sorting nexing 24 |
| 1433614 | at | -1.08 | 0.344 | Snx27 | sorting nexin family member 27 |
| 1422480 | at | 1.06 | 0.347 | Snx3 | sorting nexin 3 |
| 1460182 | at | 1.26 | 0.023 | Snx4 | sorting nexin 4 |
| 1428527 | at | 1.46 | 0.000 | Snx7 | sorting nexin 7 |
| 1454667 | at | 1.30 | 0.002 | Snx8 | sorting nexin 8 |
| 1455142 | at | 1.12 | 0.136 | Socs4 | suppressor of cytokine signaling 4 |
| 1423350 | at | 1.05 | 0.321 | Socs5 | suppressor of cytokine signaling 5 |
| 1435492 | at | 1.06 | 0.458 | Socs6 | suppressor of cytokine signaling 6 |

Supplementary table S1

| | | | | | |
|---|---|---|---|---|---|
| 1434834 | at | 1.16 | 0.126 | Socs7 | suppressor of cytokine signaling 7 |
| 1435304 | at | -1.03 | 0.511 | Sod1 | superoxide dismutase 1, soluble |
| 1417194 | at | -1.23 | 0.013 | Sod2 | superoxide dismutase 2, mitochondrial |
| 1417633 | at | 1.28 | 0.050 | Sod3 | superoxide dismutase 3, extracellular |
| 1434416 | a_at | -1.07 | 0.557 | Solh | small optic lobes homolog (Drosophila) |
| 1455634 | at | -1.20 | 0.423 | Son | Son cell proliferation protein |
| 1453003 | at | 1.49 | 0.000 | Sorl1 | sortilin-related receptor, LDLR class A repeats-containing |
| 1423362 | at | -1.27 | 0.001 | Sort1 | sortilin 1 |
| 1434389 | at | 1.01 | 0.938 | Sos1 | Son of sevenless homolog 1 (Drosophila) |
| 1452281 | at | -1.01 | 0.914 | Sos2 | Son of sevenless homolog 2 (Drosophila) |
| 1431225 | at | -1.45 | 0.147 | Sox11 | SRY-box containing gene 11 |
| 1450191 | a_at | 1.21 | 0.210 | Sox13 | SRY-box containing gene 13 |
| 1449135 | at | -1.08 | 0.277 | Sox18 | SRY-box containing gene 18 |
| 1433575 | at | -1.34 | 0.077 | Sox4 | SRY-box containing gene 4 |
| 1427677 | a_at | -1.49 | 0.024 | Sox6 | SRY-box containing gene 6 |
| 1416564 | at | -1.17 | 0.003 | Sox7 | SRY-box containing gene 7 |
| 1451538 | at | 1.35 | 0.010 | Sox9 | SRY-box containing gene 9 |
| 1454852 | at | -1.18 | 0.811 | Sp1 | trans-acting transcription factor 1 |
| 1447213 | at | -1.21 | 0.036 | Sp100 | nuclear antigen Sp100 |
| 1417635 | at | -1.38 | 0.006 | Spa17 | sperm autoantigenic protein 17 |
| 1433940 | at | -1.17 | 0.053 | Spag7 | sperm associated antigen 7 |
| 1433648 | at | 1.07 | 0.104 | Spag9 | sperm associated antigen 9 |
| 1448392 | at | 1.35 | 0.008 | Sparc | secreted acidic cysteine rich glycoprotein |
| 1416114 | at | 1.09 | 0.369 | Sparcl1 | SPARC-like 1 (mast9, hevin) |
| 1428516 | a_at | -1.36 | 0.000 | Spata11 | spermatogenesis associated 11 |
| 1434107 | at | -1.14 | 0.039 | Spata2 | spermatogenesis associated 2 |
| 1417254 | at | -1.04 | 0.450 | Spata5 | spermatogenesis associated 5 |
| 1437464 | at | -1.06 | 0.459 | Spata7 | spermatogenesis associated 7 |
| 1460677 | at | -1.05 | 0.556 | Spats2 | spermatogenesis associated, serine-rich 2 |
| 1416220 | at | 1.13 | 0.149 | Spcs1 | signal peptidase complex subunit 1 homolog (S. cerevisiae) |
| 1423215 | at | -1.11 | 0.477 | Spcs2 | signal peptidase complex subunit 2 homolog (S. cerevisiae) |
| 1451175 | at | 1.60 | 0.000 | Spcs3 | signal peptidase complex subunit 3 homolog (S. cerevisiae) |
| 1437481 | at | -1.30 | 0.021 | Spdy1 | speedy homolog 1 (Drosophila) |
| 1424210 | at | 1.26 | 0.002 | Spfh1 | SPFH domain family, member 1 |
| 1435223 | at | 1.19 | 0.006 | Spfh2 | SPFH domain family, member 2 |
| 1451520 | at | 1.02 | 0.887 | Spg20 | spastic paraplegia 20, spartin (Troyer syndrome) homolog (human) |
| 1460400 | at | 1.23 | 0.000 | Spg4 | spastic paraplegia 4 homolog (human) |
| 1427166 | a_at | -1.20 | 0.003 | Spg7 | spastic paraplegia 7 homolog (human) |
| 1426230 | at | -1.09 | 0.821 | Sphk2 | sphingosine kinase 2 |
| 1435912 | at | -1.14 | 0.162 | Spink3 | serine protease inhibitor, Kazal type 3 |
| 1451935 | a_at | -1.44 | 0.000 | Spint2 | serine protease inhibitor, Kunitz type 2 |

Supplementary table S1

| | | | | | |
|---|---|---|---|---|---|
| 1446117 | at | -1.14 | 0.524 | Spire1 | spire homolog 1 (Drosophila) |
| 1427888 | a_at | 1.10 | 0.592 | Spna2 | spectrin alpha 2 |
| 1455226 | at | -1.11 | 0.018 | Spnb1 | spectrin beta 1 |
| 1451342 | at | -1.16 | 0.043 | Spon1 | spondin 1, (f-spondin) extracellular matrix protein |
| 1416525 | at | -1.15 | 0.375 | Spop | speckle-type POZ protein |
| 1451073 | at | -1.17 | 0.007 | Sppl3 | signal peptide peptidase 3 |
| 1432372 | a_at | -1.19 | 0.003 | Spr | sepiapterin reductase |
| 1445228 | at | 1.17 | 0.633 | Spred1 | sprouty protein with EVH-1 domain 1, related sequence |
| 1434403 | at | 1.13 | 0.200 | Spred2 | sprouty-related, EVH1 domain containing 2 |
| 1415874 | at | 1.46 | 0.013 | Spry1 | sprouty homolog 1 (Drosophila) |
| 1436584 | at | 1.07 | 0.236 | Spry2 | sprouty homolog 2 (Drosophila) |
| 1440867 | at | 1.30 | 0.015 | Spry4 | sprouty homolog 4 (Drosophila) |
| 1422690 | at | 1.19 | 0.004 | Sptlc1 | serine palmitoyltransferase, long chain base subunit 1 |
| 1435937 | at | 1.21 | 0.400 | Sptlc2 | serine palmitoyltransferase, long chain base subunit 2 |
| 1420641 | a_at | -1.28 | 0.000 | Sqrdl | sulfide quinone reductase-like (yeast) |
| 1450957 | a_at | 1.15 | 0.033 | Sqstm1 | sequestosome 1 |
| 1437423 | a_at | 1.03 | 0.301 | Sra1 | steroid receptor RNA activator 1 |
| 1451117 | a_at | 1.09 | 0.205 | Srcasm | Src activating and signaling molecule |
| 1437482 | at | 1.14 | 0.031 | Srd5a2l2 | steroid 5 alpha-reductase 2-like 2 |
| 1426690 | a_at | -1.01 | 0.939 | Srebf1 | sterol regulatory element binding factor 1 |
| 1452174 | at | -1.02 | 0.827 | Srebf2 | sterol regulatory element binding factor 2 |
| 1418256 | at | 1.21 | 0.189 | Srf | serum response factor |
| 1420510 | at | 1.05 | 0.846 | Srfbp1 | serum response factor binding protein 1 |
| 1450878 | at | 1.18 | 0.031 | Sri | sorcin |
| 1436867 | at | 1.08 | 0.001 | Srl | sarcalumenin |
| 1421260 | a_at | 1.26 | 0.041 | Srm | spermidine synthase |
| 1418567 | a_at | 1.02 | 0.716 | Srp14 | signal recognition particle 14 |
| 1450891 | at | 1.02 | 0.634 | Srp19 | signal recognition particle 19 |
| 1433708 | at | -1.05 | 0.455 | Srp68 | signal recognition particle 68 |
| 1423670 | a_at | -1.15 | 0.007 | Srpr | signal recognition particle receptor ('docking protein') |
| 1421012 | at | -1.17 | 0.233 | Srprb | signal recognition particle receptor, B subunit |
| 1454689 | at | -1.17 | 0.444 | Srrm1 | serine/arginine repetitive matrix 1 |
| 1437638 | at | -1.17 | 0.122 | Srrm2 | serine/arginine repetitive matrix 2 |
| 1436911 | at | -1.15 | 0.044 | Ss18l1 | synovial sarcoma translocation gene on chromosome 18-like 1 |
| 1436535 | at | 1.19 | 0.522 | Ssa2 | Sjogren syndrome antigen A2 |
| 1451418 | a_at | 2.27 | 0.000 | Ssb4 | SPRY domain-containing SOCS box 4 |
| 1427965 | at | 1.05 | 0.246 | Ssbp1 | single-stranded DNA binding protein 1 |
| 1429951 | at | -1.13 | 0.067 | Ssbp2 | single-stranded DNA binding protein 2 |
| 1425940 | a_at | -1.29 | 0.138 | Ssbp3 | single-stranded DNA binding protein 3 |
| 1423613 | at | 1.23 | 0.013 | Ssfa2 | sperm specific antigen 2 |
| 1455854 | a_at | 1.11 | 0.274 | Ssh1 | slingshot homolog 1 (Drosophila) |

Supplementary table S1

| | | | | | |
|---|---|---|---|---|---|
| 1455078 | at | -1.13 | 0.162 | Ssh2 | slingshot homolog 2 (Drosophila) |
| 1448643 | at | -1.01 | 0.777 | Ssna1 | Sjogren's syndrome nuclear autoantigen 1 |
| 1417763 | at | -1.07 | 0.073 | Ssr1 | signal sequence receptor, alpha |
| 1449930 | a_at | 1.04 | 0.486 | Ssr2 | signal sequence receptor, beta |
| 1415700 | a_at | -1.01 | 0.820 | Ssr3 | signal sequence receptor, gamma |
| 1417726 | at | -1.16 | 0.135 | Sssca1 | Sjogren's syndrome/scleroderma autoantigen 1 homolog (human) |
| 1449467 | at | 1.05 | 0.862 | Ssty1 | spermiogenesis specific transcript on the Y 1 |
| 1417838 | at | -1.01 | 0.962 | Ssty2 | spermiogenesis specific transcript on the Y 2 |
| 1448743 | at | 1.31 | 0.003 | Ssx2ip | synovial sarcoma, X breakpoint 2 interacting protein |
| 1450406 | a_at | -1.06 | 0.090 | St3gal3 | ST3 beta-galactoside alpha-2,3-sialyltransferase 3 |
| 1449198 | a_at | -1.03 | 0.598 | St3gal5 | ST3 beta-galactoside alpha-2,3-sialyltransferase 5 |
| 1449078 | at | 1.07 | 0.345 | St3gal6 | ST3 beta-galactoside alpha-2,3-sialyltransferase 6 |
| 1428372 | at | 1.18 | 0.247 | St5 | suppression of tumorigenicity 5 |
| 1420928 | at | 1.30 | 0.004 | St6gal1 | beta galactoside alpha 2,6 sialyltransferase 1 |
| 1417616 | at | 1.22 | 0.055 | St6galnac2 | ST6 (alpha-N-acetyl-neuraminyl-2,3-beta-galactosyl-1,3)-N-acetylgalactosaminide alpha-2,6-sialyltransferase 2 |
| 1457583 | at | -1.31 | 0.086 | St6galnac3 | ST6 (alpha-N-acetyl-neuraminyl-2,3-beta-galactosyl-1,3)-N-acetylgalactosaminide alpha-2,6-sialyltransferase 3 |
| 1418075 | at | 1.41 | 0.000 | St6galnac4 | ST6 (alpha-N-acetyl-neuraminyl-2,3-beta-galactosyl-1,3)-N-acetylgalactosaminide alpha-2,6-sialyltransferase 4 |
| 1423115 | at | -1.27 | 0.021 | St6galnac6 | ST6 (alpha-N-acetyl-neuraminyl-2,3-beta-galactosyl-1,3)-N-acetylgalactosaminide alpha-2,6-sialyltransferase 6 |
| 1418967 | a_at | 1.17 | 0.231 | St7 | Suppression of tumorigenicity 7 |
| 1427916 | at | 1.03 | 0.562 | St7l | suppression of tumorigenicity 7-like |
| 1419186 | a_at | 1.07 | 0.461 | St8sia4 | ST8 alpha-N-acetyl-neuraminide alpha-2,8-sialyltransferase 4 |
| 1450199 | a_at | 1.15 | 0.103 | Stab1 | stabilin 1 |
| 1421849 | at | 1.09 | 0.039 | Stag2 | stromal antigen 2 |
| 1416861 | at | -1.00 | 0.993 | Stam | signal transducing adaptor molecule (SH3 domain and ITAM motif) 1 |
| 1416976 | at | 1.19 | 0.784 | Stam2 | signal transducing adaptor molecule (SH3 domain and ITAM motif) 2 |
| 1448956 | at | -2.21 | 0.000 | Stard10 | START domain containing 10 |
| 1452604 | at | -1.13 | 0.051 | Stard13 | serologically defined colon cancer antigen 13 |
| 1430274 | a_at | 1.07 | 0.343 | Stard3nl | STARD3 N-terminal like |
| 1429240 | at | 1.04 | 0.902 | Stard4 | StAR-related lipid transfer (START) domain containing 4 |
| 1426317 | at | 1.12 | 0.305 | Stard6 | StAR-related lipid transfer (START) domain containing 6 |
| 1427072 | at | -1.08 | 0.326 | Stard8 | START domain containing 8 |
| 1450034 | at | -1.10 | 0.314 | Stat1 | signal transducer and activator of transcription 1 |
| 1460700 | at | 1.51 | 0.000 | Stat3 | signal transducer and activator of transcription 3 |
| 1421469 | a_at | -1.15 | 0.235 | Stat5a | signal transducer and activator of transcription 5A |
| 1422103 | a_at | -1.17 | 0.047 | Stat5b | signal transducer and activator of transcription 5B |
| 1426353 | at | 1.07 | 0.323 | Stat6 | signal transducer and activator of transcription 6 |
| 1415774 | at | -1.03 | 0.579 | Statip1 | signal transducer and activator of transcription interacting protein 1 |
| 1445186 | at | -1.50 | 0.008 | Stc2 | stanniocalcin 2 |
| 1433772 | at | 1.03 | 0.630 | Stch | stress 70 protein chaperone, microsome-associated, human homolog |
| 1434219 | at | 1.14 | 0.065 | Stim2 | stromal interaction molecule 2 |
| 1415909 | at | 1.01 | 0.943 | Stip1 | stress-induced phosphoprotein 1 |

Supplementary table S1

| | | | | | |
|---|---|---|---|---|---|
| 1417751 | at | 1.08 | 0.319 | Stk10 | serine/threonine kinase 10 |
| 1418644 | a_at | -1.07 | 0.299 | Stk11 | serine/threonine kinase 11 |
| 1426160 | a_at | -1.06 | 0.239 | Stk16 | serine/threonine kinase 16 |
| 1450997 | at | 1.11 | 0.352 | Stk17b | serine/threonine kinase 17b (apoptosis-inducing) |
| 1426247 | at | -1.20 | 0.044 | Stk24 | serine/threonine kinase 24 (STE20 homolog, yeast) |
| 1416770 | at | -1.03 | 0.648 | Stk25 | serine/threonine kinase 25 (yeast) |
| 1418512 | at | 1.18 | 0.036 | Stk3 | serine/threonine kinase 3 (Ste20, yeast homolog) |
| 1416252 | at | 1.22 | 0.004 | Stk38 | serine/threonine kinase 38 |
| 1431434 | at | -1.68 | 0.016 | Stk39 | serine/threonine kinase 39, STE20/SPS1 homolog (yeast) |
| 1434159 | at | 1.08 | 0.309 | Stk4 | serine/threonine kinase 4 |
| 1448113 | at | 1.00 | 0.999 | Stmn1 | stathmin 1 |
| 1423281 | at | 1.04 | 0.739 | Stmn2 | stathmin-like 2 |
| 1438910 | a_at | -1.22 | 0.003 | Stom | stomatin |
| 1448774 | at | -1.03 | 0.643 | Stoml2 | stomatin (Epb7.2)-like 2 |
| 1416553 | at | -1.11 | 0.043 | Stra13 | stimulated by retinoic acid 13 |
| 1416566 | at | 1.10 | 0.159 | Strap | serine/threonine kinase receptor associated protein |
| 1455022 | at | 1.13 | 0.956 | Strn | striatin, calmodulin binding protein |
| 1426209 | at | 1.10 | 0.100 | Strn4 | striatin, calmodulin binding protein 4 |
| 1421500 | at | -1.09 | 0.444 | Sts | steroid sulfatase |
| 1416580 | a_at | -1.13 | 0.040 | Stub1 | STIP1 homology and U-Box containing protein 1 |
| 1448936 | at | 1.19 | 0.037 | Stx12 | syntaxin 12 |
| 1429237 | at | 1.04 | 0.382 | Stx16 | syntaxin 16 |
| 1431295 | a_at | -1.02 | 0.704 | Stx18 | syntaxin 18 |
| 1451573 | a_at | 1.09 | 0.112 | Stx4a | syntaxin 4A (placental) |
| 1423038 | at | 1.20 | 0.042 | Stx6 | syntaxin 6 |
| 1418436 | at | 1.01 | 0.775 | Stx7 | syntaxin 7 |
| 1418088 | a_at | 1.19 | 0.120 | Stx8 | syntaxin 8 |
| 1420505 | a_at | 1.18 | 0.157 | Stxbp1 | syntaxin binding protein 1 |
| 1416653 | at | 1.05 | 0.583 | Stxbp3 | syntaxin binding protein 3 |
| 1435396 | at | -1.03 | 0.594 | Stxbp6 | syntaxin binding protein 6 (amisyn) |
| 1421664 | a_at | -1.01 | 0.858 | Styx | phosphoserine/threonine/tyrosine interaction protein |
| 1452206 | at | -1.01 | 0.691 | Sucla2 | succinate-Coenzyme A ligase, ADP-forming, beta subunit |
| 1415891 | at | -1.09 | 0.031 | Suclg1 | succinate-CoA ligase, GDP-forming, alpha subunit |
| 1427441 | a_at | -1.27 | 0.000 | Suclg2 | succinate-Coenzyme A ligase, GDP-forming, beta subunit |
| 1460564 | at | -1.20 | 0.030 | Suhw2 | suppressor of hairy wing homolog 2 (Drosophila) |
| 1427983 | at | 1.01 | 0.964 | Suhw3 | suppressor of hairy wing homolog 3 (Drosophila) |
| 1426556 | at | -1.08 | 0.161 | Suhw4 | suppressor of hairy wing homolog 4 (Drosophila) |
| 1438200 | at | 1.97 | 0.000 | Sulf1 | sulfatase 1 |
| 1430388 | a_at | 1.10 | 0.228 | Sulf2 | sulfatase 2 |
| 1427345 | a_at | -1.52 | 0.001 | Sult1a1 | sulfotransferase family 1A, phenol-preferring, member 1 |
| 1424603 | at | 1.17 | 0.132 | Sumf1 | sulfatase modifying factor 1 |

Supplementary table S1

| | | | | | |
|---|---|---|---|---|---|
| 1438289 | a_at | 1.05 | 0.448 | Sumo1 | SMT3 suppressor of mif two 3 homolog 1 (yeast) |
| 1419741 | at | 1.13 | 0.213 | Supt16h | suppressor of Ty 16 homolog (S. cerevisiae) |
| 1434963 | at | 1.13 | 0.167 | Supt3h | suppressor of Ty 3 homolog (S. cerevisiae) |
| 1424255 | at | 1.06 | 0.372 | Supt5h | suppressor of Ty 5 homolog (S. cerevisiae) |
| 1417628 | at | -1.00 | 0.945 | Supt6h | suppressor of Ty 6 homolog (S. cerevisiae) |
| 1460557 | at | 1.08 | 0.417 | Supv3l1 | suppressor of var1, 3-like 1 (S. cerevisiae) |
| 1425806 | a_at | 1.07 | 0.092 | Surb7 | SRB7 (supressor of RNA polymerase B) homolog (S. cerevisiae) |
| 1450561 | a_at | -1.17 | 0.006 | Surf1 | surfeit gene 1 |
| 1436797 | a_at | 1.03 | 0.877 | Surf4 | surfeit gene 4 |
| 1421528 | a_at | -1.05 | 0.471 | Surf5 | surfeit gene 5 |
| 1416864 | at | 1.16 | 0.069 | Surf6 | surfeit gene 6 |
| 1434999 | at | 1.03 | 0.495 | Suv420h1 | suppressor of variegation 4-20 homolog 1 (Drosophila) |
| 1449661 | at | -1.08 | 0.841 | Suz12 | suppressor of zeste 12 homolog (Drosophila) |
| 1460735 | at | 1.29 | 0.001 | Svil | supervillin |
| 1434225 | at | 1.19 | 0.032 | Swap70 | SWA-70 protein |
| 1416472 | at | 1.06 | 0.044 | Syap1 | synapse associated protein 1 |
| 1452007 | at | 1.11 | 0.624 | Sybl1 | synaptobrevin like 1 |
| 1449534 | at | 1.05 | 0.736 | Sycp3 | synaptonemal complex protein 3 |
| 1418261 | at | 1.42 | 0.003 | Syk | spleen tyrosine kinase |
| 1428673 | at | -1.01 | 0.881 | Sympk | symplekin |
| 1422118 | at | -1.07 | 0.374 | Sync | syncoilin |
| 1428546 | at | 1.20 | 0.082 | Syncrip | synaptotagmin binding, cytoplasmic RNA interacting protein |
| 1421545 | a_at | -1.30 | 0.004 | Syne1 | synaptic nuclear envelope 1 |
| 1419289 | a_at | 1.09 | 0.200 | Syngr1 | synaptogyrin 1 |
| 1417081 | a_at | -1.13 | 0.019 | Syngr2 | synaptogyrin 2 |
| 1427685 | a_at | -1.14 | 0.535 | Synj2 | synaptojanin 2 |
| 1427045 | at | 2.14 | 0.005 | Synpo | synaptopodin |
| 1452879 | at | 1.08 | 0.132 | Synpo2 | synaptopodin 2 |
| 1428295 | at | 3.27 | 0.000 | Synpo2l | synaptopodin 2-like |
| 1444331 | at | -1.29 | 0.479 | Synpr | synaptoporin |
| 1460081 | at | -1.44 | 0.013 | Syt7 | synaptotagmin 7 |
| 1416783 | at | -1.03 | 0.704 | Tac1 | tachykinin 1 |
| 1454909 | at | 1.36 | 0.658 | Tacc1 | transforming, acidic coiled-coil containing protein 1 |
| 1425745 | a_at | 1.05 | 0.242 | Tacc2 | transforming, acidic coiled-coil containing protein 2 |
| 1452310 | at | -1.14 | 0.141 | Tada2l | transcriptional adaptor 2 (ADA2 homolog, yeast)-like |
| 1417467 | a_at | -1.10 | 0.267 | Tada3l | transcriptional adaptor 3 (NGG1 homolog, yeast)-like |
| 1448784 | at | -1.11 | 0.476 | Taf10 | TAF10 RNA polymerase II, TATA box binding protein (TBP)-associated factor |
| 1451995 | at | 1.12 | 0.232 | Taf11 | TAF11 RNA polymerase II, TATA box binding protein (TBP)-associated factor |
| 1423398 | at | -1.14 | 0.045 | Taf12 | TAF12 RNA polymerase II, TATA box binding protein (TBP)-associated factor |
| 1448707 | at | 1.15 | 0.000 | Taf13 | TAF13 RNA polymerase II, TATA box binding protein (TBP)-associated factor |
| 1419702 | at | -1.05 | 0.553 | Taf1a | TATA box binding protein (Tbp)-associated factor, RNA polymerase I, A |

Supplementary table S1

| | | | | | |
|---|---|---|---|---|---|
| 1431345 | a_at | -1.09 | 0.349 | Taf1b | TATA box binding protein (Tbp)-associated factor, RNA polymerase I, B |
| 1435474 | at | -1.10 | 0.091 | Taf5 | TAF5 RNA polymerase II, TATA box binding protein (TBP)-associated factor |
| 1448195 | at | 1.05 | 0.267 | Taf5l | TAF5-like RNA polymerase II, p300/CBP-associated factor (PCAF)-associated factor |
| 1418593 | at | -1.14 | 0.341 | Taf6 | TAF6 RNA polymerase II, TATA box binding protein (TBP)-associated factor |
| 1451509 | at | 1.07 | 0.736 | Taf9 | TAF9 RNA polymerase II, TATA box binding protein (TBP)-associated factor |
| 1460453 | at | -1.02 | 0.634 | Tagap1 | T-cell activation GTPase activating protein 1 |
| 1423505 | at | 1.15 | 0.285 | Tagln | transgelin |
| 1426529 | a_at | 1.30 | 0.061 | Tagln2 | transgelin 2 |
| 1449389 | at | 1.22 | 0.106 | Tal1 | T-cell acute lymphocytic leukemia 1 |
| 1425129 | a_at | -1.06 | 0.369 | Taldo1 | transaldolase 1 |
| 1421640 | a_at | 1.03 | 0.775 | Tank | TRAF family member-associated Nf-kappa B activator |
| 1438208 | at | 1.09 | 0.625 | Taok2 | TAO kinase 2 |
| 1416016 | at | 1.08 | 0.700 | Tap1 | transporter 1, ATP-binding cassette, sub-family B (MDR/TAP) |
| 1421812 | at | -1.14 | 0.030 | Tapbp | TAP binding protein |
| 1423616 | at | 1.03 | 0.602 | Tarbp2 | TAR (HIV) RNA binding protein 2 |
| 1460323 | at | 1.30 | 0.000 | Tars | threonyl-tRNA synthetase |
| 1431125 | a_at | -1.15 | 0.107 | Tarsl1 | threonyl-tRNA synthetase-like 1 |
| 1434738 | at | -1.17 | 0.039 | Tarsl2 | threonyl-tRNA synthetase-like 2 |
| 1448033 | at | 1.22 | 0.044 | Tatdn1 | TatD DNase domain containing 1 |
| 1454769 | at | 1.19 | 0.010 | Tatdn2 | TatD DNase domain containing 2 |
| 1448399 | at | 1.03 | 0.246 | Tax1bp1 | Tax1 (human T-cell leukemia virus type I) binding protein 1 |
| 1424169 | at | 1.43 | 0.044 | Tax1bp3 | Tax1 (human T-cell leukemia virus type I) binding protein 3 |
| 1416207 | at | 1.06 | 0.339 | Taz | tafazzin |
| 1454850 | at | -3.01 | 0.000 | Tbc1d10c | TBC1 domain family, member 10c |
| 1435675 | at | 1.12 | 0.100 | Tbc1d12 | TBC1D12: TBC1 domain family, member 12 |
| 1449772 | at | 1.11 | 0.119 | Tbc1d14 | TBC1 domain family, member 14 |
| 1416061 | at | 1.12 | 0.641 | Tbc1d15 | TBC1 domain family, member 15 |
| 1435763 | at | -1.51 | 0.001 | Tbc1d16 | TBC1 domain family, member 16 |
| 1423659 | a_at | -1.05 | 0.615 | Tbc1d17 | TBC1 domain family, member 17 |
| 1424918 | at | -1.05 | 0.229 | Tbc1d19 | TBC1 domain family, member 19 |
| 1426644 | at | 1.02 | 0.749 | Tbc1d20 | TBC1 domain family, member 20 |
| 1451107 | at | 1.09 | 0.443 | Tbc1d22a | TBC1 domain family, member 22a |
| 1436053 | at | -1.26 | 0.022 | Tbc1d22b | TBC1 domain family, member 22B |
| 1455903 | at | -1.69 | 0.001 | Tbc1d4 | TBC1 domain family, member 4 |
| 1428598 | at | 1.00 | 0.983 | Tbc1d7 | TBC1 domain family, member 7 |
| 1416996 | at | -1.13 | 0.049 | Tbc1d8 | TBC1 domain family, member 8 |
| 1417652 | a_at | 1.17 | 0.012 | Tbca | tubulin cofactor a |
| 1424644 | at | 1.13 | 0.250 | Tbcc | tubulin-specific chaperone c |
| 1452194 | at | 1.08 | 0.593 | Tbcd | tubulin-specific chaperone d |
| 1428282 | at | -1.00 | 0.990 | Tbce | tubulin-specific chaperone e |
| 1422469 | at | 1.16 | 0.001 | Tbk1 | TANK-binding kinase 1 |

Supplementary table S1

| | | | | | |
|---|---|---|---|---|---|
| 1455042 | at | -1.22 | 0.454 | Tbl1x | transducin (beta)-like 1 X-linked |
| 1436603 | at | 1.03 | 0.596 | Tbl2 | transducin (beta)-like 2 |
| 1415750 | at | -1.11 | 0.128 | Tbl3 | transducin (beta)-like 3 |
| 1426469 | a_at | -1.09 | 0.386 | Tbp | TATA box binding protein |
| 1452648 | at | 1.21 | 0.000 | Tbrg1 | transforming growth factor beta regulated gene 1 |
| 1448795 | a_at | -1.06 | 0.582 | Tbrg4 | transforming growth factor beta regulated gene 4 |
| 1425158 | at | 1.14 | 0.067 | Tbx20 | T-box 20 |
| 1448029 | at | -1.52 | 0.001 | Tbx3 | T-box 3 |
| 1425695 | at | -2.58 | 0.000 | Tbx5 | T-box 5 |
| 1423145 | a_at | -1.11 | 0.117 | Tcap | titin-cap |
| 1417891 | at | -1.00 | 0.937 | Tce1 | T-complex expressed gene 1 |
| 1419258 | at | -1.14 | 0.025 | Tcea1 | transcription elongation factor A (SII) 1 |
| 1424531 | a_at | -1.50 | 0.000 | Tcea3 | transcription elongation factor A (SII), 3 |
| 1424634 | at | -1.10 | 0.449 | Tceal1 | transcription elongation factor A (SII)-like 1 |
| 1418171 | at | 1.28 | 0.024 | Tceal8 | transcription elongation factor A (SII)-like 8 |
| 1428263 | a_at | -1.04 | 0.573 | Tceb2 | transcription elongation factor B (SIII), polypeptide 2 |
| 1422591 | at | 1.08 | 0.251 | Tceb3 | transcription elongation factor B (SIII), polypeptide 3 |
| 1449592 | at | -1.10 | 0.357 | Tcf15 | transcription factor 15 |
| 1421910 | at | -1.17 | 0.086 | Tcf20 | transcription factor 20 |
| 1417447 | at | 1.04 | 0.401 | Tcf21 | transcription factor 21 |
| 1436207 | at | -1.17 | 0.252 | Tcf3 | transcription factor 3 |
| 1429428 | at | -1.33 | 0.028 | Tcf7l2 | transcription factor 7-like 2, T-cell specific, HMG-box |
| 1427046 | at | -2.56 | 0.000 | Tcfcp2l3 | transcription factor CP2-like 3 |
| 1434318 | a_at | -1.00 | 0.960 | Tcfe3 | transcription factor E3 |
| 1418284 | at | -1.16 | 0.125 | Tcfl1 | transcription factor-like 1 |
| 1418437 | a_at | -1.03 | 0.462 | Tcfl4 | transcription factor-like 4 |
| 1420635 | a_at | 1.13 | 0.024 | Tcirg1 | T-cell, immune regulator 1 |
| 1448200 | at | -1.20 | 0.002 | Tcn2 | transcobalamin 2 |
| 1448122 | at | -1.00 | 0.967 | Tcp1 | t-complex protein 1 |
| 1424068 | at | 1.12 | 0.101 | Tcta | T-cell leukemia translocation altered gene |
| 1449929 | at | 1.07 | 0.143 | Tcte1l | t-complex-associated-testis-expressed 1-like |
| 1416790 | a_at | 1.13 | 0.204 | Tdg | thymine DNA glycosylase |
| 1426716 | at | 1.08 | 0.245 | Tdrd7 | tudor domain containing 7 |
| 1423434 | at | 1.63 | 0.000 | Tead1 | TEA domain family member 1 |
| 1460221 | at | -1.06 | 0.257 | Tebp | telomerase binding protein, p23 |
| 1460204 | at | 1.01 | 0.891 | Tec | cytoplasmic tyrosine kinase, Dscr28C related (Drosophila) |
| 1424175 | at | 1.05 | 0.729 | Tef | thyrotroph embryonic factor |
| 1451586 | at | -1.07 | 0.149 | Tegt | testis enhanced gene transcript |
| 1418788 | at | -1.03 | 0.692 | Tek | endothelial-specific receptor tyrosine kinase |
| 1452264 | at | -1.16 | 0.052 | Tenc1 | tensin like C1 domain-containing phosphatase |
| 1418196 | at | -1.04 | 0.533 | Tep1 | telomerase associated protein 1 |

Supplementary table S1

| | | | | | |
|---|---|---|---|---|---|
| 1448126 | at | 1.16 | 0.088 | Tera | teratocarcinoma expressed, serine rich |
| 1421147 | at | -1.09 | 0.160 | Terf2 | telomeric repeat binding factor 2 |
| 1426049 | a_at | -1.17 | 0.009 | Terf2ip | telomeric repeat binding factor 2, interacting protein |
| 1455054 | a_at | 0.58 | 0.066 | Tes3 | testis derived transcript 3 |
| 1418743 | a_at | -1.26 | 0.000 | Tesc | tescalcin |
| 1450662 | at | -1.12 | 0.218 | Tesk1 | testis specific protein kinase 1 |
| 1452213 | at | 1.07 | 0.081 | Tex2 | testis expressed gene 2 |
| 1425068 | a_at | 1.05 | 0.300 | Tex264 | testis expressed gene 264 |
| 1423884 | at | 1.11 | 0.085 | Tex292 | testis expressed gene 292 |
| 1456215 | at | 1.11 | 0.238 | Tfam | transcription factor A, mitochondrial |
| 1448224 | at | 1.08 | 0.484 | Tfam | transcription factor A, mitochondrial |
| 1435924 | at | -1.08 | 0.191 | Tfb1m | transcription factor B1, mitochondrial |
| 1423441 | at | -1.05 | 0.364 | Tfb2m | transcription factor B2, mitochondrial |
| 1423241 | a_at | -1.02 | 0.654 | Tfdp1 | transcription factor Dp 1 |
| 1437174 | at | -1.35 | 0.049 | Tfdp2 | transcription factor Dp 2 |
| 1415887 | at | -1.07 | 0.057 | Tfg | Trk-fused gene |
| 1448976 | at | -1.09 | 0.314 | Tfip11 | tuftelin interacting protein 11 |
| 1438530 | at | -1.28 | 0.559 | Tfpi | tissue factor pathway inhibitor |
| 1418064 | at | -1.30 | 0.007 | Tfpt | TCF3 (E2A) fusion partner |
| 1422966 | a_at | 1.45 | 0.004 | Tfrc | transferrin receptor |
| 1424526 | a_at | 1.07 | 0.450 | Tgds | TDP-glucose 4,6-dehydratase |
| 1418136 | at | 1.29 | 0.059 | Tgfb1i1 | transforming growth factor beta 1 induced transcript 1 |
| 1450923 | at | 2.72 | 0.000 | Tgfb2 | transforming growth factor, beta 2 |
| 1415871 | at | 1.49 | 0.003 | Tgfbi | transforming growth factor, beta induced |
| 1420895 | at | 1.40 | 0.001 | Tgfbr1 | transforming growth factor, beta receptor I |
| 1426397 | at | 1.24 | 0.001 | Tgfbr2 | transforming growth factor, beta receptor II |
| 1433795 | at | -1.57 | 0.007 | Tgfbr3 | transforming growth factor, beta receptor III |
| 1426004 | a_at | 1.38 | 0.051 | Tgm2 | transglutaminase 2, C polypeptide |
| 1438341 | at | 1.16 | 0.288 | Tgm4 | transglutaminase 4 (prostate) |
| 1423309 | at | 1.03 | 0.775 | Tgoln2 | trans-golgi network protein 2 |
| 1449009 | at | -1.07 | 0.776 | Tgtp | T-cell specific GTPase |
| 1416198 | at | 1.03 | 0.470 | Th1l | TH1-like homolog (Drosophila) |
| 1455003 | at | 1.11 | 0.033 | Thap1 | THAP domain containing, apoptosis associated protein 1 |
| 1416428 | at | -1.10 | 0.089 | Thap11 | THAP domain containing 11 |
| 1428521 | at | -1.02 | 0.806 | Thap3 | THAP domain containing, apoptosis associated protein 3 |
| 1424052 | at | -1.35 | 0.000 | Thap4 | THAP domain containing 4 |
| 1452069 | a_at | 1.07 | 0.257 | Thap7 | THAP domain containing 7 |
| 1448529 | at | 1.06 | 0.245 | Thbd | thrombomodulin |
| 1421811 | at | 2.90 | 0.000 | Thbs1 | thrombospondin 1 |
| 1450663 | at | -1.39 | 0.009 | Thbs2 | thrombospondin 2 |
| 1449388 | at | 6.04 | 0.000 | Thbs4 | thrombospondin 4 |

Supplementary table S1

| | | | | | |
|---|---|---|---|---|---|
| 1429267 | at | -1.11 | 0.177 | Thea | thioesterase, adipose associated |
| 1417316 | at | 1.01 | 0.822 | Them2 | thioesterase superfamily member 2 |
| 1452605 | at | -1.17 | 0.060 | Thnsl1 | threonine synthase-like 1 (bacterial) |
| 1455829 | at | 1.20 | 0.136 | Thoc1 | THO complex 1 |
| 1423970 | at | 1.03 | 0.513 | Thoc3 | THO complex 3 |
| 1436211 | at | -1.05 | 0.482 | Thoc4 | THO complex 4 |
| 1454675 | at | -1.33 | 0.063 | Thra | thyroid hormone receptor alpha |
| 1434603 | at | -1.18 | 0.044 | Thrap2 | thyroid hormone receptor associated protein 2 |
| 1460545 | at | -1.19 | 0.110 | Thrap3 | thyroid hormone receptor associated protein 3 |
| 1424180 | a_at | 1.00 | 0.970 | Thrap4 | thyroid hormone receptor associated protein 4 |
| 1438416 | at | -1.01 | 0.905 | Thrap5 | thyroid hormone receptor associated protein 5 |
| 1448917 | at | 1.07 | 0.191 | Thrap6 | thyroid hormone receptor associated protein 6 |
| 1424737 | at | -1.11 | 0.201 | Thrsp | thyroid hormone responsive SPOT14 homolog (Rattus) |
| 1438266 | at | 1.39 | 0.001 | Thsd6 | thrombospondin, type I domain containing 6 |
| 1436007 | a_at | 1.10 | 0.375 | Thumpd1 | THUMP domain containing 1 |
| 1417684 | at | 1.22 | 0.001 | Thumpd3 | THUMP domain containing 3 |
| 1416814 | at | -1.16 | 0.165 | Tia1 | cytotoxic granule-associated RNA binding protein 1 |
| 1451422 | at | 1.01 | 0.899 | Tiaf1 | TGF-beta1-induced anti-apoptotic factor 1 |
| 1420146 | at | -1.09 | 0.229 | Tiam2 | T-cell lymphoma invasion and metastasis 2 |
| 1416238 | at | -1.00 | 0.986 | Tie1 | tyrosine kinase receptor 1 |
| 1416029 | at | 1.26 | 0.026 | Tieg1 | TGFB inducible early growth response 1 |
| 1437241 | at | -1.09 | 0.363 | Tieg3 | TGFB inducible early growth response 3 |
| 1429194 | at | -1.11 | 0.106 | Tigd2 | tigger transposable element derived 2 |
| 1417586 | at | 1.52 | 0.004 | Timeless | timeless homolog (Drosophila) |
| 1449886 | a_at | -1.14 | 0.225 | Timm10 | translocase of inner mitochondrial membrane 10 homolog (yeast) |
| 1417499 | at | 1.06 | 0.247 | Timm13a | translocase of inner mitochondrial membrane 13 homolog a (yeast) |
| 1426256 | at | 1.01 | 0.639 | Timm17a | translocator of inner mitochondrial membrane 17a |
| 1448517 | at | 1.05 | 0.550 | Timm22 | translocase of inner mitochondrial membrane 22 homolog (yeast) |
| 1416485 | at | -1.01 | 0.792 | Timm23 | translocase of inner mitochondrial membrane 23 homolog (yeast) |
| 1455456 | a_at | -1.04 | 0.710 | Timm50 | translocase of inner mitochondrial membrane 50 homolog (yeast) |
| 1416345 | at | 1.11 | 0.119 | Timm8a | translocase of inner mitochondrial membrane 8 homolog a (yeast) |
| 1431665 | a_at | -1.10 | 0.052 | Timm8b | translocase of inner mitochondrial membrane 8 homolog b (yeast) |
| 1455211 | a_at | -1.13 | 0.017 | Timm9 | translocase of inner mitochondrial membrane 9 homolog (yeast) |
| 1450040 | at | 1.43 | 0.018 | Timp2 | tissue inhibitor of metalloproteinase 2 |
| 1449334 | at | 1.06 | 0.444 | Timp3 | tissue inhibitor of metalloproteinase 3 |
| 1435833 | at | -1.72 | 0.062 | Timp4 | tissue inhibitor of metalloproteinase 4 |
| 1417109 | at | 1.01 | 0.900 | Tinagl | tubulointerstitial nephritis antigen-like |
| 1451163 | at | 1.05 | 0.401 | Tinf2 | Terf1 (TRF1)-interacting nuclear factor 2 |
| 1426612 | at | 1.04 | 0.454 | Tipin | timeless interacting protein |
| 1457676 | at | 1.01 | 0.920 | Tirap | toll-interleukin 1 receptor (TIR) domain-containing adaptor protein |
| 1417749 | a_at | 1.00 | 0.959 | Tjp1 | tight junction protein 1 |

Supplementary table S1

| | | | | | |
|---|---|---|---|---|---|
| 1450985 | a_at | 1.19 | 0.042 | Tjp2 | tight junction protein 2 |
| 1451015 | at | 1.03 | 0.758 | Tkt | transketolase |
| 1419654 | at | 1.17 | 0.138 | Tle3 | transducin-like enhancer of split 3, homolog of Drosophila E(spl) |
| 1450853 | at | 1.09 | 0.347 | Tle4 | transducin-like enhancer of split 4, homolog of Drosophila E(spl) |
| 1431827 | a_at | 1.06 | 0.731 | Tlk2 | tousled-like kinase 2 (Arabidopsis) |
| 1448402 | at | 1.34 | 0.005 | Tln1 | talin 1 |
| 1418162 | at | 1.79 | 0.000 | Tlr4 | toll-like receptor 4 |
| 1439925 | at | 1.07 | 0.092 | Tm4sf1 | transmembrane 4 superfamily member 1 |
| 1452664 | a_at | 1.12 | 0.223 | Tm7sf3 | transmembrane 7 superfamily member 3 |
| 1418198 | a_at | -1.04 | 0.380 | Tm9sf1 | transmembrane 9 superfamily member 1 |
| 1423205 | at | -1.10 | 0.148 | Tm9sf4 | transmembrane 9 superfamily protein member 4 |
| 1430818 | at | -1.20 | 0.200 | Tmc1 | transmembrane channel-like gene family 1 |
| 1456981 | at | -1.80 | 0.001 | Tmc7 | transmembrane channel-like gene family 7 |
| 1452666 | a_at | 1.02 | 0.786 | Tmcc2 | transmembrane and coiled-coil domains 2 |
| 1435554 | at | -1.01 | 0.593 | Tmcc3 | transmembrane and coiled coil domains 3 |
| 1451327 | a_at | -1.06 | 0.107 | Tmed1 | transmembrane emp24 domain containing 1 |
| 1416108 | a_at | 1.10 | 0.247 | Tmed3 | transmembrane emp24 domain containing 3 |
| 1427314 | at | 1.05 | 0.598 | Tmed7 | transmembrane emp24 protein transport domain containing 7 |
| 1428854 | at | 1.15 | 0.014 | Tmed8 | transmembrane emp24 domain containing 8 |
| 1448268 | at | 1.04 | 0.345 | Tmed9 | transmembrane emp24 protein transport domain containing 9 |
| 1426649 | at | -1.11 | 0.306 | Tmeff1 | transmembrane protein with EGF-like and two follistatin-like domains 1 |
| 1416479 | a_at | -1.07 | 0.034 | Tmem14c | transmembrane protein 14C |
| 1424366 | at | -1.01 | 0.915 | Tmem15 | transmembrane protein 15 |
| 1442143 | at | -1.41 | 0.030 | Tmem16d | transmembrane protein 16D (eight membrane-spanning domains) |
| 1455309 | at | 1.12 | 0.691 | Tmem16f | transmembrane protein 16F |
| 1454836 | at | -1.08 | 0.168 | Tmem18 | transmembrane protein 18 |
| 1416261 | at | -1.08 | 0.204 | Tmem19 | transmembrane protein 19 |
| 1424711 | at | 1.29 | 0.011 | Tmem2 | transmembrane protein 2 |
| 1428095 | a_at | 1.31 | 0.016 | Tmem24 | transmembrane protein 24 |
| 1452962 | at | -1.42 | 0.000 | Tmem25 | transmembrane protein 25 |
| 1436705 | at | -1.14 | 0.017 | Tmem32 | transmembrane protein 32 |
| 1426628 | at | 1.20 | 0.002 | Tmem34 | transmembrane protein 34 |
| 1424178 | at | -1.12 | 0.147 | Tmem38a | transmembrane protein 38a |
| 1423316 | at | 1.02 | 0.801 | Tmem39a | transmembrane protein 39a |
| 1448372 | a_at | -1.07 | 0.103 | Tmem4 | transmembrane protein 4 |
| 1424191 | a_at | -1.12 | 0.248 | Tmem41a | transmembrane protein 41a |
| 1460122 | at | 1.09 | 0.296 | Tmem41b | transmembrane protein 41B |
| 1421127 | at | 1.04 | 0.518 | Tmem42 | transmembrane protein 42 |
| 1420514 | at | -1.37 | 0.003 | Tmem47 | transmembrane protein 47 |
| 1423722 | at | 1.26 | 0.003 | Tmem49 | transmembrane protein 49 |
| 1424042 | at | 1.12 | 0.095 | Tmem5 | transmembrane protein 5 |

Supplementary table S1

| | | | | | |
|---|---|---|---|---|---|
| 1452472 | at | -1.01 | 0.954 | Tmem7 | transmembrane protein 7 |
| 1422705 | at | 1.64 | 0.002 | Tmepai | transmembrane, prostate androgen induced RNA |
| 1420725 | at | 1.13 | 0.342 | Tmlhe | trimethyllysine hydroxylase, epsilon |
| 1422754 | at | -1.05 | 0.086 | Tmod1 | tropomodulin 1 |
| 1423088 | at | 1.04 | 0.624 | Tmod3 | tropomodulin 3 |
| 1449969 | at | -1.53 | 0.001 | Tmod4 | tropomodulin 4 |
| 1435031 | at | -1.02 | 0.752 | Tmpit | transmembrane protein induced by tumor necrosis factor alpha |
| 1452036 | a_at | -1.12 | 0.261 | Tmpo | thymopoietin |
| 1415906 | at | 1.06 | 0.135 | Tmsb4x | thymosin, beta 4, X chromosome |
| 1417866 | at | 1.10 | 0.140 | Tnfaip1 | tumor necrosis factor, alpha-induced protein 1 (endothelial) |
| 1416273 | at | 1.07 | 0.358 | Tnfaip2 | tumor necrosis factor, alpha-induced protein 2 |
| 1416950 | at | -1.31 | 0.002 | Tnfaip8 | tumor necrosis factor, alpha-induced protein 8 |
| 1460197 | a_at | -1.11 | 0.116 | Tnfaip9 | tumor necrosis factor, alpha-induced protein 9 |
| 1418571 | at | 2.12 | 0.014 | Tnfrsf12a | tumor necrosis factor receptor superfamily, member 12a |
| 1417291 | at | 1.16 | 0.150 | Tnfrsf1a | tumor necrosis factor receptor superfamily, member 1a |
| 1439680 | at | -1.21 | 0.368 | Tnfsf10 | tumor necrosis factor (ligand) superfamily, member 10 |
| 1452440 | at | 1.24 | 0.027 | Tnfsf12 | tumor necrosis factor (ligand) superfamily, member 12 |
| 1418345 | at | -1.04 | 0.466 | Tnfsf13 | tumor necrosis factor (ligand) superfamily, member 13 |
| 1425373 | a_at | -1.07 | 0.270 | Tnfsf5ip1 | tumor necrosis factor superfamily, member 5-induced protein 1 |
| 1458403 | at | -1.43 | 0.066 | Tnik | TRAF2 and NCK interacting kinase |
| 1427689 | a_at | 1.73 | 0.001 | Tnip1 | TNFAIP3 interacting protein 1 |
| 1448297 | a_at | -1.14 | 0.257 | Tnk2 | tyrosine kinase, non-receptor, 2 |
| 1418370 | at | 1.01 | 0.220 | Tnnc1 | troponin C, cardiac/slow skeletal |
| 1422536 | at | -1.05 | 0.013 | Tnni3 | troponin I, cardiac |
| 1458482 | at | -1.30 | 0.752 | Tnni3k | TNNI3 interacting kinase |
| 1419606 | a_at | -1.64 | 0.000 | Tnnt1 | troponin T1, skeletal, slow |
| 1418726 | a_at | -1.04 | 0.007 | Tnnt2 | troponin T2, cardiac |
| 1455043 | at | 1.06 | 0.484 | Tnpo1 | transportin 1 |
| 1425585 | at | -1.09 | 0.270 | Tnrc11 | trinucleotide repeat containing 11 (THR-associated protein) |
| 1451397 | at | -1.01 | 0.899 | Tnrc15 | trinucleotide repeat containing 15 |
| 1455141 | at | -1.09 | 0.130 | Tnrc6a | trinucleotide repeat containing 6a |
| 1436600 | at | 1.11 | 0.468 | Tnrc9 | trinucleotide repeat containing 9 |
| 1449405 | at | -1.03 | 0.803 | Tns | tensin |
| 1450798 | at | 1.11 | 0.420 | Tnxb | tenascin XB |
| 1423176 | at | 1.14 | 0.144 | Tob1 | transducer of ErbB-2.1 |
| 1417309 | at | -1.44 | 0.085 | Tob2 | transducer of ERBB2, 2 |
| 1423047 | at | -1.09 | 0.031 | Tollip | toll interacting protein |
| 1435055 | a_at | 1.07 | 0.411 | Tom1 | target of myb1 homolog (chicken) |
| 1434385 | at | 1.04 | 0.627 | Tom1l2 | target of myb1-like 2 (chicken) |
| 1429103 | at | 1.06 | 0.281 | Tomm22 | translocase of outer mitochondrial membrane 22 homolog (yeast) |
| 1448345 | at | -1.06 | 0.514 | Tomm34 | translocase of outer mitochondrial membrane 34 |

Supplementary table S1

| | | | | | |
|---|---|---|---|---|---|
| 1426118 | a_at | 1.01 | 0.902 | Tomm40 | translocase of outer mitochondrial membrane 40 homolog (yeast) |
| 1428214 | at | -1.07 | 0.002 | Tomm7 | translocase of outer mitochondrial membrane 7 homolog (yeast) |
| 1426675 | at | 1.15 | 0.039 | Tomm70a | translocase of outer mitochondrial membrane 70 homolog A (yeast) |
| 1423474 | at | -1.15 | 0.334 | Top1 | topoisomerase (DNA) I |
| 1460370 | at | -1.26 | 0.037 | Top1mt | DNA topoisomerase 1, mitochondrial |
| 1416731 | at | 1.06 | 0.233 | Top2b | topoisomerase (DNA) II beta |
| 1448173 | a_at | -1.05 | 0.418 | Top3b | topoisomerase (DNA) III beta |
| 1452241 | at | 1.09 | 0.428 | Topbp1 | topoisomerase (DNA) II beta binding protein |
| 1448838 | at | 1.07 | 0.409 | Topors | topoisomerase I binding, arginine/serine-rich |
| 1426515 | a_at | 1.02 | 0.809 | Tor1a | torsin family 1, member A (torsin A) |
| 1417819 | at | -1.08 | 0.599 | Tor1b | torsin family 1, member B |
| 1423783 | at | -1.15 | 0.205 | Tor2a | torsin family 2, member A |
| 1421998 | at | 1.32 | 0.013 | Tor3a | torsin family 3, member A |
| 1445612 | at | -1.21 | 0.374 | Tox | thymocyte selection-associated HMG box gene |
| 1415741 | at | 1.04 | 0.411 | Tparl | TPA regulated locus |
| 1434930 | at | 1.02 | 0.877 | Tpcn1 | two pore channel 1 |
| 1419493 | a_at | 1.24 | 0.040 | Tpd52 | tumor protein D52 |
| 1418412 | at | -1.04 | 0.746 | Tpd52l1 | tumor protein D52-like 1 |
| 1435659 | a_at | 1.09 | 0.005 | Tpi1 | triosephosphate isomerase 1 |
| 1427260 | a_at | 1.23 | 0.003 | Tpm3 | tropomyosin 3, gamma |
| 1433883 | at | 1.26 | 0.071 | Tpm4 | tropomyosin 4 |
| 1431893 | a_at | 1.05 | 0.570 | Tprt | trans-prenyltransferase |
| 1421733 | a_at | -1.12 | 0.024 | Tpst1 | protein-tyrosine sulfotransferase 1 |
| 1416086 | at | -1.18 | 0.136 | Tpst2 | protein-tyrosine sulfotransferase 2 |
| 1416642 | a_at | 1.02 | 0.019 | Tpt1 | tumor protein, translationally-controlled 1 |
| 1415889 | a_at | -1.09 | 0.348 | Tra1 | tumor rejection antigen gp96 |
| 1452622 | a_at | 1.23 | 0.025 | Tradd | TNFRSF1A-associated via death domain |
| 1418587 | at | 1.51 | 0.001 | Traf3 | Tnf receptor-associated factor 3 |
| 1452887 | at | -1.01 | 0.792 | Traf3ip1 | TNF receptor-associated factor 3 interacting protein 1 |
| 1423732 | at | 1.00 | 0.961 | Tram1 | translocating chain-associating membrane protein 1 |
| 1439539 | at | 1.33 | 0.015 | Tram2 | translocating chain-associating membrane protein 2 |
| 1416170 | at | -1.19 | 0.021 | Trap1 | TNF receptor-associated protein 1 |
| 1432158 | a_at | 1.04 | 0.429 | Trappc2 | trafficking protein particle complex 2 |
| 1416771 | at | -1.06 | 0.295 | Trappc3 | trafficking protein particle complex 3 |
| 1415674 | a_at | 1.33 | 0.003 | Trappc4 | trafficking protein particle complex 4 |
| 1428177 | at | 1.04 | 0.475 | Trappc6b | trafficking protein particle complex 6B |
| 1450672 | a_at | -1.35 | 0.077 | Trex1 | three prime repair exonuclease 1 |
| 1425546 | a_at | 1.38 | 0.003 | Trf | transferrin |
| 1448478 | at | 1.07 | 0.573 | Trfp | Trf (TATA binding protein-related factor)-proximal protein homolog (Drosophila) |
| 1419630 | a_at | -1.01 | 0.880 | Trim11 | tripartite motif protein 11 |
| 1436061 | at | 1.14 | 0.256 | Trim12 | tripartite motif protein 12 |

Supplementary table S1

| | | | | | |
|---|---|---|---|---|---|
| 1417888 | at | -1.01 | 0.909 | Trim13 | tripartite motif protein 13 |
| 1417027 | at | 1.08 | 0.187 | Trim2 | tripartite motif protein 2 |
| 1448940 | at | 1.05 | 0.273 | Trim21 | tripartite motif protein 21 |
| 1426969 | at | 1.07 | 0.941 | Trim23 | tripartite motif protein 23 |
| 1425974 | a_at | 1.03 | 0.529 | Trim25 | tripartite motif protein 25 |
| 1415869 | a_at | -1.11 | 0.081 | Trim28 | tripartite motif protein 28 |
| 1451663 | a_at | 1.23 | 0.000 | Trim3 | tripartite motif protein 3 |
| 1451860 | a_at | 1.24 | 0.003 | Trim30 | tripartite motif protein 30 |
| 1427476 | a_at | -1.09 | 0.246 | Trim32 | tripartite motif protein 32 |
| 1426093 | at | 1.03 | 0.698 | Trim34 | tripartite motif protein 34 |
| 1454650 | at | 1.20 | 0.008 | Trim35 | tripartite motif-containing 35 |
| 1436393 | a_at | 1.09 | 0.133 | Trim37 | tripartite motif protein 37 |
| 1418851 | at | -1.04 | 0.427 | Trim39 | tripartite motif protein 39 |
| 1421870 | at | 1.11 | 0.127 | Trim44 | tripartite motif-containing 44 |
| 1426784 | at | 1.73 | 0.001 | Trim47 | tripartite motif protein 47 |
| 1419440 | at | 1.07 | 0.359 | Trim54 | tripartite motif-containing 54 |
| 1441320 | a_at | -1.02 | 0.671 | Trim56 | tripartite motif-containing 56 |
| 1442004 | at | 1.02 | 0.879 | Trim65 | tripartite motif-containing 65 |
| 1455124 | at | -1.11 | 0.191 | Trim68 | tripartite motif containing 68 |
| 1439352 | at | -1.26 | 0.086 | Trim7 | tripartite motif protein 7 |
| 1418577 | at | -1.01 | 0.847 | Trim8 | tripartite motif protein 8 |
| 1449041 | a_at | -1.07 | 0.463 | Trip6 | thyroid hormone receptor interactor 6 |
| 1424489 | a_at | 1.18 | 0.007 | Trit1 | tRNA isopentenyltransferase 1 |
| 1452786 | at | -1.02 | 0.721 | Trmt1 | tRNA (5-methylaminomethyl-2-thiouridylate)-methyltransferase 1 |
| 1425561 | at | -1.09 | 0.292 | Trnt1 | tRNA nucleotidyl transferase, CCA-adding, 1 |
| 1433660 | at | 1.15 | 0.315 | Trp53bp1 | transformation related protein 53 binding protein 1 |
| 1416926 | at | 1.05 | 0.622 | Trp53inp1 | transformation related protein 53 inducible nuclear protein 1 |
| 1452646 | at | 1.03 | 0.150 | Trp53inp2 | tumor protein p53 inducible nuclear protein 2 |
| 1421096 | at | 1.00 | 0.991 | Trpc1 | transient receptor potential cation channel, subfamily C, member 1 |
| 1417577 | at | -1.18 | 0.096 | Trpc3 | transient receptor potential cation channel, subfamily C, member 3 |
| 1460720 | at | -1.15 | 0.041 | Trpc4ap | transient receptor potential cation channel, subfamily C, member 4 associated protein |
| 1435549 | at | -1.23 | 0.181 | Trpm4 | transient receptor potential cation channel, subfamily M, member 4 |
| 1416532 | at | -1.01 | 0.935 | Trrap | transformation/transcription domain-associated protein |
| 1424610 | at | 1.25 | 0.004 | Trub2 | TruB pseudouridine (psi) synthase homolog 2 (E. coli) |
| 1455252 | at | -1.00 | 0.962 | Tsc1 | tuberous sclerosis 1 |
| 1452105 | a_at | -1.08 | 0.229 | Tsc2 | tuberous sclerosis 2 |
| 1430535 | at | 1.11 | 0.196 | Tsc22d2 | TSC22 domain family 2 |
| 1448412 | a_at | -1.08 | 0.092 | Tsc22d4 | TSC22 domain family 4 |
| 1435118 | at | 1.11 | 0.108 | Tsen2 | tRNA splicing endonuclease 2 homolog (SEN2, S. cerevisiae) |
| 1452698 | at | -1.07 | 0.355 | Tsfm | Ts translation elongation factor, mitochondrial |
| 1417004 | at | -1.00 | 0.983 | Tsg101 | tumor susceptibility gene 101 |

Supplementary table S1

| | | | | | |
|---|---|---|---|---|---|
| 1436045 | at | 1.08 | 0.165 | Tsga10 | testis specific 10 |
| 1418097 | a_at | 1.36 | 0.005 | Tslpr | thymic stromal-derived lymphopoietin, receptor |
| 1423395 | at | -1.12 | 0.042 | Tsnax | translin-associated factor X |
| 1460239 | at | -1.20 | 0.701 | Tspan13 | tetraspanin 13 |
| 1424568 | at | -1.14 | 0.250 | Tspan2 | tetraspan 2 |
| 1416009 | at | -1.05 | 0.090 | Tspan3 | tetraspanin 3 |
| 1448276 | at | 1.26 | 0.189 | Tspan4 | tetraspanin 4 |
| 1448501 | at | 1.33 | 0.008 | Tspan6 | tetraspanin 6 |
| 1448737 | at | -1.05 | 0.769 | Tspan7 | tetraspanin 7 |
| 1428197 | at | 1.80 | 0.002 | Tspan9 | tetraspanin 9 |
| 1415908 | at | 1.01 | 0.940 | Tspyl1 | testis-specific protein, Y-encoded-like 1 |
| 1451932 | a_at | -1.03 | 0.695 | Tsrc1 | thrombospondin repeat containing 1 |
| 1427661 | a_at | 1.04 | 0.673 | Tssc4 | tumor-suppressing subchromosomal transferable fragment 4 |
| 1448609 | at | -1.32 | 0.013 | Tst | thiosulfate sulfurtransferase, mitochondrial |
| 1448495 | at | 1.01 | 0.973 | Tsta3 | tissue specific transplantation antigen P35B |
| 1416994 | at | 1.12 | 0.044 | Ttc1 | tetratricopeptide repeat domain 1 |
| 1416764 | at | -1.08 | 0.031 | Ttc11 | tetratricopeptide repeat domain 11 |
| 1435211 | at | -1.32 | 0.000 | Ttc12 | tetratricopeptide repeat domain 12 |
| 1451351 | at | 1.28 | 0.001 | Ttc13 | tetratricopeptide repeat domain 13 |
| 1428862 | at | -1.23 | 0.016 | Ttc17 | tetratricopeptide repeat domain 17 |
| 1427114 | at | -1.21 | 0.189 | Ttc19 | tetratricopeptide repeat domain 19 |
| 1460077 | at | -1.16 | 0.154 | Ttc3 | tetratricopeptide repeat domain 3 |
| 1451192 | a_at | -1.06 | 0.160 | Ttc4 | tetratricopeptide repeat domain 4 |
| 1435334 | at | -1.59 | 0.000 | Ttc7 | tetratricopeptide repeat domain 7 |
| 1418155 | at | 1.84 | 0.000 | Ttid | titin immunoglobulin domain protein (myotilin) |
| 1427112 | at | 1.33 | 0.000 | Ttl | tubulin tyrosine ligase |
| 1426427 | at | -1.71 | 0.000 | Ttll1 | tubulin tyrosine ligase-like 1 |
| 1428911 | at | -1.03 | 0.730 | Ttll4 | tubulin tyrosine ligase-like family, member 4 |
| 1448706 | at | -1.03 | 0.638 | Ttrap | Traf and Tnf receptor associated protein |
| 1434500 | at | 1.08 | 0.427 | Ttyh2 | tweety homolog 2 (Drosophila) |
| 1417373 | a_at | -0.03 | 0.282 | Tuba4 | tubulin, alpha 4 |
| 1419518 | at | -1.13 | 0.308 | Tuba8 | tubulin, alpha 8 |
| 1452571 | at | 1.46 | 0.012 | Tuba-rs1 | tubulin alpha, related sequence 1 |
| 1427838 | at | 1.59 | 0.002 | Tubb2 | tubulin, beta 2 |
| 1455719 | at | 1.13 | 0.167 | Tubb5 | tubulin, beta 5 |
| 1446353 | at | 1.25 | 0.135 | Tubb6 | tubulin, beta 6 |
| 1421748 | a_at | -1.27 | 0.016 | Tubd1 | tubulin, delta 1 |
| 1417144 | at | -1.21 | 0.082 | Tubg1 | tubulin, gamma 1 |
| 1455868 | a_at | -1.25 | 0.000 | Tubgcp2 | tubulin, gamma complex associated protein 2 |
| 1426707 | at | -1.00 | 0.989 | Tubgcp3 | tubulin, gamma complex associated protein 3 |
| 1426518 | at | -1.09 | 0.282 | Tubgcp5 | tubulin, gamma complex associated protein 5 |

Supplementary table S1

| | | | | | |
|---|---|---|---|---|---|
| 1433952 | at | -1.12 | 0.092 | Tufm | Tu translation elongation factor, mitochondrial |
| 1449008 | at | 1.16 | 0.108 | Tulp3 | tubby-like protein 3 |
| 1417593 | at | 1.03 | 0.663 | Tusc2 | tumor suppressor candidate 2 |
| 1417046 | at | -1.11 | 0.148 | Tusc4 | tumor suppressor candidate 4 |
| 1452852 | at | 1.01 | 0.834 | Twistnb | TWIST neighbor |
| 1454861 | at | 1.18 | 0.114 | Txln | taxilin |
| 1416119 | at | 1.11 | 0.001 | Txn1 | thioredoxin 1 |
| 1452782 | a_at | -1.16 | 0.015 | Txn2 | thioredoxin 2 |
| 1437143 | a_at | 1.03 | 0.802 | Txndc1 | thioredoxin domain containing 1 |
| 1440279 | at | 1.04 | 0.671 | Txndc10 | thioredoxin domain containing 10 |
| 1448782 | at | -1.34 | 0.000 | Txndc11 | thioredoxin domain containing 11 |
| 1423246 | at | -1.03 | 0.527 | Txndc4 | thioredoxin domain containing 4 (endoplasmic reticulum) |
| 1451091 | at | 1.26 | 0.024 | Txndc5 | thioredoxin domain containing 5 |
| 1423648 | at | -1.11 | 0.270 | Txndc7 | thioredoxin domain containing 7 |
| 1424512 | a_at | 1.13 | 0.069 | Txndc9 | thioredoxin domain containing 9 |
| 1415996 | at | -1.31 | 0.372 | Txnip | thioredoxin interacting protein |
| 1417340 | at | -1.07 | 0.095 | Txnl2 | thioredoxin-like 2 |
| 1423034 | at | 1.13 | 0.032 | Txnl5 | thioredoxin-like 5 |
| 1421529 | a_at | 1.05 | 0.228 | Txnrd1 | thioredoxin reductase 1 |
| 1449097 | at | -1.08 | 0.562 | Txnrd2 | thioredoxin reductase 2 |
| 1449623 | at | 1.23 | 0.050 | Txnrd3 | thioredoxin reductase 3 |
| 1417306 | at | 1.10 | 0.294 | Tyk2 | tyrosine kinase 2 |
| 1450484 | a_at | 1.07 | 0.455 | Tyki | thymidylate kinase family LPS-inducible member |
| 1450792 | at | 1.39 | 0.012 | Tyrobp | TYRO protein tyrosine kinase binding protein |
| 1428689 | at | 1.04 | 0.364 | Tysnd1 | trypsin domain containing 1 |
| 1422509 | at | -1.11 | 0.111 | U2af1 | U2 small nuclear ribonucleoprotein auxiliary factor (U2AF) 1 |
| 1434305 | at | -1.08 | 0.036 | U2af1l4 | U2 small nuclear RNA auxiliary factor 1-like 4 |
| 1449354 | at | -1.26 | 0.001 | U2af1-rs1 | U2 small nuclear ribonucleoprotein auxiliary factor (U2AF) 1, related sequence 1 |
| 1417260 | at | -1.02 | 0.755 | U2af2 | U2 small nuclear ribonucleoprotein auxiliary factor (U2AF) 2 |
| 1452985 | at | 1.22 | 0.114 | Uaca | uveal autoantigen with coiled-coil domains and ankyrin repeats |
| 1416743 | at | 1.13 | 0.102 | Uap1 | UDP-N-acetylglucosamine pyrophosphorylase 1 |
| 1416624 | a_at | -1.00 | 0.953 | Uba52 | ubiquitin A-52 residue ribosomal protein fusion product 1 |
| 1416963 | at | -1.10 | 0.083 | Ubadc1 | ubiquitin associated domain containing 1 |
| 1424281 | at | 1.17 | 0.050 | Ubap2 | ubiquitin-associated protein 2 |
| 1437532 | at | -1.22 | 0.025 | Ubce7ip1 | ubiquitin conjugating enzyme 7 interacting protein 1 |
| 1431746 | a_at | 1.10 | 0.232 | Ube1c | ubiquitin-activating enzyme E1C |
| 1451973 | at | 1.07 | 0.596 | Ube1dc1 | ubiquitin-activating enzyme E1-domain containing 1 |
| 1448116 | at | -1.03 | 0.616 | Ube1x | ubiquitin-activating enzyme E1, Chr X |
| 1417609 | at | -1.04 | 0.493 | Ube2a | ubiquitin-conjugating enzyme E2A, RAD6 homolog (S. cerevisiae) |
| 1423107 | at | -1.07 | 0.144 | Ube2b | ubiquitin-conjugating enzyme E2B, RAD6 homology (S. cerevisiae) |
| 1424062 | at | 1.07 | 0.571 | Ube2d1 | ubiquitin-conjugating enzyme E2D 1, UBC4/5 homolog (yeast) |

Supplementary table S1

| | | | | | |
|---|---|---|---|---|---|
| 1448356 | at | 1.19 | 0.285 | Ube2d2 | ubiquitin-conjugating enzyme E2D 2 |
| 1423112 | at | 1.13 | 0.391 | Ube2d3 | ubiquitin-conjugating enzyme E2D 3 (UBC4/5 homolog, yeast) |
| 1434328 | at | 1.11 | 0.037 | Ube2e1 | ubiquitin-conjugating enzyme E2E 1, UBC4/5 homolog (yeast) |
| 1424358 | at | 1.37 | 0.012 | Ube2e2 | ubiquitin-conjugating enzyme E2E 2 (UBC4/5 homolog, yeast) |
| 1440969 | at | -1.20 | 0.062 | Ube2e3 | ubiquitin-conjugating enzyme E2E 3, UBC4/5 homolog (yeast) |
| 1415688 | at | 1.14 | 0.003 | Ube2g1 | ubiquitin-conjugating enzyme E2G 1 (UBC7 homolog, C. elegans) |
| 1417032 | at | 1.15 | 0.016 | Ube2g2 | ubiquitin-conjugating enzyme E2G 2 |
| 1418631 | at | -1.18 | 0.738 | Ube2h | ubiquitin-conjugating enzyme E2H |
| 1426138 | a_at | -1.06 | 0.212 | Ube2j2 | ubiquitin-conjugating enzyme E2, J2 homolog (yeast) |
| 1429311 | at | -1.15 | 0.457 | Ube2q | ubiquitin-conjugating enzyme E2Q (putative) |
| 1415768 | a_at | 1.00 | 0.976 | Ube2r2 | ubiquitin-conjugating enzyme E2R 2 |
| 1417983 | a_at | 1.06 | 0.153 | Ube2v2 | ubiquitin-conjugating enzyme E2 variant 2 |
| 1452149 | at | -1.05 | 0.538 | Ube3b | ubiquitin protein ligase E3B |
| 1437747 | at | 1.03 | 0.642 | Ube4a | ubiquitination factor E4A, UFD2 homolog (S. cerevisiae) |
| 1448410 | at | -1.06 | 0.482 | Ube4b | ubiquitination factor E4B, UFD2 homolog (S. cerevisiae) |
| 1423461 | a_at | -1.01 | 0.933 | Ubl3 | ubiquitin-like 3 |
| 1416848 | at | -1.48 | 0.009 | Ubl5 | ubiquitin-like 5 |
| 1451465 | at | -1.06 | 0.334 | Ubl7 | ubiquitin-like 7 (bone marrow stromal cell-derived) |
| 1416443 | a_at | 1.18 | 0.003 | Uble1a | ubiquitin-like 1 (sentrin) activating enzyme E1A |
| 1448283 | a_at | 1.18 | 0.010 | Uble1b | ubiquitin-like 1 (sentrin) activating enzyme E1B |
| 1433780 | at | -1.05 | 0.759 | Ubn1 | ubinuclein 1 |
| 1460655 | a_at | 1.05 | 0.342 | Ubp1 | upstream binding protein 1 |
| 1419385 | a_at | 1.04 | 0.411 | Ubqln1 | ubiquilin 1 |
| 1450021 | at | -1.05 | 0.495 | Ubqln2 | ubiquilin 2 |
| 1448692 | at | 1.16 | 0.221 | Ubqln4 | ubiquilin 4 |
| 1450066 | at | 1.06 | 0.171 | Ubr1 | ubiquitin protein ligase E3 component n-recognin 1 |
| 1453097 | a_at | -1.00 | 0.951 | Ubtf | upstream binding transcription factor, RNA polymerase I |
| 1417557 | at | -1.09 | 0.041 | Ubxd1 | UBX domain containing 1 |
| 1426007 | a_at | -1.98 | 0.001 | Ubxd3 | UBX domain containing 3 |
| 1425020 | at | -1.09 | 0.817 | Ubxd4 | UBX domain containing 4 |
| 1448260 | at | 2.02 | 0.001 | Uchl1 | ubiquitin carboxy-terminal hydrolase L1 |
| 1419452 | at | -1.00 | 0.955 | Uchl5 | ubiquitin carboxyl-terminal esterase L5 |
| 1426909 | at | 3.37 | 0.000 | Uck2 | uridine-cytidine kinase 2 |
| 1424646 | at | 1.17 | 0.144 | Uckl1 | uridine-cytidine kinase 1-like 1 |
| 1448188 | at | -1.04 | 0.636 | Ucp2 | uncoupling protein 2 (mitochondrial, proton carrier) |
| 1420657 | at | -1.44 | 0.138 | Ucp3 | uncoupling protein 3 (mitochondrial, proton carrier) |
| 1418087 | at | 1.04 | 0.824 | Ufd1l | ubiquitin fusion degradation 1 like |
| 1435133 | at | 1.03 | 0.568 | Ugcg | UDP-glucose ceramide glucosyltransferase |
| 1416308 | at | 1.32 | 0.009 | Ugdh | UDP-glucose dehydrogenase |
| 1426461 | at | 1.32 | 0.005 | Ugp2 | UDP-glucose pyrophosphorylase 2 |
| 1433737 | at | -1.00 | 0.918 | Uhmk1 | U2AF homology motif (UHM) kinase 1 |

Supplementary table S1

| | | | | | |
|---|---|---|---|---|---|
| 1448370 | at | -1.06 | 0.206 | Ulk1 | Unc-51 like kinase 1 (C. elegans) |
| 1455832 | a_at | -1.00 | 0.997 | Umps | uridine monophosphate synthetase |
| 1417757 | at | -1.01 | 0.939 | Unc13b | unc-13 homolog B (C. elegans) |
| 1421982 | a_at | 1.27 | 0.045 | Unc50 | unc-50 homolog (C. elegans) |
| 1426666 | a_at | -1.03 | 0.540 | Unc84a | unc-84 homolog A (C. elegans) |
| 1425753 | a_at | 1.00 | 0.959 | Ung | uracil-DNA glycosylase |
| 1452971 | at | 1.26 | 0.002 | Upf3a | UPF3 regulator of nonsense transcripts homolog A (yeast) |
| 1448292 | at | -1.12 | 0.001 | Uqcr | ubiquinol-cytochrome c reductase (6.4kD) subunit |
| 1428782 | a_at | -1.10 | 0.007 | Uqcrc1 | ubiquinol-cytochrome c reductase core protein 1 |
| 1428631 | a_at | -1.06 | 0.304 | Uqcrc2 | ubiquinol cytochrome c reductase core protein 2 |
| 1450968 | at | 1.02 | 0.358 | Uqcrfs1 | ubiquinol-cytochrome c reductase, Rieske iron-sulfur polypeptide 1 |
| 1452133 | at | -1.03 | 0.001 | Uqcrh | ubiquinol-cytochrome c reductase hinge protein |
| 1417206 | at | -1.13 | 0.050 | Urod | uroporphyrinogen decarboxylase |
| 1423482 | at | -1.10 | 0.018 | Uros | uroporphyrinogen III synthase |
| 1418241 | at | -1.28 | 0.024 | Usf2 | upstream transcription factor 2 |
| 1451500 | at | -1.28 | 0.063 | Ushbp1 | Usher syndrome 1C binding protein 1 |
| 1422307 | at | -1.35 | 0.004 | Usmg3 | upregulated during skeletal muscle growth 3 |
| 1417626 | at | 1.03 | 0.489 | Usmg4 | upregulated during skeletal muscle growth 4 |
| 1448179 | at | -1.08 | 0.000 | Usmg5 | upregulated during skeletal muscle growth 5 |
| 1451080 | at | 1.18 | 0.042 | Usp1 | ubiquitin specific protease 1 |
| 1448230 | at | -1.07 | 0.274 | Usp10 | ubiquitin specific protease 10 |
| 1434483 | at | 1.24 | 0.001 | Usp12 | ubiquitin specific protease 12 |
| 1455473 | at | 1.10 | 0.109 | Usp13 | ubiquitin specific protease 13 (isopeptidase T-3) |
| 1454036 | a_at | 1.16 | 0.053 | Usp15 | ubiquitin specific protease 15 |
| 1431951 | a_at | 1.42 | 0.000 | Usp16 | ubiquitin specific protease 16 |
| 1418191 | at | 1.34 | 0.116 | Usp18 | ubiquitin specific protease 18 |
| 1455205 | a_at | 1.07 | 0.320 | Usp19 | ubiquitin specific protease 19 |
| 1417168 | a_at | -1.38 | 0.022 | Usp2 | ubiquitin specific protease 2 |
| 1423777 | at | -1.05 | 0.621 | Usp20 | ubiquitin specific protease 20 |
| 1454034 | a_at | -1.13 | 0.120 | Usp21 | ubiquitin specific protease 21 |
| 1437151 | at | 1.07 | 0.197 | Usp22 | ubiquitin specific protease 22 |
| 1452868 | at | 1.19 | 0.022 | Usp24 | ubiquitin specific protease 24 |
| 1445655 | at | 1.30 | 0.180 | Usp28 | ubiquitin specific protease 28 |
| 1425022 | at | -1.06 | 0.858 | Usp3 | ubiquitin specific protease 3 |
| 1434724 | at | -1.09 | 0.065 | Usp31 | ubiquitin specific protease 31 |
| 1426709 | a_at | -1.18 | 0.044 | Usp33 | ubiquitin specific protease 33 |
| 1455237 | at | -1.05 | 0.677 | Usp36 | ubiquitin specific protease 36 |
| 1460209 | at | 1.15 | 0.162 | Usp39 | ubiquitin specific protease 39 |
| 1450892 | a_at | 1.10 | 0.042 | Usp4 | ubiquitin specific protease 4 (proto-oncogene) |
| 1434782 | at | -1.05 | 0.255 | Usp42 | ubiquitin specific protease 42 |
| 1427990 | at | -1.02 | 0.818 | Usp45 | ubiquitin specific protease 45 |

Supplementary table S1

| | | | | | |
|---|---|---|---|---|---|
| 1426976 | at | 1.26 | 0.001 | Usp47 | ubiquitin specific protease 47 |
| 1419278 | at | -1.20 | 0.057 | Usp48 | ubiquitin specific protease 48 |
| 1448311 | at | 1.15 | 0.112 | Usp5 | ubiquitin specific protease 5 (isopeptidase T) |
| 1426700 | a_at | -1.12 | 0.253 | Usp52 | ubiquitin specific protease 52 |
| 1428731 | at | -1.03 | 0.776 | Usp54 | ubiquitin specific protease 54 |
| 1454948 | at | 1.01 | 0.883 | Usp7 | ubiquitin specific protease 7 |
| 1422883 | at | 1.19 | 0.028 | Usp8 | ubiquitin specific protease 8 |
| 1435099 | at | 1.08 | 0.562 | Utp14a | UTP14, U3 small nucleolar ribonucleoprotein, homolog A (yeast) |
| 1452222 | at | -1.09 | 0.563 | Utrn | utrophin |
| 1454706 | at | -1.01 | 0.818 | Uvrag | UV radiation resistance associated gene |
| 1426275 | a_at | 1.29 | 0.040 | Uxs1 | UDP-glucuronate decarboxylase 1 |
| 1418986 | a_at | -1.14 | 0.007 | Uxt | ubiquitously expressed transcript |
| 1416231 | at | 1.10 | 0.067 | Vac14 | Vac14 homolog (S. cerevisiae) |
| 1421863 | at | -1.07 | 0.348 | Vamp1 | vesicle-associated membrane protein 1 |
| 1420834 | at | -1.06 | 0.785 | Vamp2 | vesicle-associated membrane protein 2 |
| 1433916 | at | 1.03 | 0.676 | Vamp3 | vesicle-associated membrane protein 3 |
| 1430522 | a_at | 1.23 | 0.005 | Vamp5 | vesicle-associated membrane protein 5 |
| 1420624 | a_at | -1.11 | 0.005 | Vamp8 | vesicle-associated membrane protein 8 |
| 1427234 | at | 1.10 | 0.077 | Vangl1 | vang, van gogh-like 1 (Drosophila) |
| 1423152 | at | 1.08 | 0.069 | Vapb | vesicle-associated membrane protein, associated protein B and C |
| 1434349 | at | -1.08 | 0.455 | Vars2l | valyl-tRNA synthetase 2-like |
| 1451097 | at | 1.22 | 0.003 | Vasp | vasodilator-stimulated phosphoprotein |
| 1423726 | at | 1.33 | 0.008 | Vat1 | vesicle amine transport protein 1 homolog (T californica) |
| 1435244 | at | 1.08 | 0.279 | Vav2 | Vav2 oncogene |
| 1421750 | a_at | 1.01 | 0.757 | Vbp1 | von Hippel-Lindau binding protein 1 |
| 1448162 | at | 1.24 | 0.037 | Vcam1 | vascular cell adhesion molecule 1 |
| 1416156 | at | 1.07 | 0.081 | Vcl | vinculin |
| 1423030 | at | 1.05 | 0.484 | Vcp | valosin containing protein |
| 1415998 | at | -1.05 | 0.387 | Vdac1 | voltage-dependent anion channel 1 |
| 1415990 | at | 1.02 | 0.394 | Vdac2 | voltage-dependent anion channel 2 |
| 1416175 | a_at | -1.08 | 0.034 | Vdac3 | voltage-dependent anion channel 3 |
| 1424274 | at | 1.03 | 0.484 | Vdp | vesicle docking protein |
| 1417844 | at | -1.05 | 0.506 | Vdrip | vitamin D receptor interacting protein |
| 1451959 | a_at | 1.23 | 0.942 | Vegfa | vascular endothelial growth factor A |
| 1451803 | a_at | -1.42 | 0.001 | Vegfb | vascular endothelial growth factor B |
| 1419417 | at | 1.24 | 0.030 | Vegfc | vascular endothelial growth factor C |
| 1434708 | at | -1.01 | 0.834 | Vhlh | von Hippel-Lindau syndrome homolog |
| 1450850 | at | -1.10 | 0.054 | Vil2 | villin 2 |
| 1456292 | a_at | 1.25 | 0.029 | Vim | vimentin |
| 1452770 | at | 1.16 | 0.078 | Vkorc1 | vitamin K epoxide reductase complex, subunit 1 |
| 1434916 | at | 1.10 | 0.147 | Vkorc1l1 | vitamin K epoxide reductase complex, subunit 1-like 1 |

Supplementary table S1

| | | | | | |
|---|---|---|---|---|---|
| 1438258 | at | -1.29 | 0.005 | Vldlr | very low density lipoprotein receptor |
| 1428109 | at | 1.07 | 0.392 | Vps11 | vacuolar protein sorting 11 (yeast) |
| 1457674 | at | 1.37 | 0.074 | Vps13a | vacuolar protein sorting 13A (yeast) |
| 1434571 | at | -1.03 | 0.492 | Vps13b | vacuolar protein sorting 13B (yeast) |
| 1416254 | a_at | 1.06 | 0.393 | Vps16 | vacuolar protein sorting 16 (yeast) |
| 1428165 | at | 1.28 | 0.005 | Vps24 | vacuolar protein sorting 24 (yeast) |
| 1420495 | a_at | -1.05 | 0.316 | Vps26 | vacuolar protein sorting 26 (yeast) |
| 1448447 | at | -1.10 | 0.010 | Vps28 | vacuolar protein sorting 28 (yeast) |
| 1417659 | at | 1.32 | 0.000 | Vps29 | vacuolar protein sorting 29 (S. pombe) |
| 1434816 | at | -1.04 | 0.327 | Vps33a | vacuolar protein sorting 33A (yeast) |
| 1415784 | at | 1.12 | 0.913 | Vps35 | vacuolar protein sorting 35 |
| 1451720 | at | 1.11 | 0.228 | Vps39 | vacuolar protein sorting 39 (yeast) |
| 1437901 | a_at | -1.06 | 0.461 | Vps41 | vacuolar protein sorting 41 (yeast) |
| 1418143 | at | 1.12 | 0.110 | Vps45 | vacuolar protein sorting 45 (yeast) |
| 1417510 | at | -1.06 | 0.257 | Vps4a | vacuolar protein sorting 4a (yeast) |
| 1417007 | a_at | 1.14 | 0.013 | Vps4b | vacuolar protein sorting 4b (yeast) |
| 1437078 | at | -1.03 | 0.617 | Vps52 | vacuolar protein sorting 52 (yeast) |
| 1449095 | at | 1.13 | 0.155 | Vps54 | vacuolar protein sorting 54 (yeast) |
| 1418121 | at | -1.09 | 0.060 | Vrk3 | vaccinia related kinase 3 |
| 1422634 | a_at | -1.23 | 0.062 | Vsig2 | V-set and immunoglobulin domain containing 2 |
| 1445374 | at | -1.31 | 0.035 | Vti1a | vesicle transport through interaction with t-SNAREs homolog 1A (yeast) |
| 1449003 | a_at | 1.11 | 0.002 | Vti1b | vesicle transport through interaction with t-SNAREs 1B homolog |
| 1455098 | a_at | -1.35 | 0.006 | Vtn | vitronectin |
| 1435386 | at | 1.34 | 0.002 | Vwf | Von Willebrand factor homolog |
| 1415694 | at | 1.02 | 0.827 | Wars | tryptophanyl-tRNA synthetase |
| 1454673 | at | 1.13 | 0.116 | Wasf2 | WAS protein family, member 2 |
| 1426776 | at | -1.04 | 0.339 | Wasl | Wiskott-Aldrich syndrome-like (human) |
| 1449441 | a_at | -1.14 | 0.010 | Wbp1 | WW domain binding protein 1 |
| 1415799 | at | 1.03 | 0.537 | Wbp11 | WW domain binding protein 11 |
| 1448121 | at | -1.06 | 0.122 | Wbp2 | WW domain binding protein 2 |
| 1419374 | at | -1.05 | 0.948 | Wbp4 | WW domain binding protein 4 |
| 1451230 | a_at | 1.08 | 0.205 | Wbp5 | WW domain binding protein 5 |
| 1419185 | a_at | 1.07 | 0.550 | Wbscr14 | Williams-Beuren syndrome chromosome region 14 homolog (human) |
| 1416622 | at | 1.00 | 0.988 | Wbscr16 | Williams-Beuren syndrome chromosome region 16 homolog (human) |
| 1418081 | at | 1.02 | 0.686 | Wbscr18 | Williams-Beuren syndrome chromosome region 18 homolog (human) |
| 1428428 | at | -1.01 | 0.822 | Wbscr21 | Williams Beuren syndrome chromosome region 21 |
| 1423575 | a_at | 1.04 | 0.394 | Wbscr22 | Williams Beuren syndrome chromosome region 22 |
| 1437358 | at | 1.36 | 0.040 | Wdfy1 | WD repeat and FYVE domain containing 1 |
| 1434517 | at | -1.05 | 0.362 | Wdfy2 | WD repeat and FYVE domain containing 2 |
| 1428517 | at | -1.03 | 0.842 | Wdfy3 | WD repeat and FYVE domain containing 3 |
| 1450851 | at | 1.37 | 0.011 | Wdr1 | WD repeat domain 1 |

Supplementary table S1

| | | | | | |
|---|---|---|---|---|---|
| 1452820 | at | 1.21 | 0.000 | Wdr11 | WD repeat domain 11 |
| 1451728 | at | 1.19 | 0.011 | Wdr13 | WD repeat domain 13 |
| 1455702 | at | 1.08 | 0.359 | Wdr22 | WD repeat domain 22 |
| 1460189 | at | -1.17 | 0.000 | Wdr23 | WD repeat domain 23 |
| 1433746 | at | 1.00 | 0.960 | Wdr3 | WD repeat domain 3 |
| 1438727 | at | 1.42 | 0.000 | Wdr32 | WD repeat domain 32 |
| 1453322 | at | -1.17 | 0.593 | Wdr33 | WD repeat domain 33 |
| 1454790 | at | -1.42 | 0.001 | Wdr35 | WD repeat domain 35 |
| 1451087 | at | 1.15 | 0.020 | Wdr36 | WD repeat domain 36 |
| 1434077 | at | 1.09 | 0.293 | Wdr37 | WD repeat domain 37 |
| 1450953 | at | 1.04 | 0.615 | Wdr39 | WD repeat domain 39 |
| 1429393 | at | 1.04 | 0.877 | Wdr40a | WD repeat domain 40A |
| 1434170 | at | -1.63 | 0.006 | Wdr40b | WD repeat domain 40B |
| 1434134 | at | 1.36 | 0.038 | Wdr42a | WD repeat domain 42A |
| 1428390 | at | 1.24 | 0.083 | Wdr43 | WD repeat domain 43 |
| 1451754 | a_at | -1.09 | 0.335 | Wdr45 | WD repeat domain 45 |
| 1416281 | at | -1.02 | 0.797 | Wdr45l | Wdr45 like |
| 1434558 | at | 1.44 | 0.000 | Wdr47 | WD repeat domain 47 |
| 1426723 | at | -1.11 | 0.017 | Wdr48 | WD repeat domain 48 |
| 1416581 | at | -1.06 | 0.189 | Wdr5 | WD repeat domain 5 |
| 1454817 | at | -1.03 | 0.736 | Wdr50 | WD repeat domain 50 |
| 1428839 | at | -1.36 | 0.001 | Wdr53 | WD repeat domain 53 |
| 1427568 | a_at | -1.02 | 0.823 | Wdr56 | WD repeat domain 56 |
| 1428264 | at | 1.17 | 0.911 | Wdr57 | WD repeat domain 57 (U5 snRNP specific) |
| 1415770 | at | -1.10 | 0.201 | Wdr6 | WD repeat domain 6 |
| 1423923 | a_at | -1.13 | 0.233 | Wdr8 | WD repeat domain 8 |
| 1437250 | at | -1.03 | 0.650 | Wdt2 | whn-dependent transcript 2 |
| 1434560 | at | -1.33 | 0.019 | Wdtc1 | WD and tetratricopeptide repeats 1 |
| 1416773 | at | 1.04 | 0.724 | Wee1 | wee 1 homolog (S. pombe) |
| 1448411 | at | 1.03 | 0.606 | Wfs1 | Wolfram syndrome 1 homolog (human) |
| 1447931 | at | 1.16 | 0.041 | Whsc1l1 | Wolf-Hirschhorn syndrome candidate 1-like 1 (human) |
| 1449353 | at | 1.30 | 0.050 | Wig1 | wild-type p53-induced gene 1 |
| 1419015 | at | 5.11 | 0.000 | Wisp2 | WNT1 inducible signaling pathway protein 2 |
| 1434280 | at | 1.08 | 0.717 | Wiz | widely-interspaced zinc finger motifs |
| 1448818 | at | -1.58 | 0.004 | Wnt5a | wingless-related MMTV integration site 5A |
| 1436978 | at | 1.20 | 0.028 | Wnt9a | wingless-type MMTV integration site 9A |
| 1460446 | at | -1.13 | 0.121 | Wrb | tryptophan rich basic protein |
| 1425074 | at | -1.05 | 0.485 | Wrn | Werner syndrome homolog (human) |
| 1448376 | at | -1.28 | 0.000 | Wrnip1 | Werner helicase interacting protein 1 |
| 1425241 | a_at | 1.14 | 0.131 | Wsb1 | WD repeat and SOCS box-containing 1 |
| 1452401 | at | -1.05 | 0.419 | Wtap | Wilms' tumour 1-associating protein |

Supplementary table S1

| | | | | | |
|---|---|---|---|---|---|
| 1448146 | at | 1.21 | 0.010 | Wwp2 | WW domain containing E3 ubiquitin protein ligase 2 |
| 1448134 | at | 1.08 | 0.359 | X99384 | cDNA sequence X99384 |
| 1417390 | at | -1.00 | 0.995 | Xab1 | XPA binding protein 1 |
| 1448278 | at | -1.01 | 0.907 | Xab2 | XPA binding protein 2 |
| 1420886 | a_at | -1.22 | 0.022 | Xbp1 | X-box binding protein 1 |
| 1451006 | at | -1.17 | 0.052 | Xdh | xanthine dehydrogenase |
| 1434113 | a_at | 1.11 | 0.008 | Xpmc2h | XPMC2 prevents mitotic catastrophe 2 homolog (Xenopus laevis) |
| 1422443 | at | 1.20 | 0.014 | Xpnpep1 | X-prolyl aminopeptidase (aminopeptidase P) 1, soluble |
| 1456396 | at | 1.24 | 0.001 | Xpo4 | exportin 4 |
| 1428560 | at | 1.04 | 0.429 | Xpo5 | exportin 5 |
| 1422759 | a_at | 1.02 | 0.851 | Xpo6 | exportin 6 |
| 1439411 | a_at | -1.09 | 0.533 | Xpo7 | exportin 7 |
| 1428949 | at | 1.13 | 0.179 | Xpot | exportin, tRNA (nuclear export receptor for tRNAs) |
| 1426993 | at | 1.22 | 0.865 | Xpr1 | xenotropic and polytropic retrovirus receptor 1 |
| 1416587 | a_at | 1.05 | 0.706 | Xrcc1 | X-ray repair complementing defective repair in Chinese hamster cells 1 |
| 1455335 | at | -1.18 | 0.029 | Xrcc2 | X-ray repair complementing defective repair in Chinese hamster cells 2 |
| 1424601 | at | -1.02 | 0.762 | Xrcc4 | X-ray repair complementing defective repair in Chinese hamster cells 4 |
| 1451968 | at | 1.02 | 0.810 | Xrcc5 | X-ray repair complementing defective repair in Chinese hamster cells 5 |
| 1417437 | at | 1.38 | 0.000 | Xrcc6 | X-ray repair complementing defective repair in Chinese hamster cells 6 |
| 1437475 | at | 1.06 | 0.549 | Xrn1 | 5'-3' exoribonuclease 1 |
| 1422842 | at | 1.02 | 0.986 | Xrn2 | 5'-3' exoribonuclease 2 |
| 1423399 | a_at | 1.10 | 0.124 | Yaf2 | YY1 associated factor 2 |
| 1448363 | at | 1.08 | 0.737 | Yap1 | yes-associated protein 1 |
| 1460638 | at | 1.32 | 0.020 | Yars | tyrosyl-tRNA synthetase |
| 1423105 | a_at | 1.07 | 0.498 | Yeats4 | YEATS domain containing 4 |
| 1418470 | at | | 0.142 | Yes1 | Yamaguchi sarcoma viral (v-yes) oncogene homolog 1 |
| 1423939 | a_at | 1.09 | 0.180 | Yif1 | Yip1 interacting factor homolog (S. cerevisiae) |
| 1428445 | at | -1.03 | 0.535 | Yif1b | Yip1 interacting factor homolog B (S. cerevisiae) |
| 1424532 | at | -1.07 | 0.235 | Ylpm1 | YLP motif containing 1 |
| 1423360 | at | -0.40 | 0.169 | Yme1l1 | YME1-like 1 (S. cerevisiae) |
| 1426624 | a_at | -1.21 | 0.220 | Ypel3 | yippee-like 3 (Drosophila) |
| 1433593 | at | 1.10 | 0.147 | Ypel5 | yippee-like 5 (Drosophila) |
| 1416539 | at | -1.25 | 0.045 | Ysg2 | yolk sac gene 2 |
| 1415767 | at | -1.07 | 0.437 | Ythdf1 | YTH domain family 1 |
| 1460357 | at | -1.02 | 0.902 | Ythdf2 | YTH domain family 2 |
| 1420880 | a_at | 1.09 | 0.257 | Ywhab | tyrosine 3-monooxygenase/tryptophan 5-monooxygenase activation protein, beta polypeptide |
| 1420817 | at | 1.01 | 0.907 | Ywhag | 3-monooxygenase/tryptophan 5-monooxygenase activation protein, gamma polypeptide |
| 1416004 | at | 1.09 | 0.077 | Ywhah | tyrosine 3-monooxygenase/tryptophan 5-monooxygenase activation protein, eta polypeptide |
| 1420829 | a_at | 1.21 | 0.023 | Ywhaq | tyrosine 3-monooxygenase/tryptophan 5-monooxygenase activation protein, theta polypeptide |
| 1416103 | at | -1.13 | 0.604 | Ywhaz | tyrosine 3-monooxygenase/tryptophan 5-monooxygenase activation protein, zeta polypeptide |
| 1416083 | at | -1.10 | 0.513 | Za20d2 | zinc finger, A20 domain containing 2 |

Supplementary table S1

| | | | | | |
|---|---|---|---|---|---|
| 1451277 | at | -1.26 | 0.000 | Zadh2 | zinc binding alcohol dehydrogenase, domain containing 2 |
| 1427090 | at | 1.01 | 0.928 | Zbed4 | zinc finger, BED domain containing 4 |
| 1424750 | at | 1.03 | 0.704 | Zbtb1 | zinc finger and BTB domain containing 1 |
| 1436068 | at | -1.14 | 0.328 | Zbtb10 | zinc finger and BTB domain containing 10 |
| 1454826 | at | 1.02 | 0.743 | Zbtb11 | zinc finger and BTB domain containing 11 |
| 1442026 | at | -1.32 | 0.471 | Zbtb16 | zinc finger and BTB domain containing 16 |
| 1443471 | at | -1.38 | 0.036 | Zbtb20 | zinc finger and BTB domain containing 20 |
| 1434629 | at | -1.14 | 0.278 | Zbtb26 | zinc finger and BTB domain containing 26 |
| 1434022 | at | -1.04 | 0.651 | Zbtb33 | zinc finger and BTB domain containing 33 |
| 1438269 | at | -1.13 | 0.262 | Zbtb38 | zinc finger and BTB domain containing 38 |
| 1452932 | at | 1.26 | 0.007 | Zbtb7a | zinc finger and BTB domain containing 7a |
| 1436183 | at | 1.00 | 0.951 | Zc3hav1 | zinc finger CCCH type, antiviral 1 |
| 1426654 | at | 1.00 | 0.954 | Zc3hc1 | zinc finger, C3HC type 1 |
| 1426774 | at | -1.10 | 0.035 | Zc3hdc1 | zinc finger CCCH type domain containing 1 |
| 1419899 | at | -1.02 | 0.647 | Zc3hdc7 | zinc finger CCCH type domain containing 7 |
| 1418495 | at | -1.31 | 0.002 | Zc3hdc8 | zinc finger CCCH type domain containing 8 |
| 1418170 | a_at | -1.21 | 0.178 | Zcchc14 | zinc finger, CCHC domain containing 14 |
| 1428402 | at | -1.15 | 0.761 | Zcchc3 | zinc finger, CCHC domain containing 3 |
| 1433910 | at | 1.14 | 0.004 | Zcchc6 | zinc finger, CCHC domain containing 6 |
| 1417321 | at | 1.10 | 0.220 | Zcchc7 | zinc finger, CCHC domain containing 7 |
| 1423755 | at | -1.04 | 0.555 | Zcchc8 | zinc finger, CCHC domain containing 8 |
| 1424557 | at | -1.18 | 0.068 | Zcchc9 | zinc finger, CCHC domain containing 9 |
| 1455268 | at | -1.02 | 0.730 | Zcsl2 | zinc finger, CSL domain containing 2 |
| 1429532 | at | -1.12 | 0.112 | Zcwcc1 | zinc finger, CW-type with coiled-coil domain 1 |
| 1434436 | at | -1.03 | 0.766 | Zcwcc2 | zinc finger, CW-type with coiled-coil domain 2 |
| 1420092 | at | 1.08 | 0.640 | Zcwcc3 | zinc finger, CW-type with coiled-coil domain 3 |
| 1428307 | at | -1.13 | 0.335 | Zdhhc13 | zinc finger, DHHC domain containing 13 |
| 1423668 | at | -1.38 | 0.002 | Zdhhc14 | zinc finger, DHHC domain containing 14 |
| 1421115 | a_at | 1.14 | 0.016 | Zdhhc16 | zinc finger, DHHC domain containing 16 |
| 1434397 | at | -1.02 | 0.747 | Zdhhc17 | zinc finger, DHHC domain containing 17 |
| 1452654 | at | 1.05 | 0.490 | Zdhhc2 | zinc finger, DHHC domain containing 2 |
| 1434236 | at | 1.29 | 0.001 | Zdhhc20 | zinc finger, DHHC domain containing 20 |
| 1423646 | at | 1.01 | 0.921 | Zdhhc3 | zinc finger, DHHC domain containing 3 |
| 1423300 | at | 1.08 | 0.069 | Zdhhc6 | zinc finger, DHHC domain containing 6 |
| 1416545 | at | -1.05 | 0.556 | Zdhhc7 | zinc finger, DHHC domain containing 7 |
| 1418408 | at | 1.04 | 0.567 | Zfand1 | zinc finger, AN1-type domain 1 |
| 1417791 | a_at | -1.21 | 0.948 | Zfml | zinc finger, matrin-like |
| 1450090 | at | -1.13 | 0.470 | Zfp101 | zinc finger protein 101 |
| 1425332 | at | 1.22 | 0.086 | Zfp106 | zinc finger protein 106 |
| 1450998 | at | -1.03 | 0.617 | Zfp110 | zinc finger protein 110 |
| 1421458 | at | -1.06 | 0.730 | Zfp112 | zinc finger protein 112 |

Supplementary table S1

| | | | | | |
|---|---|---|---|---|---|
| 1421519 | a_at | -1.17 | 0.140 | Zfp120 | zinc finger protein 120 |
| 1427536 | at | -1.05 | 0.874 | Zfp125 | zinc finger protein 125 |
| 1428616 | at | 1.05 | 0.308 | Zfp131 | zinc finger protein 131 |
| 1441612 | at | -1.13 | 0.163 | Zfp148 | zinc finger protein 148 |
| 1420865 | at | -1.07 | 0.482 | Zfp161 | zinc finger protein 161 |
| 1423750 | a_at | -1.05 | 0.354 | Zfp162 | zinc finger protein 162 |
| 1428170 | at | 1.10 | 0.155 | Zfp180 | zinc finger protein 180 |
| 1426895 | at | -1.21 | 0.129 | Zfp191 | zinc finger protein 191 |
| 1455778 | at | -1.09 | 0.302 | Zfp192 | zinc finger protein 192 |
| 1452970 | at | -1.18 | 0.001 | Zfp198 | zinc finger protein 198 |
| 1449913 | at | -1.13 | 0.415 | Zfp2 | zinc finger protein 2 |
| 1451292 | at | -1.11 | 0.070 | Zfp212 | Zinc finger protein 212 |
| 1424301 | at | -1.12 | 0.309 | Zfp219 | zinc finger protein 219 |
| 1449329 | at | -1.36 | 0.011 | Zfp235 | zinc finger protein 235 |
| 1455153 | at | -1.13 | 0.084 | Zfp236 | zinc finger protein 236 |
| 1417010 | at | -1.13 | 0.078 | Zfp238 | zinc finger protein 238 |
| 1453034 | at | -1.15 | 0.052 | Zfp251 | zinc finger protein 251 |
| 1438685 | at | -1.09 | 0.022 | Zfp258 | zinc finger protein 258 |
| 1419281 | a_at | 1.19 | 0.003 | Zfp259 | zinc finger protein 259 |
| 1427120 | at | 1.00 | 0.949 | Zfp26 | zinc finger protein 26 |
| 1419164 | at | 1.03 | 0.853 | Zfp260 | zinc finger protein 260 |
| 1417794 | at | 1.00 | 0.970 | Zfp261 | zinc finger protein 261 |
| 1454652 | at | 1.13 | 0.003 | Zfp265 | zinc finger protein 265 |
| 1418397 | at | -1.12 | 0.202 | Zfp275 | Zinc finger protein 275 |
| 1418312 | at | -1.24 | 0.098 | Zfp276 | zinc finger protein (C2H2 type) 276 |
| 1433953 | at | 1.04 | 0.515 | Zfp277 | zinc finger protein 277 |
| 1429739 | a_at | -1.03 | 0.727 | Zfp278 | zinc finger protein 278 |
| 1426377 | at | -1.05 | 0.836 | Zfp281 | zinc finger protein 281 |
| 1457842 | at | -1.23 | 0.084 | Zfp292 | zinc finger protein 292 |
| 1441973 | at | 1.11 | 0.342 | Zfp295 | zinc finger protein 295 |
| 1448705 | at | 1.16 | 0.014 | Zfp297 | zinc finger protein 297 |
| 1435900 | at | 1.06 | 0.407 | Zfp297b | zinc finger protein 297B |
| 1422551 | at | -1.09 | 0.228 | Zfp306 | zinc finger protein 306 |
| 1427874 | at | -1.05 | 0.218 | Zfp313 | zinc finger protein 313 |
| 1430558 | at | 1.11 | 0.058 | Zfp318 | zinc finger protein 318 |
| 1455638 | at | -1.41 | 0.050 | Zfp319 | zinc finger protein 319 |
| 1439035 | at | 1.16 | 0.212 | Zfp322a | zinc finger protein 322a |
| 1419181 | at | 1.24 | 0.019 | Zfp326 | zinc finger protein 326 |
| 1451216 | at | 1.16 | 0.021 | Zfp330 | zinc finger protein 330 |
| 1428348 | at | 1.03 | 0.525 | Zfp336 | zinc finger protein 336 |
| 1417778 | at | -1.05 | 0.286 | Zfp35 | zinc finger protein 35 |

Supplementary table S1

| | | | | | |
|---|---|---|---|---|---|
| 1439044 | at | 1.03 | 0.686 | Zfp354c | zinc finger protein 354C |
| 1456151 | at | -1.23 | 0.000 | Zfp358 | zinc finger protein 358 |
| 1452519 | a_at | -1.02 | 0.923 | Zfp36 | zinc finger protein 36 |
| 1423615 | at | 1.66 | 0.000 | Zfp364 | zinc finger protein 364 |
| 1433623 | at | 1.18 | 0.070 | Zfp367 | zinc finger protein 367 |
| 1455681 | at | 1.00 | 0.974 | Zfp369 | zinc finger protein 369 |
| 1450644 | at | 1.03 | 0.581 | Zfp36l1 | zinc finger protein 36, C3H type-like 1 |
| 1437626 | at | 1.25 | 0.028 | Zfp36l2 | zinc finger protein 36, C3H type-like 2 |
| 1451146 | at | -1.26 | 0.012 | Zfp386 | zinc finger protein 386 (Kruppel-like) |
| 1441324 | at | -1.44 | 0.013 | Zfp395 | zinc finger protein 395 |
| 1435664 | at | -1.22 | 0.001 | Zfp397 | zinc finger protein 397 |
| 1439979 | at | -1.30 | 0.073 | Zfp398 | zinc finger protein 398 |
| 1451150 | at | -1.09 | 0.384 | Zfp410 | zinc finger protein 410 |
| 1451244 | a_at | -1.17 | 0.046 | Zfp422 | zinc finger protein 422 |
| 1451997 | at | -1.03 | 0.607 | Zfp426 | zinc finger protein 426 |
| 1454774 | at | -1.11 | 0.478 | Zfp445 | zinc finger protein 445 |
| 1439945 | at | -1.15 | 0.007 | Zfp449 | zinc finger protein 449 |
| 1427208 | at | 1.09 | 0.915 | Zfp451 | zinc finger protein 451 |
| 1445411 | at | -1.07 | 0.767 | Zfp458 | zinc finger protein 458 |
| 1425231 | a_at | -1.26 | 0.032 | Zfp46 | zinc finger protein 46 |
| 1456789 | at | 1.10 | 0.412 | Zfp462 | zinc finger protein 462 |
| 1439550 | at | -1.44 | 0.009 | Zfp469 | zinc finger protein 469 |
| 1455273 | at | 1.12 | 0.031 | Zfp482 | zinc finger protein 482 |
| 1434088 | at | 1.01 | 0.829 | Zfp496 | zinc finger protein 496 |
| 1460572 | a_at | -1.05 | 0.733 | Zfp511 | zinc finger protein 511 |
| 1452689 | at | -1.01 | 0.922 | Zfp512 | zinc finger protein 512 |
| 1426471 | at | 1.33 | 0.003 | Zfp52 | zinc finger protein 52 |
| 1451332 | at | -1.04 | 0.619 | Zfp521 | zinc finger protein 521 |
| 1435669 | at | -1.07 | 0.284 | Zfp532 | zinc finger protein 532 |
| 1448920 | at | 1.04 | 0.657 | Zfp535 | zinc finger protein 535 |
| 1435337 | at | 1.20 | 0.068 | Zfp537 | zinc finger protein 537 |
| 1426564 | at | -1.22 | 0.004 | Zfp553 | zinc finger protein 553 |
| 1434128 | a_at | 1.13 | 0.198 | Zfp574 | zinc finger protein 574 |
| 1460620 | at | 1.03 | 0.666 | Zfp592 | zinc finger protein 592 |
| 1435556 | at | -1.12 | 0.013 | Zfp597 | zinc finger protein 597 |
| 1460691 | at | 1.21 | 0.050 | Zfp598 | zinc finger protein 598 |
| 1427104 | at | -1.66 | 0.004 | Zfp612 | zinc finger protein 612 |
| 1425030 | at | 1.21 | 0.000 | Zfp622 | zinc finger protein 622 |
| 1426625 | at | 1.21 | 0.004 | Zfp623 | zinc finger protein 623 |
| 1436537 | at | -1.67 | 0.000 | Zfp629 | zinc finger protein 629 |
| 1434689 | at | -1.01 | 0.826 | Zfp637 | zinc finger protein 637 |

Supplementary table S1

| | | | | | |
|---|---|---|---|---|---|
| 1424237 | at | -1.02 | 0.846 | Zfp639 | zinc finger protein 639 |
| 1424261 | at | 1.02 | 0.555 | Zfp672 | zinc finger protein 672 |
| 1417549 | at | -1.09 | 0.324 | Zfp68 | zinc finger protein 68 |
| 1453230 | at | -1.03 | 0.512 | Zfp74 | zinc finger protein 74 |
| 1439558 | at | -1.02 | 0.718 | Zfp75 | zinc finger protein 75 |
| 1456483 | at | 1.04 | 0.359 | Zfp9 | zinc finger protein 9 |
| 1429615 | at | 1.16 | 0.002 | Zfp91 | zinc finger protein 91 |
| 1424892 | at | 1.24 | 0.032 | Zfp95 | zinc finger protein 95 |
| 1451281 | at | 1.00 | 0.938 | Zfp96 | zinc finger protein 96 |
| 1449314 | at | -1.15 | 0.059 | Zfpm2 | zinc finger protein, multitype 2 |
| 1437542 | at | -1.24 | 0.006 | Zfpn1a2 | zinc finger protein, subfamily 1A, 2 (Helios) |
| 1436554 | at | -1.07 | 0.868 | Zfpn1a5 | zinc finger protein, subfamily 1A, 5 |
| 1458450 | at | -1.00 | 0.989 | Zfr | zinc finger RNA binding protein |
| 1447947 | at | 1.63 | 0.000 | Zfyve16 | zinc finger, FYVE domain containing 16 |
| 1451380 | at | 1.07 | 0.272 | Zfyve19 | zinc finger, FYVE domain containing 19 |
| 1424669 | at | -1.45 | 0.000 | Zfyve21 | zinc finger, FYVE domain containing 21 |
| 1440348 | at | -1.04 | 0.571 | Zfyve9 | zinc finger, FYVE domain containing 9 |
| 1424548 | at | -1.07 | 0.167 | Zgpat | zinc finger, CCCH-type with G patch domain |
| 1448875 | at | 1.06 | 0.341 | Zhx1 | zinc fingers and homeoboxes protein 1 |
| 1455210 | at | -1.04 | 0.690 | Zhx2 | zinc fingers and homeoboxes protein 2 |
| 1452977 | at | -1.12 | 0.203 | Zhx3 | zinc fingers and homeoboxes 3 |
| 1417536 | at | 1.25 | 0.000 | Zmat2 | zinc finger, matrin type 2 |
| 1427923 | at | 1.02 | 0.550 | Zmpste24 | zinc metalloproteinase, STE24 homolog (S. cerevisiae) |
| 1428052 | a_at | 1.07 | 0.380 | Zmym1 | zinc finger, MYM domain containing 1 |
| 1425531 | at | -1.09 | 0.029 | Znhit1 | zinc finger, HIT domain containing 1 |
| 1416651 | at | -1.18 | 0.225 | Znhit2 | zinc finger, HIT domain containing 2 |
| 1422517 | a_at | 1.05 | 0.552 | Znrd1 | zinc ribbon domain containing, 1 |
| 1420950 | at | -1.38 | 0.001 | Znrf1 | zinc and ring finger 1 |
| 1434016 | at | 1.15 | 0.138 | Znrf2 | zinc and ring finger 2 |
| 1445389 | at | 1.29 | 0.405 | Zp3r | zona pellucida 3 receptor |
| 1451189 | at | 1.06 | 0.257 | Zswim1 | zinc finger, SWIM domain containing 1 |
| 1434967 | at | -1.09 | 0.211 | Zswim6 | zinc finger, SWIM domain containing 6 |
| 1424891 | a_at | 1.06 | 0.423 | Zw10 | ZW10 homolog (Drosophila), centromere/kinetochore protein |
| 1423724 | at | 1.03 | 0.772 | Zwint | ZW10 interactor |
| 1417240 | at | 1.55 | 0.005 | Zyx | zyxin |
| 1453583 | at | 1.03 | 0.838 | Zzef1 | zinc finger, ZZ-type with EF hand domain 1 |
| 1434332 | at | 1.02 | 0.773 | Zzz3 | zinc finger, ZZ domain containing 3 |
| | | 1.04 | | | |
