## Supplemental tables S2-S3 for "A phenotypic screen identified KEAP1-kelch domain blockade as a mechanism to restore cardiac function in the setting of chronic severe hemodynamic stress"

**Table S2: Phenotypic screen hits.** A hit was defined as any compound with a score above the vehicle score variability.

| Compound | Score | P value | Score Rank | Compound target annotation | Mechanism of action |
|---|---|---|---|---|---|
| GSK1326341A | 1.71 | 1.8E-04 | 1 | Cathepsin B | Inhibitor |
| GSK929027A | 1.60 | 2.1E-04 | 2 | Glucocorticoid receptor (NR3C1) | Agonist |
| GW671021B | 1.44 | 2.6E-04 | 3 | Prostaglandin E receptor 3 (EP3) | Antagonist |
| GSK1580339A | 1.42 | 2.7E-04 | 4 | Solute carrier organic anion transporter family member 1B1 | Antagonist |
| GSK1436028A | 1.17 | 4.2E-04 | 5 | Mechanistic target of rapamycin kinase (MTOR) | Inhibitor |
| GSK2287329C | 1.16 | 4.3E-04 | 6 | TNNI3 interacting kinase (TNNI3K) | Inhibitor |
| GW297251A | 1.16 | 4.3E-04 | 7 | Protein kinase AMP-activated catalytic subunit α1 (AMPK) | Inhibitor |
| GW709135X | 1.16 | 4.3E-04 | 8 | Pan-peroxisome proliferator activated receptor | Agonist |
| SB-616234-B | 1.15 | 4.4E-04 | 9 | 5-hydroxytrptamine receptor 1B (5-HT1B) | Antagonist |
| GSK2366910A | 1.12 | 4.6E-04 | 10 | Kelch-like ECH associated protein 1 (KEAP1) | Inhibitor |
| GSK1645879A | 1.10 | 8.4E-04 | 11 | Checkpoint kinase 1 | Inhibitor |
| GW855266X | 1.09 | 5.0E-04 | 12 | Peroxisome proliferator activated receptor-γ | Agonist |
| GSK2702922A | 1.07 | 5.1E-04 | 13 | Protein phosphatase 1D (WIP-1) | Inhibitor |
| GSK1847058A | 1.06 | 5.2E-04 | 14 | Protein kinase Cα (PKCα) | Inhibitor |
| GSK2115363A | 1.06 | 5.3E-04 | 15 | p21 activated kinase 5 (PAK5) | Inhibitor |
| GI166507X | 1.05 | 5.4E-04 | 16 | Ca++ channel, L-type and T-type | Antagonist |
| GW786008A | 1.00 | 6.0E-04 | 17 | Leukotriene B4 receptor 2 | Antagonist |
| GSK1869056B | 1.00 | 6.0E-04 | 18 | Mitogen-activated protein kinase 3 (MAPK3) | Inhibitor |
| GW551106X | 1.00 | 6.1E-04 | 19 | Prostaglandin G/H synthase 2 (COX2) | Inhibitor |
| GSK1522082A | 0.99 | 6.1E-04 | 20 | Prostaglandin D2 receptor (CRTH2) | Antagonist |
| GSK1998042A | 0.97 | 6.5E-04 | 21 | Phosphatidylinositol 4,5 biphosphate 3-kinase catalytic subunit-β | Inhibitor |
| GSK1900952A | 0.97 | 6.5E-04 | 22 | Phosphatidylinositol 3-kinase regulatory subunit-α (PIK3R1) | Inhibitor |
| GSK1866180A | 0.96 | 6.6E-04 | 23 | Protein kinase C epsilon type (PRKCE) | Inhibitor |
| GSK1079865A | 0.95 | 6.7E-04 | 24 | Cathepsin D | Inhibitor |

| | | | | | |
|---|---|---|---|---|---|
| GSK1973832A | 0.95 | 6.7E-04 | 25 | Phosphatidylinositol 4,5 biphosphate 3-kinase catalytic subunit-δ | Inhibitor |
| GSK2250262A | 0.94 | 7.0E-04 | 26 | Oxysterols receptor LXR-alpha (NR1H3) | Agonist |
| GSK2616607A | 0.93 | 7.0E-04 | 27 | Nuclear receptor ROR-gamma (RORC) | Antagonist |
| GSK1642651A | 0.93 | 7.1E-04 | 28 | Calcitonin receptor-like receptor (CRLR) | Antagonist |
| SB-706991 | 0.93 | 7.1E-04 | 29 | Vascular endothelial growth factor receptor (sorafenib) | Inhibitor |
| GSK2831564A | 0.92 | 7.2E-04 | 30 | Nuclear receptor ROR-gamma (RORC) | Antagonist |
| SB-502424-A | 0.92 | 7.2E-04 | 31 | Nociceptin receptor (ORL1) | Antagonist |
| GSK2696294A | 0.89 | 7.9E-04 | 32 | Calcium channel, voltage-dependent, L-type, alpha 1C (CaV1.2) | Antagonist |
| GW664267X | 0.88 | 8.1E-04 | 33 | Nuclear receptor ROR-alpha | Antagonist |
| GSK1943275A | 0.87 | 8.2E-04 | 34 | Phosphatidylinositol 4,5 biphosphate 3-kinase catalytic subunit-β | Inhibitor |
| GSK1995018A | 0.85 | 8.7E-04 | 35 | Mechanistic target of rapamycin kinase (MTOR) | Inhibitor |
| SKF-60368 | 0.84 | 8.8E-04 | 36 | Calcium channel, voltage-dependent, L-type, alpha 1C (CaV1.2) | Antagonist |
| GSK2261818A | 0.84 | 8.9E-04 | 37 | Mitogen-activated protein kinase kinase-kinase 5 (ASK1) | Inhibitor |
| GSK1993774A | 0.84 | 9.0E-04 | 38 | Lysine-specific histone demethylase 1A (LSD-1) | Inhibitor |
| GSK1559549A | 0.83 | 9.1E-04 | 39 | NACHT, LRR and PYD domains-containing protein 3 (NLRP3) | Inhibitor |
| GSK2376536A | 0.83 | 9.1E-04 | 40 | Phosphatidylinositol 4,5 biphosphate 3-kinase catalytic subunit-β | Inhibitor |
| GSK344736A | 0.82 | 9.4E-04 | 41 | Gamma-aminobutyric acid B receptor (GABAB) | Modulator |
| GSK207933A | 0.80 | 9.8E-04 | 42 | TGF-beta receptor type-1 (ALK5) | Inhibitor |
| GSK2242446A | 0.79 | 1.0E-03 | 43 | Spleen tyrosine kinase (SYK) | Inhibitor |
| SKF-94212 | 0.76 | 1.1E-03 | 44 | Thyroid receptor-β (TR-β) | Agonist |
| GSK1322660A | 0.76 | 1.1E-03 | 45 | Gastric inhibitory polypeptide receptor (GIPR) | Antagonist |
| SKF-105570 | 0.75 | 1.1E-03 | 46 | cAMP-specific 3',5'-cyclic phosphodiesterase 4B | Inhibitor |
| GSK245351A | 0.74 | 1.2E-03 | 47 | Polo-like kinase 3 (PLK3) | Inhibitor |
| GSK2923214A | 0.74 | 1.2E-03 | 48 | Coactivator-associated arginine methyltransferase 1 (CARM1) | Inhibitor |
| GSK249849A | 0.74 | 1.2E-03 | 49 | Dual specificity tyrosine-phosphorylation-regulated kinase-2 (DYRK2) | Inhibitor |
| GSK350165A | 0.74 | 1.2E-03 | 50 | ATP citrate lyase (ACLY) | Inhibitor |
| GW837331X | 0.72 | 1.3E-03 | 51 | Polo-like kinase 1 (PLK1) | Inhibitor |
| GSK1418002B | 0.70 | 1.3E-03 | 52 | Checkpoint kinase 2 (CHK2) | Inhibitor |
| GSK2105314A | 0.69 | 1.3E-03 | 53 | Protein kinase C-θ type | Antagonist |
| GSK1304315A | 0.68 | 1.3E-03 | 54 | G-protein coupled estrogen receptor | Agonist |

| | | | | | |
|---|---|---|---|---|---|
| GSK200763A | 0.67 | 1.3E-03 | 55 | Vascular endothelial growth factor receptor 2 | Inhibitor |
| GSK2467955A | 0.66 | 1.3E-03 | 56 | 5'-AMP-activated protein kinase catalytic subunit α-2 | Activator |
| GSK2506758A | 0.64 | 1.3E-03 | 57 | Histone-lysine N-methyltransferase EHMT2 | Inhibitor |
| GSK2655742B | 0.64 | 1.3E-03 | 58 | Acetylcholine receptor subunit alpha | Antagonist |
| GSK994817A | 0.64 | 1.3E-03 | 59 | Insulin-like growth factor 1 receptor | Inhibitor |
| GW848687X | 0.64 | 1.3E-03 | 60 | Prostaglandin E2 receptor EP1 subtype | Antagonist |
| GSK1061445A | 0.64 | 1.3E-03 | 61 | Cathepsin D | Inhibitor |
| GSK2603613A | 0.63 | 1.3E-03 | 62 | Histone-lysine N-methyltransferase EHMT2 | Inhibitor |
| GSK2376537A | 0.63 | 1.3E-03 | 63 | Phosphatidylinositol 4,5-bisphosphate 3-kinase catalytic subunit-β | Inhibitor |
| GSK838592A | 0.63 | 1.3E-03 | 64 | Tissue plasminogen activator | Inhibitor |
| GW710648X | 0.63 | 1.3E-03 | 65 | Glucocorticoid receptor alpha | Agonist |
| GSK2827408A | 0.62 | 1.3E-03 | 66 | Proto-oncogene tyrosine-protein kinase receptor Ret | Inhibitor |
| GSK590780A | 0.62 | 1.3E-03 | 67 | Serine/threonine-protein kinase 4 | Inhibitor |
| GSK2660524A | 0.61 | 1.3E-03 | 68 | Histone-lysine N-methyltransferase EZH1 | Inhibitor |
| GSK1995046A | 0.61 | 1.3E-03 | 69 | Anaplastic Lymphoma Kinase | Inhibitor |
| SB-711845-Z | 0.61 | 1.3E-03 | 70 | HCV Nonstructural protein 5B | Inhibitor |
| GW381086A | 0.60 | 1.3E-03 | 71 | Serine/threonine-protein kinase pim-2 | Inhibitor |
| GSK570050A | 0.60 | 1.3E-03 | 72 | Glucocorticoid Receptor alpha | Agonist |
| GSK1343527A | 0.60 | 1.3E-03 | 73 | Catechol O-Methyltransferase | Inhibitor |
| GW842470X | 0.59 | 1.3E-03 | 74 | cAMP-specific 3',5'-cyclic phosphodiesterase 4 (pan) | Inhibitor |
| SB-423557-A | 0.59 | 1.3E-03 | 75 | Extracellular calcium-sensing receptor | Antagonist |
| GSK577732A | 0.59 | 1.3E-03 | 76 | Cathepsin Z | Inhibitor |
| GSK2434885A | 0.59 | 1.3E-03 | 77 | Leukotriene A-4 hydrolase | Antagonist |
| GSK183390A | 0.57 | 1.3E-03 | 78 | Peroxisome proliferator activated receptor α/γ | Agonist |
| GSK1973440A | 0.57 | 1.3E-03 | 79 | Lymphocyte-specific protein tyrosine kinase | Inhibitor |
| SB-342460-A | 0.57 | 1.3E-03 | 80 | Dual specificity protein phosphatase 6 | Inhibitor |
| SKF-9439-A | 0.56 | 1.3E-03 | 81 | 2'-5' oligoadenylate synthetase-dependent ribonuclease | Inhibitor |
| GSK2251187A | 0.56 | 1.3E-03 | 82 | Spleen tyrosine kinase, | Inhibitor |
| GSK2593467A | 0.55 | 1.3E-03 | 83 | ADP-ribosyl cyclase/cyclic ADP-ribose hydrolase 1 | Inhibitor |
| GSK1598306A | 0.55 | 1.3E-03 | 84 | PC4 and SFRS1-interacting protein | Antagonist |

| | | | | | |
|---|---|---|---|---|---|
| GSK2391846A | 0.55 | 1.3E-03 | 85 | Serine/threonine-protein kinase pak-1 | Inhibitor |
| SB-799360 | 0.55 | 1.3E-03 | 86 | Carnitine O-palmitoyl-transferase 1, muscle isoform | Activator |
| GSK2894856A | 0.54 | 1.3E-03 | 87 | Lysine-specific demethylase 5B | Inhibitor |
| GSK2645771A | 0.54 | 1.3E-03 | 88 | Fatty acid-binding protein, adipocyte | Inhibitor |
| GSK1160849A | 0.53 | 1.3E-03 | 89 | Secreted embryonic alkaline phosphatase | Inhibitor |
| GSK2661339A | 0.53 | 1.3E-03 | 90 | Lysine-specific demethylase 4D | Inhibitor |
| GW854991X | 0.53 | 1.3E-03 | 91 | Kelch-like ECH associated protein 1 (KEAP1) | Inhibitor |
| GSK2395671A | 0.53 | 1.3E-03 | 92 | Diacylglycerol O-acyltransferase 1 | Inhibitor |
| GSK1367131A | 0.53 | 1.3E-03 | 93 | Lymphocyte-specific protein tyrosine kinase | Inhibitor |
| GSK2132316B | 0.51 | 1.3E-03 | 94 | Ribosomal protein S6 kinase beta-1 | Inhibitor |
| GSK1271831A | 0.51 | 1.3E-03 | 95 | cAMP-specific 3',5'-cyclic phosphodiesterase 4B | Inhibitor |
| SB-737542-V | 0.51 | 1.3E-03 | 96 | Mas-related G-protein coupled receptor member X1 | Agonist |
| SB-324691 | 0.50 | 1.3E-03 | 97 | Probable G-protein coupled receptor 101 | Agonist |
| GW629712X | 0.50 | 1.3E-03 | 98 | Integrin alpha 4 beta 1 | Inhibitor |
| GSK2657108A | 0.50 | 1.3E-03 | 99 | Lysine-specific demethylase 4D | Inhibitor |
| GSK2329198A | 0.49 | 1.3E-03 | 100 | Sodium channel protein type 5 subunit alpha (Nav1.5) | Inhibitor |

### Table S3: Initial TAC screening study data

| | Sham | TAC | | | | Sham2 | TAC2 | | | | |
|---|---|---|---|---|---|---|---|---|---|---|---|
| Primary Endpoints | Veh | Veh | SKF-92837 (0.14 mg/kg) | SKF-92837 (0.42 mg/kg) | Thyroid hormone (T3) | Veh | Veh | Bardox methyl (5.0 mg-QD) | GSK3175696A (40 mg-BID) | GSK3161593 (50 mg-QD) | GSK3161593 (50 mg-BID) |
| Ejection fraction (%) | 68 (2.3) | 34 (5.3) | 39 (6.4) | 44 (5.0) | 36 (5.2) | 67 (1.4) | 42 (2.5) | 46 (3.2) | 47 (4.0) | 49 (2.6) | 56 (2.7)* |
| LV weight /body weight | 3.3 (0.1) | 6.5 (0.4) | 5.8 (0.5) | 5.5 (0.3) | 5.4 (0.5) | 2.7 (0.1) | 4.8 (0.2) | 5.1 (0.4) | 4.6 (0.3) | 4.5 (0.1) | 4.2 (0.2) |
| LVEDP (mmHg) | 2.9 (1.1) | 21.1 (1.7) | 18.6 (2.2) | 15.3 (2.2) | 14.9 (3.9) | 0.5 (2.1) | 10 (1.9) | 7.9 (1.7) | 7.2 (1.3) | 7.8 (2.3) | 4.5 (1.3) |
| heart rate (bpm) | 552 (8.6) | 500 (10.7) | 525 (12.7) | 543 (9.0)* | 577 (19.3)* | 439 (15.0) | 497 (17.0) | 478 (27.0) | 462 (9.0) | 468 (11.0) | 441 (10.0)* |

| | |
|---|---|
| SKF-92837 | Thyroid receptor-β agonist, administered in chow day of TAC surgery |
| Thyroid hormone (T3) | Thyroid receptor(s) agonist (native), administered in chow day of TAC surgery |
| Bardoxolone methyl # | Keap1-BTB domain modulator, administered 5mg/kg by oral gavage, initiated 1-week post-TAC |
| GSK3175696 # | Keap1-kelch domain blocker, administered 40mg/kg subcutaneously twice daily, initiated 1-week post-TAC |
| GSK3161593 # | Keap1-kelch domain blocker, administered 50mg/kg, subcutanously, once (QD) or twice (BID) daily, initiated 1-week post-TAC |

All values mean (+/- standard error of the mean)

**p < 0.05 vs vehicle, Oneway ANOVA, Dunnett's multiple comparison test vs vehicle*

*# Compounds dissolved in 20% hydroxypropyl-β-cyclodextrin (w/v) with 2% dimethyl sulfoxide, pH 8.5.*
